# Large-scale GWAS in sorghum reveals common genetic control of grain size among cereals

**DOI:** 10.1101/710459

**Authors:** Yongfu Tao, Xianrong Zhao, Xuemin Wang, Adrian Hathorn, Colleen Hunt, Alan W. Cruickshank, Erik J. van Oosterom, Ian D. Godwin, Emma S. Mace, David R. Jordan

**Author notes:** Corresponding authors: Yongfu Tao, Emma Mace, David Jordan.

## Abstract

- Grain size is a key yield component of cereal crops and a major quality attribute. It is determined by a genotype’s genetic potential and its capacity to fill the grains.
- This study aims to dissect the genetic architecture of grain size in sorghum via an integrated genome wide association study (GWAS) using a diversity panel of 837 individuals and a BC-NAM population of 1,421 individuals.
- In order to isolate genetic effects associated with grain size, rather than the genotype’s capacity to fill grain, a field treatment of removing half of the panicle during flowering was imposed. Extensive variation in grain size with high heritability was observed in both populations across 5 field trials. Subsequent GWAS analyses uncovered 92 grain size QTL, which were significantly enriched for orthologues of known grain size genes in rice and maize. Significant overlap between the 92 QTL and grain size QTL in rice and maize was also found, supporting common genetic control of this trait among cereals. Further analysis found grain size genes with opposite effect on grain number were less likely to overlap with the grain size QTL from this study, indicating the treatment facilitated identification of genetic regions related to the genetic potential of grain size rather than the capacity to fill the grain.
- These results enhance understanding of the genetic architecture of grain size in cereal, and pave the way for exploration of underlying molecular mechanisms in cereal crops and manipulation of this trait in breeding practices.

## Introduction

Cereal crops, including maize, rice, wheat, barley, and sorghum, supply more than 75% of the calories consumed by humans (Sands *et al*., 2009). They are critical to address global food security, which is under threat of population expansion and climate change. Among cereals, sorghum is an important source of food, fibre, feed, and biofuel, and provides staple food for over 500 million people in the semi-arid tropics of Africa and Asia. Cereal crops share a close ancestry, and strong gene co-linearity among them supports comparative genomics approaches for exploring the genetic bases of agronomical traits at a cross-species level (Bolot *et al*., 2009). Grain size is a key yield component of grain yield in cereal crops, crops and is a major quality attribute affecting planting, harvesting, and processing activities (Tao *et al*., 2017). A better understanding of the genetic basis of grain size, including underlying molecular mechanisms, will provide new targets for improving yield and grain quality in cereal breeding.

Grain size in cereals is a function of both the potential maximum grain size and the capacity of the plant to fill the grain. The weight of individual grains is determined by the rate and duration of grain filling. In sorghum, the grain filling rate is highly correlated with the ovary volume at anthesis, which in turn is associated with the size of the meristematic dome during early floret development (Yang *et al*., 2009). While the potential maximum grain size is largely genetically pre-determined, the capacity to fill the grain is strongly affected by assimilate availability (Gambín & Borrás, 2010). Grains are the key sink for carbon demand post-anthesis, and the amount of available assimilate per grain is determined by grain number and assimilate supply, which are associated with a range of genetic and environmental factors (Gambín & Borrás, 2010; Wardlaw, 1990). Negative correlations between grain number and grain size are commonly observed in cereal crops (Acreche & Slafer, 2006; Peltonen-Sainio *et al.*, 2007; Sadras, 2007; Tao *et al.*, 2018). In sorghum, grain number reduction at early stages of flowering has been observed to result in a larger grain weight (Sharma *et al.*, 2002). This indicates that demand for carbohydrate to fill grains exceeds the available supply.

In cereals, the genetic architecture of grain size is complex and involves numerous genes. In rice and maize, around 110 genes affecting grain size have been cloned (Tao *et al*., 2017). Given the conservation of gene function among closely related cereal species, this presents opportunities for comparative genomics studies to investigate the genetics of grain size in other cereal crops using knowledge gained from gene cloning studies in maize and rice. The sorghum QTL atlas of Mace *et al.* (2018) details around 100 grain weight QTL in sorghum that have been identified through linkage analysis in 18 studies (Paterson *et al.*, 1995; Tuinstra *et al.*, 1997; Rami *et al.*, 1998; Brown *et al.*, 2006; Feltus *et al.*, 2006; Murray *et al.*, 2008; Srinivas *et al.*, 2009; Phuong *et al*., 2013; Rajkumar *et al*., 2013; Reddy *et al*., 2013; Han *et al*., 2015; Mocoeur *et al*., 2015; Shehzad & Okuno, 2015; Gelli et al., 2016; Spagnolli et al., 2016; Bai et al., 2017; Boyles et al., 2017;Tao *et al.*, 2018). A further 12 markers associated with grain weight have been identified in three GWAS studies (Upadhyaya *et al.*, 2012; Zhang *et al.*, 2015; Boyles *et al.*, 2016). Given the number of genomic regions identified using standard genetic linkage mapping approaches, the limited number of marker-trait associations identified in the GWAS studies is likely due to the small population sizes employed in these studies (n = 242 – 390). All of the previous genetic studies on grain size in sorghum have focused on the dissection of the genetic basis of grain size under natural conditions, which also include environmental impacts and Genotype × Environment (G×E) interactions. However, minimisation of variation in grain-filling capacity could limit the environment effects on variation in grain size, and thereby facilitate identification of genetic factors underlying variation of this trait.

This study aims to 1) investigate the genetic architecture of grain size in sorghum by conducting an integrated GWAS in two large populations, a diversity panel of 837 individuals and a backcross nested association mapping population (BC-NAM) population of 1421 individuals, and 2) compare genetic loci affecting grain size identified in this study to those reported in rice and maize. To minimize variation in grain size caused by assimilate variability, a field treatment of removing half of the panicle during flowering time was imposed in this study in order to maximise assimilate availability for the remaining seeds during grain filling.

## Materials and methods

### Plant material

Two populations, together comprising over 2000 individuals, were used in this study: a diversity panel (DP, n=837) (Table S1) and a BC-NAM consisting of 30 interrelated families (n=1421) (Figure S1A; Table S2). The diversity panel has around 225 genotypes in common with the US sorghum association panel. It consists predominantly of lines developed by the sorghum conversion program conducted by Texas Agricultural Experiment Station, which took diverse sorghum lines from the world collection and converted tall, late, or photoperiod sensitive sorghums from the tropics into short, early, photoperiod insensitive types that could be used by breeders in temperate regions (Rosenow *et al*., 1997). The program involved repeated backcrossing to the exotic line combined with selection for height and maturity in temperate environments. The resulting material has been reported to contain >4% genome introgression from the temperate donor with the remainder from the exotic parent and has a much narrower range of height and maturity than the original exotic lines (Thurber *et al.*, 2013).

The BC-NAM population was previously developed by the Department of Agriculture and Fisheries (DAF), Queensland, Australia and described in (Jordan *et al.*, 2011). In summary, each BC-NAM family was produced by crossing a single elite parent (R931945-2-2 or R986087-2-4-1) with a diverse, exotic line (the non-recurrent parent) and backcrossing the resulting F_1_ to the elite parent to produce a large BC_1_F_1_ population. Variable numbers of plants were selected from each BC_1_F_1_ family and using single-seed descent >5 generations of self-pollination were generated to produce BC_1_F_6_ and beyond. During generation advance, selection was imposed for height and maturity of the recurrent parental type. The 30 BC-NAM families used in the current study are listed in Table S2. In total 27 unique non-recurrent parental lines and 2 elite recurrent parental lines were used for population development. These included 10 diverse lines from advanced breeding programs and 17 landraces, 7 of which were converted to temperate adaptation through the Sorghum Conversion Program (Stephens *et al.*, 1967). Fourteen of the 27 non-recurrent parental lines were also included in the diversity panel (Figure S1B).

### Field Trials and phenotypic evaluation

A total of 5 field trials were planted at Hermitage Research Facility (HER), Warwick, Queensland, Australia (28°12ʹS, 152°5ʹE, 470 m above sea level) and Gatton Research Facility (GAT), Gatton, Queensland, Australia (27°33ʹS, 152°20ʹE, 94 m above sea level) during the Australian summer cropping seasons of 2014/15 and 2015/16. Two trials of the diversity panel were grown in the 2015/16 season at Hermitage (DPHER16) and Gatton (DPGAT16). The BC-NAM were planted in three trials at Hermitage in 2014/15 (NAMHER15), and Gatton in 2014/15 (NAMGAT15) and 2015/16 (NAMGAT16). All trials were planted between November and February, using a row column design with partial replication. Each plot consisted of two 6 m rows with a row spacing of 0.75 m. Differences in plant height within a plot was minimal. Different numbers of genotypes were grown in each trial due to seed availability (Table S3). Standard agronomic practices and pest-control practices were applied.

A treatment of removing half of each of two panicles in each plot was imposed when each panicle commenced flowering, before significant grain development had occurred (Figure 1A). Once physiological maturity was reached, the remaining half panicle (hereafter, referred as half head) was hand harvested and threshed using a mechanical threshing machine. In DPGAT16, a full head panicle in each plot was also harvested and threshed for comparison with the half head panicles from the same plot. An aspirator was used to remove any debris from each sample before measurements were taken. Although sorghum grain is typically tending toward spherical, considerable phenotypic variation in length, width, and thinkness does exist. Therefore, grain size parameters, including thousand kernel weight (TKW), and the length, width, thickness, and volume of grains were measured using Seedcount SC5000 (Next Instruments, Condell Park, NSW, Australia) and a digital balance.

**Figure 1.**
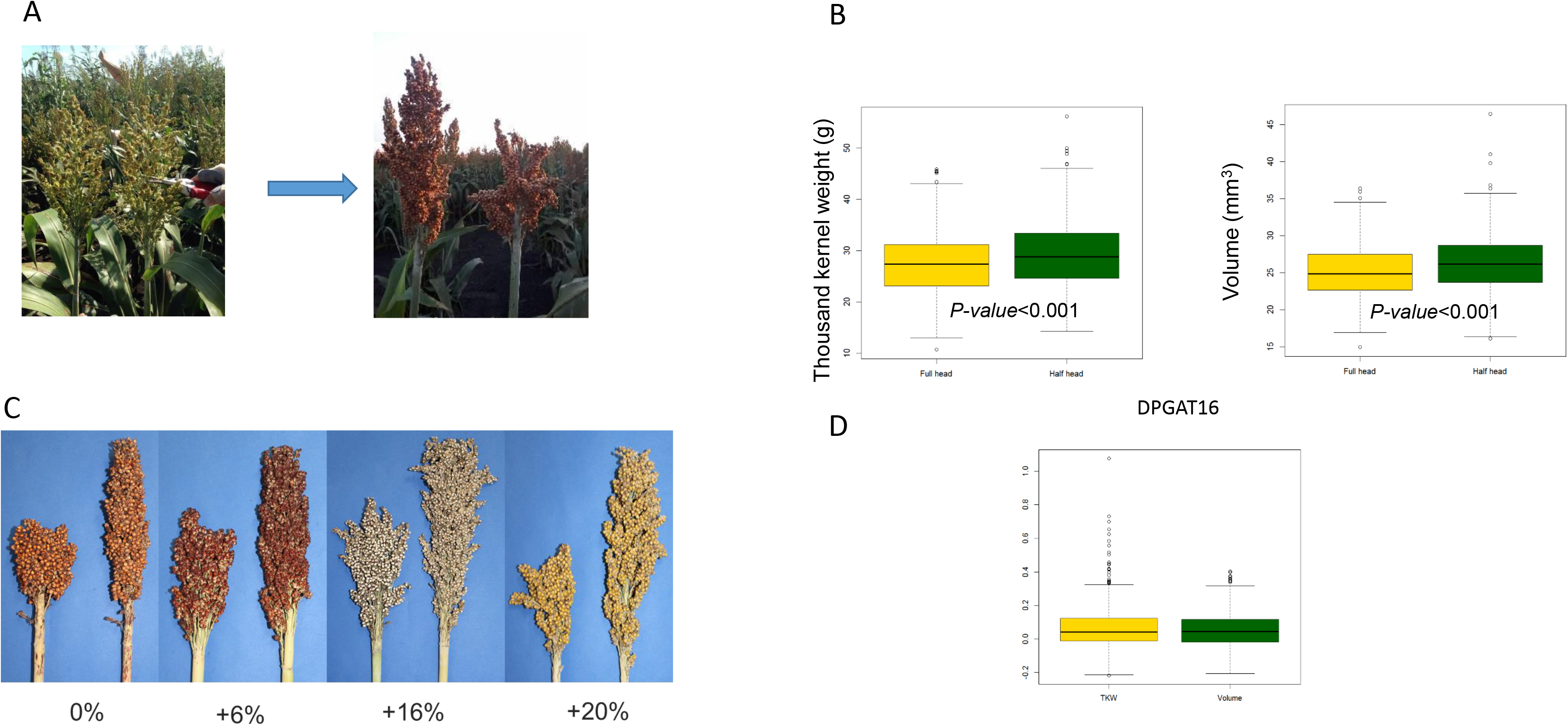
Field treatment and its effect on grain size. A) Field treatment of removing half of panicle imposed during flowering time. B) Significant increase of TKW and volume in half heads compared to full heads in DPGAT16. C) Varying increase percentage in TKW of half heads observed across genotypes. D) box-plots show variations of increase in TKW and volume in half heads across genotypes in DPGAT16.

### Statistical analysis

Due to the large number of genotypes included in this study, partial replication was used in all trial designs (Cullis *et al.*, 2006), with 30 % of the genotypes replicated two or more times and the remaining 70 % represented by single plots. The total number of plots in each trial ranged from 880 to 1521, with the total number of genotypes planted in each trial ranging from 658 to 1164 (Table S3). A customized design was used to minimise spatial error effects within each trial. The concurrence of genotypes and populations across the two seasons allowed the DP and BC-NAM trials to be analysed as two multi-environment trials (METs) for each of the five measured grain size parameters, comprising of two trials for the DP and three for the BC-NAM. Each MET was analysed by fitting a linear mixed model using the package ASReml (Butler *et al*., 2009) and the R statistical software. Each model consisted of a fixed effect for the targeted trait at each trial, random effects for genotype within trial, and spatial error for each individual trial (Smith *et al*., 2001). The G×E interaction was analysed by fitting a third-order factor analytic structure to the trial × genotype interaction, in this case the structure also modelled the 3-way interaction between site, trait, and genotype. The analysis resulted in genetic variances for each trial and loading values representing factor analytic loadings.

Generalised repeatability estimates were calculated for grain size parameters using the method proposed by Cullis *et al*. (2006). High correlations were observed among grain size parameters, and hence a principal components analysis was conducted to further investigate the interrelationships among TKW and grain length, width, and thickness (Figure S3 and S4). Because it is possible that a constructed trait such as a principal component may explain the data more effectively, the main principal components were used as additional traits in the association analysis.

### Genotyping and imputation

The 2 populations were genotyped using medium to high density genome-wide SNPs provided by Diversity Arrays Technology Pty Ltd (www.diversityarrays.com). DNA was extracted from bulked young leaves of five plants from a plot of each genotype in the 2 populations using a previously described CTAB method (Doyle, 1987). The DNA samples were then genotyped following an integrated DArT and genotyping-by-sequencing (GBS) methodology, which involves complexity reduction of the genomic DNA to remove repetitive sequences using methylation sensitive restriction enzymes prior to sequencing on Next Generation sequencing platforms. The sequence data generated were then aligned to version v3.1.1 of the sorghum reference genome sequence (McCormick *et al*., 2018) to identify SNPs (Single Nucleotide Polymorphisms).

In total, 111,089 SNPs were identified in the diversity panel and 31,478 SNPs in the BC-NAM population. The overall proportion of missing data reported in the raw genotypic data sets was approximately 10% and 5% for the diversity panel and BC-NAM respectively. Individual SNP markers with >50% missing data were removed from further analysis and the remaining missing values were phased and imputed using Beagle v4.1 (Browning & Browning, 2016). An average imputation accuracy of 96% was achieved across both populations.

### Population statistics, GWAS analysis and QTL identification

Pairwise linkage disequilibrium (LD) (r^2^) was calculated using PopLDdecay (Zhang *et al.*, 2018) for the diversity (Figure S4A) and the BC-NAM (Figure S1C). The population structure in the diversity panel was analysed used LEA in R (François, 2016). The structure analysis identified five groups that corresponded to 4 racial groups (kafir, caudatum, guinea, durras of Asian origin, durras of East African origin) (Figure S4B) based on previous racial information and geographical origin for a subset of lines. Each individual line with ≥95 % of genetic identity from one of the five identified groups was designated as a representative of the corresponding racial group (Figure S4C, Table S4). The remaining lines were considered as admixtures of multiple groups.

For the GWAS analysis, imputed SNP data sets for both populations were filtered for minor allele frequency (MAF) >0.005. The final number of SNPs used for GWAS was 56,412 for the diversity panel (Figure S5) and 26,926 for the BC-NAM population (Figure S6). For the diversity panel, a principal component analysis (PCA) was conducted using a pruned SNP data set to control for population structure. Marker pruning was conducted using PLINK (Purcell *et al.*, 2007) with a sliding window of 100 SNPs, a step size of 50 SNPs and *r*^2^ of 0.5. For the BC-NAM population, pedigree was used to control population structure. GWAS was performed using FarmCPU (Liu *et al.*, 2016) for both populations for the five measured grain size parameters and derived PCs. Thresholds for defining significant and suggestive associations were set using the Bonferroni correction based on the number of independent tests, calculated using the GEC software package (Li *et al.*, 2012). Marker-trait associations were then identified using a two-step process. Firstly, SNPs were identified that were significantly associated with grain size parameters and derived PCs in both populations separately (p<1.45E-06 for the diversity panel; p<5.83E-06 for the BC-NAM). Secondly, SNPs were identified that were only suggestively associated with grain size parameters and derived PCs in one population (P< 2.91E-05 for the diversity panel, 1.17E-04 for the BC-NAM), but that had either suggestive or significant support in the second population and were also within a specified distance of the original SNP (<1cM in the diversity panel and <2cM in the BC-NAM population, calculated from the sorghum genetic linkage consensus map; Mace *et al.*, 2009). Identified marker-trait associations were further clustered into QTL regions based on the predicted genetic positions of SNPs from the sorghum consensus map across traits within each population (Mace *et al.*, 2018). Finally QTL in common between the two populations were combined based on a 2cM window. The effects of identified QTL on all grain size parameters in each population were assessed using a linear model, with population structure accounted for by PCs for the diversity panel and by pedigrees in the BC-NAM population.

A candidate region based approach was used to further investigate associations between population specific QTL and grain size parameters in the alternative population. SNPs within a 0.5 cM window of a population specific QTL were extracted from the alternative population. To examine associations between SNPs in the candidate regions and the grain size parameters, GLM in TASSEL 5.0 was run, using either PCA in the diversity panel or pedigree in the BC-NAM population as a covariate to control for population structure (Bradbury *et al.*, 2007). The Bonferroni correction (0.05/number of markers) was used to control for false positives.

Using the BC-NAM population, allele effects of the exotic parent were estimated compared to the recurrent parent. At each QTL, the numbers of exotic alleles with larger and smaller values than the recurrent parent were determined, and a test conducted to assess if the numbers of positive and negative alleles relative to the recurrent parent varied. Families with <20 individuals were excluded from this analysis.

### Collection of previously reported grain weight QTL

Eighty-five grain weight QTL were collated from previously published studies (Table S5). The locations of these QTL were projected onto both the sorghum reference genome v3.1.1 and the sorghum consensus map following methods described by Mace et al. (2009, 2019). To be conservative, QTL with confidence interval > 20 cM were excluded during this step.

### Candidate gene analysis

Candidate genes for grain size in sorghum were identified using the methods described by Tao *et al. (*2017). Firstly, genes functionally determined to control grain size in rice, maize, and sorghum were collated through a literature search (Table S6). Subsequently, corresponding sorghum orthologous of grain size genes in rice and maize were identified using a combination of syntenic and bidirectional best hit (BBH) approaches. Syntenic gene sets among rice, maize and sorghum were downloaded from PGDD (http://chibba.agtec.uga.edu/duplication/) to identify sytenic orthologues of known grain size genes from rice and maize. Local blast was performed to identify best BLAST hits of pairs of genes from two genomes, using BLASTP.

## Results

### Minimizing environmental impact on grain size through the removal of half of the panicle during flowering time

In the DPGAT16 trial, a significant increase in average grain size of 8.54% was observed in half heads compared to full heads (Figure 1B). The magnitude of this increase varied significantly among genotypes, indicating the presence of genotypic variation in the degree of source limitation for grain filling in the full head (Figure 1C and 1D). Nonetheless, TKW of half-head and full-head treatments were highly correlated (r^2^~0.70) (Figure S7). Since the greater average TKW of the half head treatment compared with the full head treatment indicated that this treatment was effective in removing some of the variation in grain size due to restrictions in grain-filling capacity, data collected from plants with half head phenotypes were used in this study.

### Phenotypic variation of grain size

Substantial variation in TKW and the volume, length, width, and thickness of grains was observed in half head samples across the five trials and two populations (Tables 1 and 2, Figure 2). The diversity panel exhibited much broader variation in these grain size parameters than the BC-NAM population, reflecting its larger genetic diversity. For example, in the trials conducted at Gatton in 2016, the range in TKW was 6.5-58.4g in the diversity panel, compared to 16.9-44.5g in the BC-NAM population. All parameters had low levels of G×E interaction and were highly correlated across trials in both the diversity panel (r>0.82, average r=0.85) and the BC-NAM population (r>0.79, average r=0.89) (Tables 1 and 2). Hence, the combined cross-trial BLUPs were used for subsequent analyses of these parameters. The high cross-trial correlations of the individual grain size parameters were consistent with their medium to high heritability (Table 1 and 2).

**Table 1.** Summary of phenotypic data collected from half head samples across trials in the diversity panel

| Trait | Trial | Trait Mean | Range | Hsq | Correlation between Sites |
| --- | --- | --- | --- | --- | --- |
| TKW (g) | DPGAT16 | 28.8 | 6.5 - 58.4 | 0.79 | 0.87 |
| TKW (g) | DPHER16 | 28.7 | 6.7 - 56.8 | 0.80 |  |
| Volume (mm<sup>3</sup>) | DPGAT16 | 26.2 | 8.4 - 55.0 | 0.67 | 0.85 |
| Volume (mm<sup>3</sup>) | DPHER16 | 26.1 | 9.9 - 55.6 | 0.70 |  |
| Length (mm) | DPGAT16 | 4.41 | 2.93 - 5.85 | 0.68 | 0.88 |
| Length (mm) | DPHER16 | 4.38 | 3.07 - 5.88 | 0.70 |  |
| Width (mm) | DPGAT16 | 3.50 | 2.29 - 4.74 | 0.65 | 0.84 |
| Width (mm) | DPHER16 | 3.46 | 2.34 - 4.56 | 0.67 |  |
| Thickness (mm) | DPGAT16 | 3.22 | 2.4 - 4.05 | 0.67 | 0.82 |
| Thickness (mm) | DPHER16 | 3.25 | 2.64 - 3.96 | 0.75 |  |

**Table 2.** Summary of the phenotypic data collected from half head samples across trials in the BC-NAM population

| Trait | Trial | Mean | Range | Hsq | Correlations across trials |  |  |
| --- | --- | --- | --- | --- | --- | --- | --- |
|  |  |  |  |  | NAMGAT15 | NAMGAT16 | NAMHER15 |
| TKW (g) | NAMGAT15 | 28.2 | 15.6 - 41.2 | 0.77 | 1 | 0.89 | 0.87 |
| TKW (g) | NAMGAT16 | 29.9 | 16.9 - 44.5 | 0.53 | 0.89 | 1 | 0.98 |
| TKW (g) | NAMHER15 | 30.8 | 13.9 - 45.6 | 0.66 | 0.87 | 0.98 | 1 |
| Volume (mm<sup>3</sup>) | NAMGAT15 | 25.5 | 14.6 - 40.5 | 0.36 | 1 | 0.91 | 0.89 |
| Volume (mm<sup>3</sup>) | NAMGAT16 | 25.3 | 15.0 - 38.7 | 0.37 | 0.91 | 1 | 0.81 |
| Volume (mm<sup>3</sup>) | NAMHER15 | 25.7 | 15.4 - 36.4 | 0.41 | 0.89 | 0.81 | 1 |
| Length (mm) | NAMGAT15 | 4.18 | 3.20 - 5.29 | 0.41 | 1 | 0.92 | 0.91 |
| Length (mm) | NAMGAT16 | 4.17 | 3.29 - 5.31 | 0.45 | 0.92 | 1 | 0.85 |
| Length (mm) | NAMHER15 | 4.18 | 3.31 - 5.07 | 0.40 | 0.91 | 0.85 | 1 |
| Width (mm) | NAMGAT15 | 3.47 | 2.58 - 4.32 | 0.29 | 1 | 0.91 | 0.87 |
| Width (mm) | NAMGAT16 | 3.47 | 2.65 - 4.26 | 0.39 | 0.91 | 1 | 0.79 |
| Width (mm) | NAMHER15 | 3.48 | 2.76 - 4.15 | 0.29 | 0.87 | 0.79 | 1 |
| Thickness (mm) | NAMGAT15 | 3.32 | 2.71 - 3.72 | 0.54 | 1 | 0.93 | 0.89 |
| Thickness (mm) | NAMGAT16 | 3.31 | 2.85 - 3.69 | 0.41 | 0.93 | 1 | 0.86 |
| Thickness (mm) | NAMHER15 | 3.35 | 2.9 - 3.78 | 0.59 | 0.89 | 0.86 | 1 |

**Figure 2:**
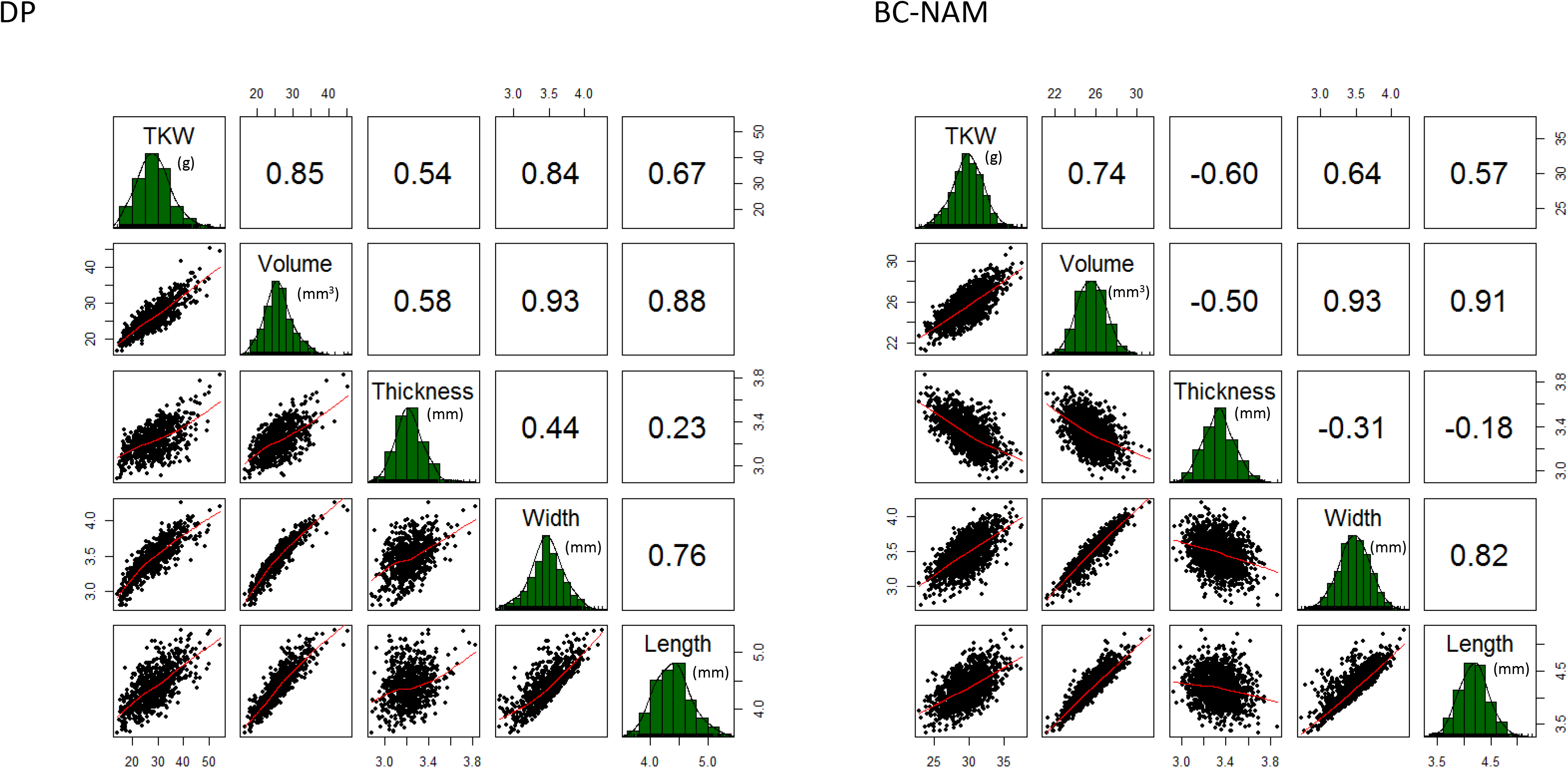
Phenotypic distribution and correlations of 5 grain size parameters across the diversity panel and BC-NAM population. The plots on the diagonal show the phenotypic distribution of each trait. The values above the diagonal are pairwise correlation coefficients between traits, and the plots below the diagonal are scatter plots of compared traits.

All five grain size parameters displayed near normal distributions in both populations (Figure 2), which in combination with the medium to high heritability and low G×E observed, indicated that variation in the parameters was controlled by multiple genetic loci. High correlations among the five grain size parameters were also observed in both populations, implying a high degree of shared genetic control among them (Figure 2). The principal components analysis determined that 3 PCs accounted for the majority (>85%) of the variation in the grain size parameters observed across multiple trials in both populations (Figure S2 and S3). The direction of the correlations between traits was positive in all contrasts with the exception of grain thickness, which displayed negative correlations with the other four grain size parameters in the BC-NAM population (r=−0.39), as opposed to positive correlations (r=+0.44) in the diversity panel (Figure 2).

To investigate possible correlations between racial groups and grain size parameters in sorghum, structure analysis of the diversity panel was conducted, which revealed 5 groups that corresponded to different racial groups of sorghum (Figure S4B, Table S4). All grain size parameters showed significant differences among these racial groups (ANOVA, *p*-value <0.05), with representatives of the caudatum and guinea racial groups having the largest and heaviest grains and east-African durras the smallest and lightest (Table S7).

### Identification of genetic loci associated with variation of grain size

In the half head panicles from the diversity panel, GWAS identified 84 marker-trait associations, with 71 SNPs significantly associated with between 1-4 of the grain size parameters (Figure 3, Figure S8, and Table S8). LD in the diversity panel decayed to the background level at ~ 200 kb, equivalent to 1 cM (Figure S4A). Thus, a 1cM window was used to cluster these 71 SNPs into 54 QTL. The majority (>80%) of these 54 QTL were significantly associated with multiple grain size parameters (Table S9). However, phenotypic variation explained by individual QTL was generally small and the majority of QTL explained < 5% of phenotypic variation each. In the BC-NAM population, 131 significant marker-trait associations were identified with 106 SNPs associated with between 1 – 4 grain size parameters (Figure 4, Figure S9, and Table S10). Because the extent of LD was greater in the BC-NAM than in the diversity panel (Figure S1C), a 2 cM window was used to cluster these associated SNPs into 66 QTL regions. Similar to observations for the diversity panel, the vast majority (~85%) of the QTL were significantly associated with multiple grain size parameters and phenotypic variation explained by individual QTL was generally small, with the majority of QTL explaining < 5% of the phenotypic variation of a given grain size parameter (Table S11). Within the BC-NAM, it was possible to observe the distribution of allelic effects of the exotic parents compared to the recurrent parent (Figure 5). In general, alleles were observed that were both larger and smaller than the elite parent, which was supportive of the presence of multiple alleles at the majority of QTL.

**Figure 3:**
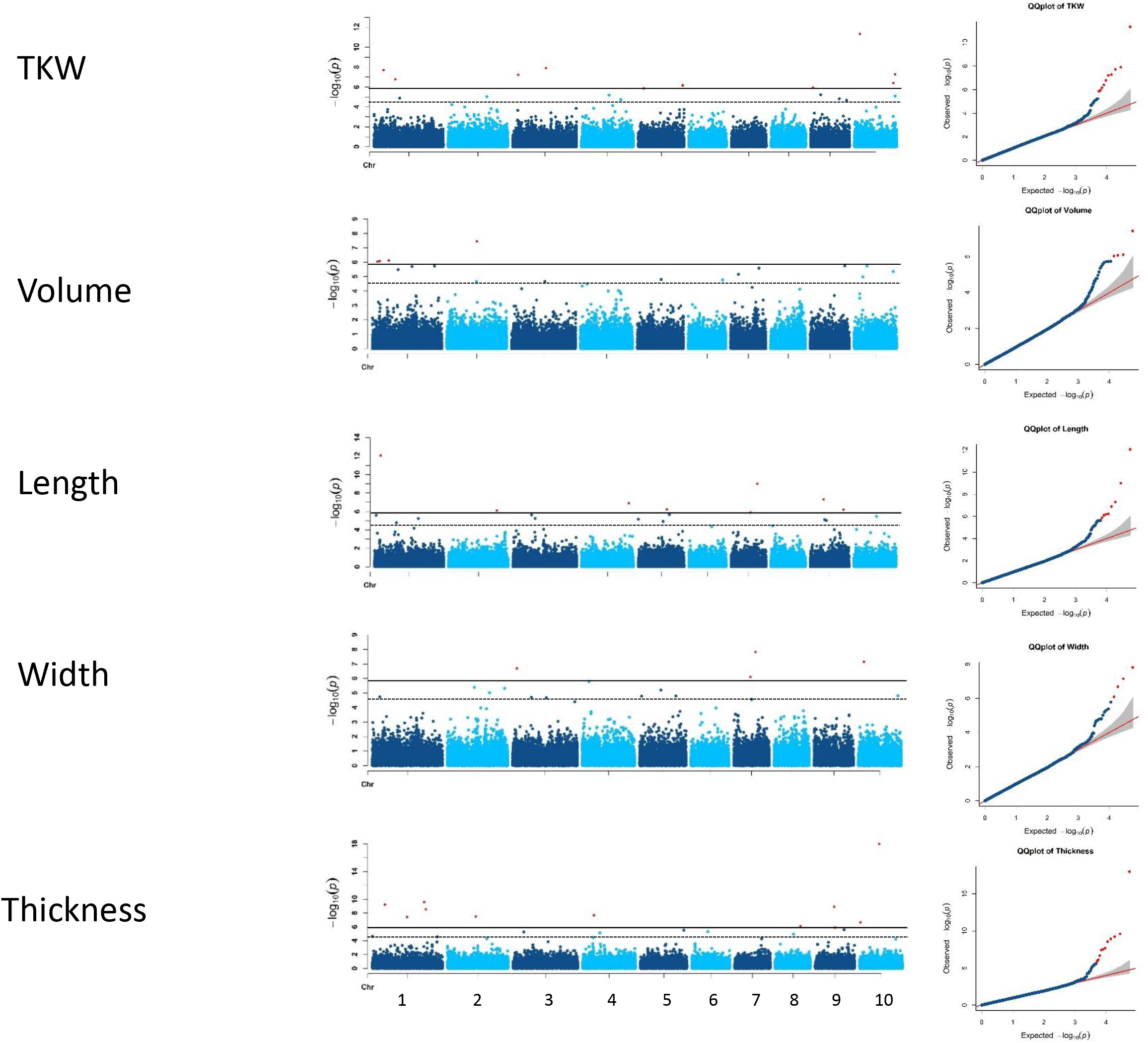
GWAS of 5 grain size parameters in the diversity panel. GWAS results were displayed in Manhattan plots and Q-Q plots. Significant threshold (solid black line) and suggest threshold (dash black line) were 1.45E-06 and 2.91E-05, respectively, according to GEC test.

**Figure 4:**
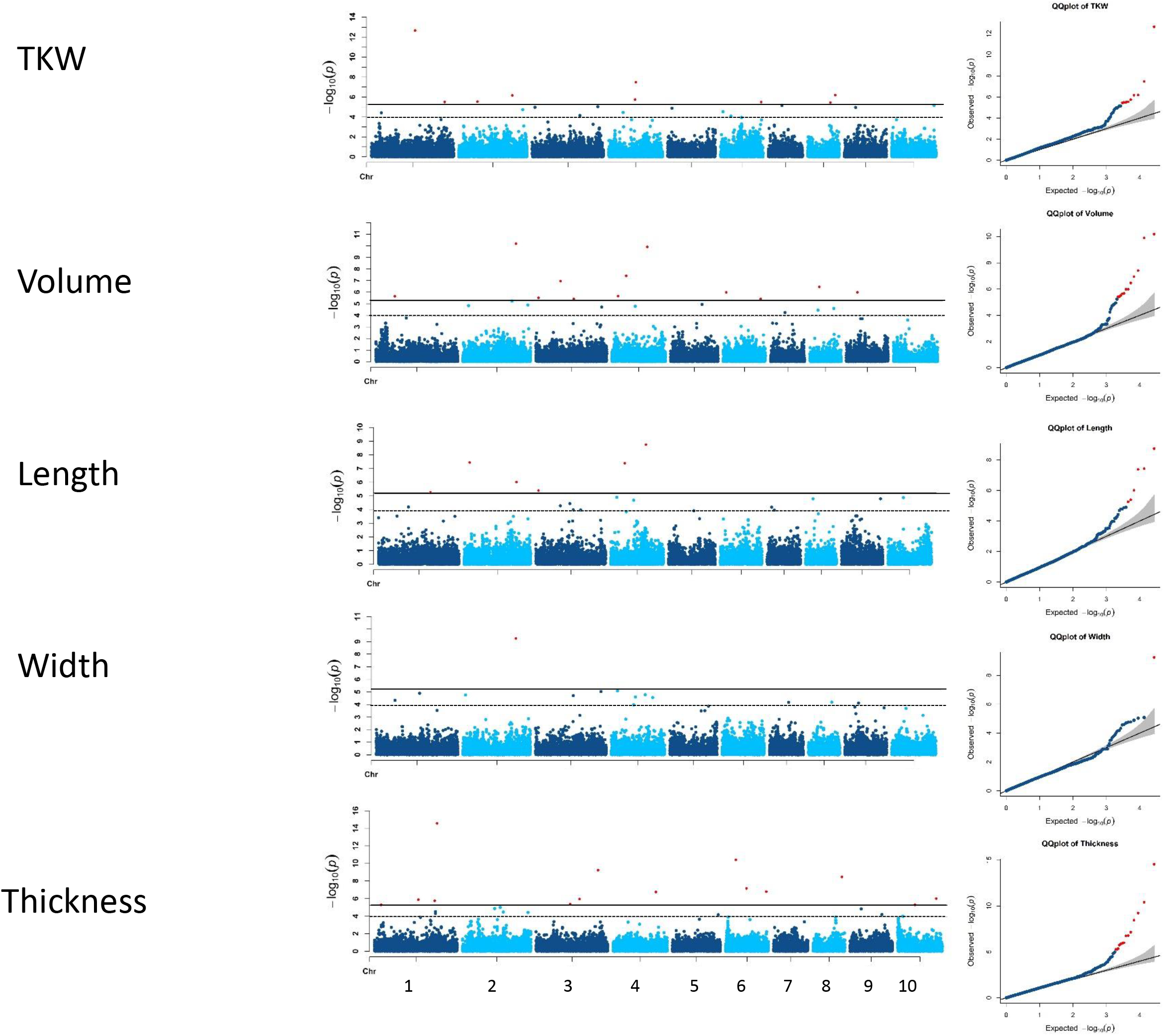
GWAS of 5 grain size parameters in the BC-NAM population. GWAS results were displayed in Manhattan plots and Q-Q plots. Significant threshold (solid black line) and suggest threshold (dash black line) were 5.83E-06 and 1.17E-04, respectively, according to GEC test.

**Figure 5:**
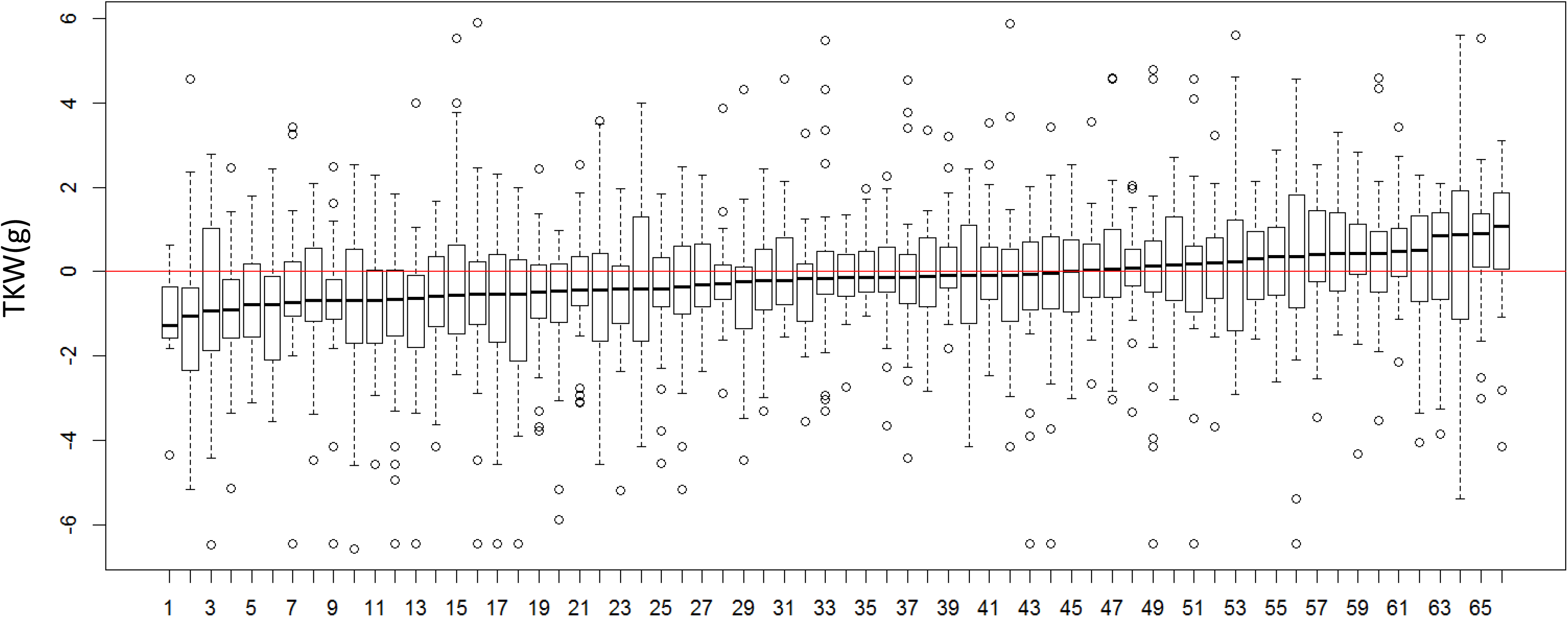
Effects of landraces alleles of grain size QTL relative to elite recurrent parents in the BC-NAM population. Each box plot illustrates the effects of exotic alleles of grain size QTL across 17 BC-NAM families derived from crosses between landraces and elite recurrent parental lines. Within each family, effects of a QTL was calculated as difference between average of the exotic allele and average of allele of the recurrent parental line. Sixty-six QTL identified in BC-NAM were ordered based on the median effect of the exotic alleles. Correspondence between numbers on x-axis and QTL identity is provided in Table S16.

A comparison of the QTL identified initially in both populations separately revealed 28 QTL in common, with 26 QTL being unique to the diversity panel and 38 QTL unique to the BC-NAM population (Figure 6), resulting in 92 QTL identified in total across populations. These 92 QTL were distributed across all 10 sorghum chromosomes (Table 3). Further examination revealed that the majority of the population specific QTL (22 out of 26 QTL unique to the diversity panel and 34 out of 38 QTL specific to BC-NAM population) were significantly associated with grain size parameters in the alternative population using the candidate region based association analysis (Table S12).

**Table 3.** Summary of the grain size QTL identified in the diversity and BC-NAM populations

| QTL ID | Chr | P-start | P-end | G-start | G-end | Traits affected |  | Overlap |
| --- | --- | --- | --- | --- | --- | --- | --- | --- |
|  |  |  |  |  |  | DP | BC-NAM |  |
| qGS1.1 | 1 | 1197434 | 1197434 | 3.18 | 3.18 | _VL__ | TVL__ | Y |
| qGS1.2 | 1 | 3551580 | 5918355 | 10.35 | 13.66 | TVLWH | TVLWH | Y |
| qGS1.3 | 1 | 6907712 | 7312784 | 16.89 | 18.27 | TVLWH | TVLWH | Y |
| qGS1.4 | 1 | 9747355 | 9819344 | 25.45 | 25.48 | TVLWH | _____ | Y |
| qGS1.5 | 1 | 12613546 | 13193518 | 31.25 | 33.54 | _V___ | TVLWH | Y |
| qGS1.6 | 1 | 17427801 | 17638035 | 43.49 | 43.60 | TVLW_ | TV__H | Y |
| qGS1.7 | 1 | 21347462 | 21457138 | 46.68 | 46.73 | TVLW_ | __L_H | N |
| qGS1.8 | 1 | 53201999 | 53936261 | 59.15 | 60.07 | TV__H | TVLWH | N |
| qGS1.9 | 1 | 62886564 | 62899303 | 94.75 | 94.76 | TVLW_ | TVLWH | N |
| qGS1.10 | 1 | 64100851 | 64100851 | 98.57 | 98.57 | TVLWH | TVLW_ | N |
| qGS1.11 | 1 | 67340797 | 67575811 | 115.03 | 116.85 | T_L__ | _V__H | Y |
| qGS1.12 | 1 | 68155790 | 68372488 | 123.06 | 125.38 | ____H | TVLWH | Y |
| qGS1.13 | 1 | 69803971 | 69803971 | 129.89 | 129.89 | ____H | __L_H | N |
| qGS1.14 | 1 | 75731963 | 75731963 | 156.66 | 156.66 | T_LWH | TVLW_ | Y |
| qGS2.1 | 2 | 1044805 | 1044805 | 0.10 | 0.10 | TV_WH | _VLW_ | N |
| qGS2.2 | 2 | 2902001 | 2977700 | 13.86 | 13.93 | TV_WH | TVLWH | N |
| qGS2.3 | 2 | 10134643 | 10134643 | 56.44 | 56.44 | ____H | TV__H | Y |
| qGS2.4 | 2 | 10821637 | 10821637 | 59.55 | 59.55 | TVLWH | TVLWH | Y |
| qGS2.5 | 2 | 57031598 | 57031598 | 88.94 | 88.94 | TV_WH | ____H | N |
| qGS2.6 | 2 | 57512343 | 57512343 | 93.17 | 93.17 | TV_WH | TV_W_ | N |
| qGS2.7 | 2 | 59194799 | 59194799 | 106.68 | 106.68 | _VL__ | TVLWH | N |
| qGS2.8 | 2 | 60589218 | 61587359 | 121.99 | 123.92 | TVLWH | TVLWH | N |
| qGS2.9 | 2 | 62351052 | 62351052 | 134.53 | 134.53 | T_____ | __LWH | N |
| qGS2.10 | 2 | 66339464 | 66573617 | 148.73 | 149.23 | ___W_ | ____H | Y |
| qGS2.11 | 2 | 67288493 | 67673759 | 152.32 | 152.63 | __L__ | TVLW_ | Y |
| qGS2.12 | 2 | 69049000 | 69049000 | 159.86 | 159.86 | TV_WH | TVLW_ | N |
| qGS2.13 | 2 | 70379875 | 70379875 | 166.28 | 166.28 | TVLW_ | TVLW_ | Y |
| qGS2.14 | 2 | 73307187 | 74155649 | 180.56 | 182.44 | TVLWH | __LWH | Y |
| qGS2.15 | 2 | 75213858 | 75310814 | 188.18 | 189.09 | TV_WH | TVLWH | Y |
| qGS3.1 | 3 | 617297 | 617297 | 0.68 | 0.68 | _____ | TVLWH | N |
| qGS3.2 | 3 | 1678176 | 1681138 | 4.04 | 4.05 | _____ | TVLW_ | N |
| qGS3.3 | 3 | 2972919 | 2972919 | 9.19 | 9.19 | TV_W_ | T__H | Y |
| qGS3.4 | 3 | 3597488 | 3597488 | 11.06 | 11.06 | TVLWH | TVLWH | Y |
| qGS3.5 | 3 | 7459268 | 7459268 | 31.12 | 31.12 | TV_W_ | TVL_H | N |
| qGS3.6 | 3 | 11842474 | 13099669 | 46.52 | 48.31 | TVLW_ | TVLWH | N |
| qGS3.7 | 3 | 27806456 | 52373180 | 56.79 | 58.13 | TV_WH | T_LWH | Y |
| qGS3.8 | 3 | 54907952 | 55299538 | 70.31 | 72.25 | TVLWH | TVLW_ | Y |
| qGS3.9 | 3 | 58595834 | 58595834 | 102.06 | 102.06 | T_WH | T_L_H | Y |
| qGS3.10 | 3 | 70185356 | 70185356 | 149.33 | 149.33 | TV_WH | TV_WH | Y |
| qGS3.11 | 3 | 70935875 | 70935875 | 151.89 | 151.89 | TVLWH | T__H | Y |
| qGS3.12 | 3 | 71518340 | 71802981 | 154.39 | 155.80 | TVLWH | TVLWH | Y |
| qGS4.1 | 4 | 4223889 | 4229830 | 27.71 | 27.75 | TV_WH | TVL__ | N |
| qGS4.2 | 4 | 4559844 | 4559844 | 29.99 | 29.99 | TVLWH | _____ | N |
| qGS4.3 | 4 | 7553755 | 7553755 | 55.81 | 55.81 | ____H | _____ | Y |
| qGS4.4 | 4 | 13834090 | 49255608 | 71.37 | 73.60 | TVLW_ | TVLWH | Y |
| qGS4.5 | 4 | 51365085 | 52404369 | 82.14 | 84.01 | ____H | TVLWH | Y |
| qGS4.6 | 4 | 54000479 | 54337837 | 94.49 | 94.71 | TVLW_ | TVLWH | Y |
| qGS4.7 | 4 | 57705140 | 58486661 | 106.83 | 106.99 | TV_WH | TVLW_ | Y |
| qGS4.8 | 4 | 61396481 | 61772353 | 121.78 | 123.63 | TVLW_ | TVLWH | Y |
| qGS4.9 | 4 | 66127254 | 66127254 | 152.27 | 152.27 | TVLW_ | __L__ | N |
| qGS5.1 | 5 | 3301226 | 3575220 | 22.87 | 23.25 | TVLWH | TVLW_ | Y |
| qGS5.2 | 5 | 38549008 | 54107776 | 64.58 | 65.76 | _VLW_ | TVLW_ | Y |
| qGS5.3 | 5 | 61218249 | 61218249 | 70.70 | 70.70 | _VL__ | _VL_H | N |
| qGS5.4 | 5 | 62745873 | 63339491 | 73.38 | 73.89 | TVLWH | TVLW_ | Y |
| qGS5.5 | 5 | 70963740 | 71211419 | 118.00 | 119.00 | TVLW_ | T_____ | N |
| qGS6.1 | 6 | 2325384 | 2491386 | 28.20 | 30.03 | _____ | TVLWH | N |
| qGS6.2 | 6 | 3407678 | 3594201 | 32.68 | 33.68 | T____H | TVLWH | Y |
| qGS6.3 | 6 | 39213130 | 43198259 | 50.50 | 52.75 | ___W_ | TVLWH | Y |
| qGS6.4 | 6 | 49661049 | 49672525 | 89.27 | 89.28 | __L__ | ____H | Y |
| qGS6.5 | 6 | 54391413 | 55406078 | 125.43 | 125.64 | __LWH | ____H | N |
| qGS6.6 | 6 | 58659162 | 58659162 | 157.96 | 157.96 | ___WH | __LW_ | Y |
| qGS6.7 | 6 | 59994049 | 59994049 | 162.94 | 162.94 | T__WH | TVLWH | Y |
| qGS7.1 | 7 | 1463261 | 1463261 | 16.76 | 16.76 | TVLW_ | _____ | N |
| qGS7.2 | 7 | 4810838 | 5140654 | 58.13 | 58.36 | TVLWH | _VLW_ | N |
| qGS7.3 | 7 | 14300295 | 38985351 | 72.21 | 72.30 | TVLWH | TV_W_ | Y |
| qGS7.4 | 7 | 52822136 | 54307603 | 74.35 | 75.24 | __LW_ | TV_W_ | Y |
| qGS7.5 | 7 | 55675278 | 55675278 | 78.67 | 78.67 | TVLWH | TVL__ | N |
| qGS7.6 | 7 | 60263942 | 60263942 | 106.63 | 106.63 | TVLW_ | ___W_ | Y |
| qGS8.1 | 8 | 5474894 | 5997557 | 58.65 | 59.32 | TVLWH | TVLWH | Y |
| qGS8.2 | 8 | 10712689 | 10712689 | 67.33 | 67.33 | TV_WH | _V___ | Y |
| qGS8.3 | 8 | 58069299 | 58069299 | 87.75 | 87.75 | ____H | _VLW_ | N |
| qGS8.4 | 8 | 59873105 | 59873105 | 99.35 | 99.35 | TV_H | TV_WH | N |
| qGS8.5 | 8 | 60306779 | 60394256 | 102.58 | 103.12 | TVLW_ | TVLWH | N |
| qGS8.6 | 8 | 60911363 | 60911363 | 106.38 | 106.38 | T_____ | TVLWH | N |
| qGS8.7 | 8 | 61382971 | 61382971 | 109.44 | 109.44 | ____H | T_____ | N |
| qGS8.8 | 8 | 61773861 | 61868131 | 111.98 | 112.60 | _____ | TVLWH | N |
| qGS8.9 | 8 | 61977263 | 61977263 | 118.53 | 118.53 | ____H | TVL_H | N |
| qGS8.10 | 8 | 62329242 | 62329242 | 127.16 | 127.16 | _____ | TV_H | N |
| qGS9.1 | 9 | 1188287 | 1188287 | 14.45 | 14.45 | TV_WH | T_____ | N |
| qGS9.2 | 9 | 8955692 | 15253529 | 67.33 | 69.39 | TVLW_ | TVLWH | Y |
| qGS9.3 | 9 | 42995635 | 45347175 | 75.64 | 76.17 | TVLWH | TVLWH | Y |
| qGS9.4 | 9 | 51441730 | 51441730 | 94.86 | 94.86 | TV_WH | __L_H | Y |
| qGS9.5 | 9 | 52757861 | 52795444 | 104.82 | 104.93 | T____H | T____H | N |
| qGS9.6 | 9 | 53947680 | 53959901 | 108.19 | 108.23 | T__W_ | TVLWH | N |
| qGS9.7 | 9 | 56099631 | 56099631 | 113.99 | 113.99 | _VL__ | TVLWH | N |
| qGS9.8 | 9 | 58170657 | 58198133 | 128.11 | 128.33 | TV_WH | TVLW_ | N |
| qGS10.1 | 10 | 777170 | 777170 | 13.22 | 13.22 | _V__H | T_L_H | N |
| qGS10.2 | 10 | 3158490 | 3445290 | 31.94 | 32.55 | TVLWH | T____H | Y |
| qGS10.3 | 10 | 9752387 | 12131845 | 53.93 | 56.00 | TVLWH | TVLWH | Y |
| qGS10.4 | 10 | 18074205 | 40511327 | 57.96 | 58.58 | TVLWH | TVLWH | Y |
| qGS10.5 | 10 | 56447108 | 56447108 | 89.00 | 89.00 | ___H | T___H | N |
| qGS10.6 | 10 | 58653903 | 60172861 | 102.30 | 103.56 | TVLWH | TVLW_ | N |

Chr indicates the chromosome on which the QTL is located. P-start provides the physical coordinate, based on v3.1.1 of the sorghum genome assembly, of the start of the QTL CI. P-end provides the physical coordinate of the end of the QTL CI. G-start provides the predicted cM genetic linkage position of the start of the QTL CI based on the sorghum consensus map (Mace et al 2009). G-end provides the predicted cM genetic linkage position of the end of the QTL CI. The traits affected columns indicate the traits significantly associated with the QTL in diversity panel (DP) and BC-NAM population respectively. The five traits were recorded as TVLWH, where T represents TKW, V represents volume, L represents length, W represents width and H represents thickness. The inclusion of the letters for a particular QTL means that the corresponding traits are affected by the QTL in the corresponding population. Missing letters in the order of TVLWH (represented as “\_”) mean that the corresponding traits are not affected by the QTL in corresponding population. The Overlap column indicates whether the QTL overlaps with previously published grain size QTL in sorghum. N means the QTL is not overlapped with previously published grain size QTL, while Y mean the QTL is overlapped with previously published QTL.

**Figure 6:**
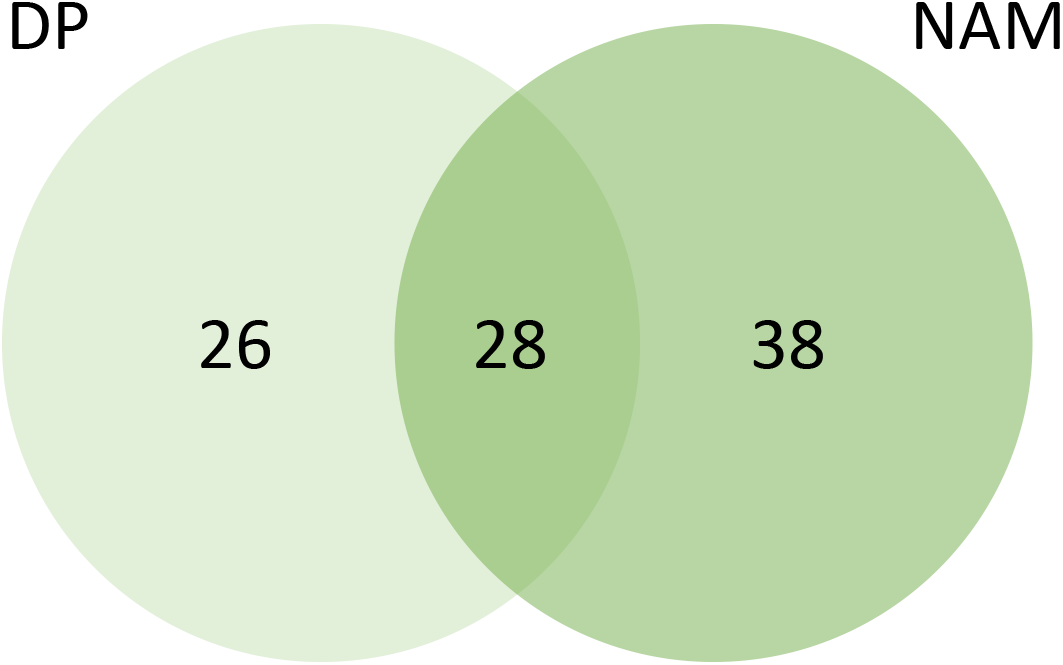
The overlap of grain size QTL identified in the diversity panel and the BC-NAM population.

The genetic basis of grain size has been the focus of 18 previous studies in sorghum, which used 21 bi-parental populations and reported 85 grain weight QTL (Table S5). Nearly three quarters (62) of these previously reported QTL co-located with QTL identified in this study. A significant overlap between the QTL identified in this study and grain size SNPs identified in rice (Huang *et al.*, 2012) was observed, with 60% of the rice grain size SNPs identified within a 1cM window of the sorghum QTL identified in this study (*p-value*<0.05, χ^2^ test), and with 85% of the rice grain size SNPs identified within 5 cM (Table S13). A similar trend was found when comparing with kernel size SNPs identified in maize (Liu *et al*., 2019), with 30% and 70% of maize kernel size SNPs identified within 1 cM and 5 cM window of sorghum QTL identified in this study (Table S13).

### Candidate genes for grain size QTL

To identify putative causative genes underlying the grain size QTL, 109 genes affecting grain size that were reported in rice and maize were collated and 111 orthologues in sorghum were identified (Table S6; Table S14). The candidate genes were enriched in the grain size QTL; 16 of the 111 candidate genes were found within a 0.1 cM window of a grain size QTL (*p-value*<0.05, χ^2^ test), and 36 identified within a 1 cM window (*p-value*<0.05, χ^2^ test) (Table S15). Some of the negative correlations between grain number and grain size are likely due to variation in the genotypes’ capacity to fill the grain as observed in this study. Given the treatment imposed in the current study to minimise variation due to grain filling capacity, we would expect that the previously reported candidate genes that have been shown to exert opposite effects on grain size and grain number would be less represented within our set of QTL. To explore this hypothesis, the candidate genes were divided into 3 groups based on whether they had been reported to exert opposite effects on grain number and grain size in the species in which they were cloned (ie whether the gene was associated with bigger grains and lower grain number). This approach identified 21 genes that showed opposite effects on grain size and grain number, 37 genes that did not show opposite effects, and 53 genes where this information was not reported (Table S15). It was found that the candidate genes that were reported to have an opposite effect on grain number and grain size were underrepresented in the overlap with the QTL from this study (Table S15).

One of the candidate genes, *SbGS3*, the orthologue of the rice grain size gene, *GS3* (*Mao et al., 2010*), co-located with *qGS1.9*, a QTL identified in both the diversity and BC-NAM populations, for grain length, weight, volume, width, and thickness. In the diversity panel, one SNP was identified that caused a C to A change in the fifth exon of *SbGS3*, turning a cysteine codon (TG**C**) to a premature stop codon (TG**A**), resulting in the 198 amino acid protein being truncated at the 140^th^ amino acid. However, the most significant SNP identified for *qGS1.9* was 11.48kb (0.01cM) away from *SbGS3*, rather than the large-effect SNP causing truncation at the 140^th^ amino acid. This may have been due to the low MAF (22 out of 837 individuals carried the premature stop codon allele) and the similar genetic background for the individuals carrying the premature stop codon allele. Transcriptomics expression data (Makita *et al.*, 2015) showed that *GS3* had comparable expression patterns in sorghum and rice, with high expression levels observed in the panicle and minimal expression in other tissue types covering a range of temporal and spatial variation (Figure S10A). Additional transgenic experiments, using RNAi, have demonstrated that inhibiting the expression of *SbGS3* increased grain weight by 9.4% on average in sorghum (Trusov et al., pers. comm.). Both positive and negative allelic effects from non-recurrent parental lines were observed in the BC-NAM compared with the recurrent parental line, R931945-2-2, indicating the existence of multiple alleles of *SbGS3* in the population (Figure S10 B).

## Discussion

Grain size is one of the most critical traits of cereal crops due to its direct contribution as a yield component, its importance as a quality attribute, and its contribution to fitness stemming from its impact on reproductive rates, emergence and establishment (Tao *et al.*, 2017; Westoby *et al.*, 1992). This study represents one of the largest and most comprehensive investigations of the genetic basis underling this trait in any cereal via a joint GWAS analysis of grain size in sorghum using a diversity panel (n=837) and a BC-NAM population (n=1421). The population size is the largest of all GWAS studies on grain size in any cereal crop, >5 times larger than the second largest GWAS studies in sorghum. As a result, a total number of 92 grain size QTL were identified. Enrichment of grain size candidate genes within these QTL was observed, and significant overlap between these 92 QTL and SNPs associated with grain size in rice and maize was found, highlighting common genetic control of this trait among cereals. In addition, the treatment of removing half the panicle during flowering facilitated minimisation of variation of grain size due to assimilate variability, and identification of genetic regions related to the genetic potential of grain size rather than the capacity to fill the grain. Therefore, these results provide an enhanced understanding of the genetic basis underlying grain size in sorghum and offer opportunities for breeders in sorghum and other cereal species to manipulate grain size to improve crop productivity and quality.

### The relationship between grain size and grain number in sorghum is mainly driven by genes acting before grain filling

A negative correlation between grain size and number is widely observed in cereals The nature of this correlation is complex (Sadras 2007) and is partially driven by the fact that potential maximum grain size and grain number are established before grain filling and by the availability of assimilates to enable seeds to reach their genetic potential (Yang *et al*., 2009; Gambín & Borrás, 2010). In this study, the half-head treatment resulted in an average increase in TKW of 8.54%, with significant variation among genotypes. The average change in seed weight is in line with previous estimates of sorghum’s capacity to compensate for loss of seeds (Sharma *et al.*, 2002). Seed weight in the full-head treatment explained ~70% of the genetic variation in grain size in the half-head treatment, indicating that while assimilate availability is a significant factor influencing grain size, most of the genetic factors driving the correlation between grain size and number in sorghum are acting prior to grain filling. Future yield gains may require that this association is better understood to determine if the negative association can be broken at some or all QTL.

### Shared genetic control of grain size dimensions

Sorghum grains vary in terms of length, thickness, width, volume and weight. Substantial genetic variation was observed for all of these parameters in both populations. Cross environment correlations of these grain size parameters within the same population were very high and the heritability of the traits was also moderate to high, indicating strong genetic control as previously observed (Prado *et al*., 2014; Tao *et al*., 2018). These findings, combined with a normal distribution of the phenotypes, indicate that all of these parameters are likely to be controlled by many genes with small to moderate effects. In addition, strong correlations were observed between the different grain size parameters in the same population, indicating they have a high degree of common genetic control. This was expected for parameters such as volume and weight and their correlations with the various measured dimensions, since the traits are linked mathematically. However, strong correlations between most of these parameters in both of the two populations indicate shared genetic control. In contrast to positive correlations between other grain size parameters in both populations, the grain thickness trait was negatively correlated with the other 4 parameters in the BC-NAM population but positively correlated with these parameters in the diversity panel. This is likely due to the different genetic backgrounds of the two populations. The diversity panel has a good representation of sorghum’s racial groups (Thurber *et al*., 2013). Although the BC-NAM also provides a good representation of the genetic variation of the racial groups of sorghum, because of the population structure (backcross derived based on a single reference parent), the genetic composition of each line is strongly biased towards the genetic background of the elite recurrent parent, which is primarily of *caudatum* origin.

### High-confidence genetic architecture of grain size revealed through independent multi-population GWAS

Population size is a key factor affecting the power of a GWAS study, as is the need to control for type I and type II errors (Visscher et al., 2017). The two populations used in this study, taken individually, are among the largest published to date in cereal crop on grain size. This size, coupled with the capacity to independently verify putative QTL in the alternative population, has provided us with a substantial advantage over previously published studies. The diversity panel contains representatives of most of the racial groups in the sorghum gene pool and is characterised by its fast LD decay. In contrast, the BC-NAM population consists of BC_1_ derived lines sharing ~80% of their genomes with the elite reference parent (Jordan *et al*., 2011), which provides an opportunity to assess exotic alleles in the genetic background of an elite inbred. The BC-NAM population has lower genetic diversity, a more balanced structure and greater LD and therefore potentially more power to detect QTL. The joint use of these complementary populations has also provided a more comprehensive understanding of the genetic architecture of grain size, and ready-to-use information on the effect of QTL alleles in elite breeding material.

The high number (92) of QTL identified in this study and their relatively small individual effects highlight the genetic complexity of grain size. Over half of the QTL identified in this study (49) were co-located with QTL identified in previous sorghum studies (Table 3) with < 30% of the previous QTL not detected in this study (Table S5). Care must be taken with interpreting this result however, since the small population sizes of the previous studies limited their power to detect QTL and resulted in large confidence intervals to the extent that they often encompassed a number of the QTL detected in this study. In addition, two of the previous studies identified QTL from crosses between wild and domestic sorghum (Paterson *et al.*, 1995; Tao *et al*., 2018), which may not have been detected in our study as wild species were not included in either population, plus none of the other studies attempted to minimise variation in the capacity to fill grain. Hence, it is likely that some of the QTL previously detected may not be segregating in our study.

Our study was internally consistent, with 83% of the QTL detected being associated with multiple traits, in line with the correlations observed between the phenotypes. In addition, 91% of the QTL were supported by evidence from the two independent populations either via co-located QTL or with support from the candidate region analysis, resulting in only 8 population-specific QTL being identified. Given that the two populations were independent and differed in both genetic composition, power and LD, the high correspondence between the two studies suggests that we identified the majority of the loci for grain size in cultivated sorghum.

### Significant allelic diversity exists in grain size QTL

The BC-NAM population provided us with the opportunity to investigate variation in the effects of different QTL alleles in an elite genetic background. Our results suggest the presence of multiple alleles at the majority of the QTL with effects of the non-recurrent parental alleles that were both greater and smaller that the recurrent parent allele observed at most loci (Figure 5). The distribution of the allele effects provides interesting insights into selection for the grain size trait in an elite breeding program. There was weak tendency for the elite parental alleles to contribute to larger grain size compared to the exotic parent. This is similar to an earlier study of flowering time in the same population where positive and negative allele effects relative to the recurrent parental allele were similarly distributed (Mace *et al*., 2013). The lack of bias observed in the current study suggests modern sorghum breeding with its focus on increasing grain yield has not selected for grain size. The lack of strong selection for grain size by modern breeders is interesting and suggests a potential physiological restriction to increasing grain yield by selection for larger grain. This is consistent with the observation that much of the progress in enhancing grain yield in modern breeding of cereals has been achieved by selection for increased grain number rather than grain size. Clearly, further research is required before breeders attempt to increase grain yield by introducing alleles for larger grain size. However, given the range of allele effects, it is likely some opportunities exist to simultaneously improve grain yield and grain size, eg through manipulation of the duration of grain filling (Yang *et al*., 2010).

### Correspondence of loci controlling grain size among cereals

Sorghum shares close ancestry with maize and rice, with orthologous genes from these species commonly sharing the same function (Bolot *et al.*, 2009; Paterson *et al*., 1995). A significant association was found between the locations of the QTL in this study and GWAS signals for grain size in rice (Huang *et al*., 2012) and maize (Liu *et al*., 2019), and orthologous of cloned grain size genes from rice and maize. Both findings strongly support a common genetic architecture underlying this trait across these cereals. This provides opportunities for breeders to exploit information from other cereals, as genes identified from large scale GWAS in one species can be targeted for allele mining or diversity creation via gene editing in another species.

As an example, the gene *SbGS3*, the orthologue of the cloned gene *Grain Size 3* in rice, was found to control variation of grain size in sorghum. In addition to the presence of a large-effect SNP in the gene causing a premature stop codon, our GWAS identified a SNP that was 11.48 kb away from this gene as significantly associated with variation of grain size. *SbGS3* showed a similar expression pattern as *GS3* in rice with high expression in early inflorescence, indicating the same role in grain development. Further evidence from transgenic experiments has shown that inhibiting the expression of *SbGS3* increased grain weight by 9.4% on average in sorghum (Trusov et al., pers. comm.). It has also been reported that the orthologue of *GS3* in maize, *ZmGS3*, was involved in kernel development (Li *et al.*, 2010). GS3 contains four domains: an OSR domain in the N terminus, a transmembrane domain, a TNFR/NGFR family cysteine-rich domain, and a VWFC in the C terminus (*Mao et al., 2010*). In rice, overexpression of a truncated cDNA sequence of *GS3* where the VWFC domain is deleted, produced shorter grains (Mao *et al.*, 2010). The premature stop change of *SbGS3* caused a truncation of the VWFC domain. Further investigation is required to see whether comparable effects caused by the truncation occur in sorghum.

## Conclusion

Grain size is a key quantitative trait of cereal crops. This study presents a comprehensive genetic dissection of grain size in sorghum using two large and complementary populations, resulting in the identification of a total of 92 QTL related to grain size. Significant overlap was found between the QTL identified in this study and grain size GWAS signals in rice and maize, and orthologues of cloned grain size genes from rice and maize, supporting a common genetic architecture underlying this trait across these cereals. These findings provide a better understanding of the genetic architecture of grain size in sorghum, and pave the way for exploration of its underlying molecular and physiological mechanisms in cereal crops and its manipulation in breeding practices.

## Acknowledgments

We acknowledge access to background IP from the Grains Research and Development Corporation and support from the University of Queensland and the Department of Agriculture and Fisheries Queensland.

## Author contributions

D.J., E.M., and I.G. conceived and designed the experiments: Y.T., X.Z., X.W. and A.C. collected data; Y.T., C.H., A.H., E.O., D.J., and E.M. analysed data; Y.T. wrote the manuscript. E.M., E.O., I.G., and D.J. revised the manuscript. All authors read and approved the final manuscript.

## Supplementary materials

Supplementary Table S1 List of the diversity panel. a, country and region information was obtained from USDA GRIN (https://npgsweb.ars-grin.gov/gringlobal/search.aspx) and personal communication with Jeffery A. Dahlberg. b, race was classified based on population structure analysis, as explained in materials and methods. c records overlap between diversity panel and USSAP according to GRIN.

Supplementary Table S2 Summary of the BC-NAM

Supplementary Table S3 Summary of field experiments

Supplementary Table S4 Racial group breakdown of the diversity panel

Supplementary Table S5 List of reported grain size QTL in previous studies

Supplementary Table S6 Lis of reported grain size genes in rice and maize

Supplementary Table S7 Effect of population structure on grain size. Statistical significance was assessed using one-way analyses of variance (ANOVAs) followed by Tukey’s HSD tests for multiple comparisons.

Supplementary Table S8 Marker Trait associations in the diversity panel. Physical position of SNPs were according to *Sorghum bicolor* v3.1.1.

Supplementary Table S9 Effects of QTL in the diversity panel

Supplementary Table S10 Marker Trait associations in BC-NAM. Physical position of SNPs were according to *Sorghum bicolor* v3.1.1.

Supplementary Table S11 Effects of QTL in BC-NAM

Supplementary Table S12 Tests of associations between population-specific QTL with grain size parameters in the alternative population

Supplementary Table S13 Comparison of grain size regions identified in rice and maize with grain size QTL identified in this study

Supplementary Table S14 List of grain size candidate genes in sorghum

Supplementary Table S15 Summary of candidate genes within close vicinities of grain size QTL

Supplementary Table S16 Correspondence between numbers on x-axis of Figure 5 and identities of QTL identified in BC-NAM.

Supplementary Figure S1 PCA and LD of BC-NAM. A, PCA of BC-NAM. B, PCA of diversity panel shows the BC-NAM parental lines capture a good proportion of diversity in the diversity panel. Dots in red colour represent non-recurrent parental lines. Dots in green represent recurrent parental lines. C, LD decay in the BC-NAM.

Supplementary Figure S2 Bi-plots show PCA analysis of grain size parameters measured in the diversity panel.

Supplementary Figure S3 Bi-plots show PCA analysis of grain size parameters traits measured in BC-NAM.

Supplementary Figure S4 LD decay, Population structure and PCA of the diversity panel. A, LD decay of the diversity panel. B, population structure of the diversity panel. C, PCA of the diversity panel, dots in colour indicate representatives for each racial group.

Supplementary Figure S5 Distribution of SNPs across sorghum genome in the diversity panel.

Supplementary Figure S6 Distribution of SNPs across sorghum genome in BC-NAM.

Supplementary Figure S7. Correlation of grain size between HH and FH. A, correlation of TKW between HH and FH. B, correlation of volume between HH and FH.

Supplementary Figure S8 Manhattan plots and Q-Q plots show GWAS analysis of PCs derived from grain size parameters in the diversity panel.

Supplementary Figure S9 Manhattan plots and Q-Q plots show GWAS analysis of PCs derived from grain size parameters in BC-NAM.

Supplementary Figure 10: *SbGS3* (Sobic.001G341700) controls grain size in sorghum. A) Expression pattern of *sbGS3* in sorghum. B) Varying effects of *sbGS3* alleles in non-recurrent parental lines compared with the recurrent parental line.

