## Supplemental Table 1 for "Large-scale GWAS in sorghum reveals common genetic control of grain size among cereals"

Table S1 List of the diversity panel

| ID | Country[a] | Region[a] | Group[b] | TKW | Volume | Length | Thickness | Width | Parental line of BC-NAM | USSAP[c] |
|---|---|---|---|---|---|---|---|---|---|---|
| B599 | NA | NA | mixed | 29.93 | 22.80 | 4.15 | 3.19 | 3.28 | No | No |
| Early hegari | NA | NA | mixed | 26.03 | 25.10 | 4.30 | 3.16 | 3.49 | No | 642992 |
| B010054 | NA | NA | mixed | 27.81 | 25.41 | 4.21 | 3.29 | 3.49 | No | No |
| B35 | NA | NA | mixed | 32.89 | 26.02 | 4.14 | 3.40 | 3.53 | No | BTx642 |
| B923296 | NA | NA | mixed | 33.38 | 28.96 | 4.54 | 3.39 | 3.60 | No | No |
| B963676 | NA | NA | mixed | 31.45 | 29.37 | 4.52 | 3.44 | 3.61 | No | No |
| B986604 | NA | NA | mixed | 27.08 | 26.72 | 4.42 | 3.32 | 3.46 | No | No |
| BTx623 | NA | NA | mixed | 30.23 | 27.78 | 4.64 | 3.08 | 3.67 | No | 564163 |
| M35-1 | NA | NA | mixed | 29.74 | 25.72 | 4.34 | 3.26 | 3.46 | Yes | 656047 |
| RTAM422 | NA | NA | mixed | 38.29 | 34.60 | 4.83 | 3.44 | 3.93 | Yes | No |
| RTx430 | NA | NA | mixed | 35.52 | 32.96 | 4.81 | 3.37 | 3.84 | Yes | 655996 |
| RTx7000 | NA | NA | mixed | 33.66 | 30.94 | 4.65 | 3.30 | 3.81 | Yes | 655986 |
| SC10-14E | United States | Americas | mixed | 28.21 | 26.44 | 4.46 | 3.31 | 3.40 | No | No |
| SC100-14E | Nigeria | Western Africa | mixed | 34.08 | 27.41 | 4.69 | 3.07 | 3.61 | No | No |
| SC101-14E | South Africa | Southern Africa | mixed | 22.63 | 25.33 | 4.50 | 3.26 | 3.29 | No | 533954 |
| SC1014-14E | Ethiopia | Eastern Africa | mixed | 29.03 | 21.69 | 4.06 | 3.08 | 3.25 | No | 576375 |
| SC1015-14E | United States | Americas | mixed | 32.10 | 24.11 | 4.07 | 3.28 | 3.43 | No | No |
| SC1017-14E | Ethiopia | Eastern Africa | mixed | 32.53 | 25.17 | 4.16 | 3.30 | 3.44 | No | 576376 |
| SC1019-14E | United States | Americas | mixed | 35.54 | 24.89 | 4.21 | 3.19 | 3.52 | No | 656071 |
| SC1021-6 | United States | Americas | mixed | 26.49 | 24.67 | 4.28 | 3.19 | 3.42 | No | No |
| SC1022-14E | Ethiopia | Eastern Africa | Kafir | 18.40 | 21.06 | 4.20 | 3.22 | 2.98 | No | No |
| SC1024-9 | United States | Americas | mixed | 31.11 | 30.30 | 4.57 | 3.46 | 3.64 | No | No |
| SC1025-14E | Ethiopia | Eastern Africa | mixed | 16.30 | 19.31 | 4.22 | 3.07 | 2.81 | No | No |
| SC103-14E | NA | NA | mixed | 30.46 | 30.28 | 4.59 | 3.37 | 3.71 | Yes | 533752 |
| SC1031-12E | United States | Americas | mixed | 18.61 | 21.46 | 3.97 | 3.15 | 3.23 | No | No |
| SC1033-14E | Ethiopia | Eastern Africa | mixed | 33.06 | 29.57 | 4.51 | 3.41 | 3.63 | No | 576426 |
| SC1038-14E | Ethiopia | Eastern Africa | mixed | NA | NA | NA | NA | NA | No | 576381 |
| SC1039-14E | Ethiopia | Eastern Africa | Kafir | 26.57 | 24.04 | 4.33 | 3.06 | 3.43 | No | No |
| SC1040-14E | United States | Americas | mixed | 28.57 | 27.36 | 4.43 | 3.28 | 3.57 | No | No |
| SC1046-14E | Ethiopia | Eastern Africa | mixed | 36.63 | 33.91 | 4.75 | 3.45 | 3.91 | No | No |
| SC1049-14E | Ethiopia | Eastern Africa | Asian durras | 25.85 | 24.46 | 4.19 | 3.17 | 3.51 | No | No |
| SC105-14E | India | Asia | mixed | 22.99 | 20.65 | 4.00 | 2.99 | 3.23 | No | No |
| SC1055-14E | Sudan | Eastern Africa | mixed | 29.49 | 28.19 | 4.46 | 3.22 | 3.73 | No | 595739 |
| SC1057-14E | Uganda | Eastern Africa | mixed | 28.10 | 24.77 | 4.18 | 3.23 | 3.47 | No | 595740 |
| SC106-14E | United States | Americas | E African durra | 17.02 | 20.59 | 3.94 | 3.33 | 2.99 | No | No |
| SC1063-14E | NA | NA | mixed | 17.02 | 20.92 | 4.13 | 3.05 | 3.13 | No | 595741 |
| SC1065-14E | United States | Americas | mixed | 32.33 | 28.08 | 4.56 | 3.13 | 3.72 | No | No |
| SC1067-14E | Senegal | Western Africa | mixed | 32.67 | 26.51 | 4.32 | 3.27 | 3.58 | No | No |
| SC1069-14E | Nigeria | Western Africa | Caudatum | 29.58 | 26.77 | 4.53 | 3.22 | 3.51 | No | No |
| SC1070-14E | Nigeria | Western Africa | mixed | NA | NA | NA | NA | NA | No | 576385 |
| SC1072-14 | United States | Americas | mixed | 27.93 | 24.82 | 4.23 | 3.29 | 3.41 | No | No |
| SC1074-12E | United States | Americas | mixed | 29.73 | 27.02 | 4.65 | 3.07 | 3.59 | No | No |
| SC1076-12E | United States | Americas | mixed | 35.01 | 26.80 | 4.43 | 3.18 | 3.60 | No | 597960 |
| SC1077-14E | Nigeria | Western Africa | Caudatum | 29.22 | 25.83 | 4.30 | 3.14 | 3.63 | No | 597961 |

| | | | | | | | | | | |
|---|---|---|---|---|---|---|---|---|---|---|
| SC108-14E | NA | NA | Caudatum | 33.35 | 30.12 | 4.62 | 3.27 | 3.80 | Yes | 533792 |
| SC1080-14E | South Africa | Eastern | Kafir | 28.77 | 26.61 | 4.40 | 3.24 | 3.55 | No | 576422 |
| SC1082-6 | United States | Americas | mixed | 28.77 | 25.55 | 4.37 | 3.15 | 3.52 | No | No |
| SC1083-14E | Nigeria | Western Africa | mixed | NA | NA | NA | NA | NA | No | No |
| SC1084-14E | Nigeria | Western Africa | mixed | NA | NA | NA | NA | NA | No | No |
| SC1085-14E | India | Asia | mixed | 49.09 | 31.94 | 4.70 | 3.32 | 3.84 | No | 576401 |
| SC1088-14E | United States | Americas | mixed | 30.09 | 25.83 | 4.29 | 3.25 | 3.54 | No | No |
| SC109-14E | 0 | 0 | Caudatum | 31.62 | 27.07 | 4.34 | 3.30 | 3.60 | No | No |
| SC1097-3 | United States | Americas | mixed | 28.24 | 24.47 | 4.04 | 3.31 | 3.46 | No | No |
| SC110-14E | Ethiopia | Eastern Africa | Caudatum | 34.32 | 32.50 | 4.71 | 3.40 | 3.87 | No | 533794 |
| SC1101-14E | United States | Americas | mixed | 41.07 | 28.81 | 4.58 | 3.18 | 3.73 | No | No |
| SC1103-14E | Nigeria | Western Africa | mixed | 35.17 | 32.05 | 4.78 | 3.42 | 3.72 | No | No |
| SC1104-14E | Uganda | Eastern Africa | mixed | 24.39 | 25.13 | 4.45 | 3.19 | 3.36 | No | 576435 |
| SC1107-9 | United States | Americas | mixed | 19.37 | 21.99 | 4.19 | 3.09 | 3.21 | No | No |
| SC1109-14E | India | Asia | mixed | 21.85 | 21.00 | 3.99 | 3.03 | 3.30 | No | No |
| SC111-14E | United States | Americas | Caudatum | 40.08 | 29.49 | 4.54 | 3.28 | 3.75 | No | No |
| SC1111-14E | Sudan | Eastern Africa | mixed | 22.63 | 24.79 | 4.28 | 3.24 | 3.40 | No | No |
| SC1114-6 | United States | Americas | mixed | NA | NA | NA | NA | NA | No | No |
| SC1116-14E | Nigeria | Western Africa | mixed | NA | NA | NA | NA | NA | No | No |
| SC1117-14E | NA | NA | mixed | 42.06 | 35.91 | 5.23 | 3.20 | 4.00 | No | No |
| SC1118-14E | United States | Americas | mixed | 21.69 | 22.56 | 4.13 | 3.22 | 3.26 | No | No |
| SC112-14E | Ethiopia | Eastern Africa | Caudatum | 34.49 | 28.47 | 4.39 | 3.44 | 3.62 | No | No |
| SC1123-14E | Nigeria | Western Africa | Guinea | 27.91 | 25.72 | 4.41 | 3.35 | 3.33 | No | No |
| SC1124-14E | NA | NA | Guinea | 32.61 | 31.89 | 4.74 | 3.35 | 3.78 | No | 576418 |
| SC1125-14E | Nigeria | Western Africa | Guinea | NA | NA | NA | NA | NA | No | No |
| SC113-14 | United States | Americas | mixed | 22.62 | 20.45 | 3.77 | 3.28 | 3.16 | No | No |
| SC1133-6 | United States | Americas | mixed | 30.43 | 26.41 | 4.22 | 3.41 | 3.50 | No | No |
| SC114-14E | Uganda | Eastern Africa | mixed | 32.13 | 27.38 | 4.47 | 3.31 | 3.51 | No | No |
| SC115-14E | Uganda | Eastern Africa | mixed | 22.56 | 24.75 | 4.30 | 3.25 | 3.38 | No | 533965 |
| SC1154-14E | NA | NA | mixed | 27.15 | 25.56 | 4.35 | 3.25 | 3.48 | No | 595720 |
| SC1155-14E | NA | NA | mixed | 32.10 | 28.55 | 4.47 | 3.41 | 3.59 | No | 576425 |
| SC1158-5bk | Ethiopia | Eastern Africa | mixed | NA | NA | NA | NA | NA | No | 597957 |
| SC1160-14E | Ethiopia | Eastern Africa | mixed | NA | NA | NA | NA | NA | No | No |
| SC1166-6 | United States | Americas | mixed | 23.06 | 23.46 | 4.06 | 3.26 | 3.37 | No | No |
| SC1170-6 | United States | Americas | mixed | 28.36 | 24.03 | 4.14 | 3.21 | 3.44 | No | No |
| SC1172-14E | United States | Americas | mixed | 23.51 | 22.14 | 4.15 | 3.11 | 3.26 | No | No |
| SC1177-14E | United States | Americas | E African durra | 17.15 | 20.12 | 3.94 | 3.14 | 3.06 | No | No |
| SC1178-9 | United States | Americas | mixed | 21.47 | 24.38 | 4.28 | 3.22 | 3.37 | No | No |
| SC1179-6 | United States | Americas | mixed | 26.78 | 25.06 | 4.10 | 3.39 | 3.44 | No | No |
| SC118-14E | NA | NA | mixed | 25.55 | 23.39 | 4.06 | 3.23 | 3.39 | No | 533759 |
| SC1184-14E | NA | NA | mixed | NA | NA | NA | NA | NA | No | No |
| SC1186-14E | United States | Americas | mixed | 31.34 | 28.83 | 4.41 | 3.40 | 3.66 | No | No |
| SC119-14E | Zimbabwe | Eastern Africa | mixed | 26.87 | 24.16 | 4.21 | 3.12 | 3.49 | No | No |
| SC1193-3 | United States | Americas | mixed | 26.99 | 26.92 | 4.42 | 3.19 | 3.62 | No | No |
| SC12-14E | Ethiopia | Eastern Africa | mixed | 16.93 | 19.94 | 3.94 | 3.20 | 3.01 | No | No |
| SC120-14E | NA | NA | Caudatum | 27.94 | 25.99 | 4.39 | 3.17 | 3.54 | No | 533760 |
| SC1201-14E | United States | Americas | mixed | 28.81 | 26.02 | 4.62 | 3.21 | 3.35 | No | 595743 |

| | | | | | | | | | | |
|---|---|---|---|---|---|---|---|---|---|---|
| SC1201-6-3 | United States | Americas | mixed | 30.94 | 27.38 | 4.46 | 3.33 | 3.49 | No | 595743 |
| SC1203-14E | Brazil | Americas | mixed | 22.39 | 19.97 | 3.94 | 3.14 | 3.05 | No | 576437 |
| SC121-14E | South Africa | Southern Africa | mixed | 34.92 | 27.79 | 4.61 | 3.17 | 3.60 | No | 533961 |
| SC1211-14E | Guatemala | Americas | mixed | 33.46 | 22.71 | 3.87 | 3.38 | 3.31 | No | 595744 |
| SC1214-14E | Burkina Faso | Western Africa | mixed | 20.23 | 26.28 | 4.41 | 3.22 | 3.51 | No | 595745 |
| SC1215-13E | Niger | Western Africa | mixed | 24.50 | 20.74 | 3.82 | 3.18 | 3.27 | No | 656112 |
| SC1218 | United States | Americas | mixed | 42.21 | 31.81 | 5.27 | 3.05 | 3.75 | No | 656074 |
| SC123-14E | Ethiopia | Eastern Africa | mixed | 25.67 | 25.04 | 4.42 | 3.18 | 3.40 | No | No |
| SC124-14E | Ethiopia | Eastern Africa | mixed | 20.83 | 20.03 | 3.88 | 3.15 | 3.11 | No | 533919 |
| SC1261-14E | United States | Americas | Caudatum | 28.61 | 28.34 | 4.50 | 3.18 | 3.75 | No | No |
| SC1262-14E | United States | Americas | Caudatum | 26.39 | 25.89 | 4.42 | 3.14 | 3.54 | No | No |
| SC1263-14E | United States | Americas | mixed | 29.30 | 24.76 | 4.29 | 3.13 | 3.50 | No | No |
| SC1271-13E | United States | Americas | Caudatum | 32.00 | 28.19 | 4.96 | 3.14 | 3.51 | No | 656076 |
| SC1272-5 | NA | NA | mixed | 23.84 | 18.59 | 3.72 | 3.05 | 3.12 | No | No |
| SC1287-14E | India | Asia | mixed | 24.70 | 24.70 | 4.37 | 3.21 | 3.36 | No | No |
| SC1293-14E | Senegal | Western Africa | mixed | 30.38 | 27.32 | 4.38 | 3.30 | 3.61 | No | No |
| SC13-14E | Ethiopia | Eastern Africa | mixed | 16.51 | 17.87 | 3.72 | 3.16 | 2.90 | No | 534123 |
| SC1300-14E | United States | Americas | Caudatum | 36.46 | 26.65 | 4.33 | 3.25 | 3.59 | No | No |
| SC1302-14E | United States | Americas | Caudatum | 35.33 | 30.62 | 4.58 | 3.39 | 3.75 | No | No |
| SC1305-14E | United States | Americas | mixed | 33.29 | 30.37 | 4.76 | 3.24 | 3.74 | No | No |
| SC1307-14E | United States | Americas | Caudatum | 30.39 | 29.45 | 4.58 | 3.24 | 3.74 | No | No |
| SC1313-14E | United States | Americas | Caudatum | 41.16 | 30.35 | 4.54 | 3.37 | 3.74 | No | No |
| SC1314-14E | United States | Americas | mixed | 34.86 | 30.70 | 4.90 | 3.16 | 3.75 | No | No |
| SC1316-14E | Ethiopia | Eastern Africa | mixed | 34.53 | 29.94 | 4.69 | 3.31 | 3.66 | No | No |
| SC1317-14E | United States | Americas | Caudatum | 33.93 | 27.97 | 4.76 | 3.09 | 3.61 | No | No |
| SC1318-14E | United States | Americas | Caudatum | 34.43 | 29.06 | 4.44 | 3.41 | 3.68 | No | No |
| SC1319-14E | United States | Americas | Caudatum | 40.75 | 33.83 | 4.75 | 3.49 | 3.88 | No | 597964 |
| SC132-12E | NA | NA | mixed | 32.40 | 27.69 | 4.37 | 3.36 | 3.58 | No | No |
| SC1320-14E | NA | NA | Caudatum | 39.23 | 33.86 | 5.22 | 3.15 | 3.90 | No | 597967 |
| SC1321-14E | NA | NA | Caudatum | 34.05 | 29.74 | 4.56 | 3.30 | 3.74 | No | 597968 |
| SC1322-14E | NA | NA | mixed | 41.38 | 32.63 | 5.09 | 3.16 | 3.84 | No | No |
| SC1325-14E | United States | Americas | mixed | 27.92 | 26.48 | 4.49 | 3.01 | 3.67 | No | No |
| SC1328-14E | United States | Americas | mixed | 27.51 | 26.14 | 4.45 | 3.19 | 3.53 | No | 597971 |
| SC1329-14E | United States | Americas | mixed | 40.31 | 33.56 | 4.86 | 3.37 | 3.89 | No | 597972 |
| SC1330-14E | NA | NA | mixed | 34.59 | 29.67 | 4.63 | 3.23 | 3.76 | No | 597973 |
| SC1332-14E | United States | Americas | mixed | 25.16 | 23.01 | 4.39 | 2.99 | 3.32 | No | No |
| SC1333-14E | United States | Americas | mixed | NA | NA | NA | NA | NA | No | No |
| SC1334-5 | NA | NA | mixed | NA | NA | NA | NA | NA | No | No |
| SC1337-14E | NA | NA | mixed | 26.04 | 23.14 | 4.16 | 3.20 | 3.30 | No | 597976 |
| SC1339-14E | United States | Americas | mixed | 30.32 | 26.47 | 4.57 | 3.13 | 3.49 | No | No |
| SC134-14E | United States | Americas | mixed | 26.53 | 24.66 | 4.24 | 3.26 | 3.40 | No | 656114 |
| SC1341-14E | United States | Americas | mixed | 26.67 | 22.98 | 4.29 | 3.07 | 3.29 | No | No |
| SC1342-14E | United States | Americas | mixed | NA | NA | NA | NA | NA | No | No |
| SC1345-14E | United States | Americas | mixed | 32.94 | 27.85 | 4.57 | 3.14 | 3.71 | No | 597980 |
| SC135-14E | Ethiopia | Eastern Africa | mixed | 28.46 | 25.55 | 4.45 | 3.30 | 3.32 | No | 534148 |
| SC1351-14E | United States | Americas | mixed | 39.92 | 33.53 | 4.70 | 3.58 | 3.76 | No | No |
| SC1356-14E | United States | Americas | mixed | 31.14 | 29.51 | 4.69 | 3.35 | 3.57 | No | 597982 |

| | | | | | | | | | | |
|---|---|---|---|---|---|---|---|---|---|---|
| SC137-14E | Ethiopia | Eastern Africa | E African durra | 20.96 | 22.28 | 4.12 | 3.08 | 3.28 | No | No |
| SC139-14E | Ethiopia | Eastern Africa | mixed | 25.93 | 28.43 | 4.66 | 3.21 | 3.61 | No | No |
| SC140-14E | Ethiopia | Eastern Africa | mixed | 23.06 | 25.40 | 4.45 | 3.17 | 3.43 | No | No |
| SC141-14E | Ethiopia | Eastern Africa | mixed | 24.61 | 24.39 | 4.17 | 3.21 | 3.46 | No | No |
| SC142-14E | Ethiopia | Eastern Africa | mixed | 39.09 | 33.91 | 5.00 | 3.32 | 3.83 | No | No |
| SC1424-2 | NA | NA | mixed | 25.90 | 27.97 | 4.57 | 3.28 | 3.55 | No | 656078 |
| SC1426-10E | NA | NA | mixed | 27.94 | 23.79 | 4.42 | 2.99 | 3.41 | No | 656079 |
| SC1440-8 | NA | NA | mixed | 15.74 | 18.07 | 4.00 | 2.92 | 2.92 | No | 656115 |
| SC145-14E | United States | Americas | E African durra | 16.21 | 17.55 | 3.87 | 2.95 | 2.88 | No | 656082 |
| SC146-14E | Ethiopia | Eastern Africa | mixed | 31.08 | 29.30 | 4.37 | 3.51 | 3.66 | No | No |
| SC147-14E | Ethiopia | Eastern Africa | mixed | 21.32 | 22.50 | 4.24 | 3.17 | 3.18 | No | No |
| SC1476-8 | NA | NA | mixed | 35.74 | 25.42 | 4.27 | 3.19 | 3.52 | No | 656087 |
| SC15-14E | Ethiopia | Eastern Africa | mixed | 27.09 | 26.57 | 4.57 | 3.25 | 3.41 | No | 534124 |
| SC150-14E | United States | Americas | mixed | 27.12 | 25.06 | 4.38 | 3.25 | 3.36 | No | No |
| SC154-14E | Ethiopia | Eastern Africa | mixed | 17.43 | 21.01 | 4.04 | 3.08 | 3.18 | No | No |
| SC155-14E | Ethiopia | Eastern Africa | mixed | 25.44 | 26.19 | 4.50 | 3.15 | 3.51 | No | 534155 |
| SC156-14E | United States | Americas | E African durra | 25.04 | 22.20 | 4.05 | 3.24 | 3.22 | No | No |
| SC157-6 | NA | NA | mixed | 24.14 | 20.15 | 3.79 | 3.25 | 3.10 | No | No |
| SC158-14E | Ethiopia | Eastern Africa | mixed | 27.75 | 26.52 | 4.43 | 3.21 | 3.55 | No | No |
| SC159-11E | United States | Americas | mixed | 27.63 | 28.96 | 4.66 | 3.28 | 3.61 | No | No |
| SC16-14E | Ethiopia | Eastern Africa | mixed | 20.03 | 21.65 | 4.31 | 3.04 | 3.16 | No | No |
| SC161-14E | Ethiopia | Eastern Africa | mixed | 24.27 | 24.82 | 4.43 | 3.31 | 3.22 | No | No |
| SC165-14E | Ethiopia | Eastern Africa | mixed | 33.50 | 24.17 | 4.03 | 3.34 | 3.41 | No | No |
| SC166-14E | Ethiopia | Eastern Africa | E African durra | 29.49 | 26.66 | 4.31 | 3.39 | 3.48 | No | 533924 |
| SC167-14E | Ethiopia | Eastern Africa | mixed | 24.95 | 26.53 | 4.43 | 3.25 | 3.50 | No | No |
| SC17-14E | Ethiopia | Eastern Africa | mixed | 21.36 | 22.05 | 4.07 | 3.22 | 3.20 | No | 533903 |
| SC170-14E | NA | NA | mixed | 30.69 | 26.56 | 4.49 | 3.26 | 3.46 | No | 534157 |
| SC170-6-17 | NA | NA | mixed | 30.52 | 28.43 | 4.60 | 3.12 | 3.74 | No | 656068 |
| SC170-6-8 | NA | NA | mixed | 29.43 | 26.94 | 4.44 | 3.29 | 3.50 | No | No |
| SC171-14E | Ethiopia | Eastern Africa | mixed | 34.47 | 27.44 | 4.30 | 3.43 | 3.55 | No | No |
| SC172-12E | NA | NA | Caudatum | 49.47 | 36.75 | 5.31 | 3.40 | 3.86 | No | 656117 |
| SC173-14E | NA | NA | Caudatum | 34.20 | 27.43 | 4.39 | 3.28 | 3.60 | No | 533799 |
| SC175-14E | Ethiopia | Eastern Africa | Caudatum | 28.46 | 26.36 | 4.37 | 3.26 | 3.53 | No | 533800 |
| SC176-14E | Ethiopia | Eastern Africa | mixed | 26.24 | 25.10 | 4.31 | 3.14 | 3.52 | No | No |
| SC180-14E | United States | Americas | mixed | 27.45 | 27.74 | 4.54 | 3.27 | 3.54 | No | No |
| SC182-14E | Ethiopia | Eastern Africa | mixed | 18.45 | 21.06 | 3.90 | 3.17 | 3.21 | No | No |
| SC183-14E | Unknown | 0 | mixed | 20.23 | 19.95 | 3.98 | 3.03 | 3.15 | No | No |
| SC184-14E | United States | Americas | mixed | 27.83 | 26.13 | 4.25 | 3.32 | 3.53 | No | No |
| SC185-14E | Nigeria | Western Africa | mixed | 28.40 | 26.55 | 4.52 | 3.14 | 3.57 | No | No |
| SC186-14E | Nigeria | Western Africa | mixed | NA | NA | NA | NA | NA | No | No |
| SC187-14E | United States | Americas | mixed | 38.28 | 32.63 | 4.96 | 3.37 | 3.68 | No | No |
| SC188-14E | Nigeria | Western Africa | mixed | NA | NA | NA | NA | NA | No | No |
| SC189-2 | NA | NA | Asian durras | 34.53 | 30.80 | 4.84 | 3.10 | 3.91 | No | No |
| SC19-14E | Ethiopia | Eastern Africa | E African durra | NA | NA | NA | NA | NA | No | No |
| SC190-6 | United States | Americas | mixed | 38.42 | 29.43 | 4.62 | 3.23 | 3.78 | No | No |
| SC191-14E | United States | Americas | mixed | 27.29 | 24.05 | 4.18 | 3.19 | 3.45 | No | No |
| SC192-14E | India | Asia | mixed | 23.60 | 22.93 | 4.02 | 3.34 | 3.27 | No | 576390 |

| | | | | | | | | | | |
|---|---|---|---|---|---|---|---|---|---|---|
| SC193-14E | NA | NA | mixed | 45.70 | 35.67 | 4.89 | 3.46 | 3.96 | No | No |
| SC195-14E | India | Asia | mixed | 23.54 | 22.53 | 4.20 | 2.99 | 3.39 | No | No |
| SC196-14E | United States | Americas | mixed | 30.28 | 25.78 | 4.39 | 3.15 | 3.53 | No | No |
| SC2-14E | Ethiopia | Eastern Africa | mixed | 20.12 | 22.98 | 4.12 | 3.19 | 3.32 | No | No |
| SC20-14E | Ethiopia | Eastern Africa | E African durra | 19.57 | 20.45 | 4.07 | 3.19 | 2.99 | No | No |
| SC200-14E | India | Asia | Asian durras | 25.21 | 23.66 | 4.24 | 3.17 | 3.36 | No | No |
| SC201-14E | United States | Americas | mixed | 26.80 | 26.99 | 4.57 | 3.21 | 3.54 | No | No |
| SC202-14E | India | Asia | Asian durras | NA | NA | NA | NA | NA | No | No |
| SC203-14E | India | Asia | Asian durras | 31.16 | 28.10 | 4.50 | 3.34 | 3.53 | No | No |
| SC206-14E | India | Asia | Asian durras | 25.35 | 24.21 | 4.11 | 3.32 | 3.37 | No | 533814 |
| SC207-14E | India | Asia | Asian durras | 24.27 | 25.45 | 4.30 | 3.31 | 3.41 | No | No |
| SC208-14E | India | Asia | Asian durras | 20.05 | 23.01 | 4.09 | 3.22 | 3.30 | No | No |
| SC209-14E | India | Asia | mixed | 31.54 | 30.22 | 4.60 | 3.25 | 3.83 | No | No |
| SC21-14E | Ethiopia | Eastern Africa | E African durra | 20.78 | 21.91 | 4.15 | 3.26 | 3.10 | No | 534127 |
| SC210-14E | India | Asia | Asian durras | 25.80 | 24.02 | 4.04 | 3.38 | 3.33 | No | No |
| SC211-14E | India | Asia | Asian durras | 25.47 | 23.64 | 4.36 | 3.08 | 3.34 | No | No |
| SC212-14E | India | Asia | Asian durras | 27.45 | 24.02 | 4.16 | 3.27 | 3.35 | No | No |
| SC213-14E | India | Asia | mixed | 17.73 | 18.35 | 4.02 | 3.07 | 2.81 | No | 576391 |
| SC214-14E | India | Asia | mixed | 22.93 | 22.51 | 3.98 | 3.27 | 3.28 | No | 533750 |
| SC215-14E | India | Asia | Asian durras | 25.75 | 21.19 | 3.86 | 3.28 | 3.18 | No | No |
| SC216-14E | India | Asia | mixed | 40.65 | 33.65 | 4.98 | 3.34 | 3.84 | No | No |
| SC217-14E | India | Asia | Asian durras | 23.11 | 23.57 | 4.28 | 3.15 | 3.32 | No | No |
| SC218-14E | India | Asia | mixed | 38.13 | 25.77 | 4.29 | 3.21 | 3.57 | No | No |
| SC22-14E | Ethiopia | Eastern Africa | mixed | 22.68 | 23.07 | 4.17 | 3.17 | 3.32 | No | 656091 |
| SC220-14E | India | Asia | Asian durras | 25.53 | 22.37 | 3.94 | 3.28 | 3.30 | No | No |
| SC221-14E | India | Asia | Asian durras | 30.05 | 24.58 | 4.14 | 3.26 | 3.47 | No | No |
| SC223-14E | United States | Americas | mixed | 23.26 | 23.05 | 4.15 | 3.08 | 3.40 | No | 533807 |
| SC224-14E | India | Asia | E African durra | 19.01 | 21.53 | 4.12 | 3.24 | 3.06 | No | 533927 |
| SC23-14E | NA | NA | mixed | 30.03 | 25.61 | 4.11 | 3.46 | 3.41 | Yes | 534128 |
| SC230-14E | NA | NA | mixed | 24.90 | 24.26 | 4.18 | 3.17 | 3.47 | No | No |
| SC231-14E | India | Asia | mixed | 32.54 | 25.28 | 4.31 | 3.22 | 3.46 | No | No |
| SC233-14E | India | Asia | Asian durras | 17.16 | 18.44 | 3.71 | 3.11 | 3.02 | No | No |
| SC235-14E | United States | Americas | mixed | 16.08 | 16.54 | 3.69 | 2.97 | 2.80 | No | No |
| SC236-3 | United States | Americas | mixed | 36.52 | 27.19 | 4.26 | 3.40 | 3.57 | No | No |
| SC24-14E | Ethiopia | Eastern Africa | mixed | 31.28 | 30.14 | 4.64 | 3.31 | 3.72 | No | No |
| SC240-14E | India | Asia | Asian durras | 15.54 | 18.09 | 3.95 | 2.89 | 2.93 | No | 533842 |
| SC241-14E | NA | NA | mixed | 21.55 | 21.24 | 4.03 | 3.08 | 3.24 | No | 533843 |
| SC242-14E | Nepal | Asia | mixed | 16.88 | 19.08 | 4.17 | 2.97 | 2.91 | No | No |
| SC243-14E | Nepal | Asia | mixed | 18.90 | 18.73 | 3.87 | 2.99 | 3.05 | No | 533845 |
| SC244-14E | India | Asia | mixed | 21.88 | 20.32 | 4.10 | 3.03 | 3.12 | No | No |
| SC245-14E | India | Asia | mixed | NA | NA | NA | NA | NA | No | No |
| SC247-14E | United States | Americas | mixed | 16.94 | 18.78 | 3.80 | 3.02 | 3.05 | No | No |
| SC248-14E | India | Asia | mixed | 19.98 | 21.81 | 4.09 | 3.09 | 3.26 | No | No |
| SC249-14E | United States | Americas | mixed | NA | NA | NA | NA | NA | No | No |
| SC25-14E | Ethiopia | Eastern Africa | mixed | 31.05 | 27.75 | 4.42 | 3.32 | 3.61 | No | 656092 |
| SC250-14E | India | Asia | mixed | 21.96 | 20.17 | 3.96 | 2.99 | 3.23 | No | No |
| SC252-14E | India | Asia | mixed | 17.25 | 18.25 | 3.84 | 2.96 | 3.00 | No | No |

| | | | | | | | | | | |
|---|---|---|---|---|---|---|---|---|---|---|
| SC254-14E | India | Asia | mixed | 14.87 | 18.62 | 3.81 | 3.02 | 3.00 | No | No |
| SC256-14E | Nigeria | Western Africa | Guinea | 25.77 | 26.79 | 5.07 | 2.93 | 3.43 | No | No |
| SC257-14E | Zambia | Eastern Africa | mixed | 28.04 | 23.84 | 4.34 | 3.04 | 3.43 | No | No |
| SC258-14E | Tanzania | Eastern Africa | mixed | 22.53 | 21.57 | 4.07 | 3.20 | 3.15 | No | No |
| SC259-14E | Sudan | Eastern Africa | Caudatum | 25.96 | 25.71 | 4.47 | 3.15 | 3.46 | No | No |
| SC260-6 | United States | Americas | mixed | NA | NA | NA | NA | NA | No | No |
| SC261-14E | Nigeria | Western Africa | mixed | 29.09 | 26.21 | 4.43 | 3.25 | 3.47 | No | 533841 |
| SC262-14E | Mali | Western Africa | mixed | 24.30 | 24.72 | 4.55 | 3.22 | 3.21 | No | No |
| SC265-14E | Burkina Faso | Western Africa | mixed | 23.61 | 22.04 | 4.00 | 3.14 | 3.31 | No | 533766 |
| SC266-6 | United States | Americas | mixed | 28.60 | 24.15 | 4.54 | 3.11 | 3.28 | No | No |
| SC267-14E | Sudan | Eastern Africa | mixed | 34.40 | 29.75 | 4.97 | 3.18 | 3.61 | No | No |
| SC268-14E | Sudan | Eastern Africa | mixed | 29.94 | 25.93 | 4.49 | 3.19 | 3.45 | No | No |
| SC269-14E | Nigeria | Western Africa | mixed | 26.82 | 24.96 | 4.44 | 3.13 | 3.42 | No | No |
| SC27-14E | United States | Americas | mixed | 28.65 | 28.11 | 4.56 | 3.24 | 3.64 | No | No |
| SC270-14E | Nigeria | Western Africa | mixed | 36.61 | 25.87 | 4.52 | 3.15 | 3.45 | No | No |
| SC271-14E | Nigeria | Western Africa | Guinea | NA | NA | NA | NA | NA | No | No |
| SC272-14E | Nigeria | Western Africa | Guinea | 24.21 | 26.95 | 4.68 | 3.04 | 3.60 | No | No |
| SC273-14E | Nigeria | Western Africa | mixed | 31.95 | 24.58 | 4.49 | 3.00 | 3.46 | No | No |
| SC276-14E | United States | Americas | mixed | 35.83 | 28.58 | 4.64 | 3.16 | 3.67 | No | No |
| SC277-14E | Nigeria | Western Africa | Guinea | 40.26 | 32.34 | 4.91 | 3.24 | 3.85 | No | No |
| SC278-14E | India | Asia | mixed | 28.17 | 27.99 | 4.72 | 3.07 | 3.67 | No | No |
| SC279-14E | Nigeria | Western Africa | Guinea | 27.91 | 28.98 | 5.19 | 3.07 | 3.49 | No | 534070 |
| SC28-14E | Ethiopia | Eastern Africa | mixed | 31.13 | 28.88 | 4.49 | 3.35 | 3.65 | No | No |
| SC280-14E | Nigeria | Western Africa | mixed | 32.26 | 27.36 | 4.49 | 3.18 | 3.63 | No | No |
| SC281-14E | Nigeria | Western Africa | mixed | 26.15 | 24.80 | 4.46 | 3.20 | 3.31 | No | No |
| SC283-14E | Tanzania | Eastern Africa | mixed | 25.27 | 24.31 | 4.55 | 3.10 | 3.27 | No | 533869 |
| SC284-14E | Burkina Faso | Western Africa | mixed | 28.17 | 23.84 | 4.23 | 3.07 | 3.48 | No | No |
| SC285-14E | Chad | Western Africa | Guinea | 31.20 | 27.44 | 4.63 | 2.92 | 3.86 | No | No |
| SC287-14E | United States | Americas | Guinea | NA | NA | NA | NA | NA | No | No |
| SC289-14E | Nigeria | Western Africa | Guinea | 32.38 | 31.96 | 5.06 | 3.12 | 3.82 | No | No |
| SC29-14E | Ethiopia | Eastern Africa | mixed | 33.41 | 31.08 | 4.73 | 3.37 | 3.69 | No | No |
| SC290-14E | Nigeria | Western Africa | Guinea | 28.58 | 24.86 | 4.45 | 3.12 | 3.42 | No | No |
| SC291-14E | Nigeria | Western Africa | Guinea | 37.64 | 30.80 | 4.88 | 3.23 | 3.73 | No | No |
| SC293-14E | Nigeria | Western Africa | Guinea | 37.32 | 27.18 | 5.04 | 3.08 | 3.36 | No | No |
| SC295-11E | United States | Americas | Guinea | 25.48 | 23.96 | 4.61 | 3.12 | 3.26 | No | 656093 |
| SC296-14E | Nigeria | Western Africa | mixed | 22.90 | 25.54 | 4.48 | 3.17 | 3.43 | No | No |
| SC297-14E | United States | Americas | Guinea | 24.62 | 21.41 | 4.04 | 3.24 | 3.13 | No | No |
| SC299-14E | Nigeria | Western Africa | Guinea | NA | NA | NA | NA | NA | No | 533785 |
| SC3-3 | United States | Americas | mixed | 20.52 | 22.50 | 4.04 | 3.15 | 3.32 | No | No |
| SC30-14E | Ethiopia | Eastern Africa | mixed | 15.59 | 18.32 | 3.96 | 3.04 | 2.86 | No | No |
| SC300-14E | Nigeria | Western Africa | mixed | NA | NA | NA | NA | NA | No | No |
| SC301-14E | United States | Americas | mixed | 31.33 | 26.16 | 4.51 | 3.13 | 3.53 | No | 656094 |
| SC305-14E | Chad | Western Africa | mixed | 23.59 | 26.15 | 4.63 | 3.31 | 3.25 | No | 534037 |
| SC306-14E | NA | NA | E African durra | 22.43 | 24.73 | 4.30 | 3.29 | 3.33 | No | No |
| SC307-14E | India | Asia | Asian durras | 21.69 | 21.43 | 4.03 | 3.15 | 3.22 | No | No |
| SC311-14E | 0 | 0 | mixed | 27.36 | 26.09 | 4.64 | 3.15 | 3.40 | No | No |
| SC314-6 | United States | Americas | mixed | 33.06 | 29.94 | 4.91 | 3.17 | 3.63 | No | No |

| | | | | | | | | | | |
|---|---|---|---|---|---|---|---|---|---|---|
| SC319-14E | Uganda | Eastern Africa | mixed | 21.34 | 24.41 | 4.37 | 3.23 | 3.29 | No | 533833 |
| SC320-14E | Chad | Western Africa | mixed | 26.45 | 24.56 | 4.19 | 3.26 | 3.43 | No | 533863 |
| SC323-14E | Sudan | Eastern Africa | mixed | 23.94 | 25.43 | 4.51 | 3.00 | 3.57 | No | 576399 |
| SC324-14 | United States | Americas | mixed | 30.97 | 27.59 | 4.52 | 3.30 | 3.56 | No | No |
| SC325-14E | NA | NA | mixed | 23.26 | 24.64 | 4.44 | 3.11 | 3.40 | No | 533957 |
| SC326-6 | NA | NA | mixed | 23.56 | 23.66 | 4.15 | 3.22 | 3.36 | No | No |
| SC328-14E | India | Asia | Caudatum | 24.64 | 23.88 | 4.33 | 3.16 | 3.33 | No | 534112 |
| SC329-14E | Nigeria | Western Africa | mixed | NA | NA | NA | NA | NA | No | 533838 |
| SC330-14E | India | Asia | mixed | 23.75 | 21.75 | 4.04 | 3.05 | 3.36 | No | No |
| SC331-14E | Nigeria | Western Africa | mixed | 31.83 | 28.72 | 4.83 | 3.26 | 3.49 | No | 533824 |
| SC333-14E | Ethiopia | Eastern Africa | mixed | 46.51 | 39.39 | 5.15 | 3.78 | 3.81 | No | 533761 |
| SC334-14E | NA | NA | mixed | 37.83 | 31.44 | 4.71 | 3.46 | 3.67 | No | 533986 |
| SC335-14E | Sudan | Eastern Africa | mixed | 35.49 | 32.32 | 4.72 | 3.35 | 3.87 | No | No |
| SC336-14E | Sudan | Eastern Africa | mixed | NA | NA | NA | NA | NA | No | No |
| SC338-14E | Sudan | Eastern Africa | mixed | 31.77 | 32.50 | 4.96 | 3.11 | 3.96 | No | No |
| SC339-14E | Sudan | Eastern Africa | mixed | 43.13 | 36.30 | 5.00 | 3.48 | 3.99 | No | No |
| SC340-14E | Sudan | Eastern Africa | mixed | 28.19 | 24.91 | 4.48 | 3.07 | 3.43 | No | No |
| SC342-14E | Nigeria | Western Africa | mixed | 38.66 | 26.70 | 4.52 | 3.04 | 3.70 | No | No |
| SC343-9 | United States | Americas | mixed | 28.58 | 24.08 | 4.09 | 3.29 | 3.41 | No | No |
| SC344-14E | Nigeria | Western Africa | mixed | 22.42 | 20.83 | 4.15 | 3.00 | 3.19 | No | No |
| SC345-14E | Nigeria | Western Africa | mixed | 39.20 | 33.53 | 4.86 | 3.26 | 3.99 | No | No |
| SC346-14E | Nigeria | Western Africa | mixed | 18.32 | 23.23 | 4.23 | 3.05 | 3.42 | No | No |
| SC347-14E | Nigeria | Western Africa | mixed | 32.28 | 28.25 | 4.56 | 3.26 | 3.63 | No | No |
| SC348-14E | Nigeria | Western Africa | mixed | NA | NA | NA | NA | NA | No | 534075 |
| SC349-14E | United States | Americas | mixed | NA | NA | NA | NA | NA | No | No |
| SC35-14E | NA | NA | mixed | 34.52 | 30.03 | 4.51 | 3.41 | 3.72 | Yes | 534133 |
| SC350-14E | Nigeria | Western Africa | Caudatum | 25.20 | 24.04 | 4.43 | 3.02 | 3.42 | No | No |
| SC351-14E | Nigeria | Western Africa | mixed | 28.87 | 22.74 | 4.04 | 3.18 | 3.37 | No | No |
| SC352-14E | Sudan | Eastern Africa | mixed | 39.15 | 41.60 | 5.39 | 3.40 | 4.26 | No | No |
| SC353-14E | Nigeria | Western Africa | mixed | 37.51 | 29.30 | 4.61 | 3.39 | 3.54 | No | No |
| SC354-14E | Nigeria | Western Africa | mixed | 32.71 | 28.86 | 4.66 | 3.18 | 3.69 | No | No |
| SC356-14E | Nigeria | Western Africa | mixed | 37.25 | 34.37 | 5.21 | 3.18 | 3.89 | No | No |
| SC358-14E | Nigeria | Western Africa | mixed | 44.86 | 33.67 | 4.62 | 3.63 | 3.78 | No | No |
| SC36-14E | NA | NA | mixed | 28.27 | 21.56 | 3.95 | 3.24 | 3.20 | No | No |
| SC362-14E | Nigeria | Western Africa | mixed | 31.59 | 30.91 | 4.70 | 3.37 | 3.68 | No | No |
| SC366-14E | Nigeria | Western Africa | mixed | 34.86 | 33.97 | 5.12 | 3.28 | 3.84 | No | No |
| SC367-14E | Nigeria | Western Africa | mixed | 50.10 | 36.62 | 4.79 | 3.66 | 3.94 | No | No |
| SC368-14E | Nigeria | Western Africa | mixed | 54.39 | 44.26 | 5.12 | 3.83 | 4.20 | No | No |
| SC369-14E | Nigeria | Western Africa | mixed | 36.92 | 35.09 | 4.91 | 3.42 | 3.96 | No | No |
| SC37-14E | Ethiopia | Eastern Africa | mixed | 21.83 | 22.95 | 4.21 | 3.16 | 3.27 | No | No |
| SC370-14E | Nigeria | Western Africa | mixed | NA | NA | NA | NA | NA | No | 533776 |
| SC371-14E | United States | Americas | mixed | 36.65 | 33.51 | 4.79 | 3.43 | 3.86 | No | No |
| SC372-14E | NA | NA | mixed | 42.92 | 29.97 | 4.64 | 3.60 | 3.44 | No | 533878 |
| SC373-14E | NA | NA | mixed | 43.08 | 38.20 | 5.31 | 3.35 | 4.04 | No | 656095 |
| SC374-14E | Nigeria | Western Africa | Guinea | 33.26 | 29.17 | 4.73 | 3.27 | 3.58 | No | No |
| SC377-14E | Nigeria | Western Africa | Guinea | 31.18 | 27.74 | 4.37 | 3.43 | 3.53 | No | No |
| SC38-14E | Ethiopia | Eastern Africa | mixed | 29.38 | 24.92 | 4.12 | 3.33 | 3.46 | No | 534135 |

| | | | | | | | | | | |
|---|---|---|---|---|---|---|---|---|---|---|
| SC380-14E | Nigeria | Western Africa | mixed | 42.82 | 31.57 | 4.81 | 3.28 | 3.78 | No | No |
| SC386-14E | NA | NA | mixed | 43.97 | 30.47 | 4.47 | 3.48 | 3.73 | No | 656119 |
| SC387-14E | Nigeria | Western Africa | mixed | 32.75 | 32.17 | 4.85 | 3.44 | 3.67 | No | No |
| SC389-14E | Nigeria | Western Africa | Guinea | NA | NA | NA | NA | NA | No | No |
| SC391-14E | NA | NA | mixed | 39.33 | 29.61 | 4.53 | 3.42 | 3.63 | No | 656096 |
| SC392-14E | Nigeria | Western Africa | mixed | 28.38 | 27.01 | 4.37 | 3.25 | 3.61 | No | No |
| SC396-14E | Nigeria | Western Africa | mixed | NA | NA | NA | NA | NA | No | 533877 |
| SC397-14E | Nigeria | Western Africa | mixed | 25.47 | 26.44 | 4.39 | 3.38 | 3.41 | No | No |
| SC398-14E | Nigeria | Western Africa | mixed | 23.97 | 25.83 | 4.60 | 3.20 | 3.34 | No | No |
| SC399-14E | NA | NA | mixed | 42.48 | 31.29 | 4.58 | 3.43 | 3.76 | No | 533882 |
| SC402-14E | Nigeria | Western Africa | mixed | NA | NA | NA | NA | NA | No | No |
| SC403-14E | Nigeria | Western Africa | Guinea | 29.80 | 25.33 | 4.55 | 3.13 | 3.36 | No | No |
| SC405-14E | Sudan | Eastern Africa | mixed | 37.57 | 31.93 | 4.77 | 3.26 | 3.93 | No | No |
| SC406-14E | United States | Americas | mixed | NA | NA | NA | NA | NA | No | No |
| SC407-14E | Nigeria | Western Africa | mixed | 32.16 | 31.88 | 4.81 | 3.34 | 3.75 | No | No |
| SC408-14E | Nigeria | Western Africa | mixed | 24.91 | 25.74 | 4.50 | 3.24 | 3.39 | No | No |
| SC409-14E | United States | Americas | mixed | NA | NA | NA | NA | NA | No | No |
| SC41-5 | United States | Americas | mixed | 35.40 | 33.19 | 5.01 | 3.32 | 3.76 | No | No |
| SC410-14E | 0 | 0 | Asian durras | NA | NA | NA | NA | NA | No | No |
| SC410-6 | 0 | 0 | mixed | 30.48 | 22.28 | 3.88 | 3.38 | 3.24 | No | No |
| SC411-14E | Sudan | Eastern Africa | mixed | 28.10 | 27.12 | 4.49 | 3.16 | 3.64 | No | 533866 |
| SC417-14E | Senegal | Western Africa | mixed | 25.15 | 24.17 | 4.35 | 3.09 | 3.41 | No | No |
| SC418-14E | Tanzania | Eastern Africa | mixed | 34.26 | 24.23 | 4.36 | 3.00 | 3.50 | No | 533822 |
| SC42-14E | Ethiopia | Eastern Africa | mixed | 23.96 | 28.49 | 4.70 | 3.25 | 3.54 | No | 576393 |
| SC420-14E | Sudan | Eastern Africa | mixed | 16.72 | 21.47 | 4.16 | 3.00 | 3.24 | No | 533769 |
| SC422-14E | India | Asia | mixed | 21.33 | 24.35 | 4.26 | 3.19 | 3.40 | No | No |
| SC423-14 | United States | Americas | Caudatum | 27.42 | 24.44 | 4.27 | 3.20 | 3.41 | No | 533758 |
| SC424-14E | NA | NA | mixed | 23.49 | 23.56 | 4.35 | 3.35 | 3.09 | No | 533901 |
| SC425-14E | NA | NA | mixed | 45.19 | 30.04 | 4.37 | 3.57 | 3.66 | No | 533762 |
| SC43-14E | Ethiopia | Eastern Africa | E African durra | 22.61 | 23.33 | 4.14 | 3.20 | 3.33 | No | No |
| SC430-14E | Ethiopia | Eastern Africa | mixed | 31.03 | 25.13 | 4.24 | 3.34 | 3.44 | No | No |
| SC435-14E | Tanzania | Eastern Africa | Kafir | 16.59 | 19.60 | 3.88 | 3.22 | 2.99 | No | No |
| SC436-9 | United States | Americas | mixed | 29.78 | 24.02 | 4.15 | 3.19 | 3.45 | No | No |
| SC437-14E | India | Asia | mixed | 27.81 | 22.46 | 4.09 | 3.23 | 3.24 | No | No |
| SC438-3 | United States | Americas | mixed | 25.53 | 27.03 | 4.45 | 3.30 | 3.50 | No | No |
| SC44-14E | Ethiopia | Eastern Africa | E African durra | 18.69 | 20.37 | 4.08 | 3.18 | 2.99 | No | No |
| SC441-14E | India | Asia | mixed | 25.50 | 22.48 | 3.98 | 3.23 | 3.32 | No | 534009 |
| SC442-14E | India | Asia | mixed | 20.89 | 19.92 | 3.83 | 3.15 | 3.14 | No | No |
| SC445-14E | United States | Americas | mixed | 25.17 | 24.92 | 4.40 | 3.12 | 3.46 | No | No |
| SC449-14E | United States | Americas | mixed | 21.71 | 20.89 | 3.97 | 3.17 | 3.15 | No | 597950 |
| SC450-14E | India | Asia | mixed | 16.28 | 18.01 | 3.64 | 3.05 | 3.01 | No | 533852 |
| SC451-14E | India | Asia | mixed | 19.67 | 21.93 | 3.94 | 3.30 | 3.21 | No | No |
| SC452-14E | India | Asia | Asian durras | 18.46 | 21.56 | 3.95 | 3.17 | 3.25 | No | No |
| SC454-14E | India | Asia | mixed | 29.00 | 24.44 | 4.14 | 3.31 | 3.40 | No | No |
| SC456-6 | United States | Americas | mixed | 27.11 | 26.39 | 4.34 | 3.30 | 3.51 | No | No |
| SC457-14E | India | Asia | mixed | 17.04 | 19.55 | 3.93 | 3.19 | 2.95 | No | No |
| SC458-12E | United States | Americas | mixed | 29.56 | 23.09 | 4.07 | 3.25 | 3.33 | No | No |

| | | | | | | | | | | |
|---|---|---|---|---|---|---|---|---|---|---|
| SC460-14E | India | Asia | mixed | 30.73 | 26.41 | 4.47 | 3.04 | 3.66 | No | No |
| SC462-14E | India | Asia | Asian durras | 23.88 | 23.20 | 4.24 | 3.00 | 3.45 | No | No |
| SC464-14E | India | Asia | mixed | 18.86 | 20.20 | 3.85 | 3.24 | 3.07 | No | No |
| SC465-14E | India | Asia | mixed | 31.83 | 29.29 | 4.96 | 3.03 | 3.69 | No | 533997 |
| SC466-14E | India | Asia | mixed | 25.79 | 26.68 | 4.49 | 3.24 | 3.50 | No | No |
| SC467-14E | India | Asia | mixed | 26.83 | 24.10 | 4.24 | 3.20 | 3.37 | No | 533943 |
| SC468-14E | India | Asia | mixed | 24.84 | 21.70 | 3.97 | 3.19 | 3.27 | No | No |
| SC469-14E | India | Asia | Asian durras | 29.36 | 26.54 | 4.65 | 3.09 | 3.50 | No | No |
| SC470-6 | United States | Americas | mixed | 27.83 | 21.92 | 3.92 | 3.22 | 3.31 | No | No |
| SC471-14E | India | Asia | Asian durras | 23.48 | 22.22 | 4.05 | 3.13 | 3.33 | No | No |
| SC472-14E | India | Asia | Asian durras | 24.84 | 23.24 | 4.38 | 3.03 | 3.34 | No | No |
| SC473-14E | NA | NA | mixed | 24.19 | 24.24 | 4.47 | 3.13 | 3.32 | No | 534028 |
| SC475-14E | India | Asia | Asian durras | 24.08 | 19.19 | 3.82 | 3.25 | 3.01 | No | No |
| SC477-14E | India | Asia | mixed | 24.01 | 23.24 | 4.14 | 3.18 | 3.34 | No | No |
| SC479-14E | United States | Americas | Asian durras | 35.14 | 28.44 | 4.49 | 3.43 | 3.52 | No | No |
| SC48-14E | Sudan | Eastern Africa | mixed | 40.75 | 30.95 | 4.83 | 3.28 | 3.74 | No | No |
| SC480-14E | United States | Americas | Asian durras | 31.08 | 28.65 | 4.47 | 3.36 | 3.65 | No | 656097 |
| SC482-14E | India | Asia | Asian durras | 29.97 | 23.83 | 4.31 | 3.09 | 3.40 | No | No |
| SC483-14E | India | Asia | Asian durras | 23.29 | 22.95 | 4.33 | 3.18 | 3.15 | No | No |
| SC484-14E | India | Asia | mixed | 26.09 | 24.24 | 4.40 | 3.22 | 3.27 | No | No |
| SC485-14E | United States | Americas | Asian durras | 18.10 | 23.35 | 4.27 | 3.13 | 3.31 | No | No |
| SC489-14E | India | Asia | Asian durras | 25.23 | 23.32 | 4.28 | 3.26 | 3.20 | No | 533856 |
| SC490-14E | India | Asia | mixed | 23.03 | 26.15 | 4.56 | 3.26 | 3.37 | No | No |
| SC492-14E | India | Asia | Asian durras | 27.21 | 24.70 | 4.39 | 3.22 | 3.33 | No | No |
| SC493-14E | India | Asia | Kafir | 28.70 | 23.19 | 3.96 | 3.30 | 3.36 | No | No |
| SC494-14E | India | Asia | Asian durras | 25.32 | 25.46 | 4.52 | 3.24 | 3.32 | No | No |
| SC497-14E | India | Asia | Asian durras | 27.92 | 26.47 | 4.41 | 3.21 | 3.56 | No | No |
| SC498-14E | United States | Americas | Asian durras | 26.04 | 25.44 | 4.42 | 3.16 | 3.44 | No | 656099 |
| SC499-14E | India | Asia | Asian durras | 26.88 | 25.56 | 4.30 | 3.41 | 3.30 | No | No |
| SC500-9 | United States | Americas | Asian durras | 25.08 | 23.16 | 4.15 | 3.18 | 3.34 | No | 656100 |
| SC501-14E | India | Asia | mixed | 31.32 | 32.86 | 4.88 | 3.33 | 3.81 | No | No |
| SC502-14E | Sudan | Eastern Africa | mixed | 36.43 | 29.48 | 4.50 | 3.38 | 3.67 | No | 533996 |
| SC505-14E | United States | Americas | Caudatum | 30.07 | 23.66 | 4.13 | 3.22 | 3.37 | No | No |
| SC51-14E | NA | NA | mixed | 31.94 | 27.68 | 4.25 | 3.48 | 3.57 | No | 534137 |
| SC512-14E | India | Asia | mixed | 20.38 | 23.42 | 4.26 | 3.20 | 3.26 | No | No |
| SC514-14E | India | Asia | mixed | 26.84 | 24.90 | 4.43 | 3.04 | 3.52 | No | No |
| SC515-14E | India | Asia | Guinea | 29.77 | 27.84 | 4.77 | 3.08 | 3.59 | No | No |
| SC516-14E | India | Asia | mixed | 21.13 | 25.76 | 4.61 | 3.21 | 3.33 | No | No |
| SC517-9 | United States | Americas | mixed | 28.62 | 25.49 | 4.49 | 3.17 | 3.42 | No | No |
| SC519-14E | Nigeria | Western Africa | Guinea | 24.69 | 24.56 | 4.38 | 3.07 | 3.47 | No | No |
| SC520-14E | Nigeria | Western Africa | Guinea | NA | NA | NA | NA | NA | No | No |
| SC521-6 | United States | Americas | mixed | NA | NA | NA | NA | NA | No | No |
| SC523-14E | Nigeria | Western Africa | Guinea | 29.94 | 27.51 | 4.49 | 3.29 | 3.54 | No | No |
| SC525-9 | United States | Americas | mixed | 38.96 | 34.55 | 5.17 | 3.35 | 3.79 | No | 656101 |
| SC526-14E | Nigeria | Western Africa | Guinea | 37.90 | 32.06 | 5.04 | 3.17 | 3.82 | No | No |
| SC527-14E | Nigeria | Western Africa | mixed | 39.07 | 33.64 | 5.01 | 3.23 | 3.95 | No | No |
| SC529-14E | Nigeria | Western Africa | Guinea | 44.89 | 34.29 | 5.05 | 3.24 | 3.93 | No | No |

| | | | | | | | | | | |
|---|---|---|---|---|---|---|---|---|---|---|
| SC532-14E | United States | Americas | Guinea | 27.71 | 25.62 | 4.52 | 3.16 | 3.43 | No | 597951 |
| SC536-14E | Nigeria | Western Africa | Guinea | 33.05 | 29.76 | 4.87 | 3.15 | 3.68 | No | No |
| SC537-14E | United States | Americas | mixed | 28.10 | 26.59 | 4.82 | 3.08 | 3.42 | No | No |
| SC538-14E | Nigeria | Western Africa | mixed | NA | NA | NA | NA | NA | No | No |
| SC54-14E | Sudan | Eastern Africa | mixed | 33.02 | 26.87 | 4.82 | 3.02 | 3.50 | No | No |
| SC540-6 | United States | Americas | mixed | 24.31 | 25.03 | 4.38 | 3.12 | 3.51 | No | No |
| SC544-14E | Nigeria | Western Africa | Guinea | 31.62 | 26.74 | 4.54 | 3.23 | 3.50 | No | No |
| SC545-14E | Nigeria | Western Africa | Guinea | 24.87 | 24.28 | 4.46 | 3.06 | 3.36 | No | No |
| SC546-14E | Nigeria | Western Africa | mixed | 30.31 | 30.89 | 5.07 | 3.21 | 3.64 | No | No |
| SC547-6 | United States | Americas | mixed | 31.43 | 27.13 | 4.45 | 3.14 | 3.65 | No | No |
| SC55-14E | United States | Americas | mixed | 38.35 | 30.99 | 4.49 | 3.50 | 3.74 | No | 533755 |
| SC550-14E | United States | Americas | mixed | NA | NA | NA | NA | NA | No | No |
| SC553-14E | Nigeria | Western Africa | Guinea | NA | NA | NA | NA | NA | No | 534063 |
| SC556-3 | United States | Americas | Kafir | 23.80 | 25.41 | 4.27 | 3.25 | 3.48 | No | No |
| SC557-14E | Ethiopia | Eastern Africa | mixed | 14.22 | 16.50 | 3.59 | 2.89 | 2.95 | No | 533939 |
| SC558-14E | NA | NA | mixed | 23.81 | 25.30 | 4.29 | 3.25 | 3.46 | No | 533938 |
| SC56-14E | NA | NA | mixed | 33.66 | 26.30 | 4.53 | 3.15 | 3.51 | Yes | 533910 |
| SC562-14E | Sudan | Eastern Africa | mixed | 37.65 | 30.55 | 4.61 | 3.46 | 3.64 | No | 533987 |
| SC563-14E | Nigeria | Western Africa | mixed | 34.05 | 28.34 | 4.45 | 3.37 | 3.60 | No | 533876 |
| SC565-14E | Niger | Western Africa | mixed | 30.79 | 27.60 | 4.60 | 3.18 | 3.59 | No | No |
| SC566-14E | Nigeria | Western Africa | mixed | 30.58 | 25.32 | 4.29 | 3.28 | 3.43 | No | 533871 |
| SC567-14E | Nigeria | Western Africa | mixed | 31.91 | 29.21 | 4.92 | 3.11 | 3.62 | No | No |
| SC57-14E | NA | NA | Caudatum | 28.49 | 26.17 | 4.36 | 3.16 | 3.60 | No | 533789 |
| SC572-14E | NA | NA | mixed | 26.41 | 22.57 | 4.22 | 3.08 | 3.29 | No | 533980 |
| SC574-14E | Pakistan | Asia | mixed | 21.00 | 24.08 | 4.28 | 3.25 | 3.31 | No | 534114 |
| SC575-14E | Sudan | Eastern Africa | mixed | 44.68 | 37.05 | 4.94 | 3.59 | 3.95 | No | No |
| SC578-14E | Nigeria | Western Africa | mixed | 26.10 | 23.35 | 4.19 | 3.25 | 3.31 | No | No |
| SC58-14E | Sudan | Eastern Africa | mixed | 30.56 | 26.53 | 4.31 | 3.30 | 3.55 | No | 533911 |
| SC580-14E | India | Asia | Asian durras | 22.13 | 25.50 | 4.44 | 3.20 | 3.45 | No | No |
| SC582-14E | United States | Americas | mixed | 15.68 | 19.53 | 4.03 | 2.98 | 3.03 | No | No |
| SC586-14E | Nigeria | Western Africa | mixed | 27.03 | 26.97 | 4.80 | 3.15 | 3.38 | No | No |
| SC589-14E | India | Asia | Asian durras | 21.06 | 22.06 | 4.15 | 3.11 | 3.25 | No | No |
| SC59-12E | United States | Americas | mixed | 19.62 | 22.34 | 3.98 | 3.21 | 3.27 | No | 656102 |
| SC590-14E | India | Asia | mixed | 20.98 | 22.38 | 3.99 | 3.25 | 3.29 | No | No |
| SC593-14E | Ethiopia | Eastern Africa | mixed | 28.16 | 27.92 | 4.82 | 3.13 | 3.53 | No | No |
| SC598-14E | Uganda | Eastern Africa | mixed | 22.84 | 23.76 | 4.12 | 3.21 | 3.40 | No | No |
| SC599-11E | United States | Americas | mixed | 27.65 | 22.33 | 4.18 | 3.06 | 3.30 | No | 534163 |
| SC6-14E | NA | NA | mixed | 22.50 | 25.18 | 4.47 | 3.13 | 3.42 | No | 533902 |
| SC60-14E | Sudan | Eastern Africa | Caudatum | 25.03 | 23.81 | 4.06 | 3.30 | 3.38 | No | 533962 |
| SC600-14E | Sudan | Eastern Africa | mixed | 32.25 | 28.54 | 4.51 | 3.34 | 3.60 | No | No |
| SC601-14E | India | Asia | mixed | 19.38 | 22.66 | 4.14 | 3.13 | 3.32 | No | No |
| SC602-6 | 0 | 0 | mixed | 16.71 | 19.63 | 3.98 | 3.16 | 2.98 | No | No |
| SC603-14E | NA | NA | Kafir | 18.85 | 22.63 | 4.08 | 3.26 | 3.23 | No | 533936 |
| SC604-6 | United States | Americas | mixed | 24.47 | 23.75 | 4.22 | 3.16 | 3.37 | No | No |
| SC605-14E | Mali | Western Africa | mixed | 21.07 | 24.84 | 4.60 | 3.26 | 3.19 | No | 534096 |
| SC606-14E | United States | Americas | mixed | 19.63 | 22.60 | 4.16 | 3.16 | 3.28 | No | 597946 |
| SC609-14E | China | Asia | mixed | 33.22 | 28.84 | 4.50 | 3.30 | 3.71 | No | 576332 |

| | | | | | | | | | | |
|---|---|---|---|---|---|---|---|---|---|---|
| SC61-14E | Sudan | Eastern Africa | mixed | 30.64 | 29.01 | 5.05 | 3.20 | 3.42 | No | No |
| SC614-14E | Tanzania | Eastern Africa | mixed | 23.39 | 25.04 | 4.35 | 3.19 | 3.43 | No | 533940 |
| SC615-9 | United States | Americas | mixed | 17.08 | 18.83 | 3.91 | 3.14 | 2.89 | No | No |
| SC618-6 | United States | Americas | mixed | 19.55 | 20.82 | 4.13 | 2.93 | 3.22 | No | No |
| SC62-14E | NA | NA | mixed | 25.19 | 25.18 | 4.68 | 3.16 | 3.27 | Yes | 534138 |
| SC620-14E | South Africa | Southern Africa | mixed | 19.29 | 18.16 | 3.60 | 3.08 | 3.07 | No | No |
| SC621-14E | United States | Americas | mixed | 18.39 | 21.06 | 4.06 | 3.26 | 3.01 | No | 656104 |
| SC623-14E | Sudan | Eastern Africa | mixed | 17.25 | 23.90 | 4.73 | 3.19 | 3.02 | No | 533956 |
| SC624-14E | India | Asia | mixed | NA | NA | NA | NA | NA | No | 576366 |
| SC625-14E | Japan | Asia | mixed | 17.96 | 19.25 | 3.95 | 3.13 | 2.94 | No | 534097 |
| SC626-14E | Japan | Asia | mixed | 22.30 | 22.55 | 4.06 | 3.32 | 3.18 | No | No |
| SC627-14E | South Africa | Southern Africa | mixed | 17.30 | 18.34 | 3.67 | 3.18 | 2.96 | No | 576345 |
| SC628-14E | NA | NA | Kafir | 26.64 | 23.16 | 4.19 | 3.15 | 3.37 | No | 533979 |
| SC629-14E | United States | Americas | Kafir | 33.23 | 23.87 | 4.05 | 3.23 | 3.45 | No | No |
| SC63-14E | Sudan | Eastern Africa | mixed | 40.04 | 33.08 | 4.72 | 3.44 | 3.89 | No | 533912 |
| SC630-14E | NA | NA | Kafir | 26.34 | 27.00 | 4.74 | 3.31 | 3.45 | No | 533937 |
| SC631-6 | United States | Americas | Kafir | 32.61 | 26.78 | 4.32 | 3.31 | 3.58 | No | No |
| SC632-14 | United States | Americas | Kafir | 31.81 | 25.63 | 4.43 | 3.20 | 3.45 | No | No |
| SC634-14E | NA | NA | mixed | 20.25 | 23.83 | 4.44 | 3.21 | 3.20 | No | No |
| SC635-14E | United States | Americas | mixed | 31.16 | 25.94 | 4.38 | 3.22 | 3.50 | No | No |
| SC636-6 | NA | NA | mixed | 30.69 | 28.22 | 4.55 | 3.25 | 3.64 | No | No |
| SC637-14E | India | Asia | mixed | 26.79 | 28.17 | 4.56 | 3.17 | 3.69 | No | 534105 |
| SC639-6 | United States | Americas | mixed | 21.33 | 23.80 | 4.22 | 3.27 | 3.27 | No | 656105 |
| SC64-14E | Sudan | Eastern Africa | mixed | 27.24 | 25.99 | 4.40 | 3.16 | 3.55 | No | 533757 |
| SC641-14E | India | Asia | mixed | 29.21 | 27.48 | 4.49 | 3.30 | 3.53 | No | 534104 |
| SC642-14E | India | Asia | mixed | 23.20 | 24.06 | 4.14 | 3.22 | 3.44 | No | No |
| SC643-14E | NA | NA | mixed | 28.38 | 25.22 | 4.33 | 3.33 | 3.33 | No | No |
| SC644-14E | NA | NA | mixed | 25.02 | 25.48 | 4.45 | 3.14 | 3.47 | No | No |
| SC645-14E | India | Asia | mixed | 24.85 | 25.26 | 4.45 | 3.28 | 3.31 | No | 534108 |
| SC646-14E | India | Asia | Kafir | 27.10 | 27.48 | 4.59 | 3.25 | 3.50 | No | No |
| SC647-14E | NA | NA | Kafir | 30.39 | 27.55 | 4.48 | 3.33 | 3.51 | No | No |
| SC648-14E | NA | NA | Kafir | 20.62 | 24.23 | 4.61 | 3.15 | 3.20 | No | 533955 |
| SC649-14E | Zimbabwe | Eastern Africa | mixed | 30.79 | 23.62 | 4.14 | 3.28 | 3.31 | No | No |
| SC650-14E | NA | NA | Asian durras | 31.00 | 27.24 | 4.43 | 3.14 | 3.72 | No | 576340 |
| SC652-9 | United States | Americas | Kafir | 27.47 | 27.49 | 4.46 | 3.27 | 3.58 | No | No |
| SC653-14E | NA | NA | Kafir | 21.81 | 23.29 | 4.38 | 3.19 | 3.19 | No | No |
| SC654-14E | Zimbabwe | Eastern Africa | Kafir | 24.38 | 27.48 | 4.74 | 3.21 | 3.46 | No | No |
| SC655-14E | South Africa | Southern Africa | mixed | 30.29 | 26.87 | 4.39 | 3.30 | 3.53 | No | 533976 |
| SC657-14E | NA | NA | Kafir | 32.50 | 27.58 | 4.42 | 3.37 | 3.51 | No | No |
| SC659-14E | United States | Americas | mixed | 30.14 | 26.71 | 4.49 | 3.20 | 3.54 | No | 576333 |
| SC66-14E | NA | NA | mixed | 21.47 | 22.51 | 4.35 | 3.10 | 3.18 | No | 533913 |
| SC663-14E | United States | Americas | Kafir | 32.29 | 29.25 | 4.99 | 3.18 | 3.54 | No | 533948 |
| SC67-14E | Sudan | Eastern Africa | mixed | 19.15 | 19.68 | 4.14 | 3.15 | 2.89 | No | 534139 |
| SC671-14E | Kenya | Eastern Africa | mixed | 26.19 | 21.34 | 3.77 | 3.37 | 3.19 | No | 534054 |
| SC672-14E | United States | Americas | Kafir | 17.22 | 21.05 | 4.03 | 3.15 | 3.14 | No | 595702 |
| SC673-14E | Zimbabwe | Eastern Africa | Kafir | 34.61 | 26.62 | 4.58 | 3.12 | 3.52 | No | 576339 |
| SC679-14E | Sudan | Eastern Africa | mixed | 20.22 | 19.49 | 3.94 | 3.04 | 3.09 | No | 534046 |

| | | | | | | | | | | |
|---|---|---|---|---|---|---|---|---|---|---|
| SC680-14E | India | Asia | mixed | 18.07 | 22.91 | 4.33 | 3.24 | 3.12 | No | No |
| SC681-14E | Sudan | Eastern Africa | mixed | 33.11 | 26.68 | 4.41 | 3.23 | 3.51 | No | No |
| SC686-14E | India | Asia | Caudatum | 26.55 | 26.06 | 4.44 | 3.22 | 3.47 | No | No |
| SC687-14E | Sudan | Eastern Africa | mixed | 28.52 | 27.35 | 4.38 | 3.28 | 3.59 | No | No |
| SC69-14E | United States | Americas | mixed | 22.89 | 21.62 | 4.07 | 2.98 | 3.38 | No | No |
| SC690-14E | Uganda | Eastern Africa | mixed | 22.76 | 26.76 | 4.67 | 3.15 | 3.45 | No | No |
| SC691-14E | Burkina Faso | Western Africa | mixed | 26.17 | 26.77 | 4.46 | 3.20 | 3.57 | No | No |
| SC692-14E | Uganda | Eastern Africa | mixed | 31.65 | 29.77 | 4.54 | 3.33 | 3.74 | No | No |
| SC693-14E | Uganda | Eastern Africa | mixed | 27.82 | 27.20 | 4.54 | 3.21 | 3.56 | No | No |
| SC694-14E | Nigeria | Western Africa | mixed | 22.39 | 22.82 | 4.09 | 3.17 | 3.36 | No | No |
| SC695-9 | United States | Americas | mixed | 31.19 | 29.88 | 4.71 | 3.27 | 3.68 | No | 656106 |
| SC7-14E | India | Asia | E African durra | 16.00 | 18.62 | 4.01 | 3.11 | 2.84 | No | No |
| SC70-14E | Kenya | Eastern Africa | mixed | 22.75 | 26.68 | 4.68 | 3.15 | 3.45 | No | No |
| SC700-14E | South Africa | Southern Africa | mixed | 31.22 | 27.55 | 4.68 | 3.22 | 3.46 | No | No |
| SC701-14E | Sudan | Eastern Africa | mixed | 28.86 | 26.41 | 4.30 | 3.36 | 3.49 | No | 533985 |
| SC702-14E | United States | Americas | mixed | 35.57 | 27.03 | 4.32 | 3.32 | 3.58 | No | 656107 |
| SC704-14E | NA | NA | mixed | NA | NA | NA | NA | NA | No | 534099 |
| SC705-14E | Japan | Asia | mixed | 25.40 | 30.10 | 4.65 | 3.31 | 3.70 | No | No |
| SC707-14E | Unknown | 0 | mixed | 29.75 | 29.37 | 4.72 | 3.29 | 3.59 | No | No |
| SC708-14E | Uganda | Eastern Africa | mixed | 29.74 | 24.33 | 4.39 | 3.09 | 3.41 | No | 533970 |
| SC709-14E | Sudan | Eastern Africa | mixed | 23.93 | 24.71 | 4.36 | 3.09 | 3.46 | No | No |
| SC712-14E | Japan | Asia | mixed | 29.88 | 23.08 | 4.11 | 3.22 | 3.31 | No | No |
| SC715-14E | NA | NA | mixed | 25.01 | 22.68 | 3.94 | 3.28 | 3.34 | No | No |
| SC716-14E | Sudan | Eastern Africa | mixed | 35.01 | 26.59 | 4.46 | 3.03 | 3.72 | No | No |
| SC719-14E | NA | NA | mixed | NA | NA | NA | NA | NA | No | 534047 |
| SC72-9 | United States | Americas | mixed | 20.03 | 23.04 | 4.24 | 3.20 | 3.24 | No | No |
| SC721-14E | Japan | Asia | mixed | 35.37 | 30.66 | 4.56 | 3.45 | 3.71 | No | No |
| SC723-14E | Sudan | Eastern Africa | mixed | 29.82 | 25.61 | 4.32 | 3.12 | 3.61 | No | No |
| SC724-14E | Sudan | Eastern Africa | Caudatum | 27.49 | 24.40 | 4.22 | 3.19 | 3.46 | No | No |
| SC725-14E | NA | NA | mixed | 35.32 | 31.76 | 4.77 | 3.34 | 3.78 | No | 534101 |
| SC726-14E | Burkina Faso | Western Africa | mixed | 29.73 | 28.22 | 4.65 | 3.22 | 3.56 | No | No |
| SC727-14E | Sudan | Eastern Africa | mixed | 21.60 | 24.79 | 4.35 | 3.27 | 3.35 | No | No |
| SC728-14E | Uganda | Eastern Africa | mixed | 26.48 | 26.75 | 4.57 | 3.27 | 3.42 | No | No |
| SC730-14E | Sudan | Eastern Africa | Caudatum | 26.99 | 25.02 | 4.35 | 3.07 | 3.51 | No | No |
| SC733-14E | Nigeria | Western Africa | mixed | 29.42 | 25.76 | 4.21 | 3.36 | 3.47 | No | No |
| SC734-14E | Sudan | Eastern Africa | mixed | 23.25 | 23.49 | 4.18 | 3.18 | 3.36 | No | 576394 |
| SC736-14E | Sudan | Eastern Africa | mixed | 26.96 | 26.20 | 4.40 | 3.15 | 3.58 | No | No |
| SC737-14E | Sudan | Eastern Africa | mixed | 27.99 | 25.60 | 4.48 | 3.16 | 3.44 | No | No |
| SC738-14E | Sudan | Eastern Africa | mixed | 23.49 | 21.65 | 4.08 | 3.00 | 3.34 | No | 597952 |
| SC74-3 | United States | Americas | mixed | NA | NA | NA | NA | NA | No | No |
| SC741-14E | United States | Americas | mixed | 38.08 | 27.70 | 4.28 | 3.38 | 3.62 | No | No |
| SC748-14E | Sudan | Eastern Africa | mixed | 30.12 | 25.32 | 4.51 | 3.21 | 3.31 | No | 533991 |
| SC749-14E | Japan | Asia | Caudatum | 29.33 | 26.40 | 4.52 | 3.22 | 3.45 | No | 576373 |
| SC751-14E | Sudan | Eastern Africa | mixed | 30.08 | 25.42 | 4.46 | 3.18 | 3.41 | No | No |
| SC752-14 | United States | Americas | mixed | 34.07 | 29.04 | 4.74 | 3.30 | 3.56 | No | No |
| SC753-3 | United States | Americas | mixed | 27.59 | 25.44 | 4.20 | 3.38 | 3.41 | No | No |
| SC754-14 | India | Asia | mixed | 24.30 | 24.21 | 4.20 | 3.19 | 3.44 | No | No |

| | | | | | | | | | | |
|---|---|---|---|---|---|---|---|---|---|---|
| SC755-14E | United States | Americas | mixed | 19.92 | 21.85 | 4.54 | 3.05 | 3.01 | No | 576350 |
| SC756-14E | Sudan | Eastern Africa | mixed | 30.34 | 23.93 | 4.05 | 3.23 | 3.45 | No | No |
| SC757-14E | Botswana | Southern Africa | mixed | 37.13 | 28.94 | 4.37 | 3.48 | 3.65 | No | 576352 |
| SC759-6 | United States | Americas | mixed | NA | NA | NA | NA | NA | No | No |
| SC760-14E | Sudan | Eastern Africa | mixed | 26.75 | 28.07 | 4.66 | 3.13 | 3.65 | No | 533949 |
| SC761-14E | Unknown | 0 | Kafir | 36.35 | 31.26 | 4.67 | 3.32 | 3.83 | No | No |
| SC762-14 | United States | Americas | Caudatum | NA | NA | NA | NA | NA | No | No |
| SC763-14E | Sudan | Eastern Africa | mixed | 23.77 | 22.67 | 3.99 | 3.24 | 3.34 | No | No |
| SC764-14E | Zimbabwe | Eastern Africa | mixed | 26.20 | 26.02 | 4.47 | 3.15 | 3.51 | No | No |
| SC77-14E | Kenya | Eastern Africa | mixed | 21.27 | 20.36 | 3.81 | 3.18 | 3.18 | No | No |
| SC770-14E | Swaziland | Southern Africa | mixed | 34.63 | 28.09 | 4.52 | 3.14 | 3.76 | No | No |
| SC773-14E | China | Asia | mixed | 30.72 | 26.39 | 4.25 | 3.40 | 3.49 | No | No |
| SC774-14 | United States | Americas | mixed | 26.42 | 23.60 | 4.09 | 3.26 | 3.37 | No | No |
| SC779-14E | Sudan | Eastern Africa | mixed | 31.89 | 30.27 | 4.58 | 3.43 | 3.65 | No | No |
| SC78-14E | Kenya | Eastern Africa | mixed | 23.19 | 21.03 | 3.88 | 3.11 | 3.30 | No | No |
| SC780-14E | Sudan | Eastern Africa | mixed | 32.30 | 27.58 | 4.54 | 3.15 | 3.66 | No | No |
| SC781-14E | Nigeria | Western Africa | mixed | 33.05 | 28.79 | 4.53 | 3.38 | 3.60 | No | No |
| SC782-14E | India | Asia | mixed | 19.97 | 22.85 | 4.22 | 3.15 | 3.27 | No | 576364 |
| SC784-14E | Burkina Faso | Western Africa | mixed | 22.17 | 19.85 | 3.77 | 3.12 | 3.19 | No | No |
| SC787-14E | Uganda | Eastern Africa | mixed | 27.56 | 27.64 | 4.43 | 3.33 | 3.56 | No | No |
| SC79-14E | Kenya | Eastern Africa | mixed | 21.45 | 19.91 | 3.87 | 3.06 | 3.18 | No | 533915 |
| SC790-6 | United States | Americas | mixed | 33.16 | 30.93 | 4.62 | 3.36 | 3.76 | No | 656120 |
| SC797-14E | Zimbabwe | Eastern Africa | mixed | 22.83 | 21.74 | 3.92 | 3.23 | 3.26 | No | No |
| SC798-14E | NA | NA | Caudatum | 30.14 | 25.53 | 4.21 | 3.27 | 3.52 | No | 533989 |
| SC80-14E | Kenya | Eastern Africa | mixed | 27.79 | 23.13 | 4.14 | 3.11 | 3.42 | No | No |
| SC800-14E | United States | Americas | Caudatum | 34.96 | 30.49 | 4.58 | 3.37 | 3.76 | No | No |
| SC803-14E | United States | Americas | Caudatum | 32.49 | 24.23 | 4.18 | 3.19 | 3.46 | No | 533964 |
| SC804-14E | Sudan | Eastern Africa | Caudatum | 27.69 | 26.86 | 4.43 | 3.22 | 3.59 | No | No |
| SC805-14E | Uganda | Eastern Africa | mixed | 25.16 | 23.83 | 4.12 | 3.21 | 3.46 | No | 533967 |
| SC807-14E | Zimbabwe | Eastern Africa | mixed | 17.73 | 18.59 | 3.76 | 3.03 | 3.04 | No | No |
| SC808-14E | United States | Americas | mixed | 31.30 | 22.07 | 4.08 | 3.08 | 3.34 | No | No |
| SC810-14E | Sudan | Eastern Africa | mixed | 24.05 | 21.67 | 4.03 | 3.08 | 3.31 | No | No |
| SC814-9 | United States | Americas | mixed | 32.93 | 25.79 | 4.23 | 3.29 | 3.55 | No | No |
| SC817-14E | India | Asia | mixed | 29.12 | 27.40 | 4.41 | 3.24 | 3.63 | No | No |
| SC819-14E | United States | Americas | Asian durras | 22.78 | 23.01 | 4.19 | 3.15 | 3.31 | No | No |
| SC821-14E | United States | Americas | Kafir | 30.05 | 24.70 | 4.31 | 3.09 | 3.52 | No | No |
| SC823-14E | United States | Americas | E African durra | 20.33 | 18.97 | 4.14 | 3.22 | 3.21 | No | No |
| SC826-14E | India | Asia | mixed | 32.85 | 26.74 | 4.38 | 3.26 | 3.58 | No | No |
| SC827-14E | India | Asia | mixed | 31.64 | 25.04 | 4.41 | 3.06 | 3.54 | No | No |
| SC83-14E | Uganda | Eastern Africa | mixed | 21.58 | 23.82 | 4.11 | 3.32 | 3.33 | No | No |
| SC830-14E | United States | Americas | mixed | 30.43 | 22.70 | 4.12 | 3.06 | 3.42 | No | No |
| SC831-14E | India | Asia | Asian durras | 20.65 | 21.64 | 3.98 | 3.19 | 3.23 | No | No |
| SC832-14E | India | Asia | Asian durras | 33.96 | 30.63 | 4.66 | 3.38 | 3.69 | No | No |
| SC833-13E | United States | Americas | Asian durras | 22.82 | 21.51 | 4.00 | 3.11 | 3.28 | No | 656108 |
| SC834-13E | United States | Americas | mixed | 24.04 | 24.86 | 4.24 | 3.18 | 3.50 | No | No |
| SC837-6 | United States | Americas | mixed | 21.59 | 19.90 | 3.77 | 3.16 | 3.13 | No | No |
| SC839-14E | India | Asia | mixed | 22.82 | 21.33 | 3.83 | 3.35 | 3.16 | No | No |

| | | | | | | | | | | |
|---|---|---|---|---|---|---|---|---|---|---|
| SC84-14E | Uganda | Eastern Africa | mixed | 21.47 | 24.96 | 4.18 | 3.29 | 3.46 | No | 534144 |
| SC841-14E | United States | Americas | mixed | 19.56 | 20.27 | 3.96 | 3.22 | 3.04 | No | No |
| SC842-14E | India | Asia | mixed | 15.37 | 17.98 | 4.16 | 2.89 | 2.80 | No | No |
| SC846-14 | United States | Americas | Caudatum | 33.54 | 29.00 | 4.46 | 3.37 | 3.72 | No | No |
| SC847-13E | United States | Americas | mixed | 33.20 | 24.70 | 4.15 | 3.24 | 3.49 | No | No |
| SC848-13 | United States | Americas | Asian durras | 25.74 | 24.13 | 4.05 | 3.33 | 3.41 | No | No |
| SC85-6 | United States | Americas | mixed | 19.35 | 20.34 | 3.91 | 3.15 | 3.15 | No | 656109 |
| SC851-14E | Egypt | Eastern Africa | mixed | 17.10 | 21.97 | 4.19 | 3.17 | 3.17 | No | No |
| SC852-14E | Ethiopia | Eastern Africa | mixed | 24.99 | 24.30 | 4.15 | 3.29 | 3.39 | No | No |
| SC854-6 | United States | Americas | mixed | 38.95 | 31.63 | 4.68 | 3.40 | 3.77 | No | No |
| SC855-14E | United States | Americas | mixed | 32.86 | 30.27 | 4.62 | 3.28 | 3.80 | No | 597945 |
| SC858-3 | United States | Americas | mixed | 27.42 | 23.13 | 4.04 | 3.19 | 3.42 | No | No |
| SC859-14E | India | Asia | Asian durras | 26.40 | 23.51 | 4.20 | 3.07 | 3.45 | No | No |
| SC86-14E | Kenya | Eastern Africa | mixed | 22.69 | 25.60 | 4.29 | 3.35 | 3.42 | No | No |
| SC863-14E | India | Asia | Kafir | 27.00 | 27.47 | 4.67 | 3.30 | 3.40 | No | No |
| SC865-14E | India | Asia | mixed | 27.72 | 24.57 | 4.11 | 3.31 | 3.43 | No | No |
| SC868-3 | United States | Americas | mixed | 30.99 | 25.13 | 4.13 | 3.36 | 3.46 | No | No |
| SC87-14E | Kenya | Eastern Africa | mixed | 23.04 | 26.73 | 4.57 | 3.28 | 3.39 | No | No |
| SC871-6 | NA | NA | mixed | 25.15 | 24.04 | 4.10 | 3.27 | 3.42 | No | No |
| SC875-14E | India | Asia | Kafir | 32.83 | 24.90 | 4.38 | 3.25 | 3.32 | No | No |
| SC876-14E | United States | Americas | Asian durras | 28.19 | 26.07 | 4.51 | 3.14 | 3.51 | No | No |
| SC877-3 | United States | Americas | mixed | 31.31 | 23.97 | 3.92 | 3.47 | 3.35 | No | No |
| SC888-14E | United States | Americas | Asian durras | 30.57 | 27.47 | 4.37 | 3.36 | 3.56 | No | No |
| SC891-14E | India | Asia | Asian durras | 19.17 | 19.96 | 3.83 | 3.11 | 3.17 | No | No |
| SC893-3 | United States | Americas | mixed | 37.65 | 30.93 | 4.59 | 3.44 | 3.73 | No | No |
| SC895-3 | United States | Americas | mixed | 38.08 | 34.11 | 4.96 | 3.31 | 3.93 | No | No |
| SC90-14E | Zaire | Western Africa | mixed | 24.52 | 24.09 | 4.37 | 3.14 | 3.35 | No | No |
| SC902-14E | India | Asia | Kafir | 33.39 | 28.05 | 4.53 | 3.16 | 3.70 | No | No |
| SC905-14E | United States | Americas | mixed | 34.93 | 23.90 | 4.13 | 3.17 | 3.47 | No | No |
| SC906-14E | Sudan | Eastern Africa | mixed | 31.46 | 24.98 | 4.11 | 3.30 | 3.45 | No | No |
| SC91-14E | Zimbabwe | Eastern Africa | mixed | 30.70 | 27.87 | 4.99 | 3.05 | 3.49 | No | 534145 |
| SC910-14E | India | Asia | Kafir | 33.24 | 28.43 | 4.62 | 3.38 | 3.48 | No | 576359 |
| SC913-14E | Sudan | Eastern Africa | mixed | 22.29 | 27.51 | 4.71 | 3.13 | 3.56 | No | No |
| SC919-14E | India | Asia | mixed | 36.21 | 25.23 | 4.44 | 3.03 | 3.54 | No | No |
| SC921-3 | United States | Americas | mixed | 33.47 | 30.37 | 4.53 | 3.55 | 3.60 | No | No |
| SC923-3 | United States | Americas | mixed | 28.39 | 25.85 | 4.42 | 3.11 | 3.58 | No | No |
| SC924-14E | India | Asia | mixed | 25.59 | 22.61 | 4.05 | 3.25 | 3.27 | No | No |
| SC929-14E | India | Asia | Asian durras | 33.32 | 26.04 | 4.37 | 3.14 | 3.59 | No | 595699 |
| SC93-14E | Sudan | Eastern Africa | mixed | 35.45 | 30.65 | 4.60 | 3.38 | 3.74 | No | No |
| SC935-14E | United States | Americas | mixed | 26.32 | 25.50 | 4.28 | 3.34 | 3.40 | No | No |
| SC94-14E | Sudan | Eastern Africa | Guinea | 33.01 | 27.81 | 4.58 | 3.17 | 3.63 | No | No |
| SC941-14E | United States | Americas | mixed | NA | NA | NA | NA | NA | No | 576347 |
| SC942-14E | United States | Americas | E African durra | 17.88 | 18.04 | 3.63 | 3.09 | 2.99 | No | 576349 |
| SC947-13E | United States | Americas | mixed | 24.68 | 23.56 | 4.06 | 3.32 | 3.32 | No | 656121 |
| SC949-14E | United States | Americas | mixed | 19.04 | 21.95 | 4.21 | 3.17 | 3.13 | No | 533998 |
| SC950-14E | United States | Americas | mixed | 21.24 | 24.60 | 4.29 | 3.24 | 3.37 | No | No |
| SC951-14E | Sudan | Eastern Africa | mixed | 34.58 | 30.32 | 4.60 | 3.34 | 3.74 | No | No |

| | | | | | | | | | | |
|---|---|---|---|---|---|---|---|---|---|---|
| SC956-14E | Nigeria | Western Africa | mixed | NA | NA | NA | NA | NA | No | No |
| SC96-14E | Nigeria | Western Africa | mixed | NA | NA | NA | NA | NA | No | No |
| SC963-14E | South Africa | Southern Africa | mixed | 25.02 | 24.99 | 4.18 | 3.27 | 3.46 | No | No |
| SC964-14E | Uganda | Eastern Africa | Caudatum | 21.25 | 24.81 | 4.42 | 3.22 | 3.33 | No | 533972 |
| SC969-14E | NA | NA | mixed | 20.67 | 22.52 | 4.05 | 3.27 | 3.23 | No | No |
| SC97-14E | Nigeria | Western Africa | Guinea | 34.30 | 30.82 | 4.85 | 3.07 | 3.94 | No | No |
| SC971-14E | United States | Americas | mixed | 19.33 | 21.07 | 4.01 | 3.14 | 3.16 | No | 656111 |
| SC972-14E | NA | NA | Caudatum | NA | NA | NA | NA | NA | No | No |
| SC975-14E | Ethiopia | Eastern Africa | mixed | NA | NA | NA | NA | NA | No | No |
| SC979-14E | Ethiopia | Eastern Africa | Caudatum | 38.03 | 34.93 | 4.91 | 3.41 | 3.96 | No | 576428 |
| SC98-14E | Nigeria | Western Africa | Guinea | 28.87 | 28.40 | 4.86 | 3.08 | 3.60 | No | No |
| SC982-14 | United States | Americas | Caudatum | 41.67 | 34.68 | 5.10 | 3.22 | 4.01 | No | 576380 |
| SC987-14E | Ethiopia | Eastern Africa | mixed | 18.06 | 19.57 | 4.19 | 3.11 | 2.87 | No | 534116 |
| SC99-14E | Nigeria | Western Africa | mixed | 27.22 | 22.11 | 3.97 | 3.19 | 3.28 | No | No |
| SC998-14E | NA | NA | mixed | 25.68 | 29.13 | 4.81 | 3.15 | 3.64 | No | 534167 |
| SC999-14E | NA | NA | E African durra | 17.31 | 20.29 | 4.03 | 3.17 | 3.03 | No | No |
| IS13848 | NA | NA | Caudatum | 26.51 | 22.02 | 3.96 | 3.18 | 3.33 | No | No |
| IS22287 | NA | NA | mixed | 41.07 | 32.80 | 4.88 | 3.29 | 3.86 | No | No |
| IS25733 | NA | NA | mixed | 26.49 | 26.87 | 4.75 | 3.21 | 3.36 | No | No |
| IS27390 | NA | NA | mixed | 25.63 | 25.09 | 4.43 | 3.19 | 3.41 | No | No |
| IS8525 | NA | NA | Kafir | 22.99 | 26.27 | 4.77 | 3.21 | 3.28 | No | IS 8525(J) |
| Karper 669 | NA | NA | mixed | 34.97 | 30.06 | 4.58 | 3.47 | 3.59 | Yes | No |
| Kuyuma | NA | NA | Caudatum | 36.11 | 26.16 | 4.30 | 3.19 | 3.63 | No | 656044 |
| L1999B-5 | NA | NA | mixed | 38.34 | 32.57 | 4.82 | 3.27 | 3.91 | No | No |
| LR9198 | NA | NA | mixed | 38.22 | 32.22 | 4.86 | 3.33 | 3.80 | Yes | No |
| Macia | NA | NA | Caudatum | 36.36 | 29.61 | 4.64 | 3.21 | 3.79 | Yes | 565121 |
| PI525695 | NA | NA | mixed | NA | NA | NA | NA | NA | No | No |
| PI563516 | NA | NA | mixed | 40.17 | 34.89 | 5.00 | 3.37 | 3.91 | No | No |
| PI609477 | NA | NA | mixed | NA | NA | NA | NA | NA | No | No |
| PI656046 | NA | NA | mixed | 30.85 | 30.87 | 4.71 | 3.31 | 3.76 | No | 656046 |
| QL12 | NA | NA | mixed | 24.61 | 24.40 | 4.13 | 3.27 | 3.42 | Yes | No |
| R9188 | NA | NA | mixed | 24.42 | 23.67 | 4.34 | 3.14 | 3.30 | No | 656007 |
| R9247 | NA | NA | mixed | 26.15 | 25.99 | 4.23 | 3.41 | 3.43 | No | No |
| R931945-2-2 | NA | NA | mixed | 31.52 | 26.50 | 4.23 | 3.43 | 3.49 | No | No |
| R986087-2-4-1 | NA | NA | mixed | 34.46 | 27.98 | 4.31 | 3.45 | 3.59 | No | No |
| R993396 | NA | NA | mixed | 36.98 | 29.02 | 4.62 | 3.25 | 3.68 | No | No |
| R995248 | NA | NA | mixed | 34.32 | 29.54 | 4.62 | 3.19 | 3.82 | No | No |
| SC1012-8BK | Ethiopia | Eastern Africa | Caudatum | 30.15 | 23.20 | 4.17 | 3.08 | 3.42 | No | No |
| SC1013-11Ebk | Ethiopia | Eastern Africa | Caudatum | 41.82 | 34.84 | 5.21 | 3.19 | 3.92 | No | No |
| SC1018-11Ebk | Ethiopia | Eastern Africa | mixed | 32.06 | 28.13 | 4.82 | 3.07 | 3.64 | No | No |
| SC1047-11Ebk | Ethiopia | Eastern Africa | mixed | 26.69 | 27.01 | 4.31 | 3.38 | 3.53 | No | 656072 |
| SC1048-8bk | Ethiopia | Eastern Africa | mixed | 41.37 | 33.15 | 4.68 | 3.48 | 3.86 | No | No |
| SC1053-8bk | India | Asia | mixed | 37.87 | 32.00 | 4.69 | 3.43 | 3.76 | No | No |
| SC1061-8bk | Cameroon | Western Africa | mixed | 36.35 | 29.86 | 4.96 | 3.17 | 3.60 | No | No |
| SC1068-11Ebk | Nigeria | Western Africa | mixed | 27.79 | 26.23 | 4.35 | 3.22 | 3.56 | No | No |
| SC1075-8bk | Nigeria | Western Africa | mixed | 46.60 | 33.36 | 4.54 | 3.57 | 3.78 | No | No |
| SC1079-11Ebk | Sudan | Eastern Africa | mixed | 30.39 | 27.15 | 4.38 | 3.26 | 3.59 | No | 595714 |

| | | | | | | | | | | |
|---|---|---|---|---|---|---|---|---|---|---|
| SC1081-8bk | Ghana | Western Africa | mixed | 29.49 | 28.61 | 5.07 | 3.07 | 3.49 | No | No |
| SC1106-8bk | Ethiopia | Eastern Africa | mixed | 22.09 | 23.08 | 4.40 | 3.27 | 3.07 | No | No |
| SC1108-11Ebk | India | Asia | mixed | 22.97 | 21.68 | 4.30 | 2.96 | 3.24 | No | 597949 |
| SC1112-8bk | Sudan | Eastern Africa | mixed | 27.76 | 26.45 | 4.38 | 3.17 | 3.61 | No | No |
| SC1113-11Ebk | Sudan | Eastern Africa | mixed | 28.91 | 25.76 | 4.29 | 3.26 | 3.53 | No | No |
| SC1115-11Ebk | Nigeria | Western Africa | mixed | NA | NA | NA | NA | NA | No | No |
| SC1119-8bk | Nigeria | Western Africa | mixed | 26.92 | 29.59 | 4.71 | 3.29 | 3.65 | No | No |
| SC1120-8bk | Nigeria | Western Africa | mixed | 44.73 | 29.81 | 4.55 | 3.33 | 3.74 | No | No |
| SC1136-11Ebk | Kenya | Eastern Africa | mixed | 27.18 | 27.16 | 4.45 | 3.19 | 3.63 | No | No |
| SC1139-5bk | Ethiopia | Eastern Africa | mixed | 33.85 | 27.01 | 4.44 | 3.22 | 3.60 | No | No |
| SC1152-5bk | Ethiopia | Eastern Africa | mixed | 25.57 | 27.50 | 4.70 | 3.21 | 3.46 | No | No |
| SC1156-11Ebk | Ethiopia | Eastern Africa | mixed | 31.58 | 29.25 | 4.66 | 3.31 | 3.60 | No | No |
| SC1157-11Ebk | Ethiopia | Eastern Africa | mixed | 28.74 | 30.66 | 4.85 | 3.29 | 3.64 | No | No |
| SC1159-11Ebk | Ethiopia | Eastern Africa | mixed | 24.38 | 26.19 | 4.37 | 3.32 | 3.44 | No | No |
| SC1162-8bk | Ethiopia | Eastern Africa | mixed | 26.12 | 28.87 | 4.45 | 3.39 | 3.63 | No | No |
| SC1174-5bk | Ethiopia | Eastern Africa | mixed | 25.55 | 26.91 | 4.56 | 3.22 | 3.50 | No | No |
| SC1175-5bk | Ethiopia | Eastern Africa | mixed | 24.48 | 24.70 | 4.25 | 3.24 | 3.44 | No | No |
| SC1185-11Ebk | Ethiopia | Eastern Africa | mixed | 25.50 | 24.57 | 4.13 | 3.29 | 3.45 | No | No |
| SC1189-8bk | Ethiopia | Eastern Africa | mixed | 23.33 | 21.80 | 3.94 | 3.24 | 3.25 | No | No |
| SC1190-5bk | Ethiopia | Eastern Africa | mixed | 29.78 | 25.41 | 4.09 | 3.46 | 3.41 | No | No |
| SC1192-8bk | Ethiopia | Eastern Africa | mixed | 27.09 | 23.83 | 4.03 | 3.32 | 3.38 | No | No |
| SC1196-8bk | Ethiopia | Eastern Africa | mixed | 43.75 | 38.13 | 5.04 | 3.47 | 4.07 | No | No |
| SC1197-8bk | Ethiopia | Eastern Africa | mixed | 38.91 | 32.54 | 4.80 | 3.27 | 3.88 | No | No |
| SC1205-5bk | Senegal | Western Africa | mixed | 27.81 | 24.89 | 4.27 | 3.21 | 3.47 | No | 597965 |
| SC1209-5bk | Guatemala | Americas | mixed | 27.46 | 24.55 | 4.22 | 3.18 | 3.49 | No | No |
| SC1212-5bk | Venezuela | Americas | mixed | 24.44 | 22.80 | 3.98 | 3.26 | 3.33 | No | 597966 |
| SC1220-11Ebk | Chad | Western Africa | mixed | 20.83 | 21.89 | 4.01 | 3.23 | 3.22 | No | No |
| SC1222-11Ebk | Japan | Asia | mixed | 30.21 | 24.92 | 4.06 | 3.42 | 3.43 | No | No |
| SC1226-2bk | Nigeria | Western Africa | mixed | 24.03 | 24.90 | 4.24 | 3.16 | 3.52 | No | No |
| SC1227-2bk | Nigeria | Western Africa | mixed | NA | NA | NA | NA | NA | No | No |
| SC1228-2bk | United States | Americas | mixed | 32.40 | 28.88 | 4.38 | 3.42 | 3.68 | No | No |
| SC1230-5bk | Nigeria | Western Africa | mixed | 41.23 | 30.65 | 4.91 | 3.08 | 3.83 | No | No |
| SC1233-2bk | Nigeria | Western Africa | mixed | 24.57 | 27.11 | 4.40 | 3.26 | 3.60 | No | No |
| SC1234-2bk | Sudan | Eastern Africa | mixed | 29.60 | 29.72 | 4.87 | 3.24 | 3.57 | No | No |
| SC1236-2bk | Nigeria | Western Africa | mixed | 25.84 | 24.24 | 4.18 | 3.18 | 3.48 | No | No |
| SC1241-2bk | United States | Americas | mixed | 28.58 | 22.20 | 4.22 | 3.08 | 3.22 | No | No |
| SC1244-11Ebk | Uganda | Eastern Africa | mixed | 25.64 | 25.89 | 4.56 | 3.08 | 3.55 | No | No |
| SC1246-11Ebk | Chad | Western Africa | mixed | 28.26 | 26.98 | 4.59 | 3.21 | 3.50 | No | 595718 |
| SC1251-2bk | Nigeria | Western Africa | mixed | 23.12 | 25.42 | 4.22 | 3.20 | 3.55 | No | 656075 |
| SC1252-5bk | Sudan | Eastern Africa | mixed | 24.62 | 22.05 | 3.97 | 3.19 | 3.31 | No | No |
| SC1255-8bk | Sudan | Eastern Africa | mixed | 31.80 | 26.45 | 4.47 | 3.09 | 3.60 | No | No |
| SC1256-8bk | Sudan | Eastern Africa | mixed | 29.52 | 24.55 | 4.07 | 3.37 | 3.39 | No | No |
| SC1258-8bk | India | Asia | mixed | 25.08 | 23.85 | 4.22 | 3.22 | 3.37 | No | No |
| SC1260-5bk | Sudan | Eastern Africa | mixed | 29.29 | 23.03 | 4.05 | 3.14 | 3.44 | No | No |
| SC1264-5bk | Sudan | Eastern Africa | mixed | 28.37 | 27.01 | 4.47 | 3.27 | 3.53 | No | No |
| SC1267-8bk | Sudan | Eastern Africa | mixed | 27.02 | 26.07 | 4.47 | 3.10 | 3.59 | No | No |
| SC1268-5bk | Thailand | Asia | mixed | 30.07 | 26.72 | 4.39 | 3.17 | 3.65 | No | No |

| | | | | | | | | | | |
|---|---|---|---|---|---|---|---|---|---|---|
| SC1274-8bk | Ethiopia | Eastern Africa | Caudatum | 32.16 | 26.77 | 4.33 | 3.22 | 3.60 | No | No |
| SC1278-5bk | India | Asia | mixed | 29.11 | 28.79 | 5.18 | 3.05 | 3.47 | No | No |
| SC1280-5bk | Sudan | Eastern Africa | mixed | 37.50 | 34.51 | 5.00 | 3.36 | 3.89 | No | No |
| SC1281-5bk | Ethiopia | Eastern Africa | mixed | 25.74 | 26.28 | 4.46 | 3.14 | 3.56 | No | No |
| SC1282-5bk | Ethiopia | Eastern Africa | mixed | 21.28 | 23.38 | 4.38 | 3.09 | 3.28 | No | No |
| SC1286-2bk | India | Asia | mixed | 31.39 | 24.91 | 4.21 | 3.39 | 3.34 | No | No |
| SC1288-5bk | India | Asia | mixed | 24.25 | 23.11 | 4.17 | 3.14 | 3.37 | No | No |
| SC1290-5bk | India | Asia | mixed | 30.54 | 26.10 | 4.30 | 3.25 | 3.55 | No | No |
| SC1291-5bk | India | Asia | mixed | 27.41 | 25.57 | 4.29 | 3.17 | 3.59 | No | No |
| SC1294-8bk | Senegal | Western Africa | mixed | 32.99 | 25.66 | 4.26 | 3.26 | 3.49 | No | No |
| SC1295-8bk | Senegal | Western Africa | mixed | 33.20 | 30.51 | 4.80 | 3.20 | 3.75 | No | No |
| SC1296-5bk | Tanzania | Eastern Africa | mixed | 28.77 | 25.39 | 4.37 | 3.37 | 3.28 | No | No |
| SC1297-5bk | India | Asia | mixed | 26.48 | 23.29 | 4.00 | 3.26 | 3.40 | No | No |
| SC1301-5bk | Ethiopia | Eastern Africa | mixed | 33.42 | 25.84 | 4.29 | 3.30 | 3.48 | No | No |
| SC1303-2bk | Ethiopia | Eastern Africa | mixed | 33.77 | 27.99 | 4.32 | 3.45 | 3.62 | No | No |
| SC1304-5bk | Ethiopia | Eastern Africa | mixed | 34.93 | 31.96 | 4.89 | 3.29 | 3.77 | No | No |
| SC1308-2bk | Ethiopia | Eastern Africa | mixed | 31.46 | 27.79 | 4.29 | 3.47 | 3.56 | No | No |
| SC1309-5bk | Ethiopia | Eastern Africa | mixed | 38.41 | 32.55 | 4.57 | 3.55 | 3.81 | No | No |
| SC1323-5bk | Sudan | Eastern Africa | mixed | 32.46 | 30.09 | 4.73 | 3.14 | 3.82 | No | No |
| SC1326-5bk | Sudan | Eastern Africa | mixed | 44.75 | 32.81 | 4.74 | 3.37 | 3.92 | No | No |
| SC1327-2bk | Sudan | Eastern Africa | mixed | 26.81 | 24.99 | 4.24 | 3.26 | 3.44 | No | No |
| SC133-8BK | Ethiopia | Eastern Africa | mixed | 34.29 | 28.31 | 4.79 | 3.42 | 3.32 | No | No |
| SC1338-11Ebk | Mali | Western Africa | mixed | 34.63 | 26.19 | 4.45 | 3.15 | 3.57 | No | No |
| SC1340-2bk | Mali | Western Africa | mixed | 38.08 | 27.60 | 4.52 | 3.21 | 3.61 | No | No |
| SC1343-2bk | Mali | Western Africa | mixed | 23.42 | 22.64 | 4.13 | 3.24 | 3.21 | No | No |
| SC1344-2bk | Mali | Western Africa | mixed | 32.00 | 26.58 | 4.56 | 3.10 | 3.57 | No | No |
| SC1347-2bk | Sudan | Eastern Africa | mixed | 36.40 | 28.27 | 4.56 | 3.21 | 3.67 | No | No |
| SC1348-5bk | Sudan | Eastern Africa | mixed | 36.09 | 31.65 | 4.81 | 3.34 | 3.73 | No | No |
| SC1349-5bk | Sudan | Eastern Africa | mixed | 29.34 | 27.04 | 4.32 | 3.41 | 3.50 | No | No |
| SC1350-5bk | Sudan | Eastern Africa | mixed | 40.71 | 34.69 | 5.02 | 3.28 | 3.96 | No | No |
| SC1352-5bk | Sudan | Eastern Africa | mixed | 27.89 | 25.24 | 4.24 | 3.33 | 3.41 | No | No |
| SC1353-5bk | Sudan | Eastern Africa | mixed | 48.59 | 35.65 | 5.05 | 3.65 | 3.64 | No | No |
| SC1355-5bk | Sudan | Eastern Africa | mixed | 50.30 | 45.28 | 5.37 | 3.72 | 4.14 | No | No |
| SC1359-2bk | Sudan | Eastern Africa | mixed | 33.19 | 23.38 | 4.28 | 3.21 | 3.25 | No | No |
| SC1360-2bk | Sudan | Eastern Africa | mixed | 37.97 | 26.50 | 4.12 | 3.56 | 3.45 | No | No |
| SC1361-5bk | Sudan | Eastern Africa | mixed | 27.40 | 24.63 | 4.48 | 3.09 | 3.39 | No | No |
| SC1362-2bk | Honduras | Americas | mixed | 32.70 | 26.97 | 4.48 | 3.19 | 3.59 | No | No |
| SC1368-2bk | Honduras | Americas | mixed | 37.99 | 26.54 | 4.53 | 3.11 | 3.56 | No | No |
| SC1369-2bk | NA | NA | mixed | 28.55 | 26.49 | 4.41 | 3.18 | 3.56 | No | No |
| SC1371-2bk | Honduras | Americas | mixed | 22.49 | 21.73 | 4.03 | 3.08 | 3.34 | No | No |
| SC1374-2bk | Honduras | Americas | mixed | 29.81 | 25.28 | 4.24 | 3.31 | 3.46 | No | No |
| SC1375-2bk | Honduras | Americas | mixed | 28.32 | 22.86 | 4.11 | 3.19 | 3.42 | No | No |
| SC1377-2bk | Honduras | Americas | mixed | 23.82 | 23.28 | 4.27 | 3.06 | 3.38 | No | No |
| SC1380-11Ebk | NA | NA | mixed | 29.73 | 29.64 | 4.64 | 3.32 | 3.66 | No | No |
| SC1381-2bk | Honduras | Americas | mixed | 28.10 | 24.57 | 4.20 | 3.24 | 3.44 | No | No |
| SC1382-2bk | Honduras | Americas | mixed | 30.24 | 22.64 | 3.90 | 3.38 | 3.28 | No | No |
| SC1386-5bk | India | Asia | mixed | 26.70 | 25.56 | 4.15 | 3.38 | 3.46 | No | No |

| | | | | | | | | | | |
|---|---|---|---|---|---|---|---|---|---|---|
| SC1387-2bk | India | Asia | mixed | 28.30 | 25.34 | 4.18 | 3.33 | 3.46 | No | No |
| SC1388-2bk | India | Asia | mixed | 32.37 | 25.18 | 4.12 | 3.38 | 3.46 | No | No |
| SC1390-2bk | India | Asia | mixed | 33.13 | 28.56 | 4.54 | 3.27 | 3.68 | No | No |
| SC1392-5bk | India | Asia | mixed | 16.02 | 18.86 | 3.78 | 3.05 | 3.10 | No | No |
| SC1393-2bk | Honduras | Americas | mixed | 29.00 | 28.65 | 4.71 | 3.23 | 3.60 | No | No |
| SC1394-2bk | Guatemala | Americas | mixed | 29.17 | 25.02 | 4.25 | 3.29 | 3.36 | No | No |
| SC1395-2bk | Honduras | Americas | mixed | 34.86 | 27.45 | 4.24 | 3.46 | 3.57 | No | No |
| SC1396-2bk | Honduras | Americas | mixed | 28.28 | 26.64 | 4.32 | 3.34 | 3.52 | No | No |
| SC1397-2bk | Honduras | Americas | mixed | 25.83 | 21.96 | 3.94 | 3.19 | 3.31 | No | No |
| SC1398-2bk | Honduras | Americas | mixed | 31.69 | 26.01 | 4.15 | 3.45 | 3.47 | No | No |
| SC1410-2bk | Niger | Western Africa | mixed | 25.14 | 25.94 | 4.27 | 3.34 | 3.47 | No | No |
| SC1411-2bk | Niger | Western Africa | mixed | 33.06 | 26.88 | 4.40 | 3.43 | 3.40 | No | No |
| SC1412-2bk | Niger | Western Africa | mixed | 35.10 | 29.35 | 4.55 | 3.31 | 3.73 | No | No |
| SC1413-2bk | Niger | Western Africa | mixed | 41.74 | 35.92 | 4.88 | 3.39 | 4.06 | No | No |
| SC1415-2bk | Niger | Western Africa | mixed | 30.77 | 26.50 | 4.69 | 3.24 | 3.34 | No | No |
| SC1421-2bk | Mali | Western Africa | mixed | NA | NA | NA | NA | NA | No | No |
| SC1423-2bk | Mali | Western Africa | mixed | 32.62 | 26.49 | 4.44 | 3.17 | 3.58 | No | No |
| SC1430-2bk | Burkina Faso | Western Africa | mixed | 29.93 | 24.60 | 4.15 | 3.28 | 3.44 | No | No |
| SC1431-2bk | Burkina Faso | Western Africa | mixed | 24.70 | 25.59 | 4.45 | 3.22 | 3.40 | No | No |
| SC219-8BK | India | Asia | mixed | 25.03 | 23.17 | 4.05 | 3.25 | 3.35 | No | No |
| SC355-8BK | Nigeria | Western Africa | mixed | 28.58 | 23.43 | 4.42 | 3.10 | 3.28 | No | No |
| SC400-5BK | Nigeria | Western Africa | mixed | 37.83 | 31.58 | 4.59 | 3.43 | 3.81 | No | No |
| SC45-14TempBlk | NA | NA | mixed | 21.12 | 25.60 | 4.40 | 3.21 | 3.44 | No | No |
| SC513-11EBK | India | Asia | mixed | 22.94 | 22.57 | 4.44 | 3.02 | 3.19 | No | No |
| SC610-5BK | India | Asia | mixed | 26.46 | 29.47 | 4.81 | 3.22 | 3.60 | No | 656103 |
| SC674-5BK | NA | NA | mixed | 31.37 | 33.68 | 5.15 | 3.30 | 3.72 | No | No |
| SC676-5BK | South Africa | Southern Africa | mixed | 26.31 | 24.93 | 4.28 | 3.30 | 3.41 | No | No |
| SC678-5BK | South Africa | Southern Africa | mixed | 26.00 | 24.96 | 4.21 | 3.26 | 3.47 | No | No |
| SC688-8BK | Uganda | Eastern Africa | mixed | 29.06 | 28.45 | 4.53 | 3.29 | 3.61 | No | No |
| SC711-11EBK | Sudan | Eastern Africa | mixed | 19.33 | 20.80 | 4.04 | 3.17 | 3.08 | No | No |
| SC758-8BK | India | Asia | mixed | 19.49 | 22.64 | 4.20 | 3.07 | 3.32 | No | No |
| SC809-11EBK | Sudan | Eastern Africa | mixed | 39.21 | 35.01 | 4.65 | 3.62 | 3.91 | No | No |
| SC824-8BK | United States | Americas | mixed | NA | NA | NA | NA | NA | No | No |
| SC843-8BK | 0 | 0 | mixed | 27.57 | 24.12 | 4.07 | 3.33 | 3.38 | No | No |
| SC844-8BK | India | Asia | mixed | 17.39 | 22.20 | 4.01 | 3.29 | 3.17 | No | No |
| SC850-8BK | India | Asia | mixed | 32.66 | 28.72 | 4.33 | 3.47 | 3.65 | No | No |
| SC916-11EBK | India | Asia | mixed | 33.98 | 26.95 | 4.30 | 3.37 | 3.53 | No | No |
| SC92-4 | NA | NA | mixed | 18.96 | 21.86 | 4.04 | 3.23 | 3.19 | No | No |
| SC961-2BK | India | Asia | mixed | 25.83 | 27.22 | 4.52 | 3.30 | 3.50 | No | No |
| SC962-2BK | India | Asia | mixed | 23.46 | 26.65 | 4.48 | 3.26 | 3.48 | No | No |
| Sureno | NA | NA | Caudatum | 30.38 | 26.60 | 4.42 | 3.14 | 3.64 | No | 561472 |
