## Supplemental Table 2 for "Large-scale GWAS in sorghum reveals common genetic control of grain size among cereals"

### Table S2 Summary of the BC-NAM

| RP | NRP | NRP Race | NRP group | NRP origin | NRP in DP | No. progenies | Average grain size per family | | | | |
|---|---|---|---|---|---|---|---|---|---|---|---|
| | | | | | | | TKW | Length | Thickness | Volume | Width |
| R931945-2-2 | Ai4 | Mixed | Landraces | China | No | 83 | 29.87 | 4.26 | 3.34 | 25.79 | 3.49 |
| R931945-2-2 | M35-1 | Mixed | Landraces | India | Yes | 41 | 29.58 | 4.11 | 3.32 | 25.35 | 3.43 |
| R931945-2-2 | RTAM422 | Mixed | Improved varieties | NA | Yes | 20 | 29.40 | 4.17 | 3.35 | 25.57 | 3.49 |
| R931945-2-2 | RTx2737 | Mixed | Improved varieties | NA | No | 24 | 29.38 | 4.08 | 3.33 | 25.25 | 3.41 |
| R931945-2-2 | RTx430 | Mixed | Improved varieties | NA | Yes | 26 | 29.76 | 4.16 | 3.33 | 25.43 | 3.44 |
| R931945-2-2 | RTx7000 | Mixed | Improved varieties | USA | Yes | 42 | 30.48 | 4.26 | 3.28 | 26.11 | 3.53 |
| R931945-2-2 | SC103-14E | Mixed | Landraces | South Africa | Yes | 42 | 29.31 | 4.14 | 3.38 | 25.35 | 3.47 |
| R931945-2-2 | SC108-14E | Caudatum | Landraces | NA | Yes | 39 | 29.57 | 4.17 | 3.34 | 25.58 | 3.50 |
| R931945-2-2 | SC23-14E | Mixed | Landraces | NA | Yes | 40 | 29.21 | 4.15 | 3.32 | 25.49 | 3.46 |
| R931945-2-2 | SC237-14E | Caudatum | Landraces | Sudan | No | 62 | 29.71 | 4.22 | 3.34 | 25.64 | 3.47 |
| R931945-2-2 | SC35-14E | Mixed | Landraces | NA | Yes | 67 | 30.05 | 4.16 | 3.31 | 25.66 | 3.49 |
| R931945-2-2 | SC56-14E | Mixed | Landraces | Sudan | Yes | 96 | 29.75 | 4.17 | 3.36 | 25.50 | 3.47 |
| R931945-2-2 | SC62-14E | Mixed | Landraces | NA | Yes | 50 | 29.12 | 4.21 | 3.36 | 25.57 | 3.48 |
| R931945-2-2 | ICSV745 | Mixed | Improved varieties | India | No | 51 | 29.17 | 4.09 | 3.35 | 25.19 | 3.43 |
| R931945-2-2 | IS3541 | NA | Landraces | NA | No | 45 | 29.70 | 4.12 | 3.34 | 25.38 | 3.46 |
| R931945-2-2 | IS3614-2 | Mixed | Landraces | Nigeria | No | 48 | 31.95 | 4.34 | 3.23 | 26.64 | 3.60 |
| R931945-2-2 | IS9710 | Mixed | Landraces | Sudan | No | 2 | 31.40 | 4.25 | 3.39 | 25.93 | 3.59 |
| R931945-2-2 | Karper 669 | Mixed | Landraces | USA | Yes | 46 | 29.76 | 4.11 | 3.39 | 25.14 | 3.43 |
| R931945-2-2 | KS115 | Mixed | Landraces | USA | No | 44 | 30.12 | 4.22 | 3.32 | 25.90 | 3.53 |
| R931945-2-2 | LR2931-2 | Mixed | Improved varieties | NA | No | 57 | 29.50 | 4.26 | 3.38 | 25.74 | 3.51 |
| R931945-2-2 | LR9198 | Mixed | Improved varieties | China | Yes | 22 | 31.19 | 4.18 | 3.26 | 25.91 | 3.52 |
| R931945-2-2 | Macia | Caudatum | Improved varieties | Mozambique | Yes | 73 | 30.17 | 4.13 | 3.30 | 25.55 | 3.47 |
| R931945-2-2 | Malisor84-7 | NA | Landraces | Mali | No | 75 | 28.70 | 4.09 | 3.39 | 25.06 | 3.42 |
| R931945-2-2 | QL12 | Mixed | Improved varieties | Australia | Yes | 101 | 29.88 | 4.16 | 3.31 | 25.56 | 3.46 |
| R931945-2-2 | Rio | Mixed | Landraces | USA | No | 57 | 28.11 | 4.11 | 3.39 | 24.81 | 3.31 |
| R931945-2-2 | RS29 | Caudatum | Improved varieties | NA | No | 48 | 29.55 | 4.18 | 3.36 | 25.54 | 3.49 |
| R986087-2-4-1 | SC237-14E | Caudatum | Landraces | Sudan | No | 54 | 30.63 | 4.36 | 3.31 | 26.30 | 3.56 |
| R986087-2-4-1 | IS22253 | Mixed | Landraces | NA | No | 7 | 29.53 | 4.20 | 3.34 | 25.79 | 3.50 |
| R986087-2-4-1 | IS9710 | Mixed | Landraces | Sudan | No | 4 | 29.53 | 4.11 | 3.30 | 25.49 | 3.46 |
| R986087-2-4-1 | QL12 | Mixed | Improved varieties | Australia | Yes | 55 | 30.66 | 4.24 | 3.28 | 25.98 | 3.52 |
