## Supplemental Table 3 for "Large-scale GWAS in sorghum reveals common genetic control of grain size among cereals"

Table S3 Summary of field experiments

| Trial | Year | Location | Sowing date | Population | Number of plots | Number of genotypes |
|---|---|---|---|---|---|---|
| NAMGAT15 | 2014/2015 | Gatton | 22/10/2014 | BC NAM | 900 | 725 |
| NAMHER15 | 2014/2015 | Hermitage | 9/10/2014 | BC NAM | 1508 | 1164 |
| NAMGAT16 | 2015/2016 | Gatton | 24/09/2015 | BC NAM | 1521 | 1116 |
| DPGAT16 | 2015/2016 | Gatton | 24/09/2015 | DP | 880 | 658 |
| DPHER16 | 2015/2016 | Hermitage | 11/01/2016 | DP | 1400 | 888 |
