## Supplemental Table 4 for "Large-scale GWAS in sorghum reveals common genetic control of grain size among cereals"

Table S4 Racial group breakdown of the diversity panel

| ID | Caudatum | Guinea | Kafir | Asian durras | E African durras | Group |
|---|---|---|---|---|---|---|
| B599 | 0.35 | 0.00 | 0.62 | 0.03 | 0.00 | mixed |
| Early hegari | 0.90 | 0.02 | 0.00 | 0.05 | 0.03 | mixed |
| B010054 | 0.45 | 0.00 | 0.51 | 0.03 | 0.00 | mixed |
| B35 | 0.06 | 0.01 | 0.24 | 0.53 | 0.15 | mixed |
| B923296 | 0.41 | 0.00 | 0.52 | 0.05 | 0.02 | mixed |
| B963676 | 0.42 | 0.00 | 0.45 | 0.09 | 0.03 | mixed |
| B986604 | 0.43 | 0.02 | 0.54 | 0.01 | 0.01 | mixed |
| BTx623 | 0.46 | 0.00 | 0.54 | 0.00 | 0.00 | mixed |
| M35-1 | 0.86 | 0.01 | 0.12 | 0.01 | 0.00 | mixed |
| RTAM422 | 0.27 | 0.00 | 0.49 | 0.21 | 0.03 | mixed |
| RTx430 | 0.36 | 0.46 | 0.12 | 0.05 | 0.02 | mixed |
| RTx7000 | 0.04 | 0.03 | 0.63 | 0.27 | 0.02 | mixed |
| SC10-14E | 0.00 | 0.00 | 0.04 | 0.37 | 0.59 | mixed |
| SC100-14E | 0.00 | 0.76 | 0.08 | 0.04 | 0.12 | mixed |
| SC101-14E | 0.12 | 0.12 | 0.69 | 0.03 | 0.04 | mixed |
| SC1014-14E | 0.04 | 0.00 | 0.08 | 0.86 | 0.02 | mixed |
| SC1015-14E | 0.12 | 0.02 | 0.14 | 0.57 | 0.15 | mixed |
| SC1017-14E | 0.07 | 0.05 | 0.08 | 0.65 | 0.15 | mixed |
| SC1019-14E | 0.76 | 0.04 | 0.04 | 0.12 | 0.04 | mixed |
| SC1021-6 | 0.00 | 0.01 | 0.40 | 0.53 | 0.07 | mixed |
| SC1022-14E | 0.00 | 0.00 | 0.97 | 0.02 | 0.01 | Kafir |
| SC1024-9 | 0.02 | 0.03 | 0.14 | 0.67 | 0.14 | mixed |
| SC1025-14E | 0.12 | 0.14 | 0.22 | 0.27 | 0.25 | mixed |
| SC103-14E | 0.81 | 0.03 | 0.10 | 0.04 | 0.02 | mixed |
| SC1031-12E | 0.07 | 0.04 | 0.19 | 0.48 | 0.22 | mixed |
| SC1033-14E | 0.07 | 0.03 | 0.14 | 0.62 | 0.14 | mixed |
| SC1038-14E | 0.94 | 0.02 | 0.00 | 0.02 | 0.02 | mixed |
| SC1039-14E | 0.00 | 0.00 | 1.00 | 0.00 | 0.00 | Kafir |
| SC1040-14E | 0.04 | 0.09 | 0.03 | 0.65 | 0.18 | mixed |
| SC1046-14E | 0.04 | 0.06 | 0.08 | 0.67 | 0.14 | mixed |
| SC1049-14E | 0.00 | 0.00 | 0.03 | 0.97 | 0.00 | Asian durras |
| SC105-14E | 0.07 | 0.01 | 0.03 | 0.89 | 0.01 | mixed |
| SC1055-14E | 0.94 | 0.02 | 0.04 | 0.00 | 0.00 | mixed |
| SC1057-14E | 0.92 | 0.01 | 0.06 | 0.01 | 0.00 | mixed |
| SC106-14E | 0.00 | 0.00 | 0.00 | 0.00 | 1.00 | E African durras |
| SC1063-14E | 0.08 | 0.48 | 0.41 | 0.00 | 0.03 | mixed |
| SC1065-14E | 0.91 | 0.02 | 0.05 | 0.00 | 0.02 | mixed |
| SC1067-14E | 0.54 | 0.25 | 0.02 | 0.15 | 0.04 | mixed |
| SC1069-14E | 0.97 | 0.01 | 0.01 | 0.01 | 0.00 | Caudatum |
| SC1070-14E | 0.86 | 0.04 | 0.01 | 0.08 | 0.00 | mixed |
| SC1072-14 | 0.17 | 0.00 | 0.52 | 0.31 | 0.00 | mixed |
| SC1074-12E | 0.05 | 0.53 | 0.38 | 0.04 | 0.00 | mixed |
| SC1076-12E | 0.15 | 0.66 | 0.14 | 0.04 | 0.01 | mixed |
| SC1077-14E | 0.98 | 0.00 | 0.00 | 0.00 | 0.01 | Caudatum |

| | | | | | | |
|---|---|---|---|---|---|---|
| SC108-14E | 1.00 | 0.00 | 0.00 | 0.00 | 0.00 | Caudatum |
| SC1080-14E | 0.00 | 0.00 | 1.00 | 0.00 | 0.00 | Kafir |
| SC1082-6 | 0.30 | 0.32 | 0.24 | 0.04 | 0.10 | mixed |
| SC1083-14E | 0.06 | 0.51 | 0.25 | 0.00 | 0.18 | mixed |
| SC1084-14E | 0.06 | 0.47 | 0.31 | 0.00 | 0.16 | mixed |
| SC1085-14E | 0.03 | 0.02 | 0.09 | 0.84 | 0.02 | mixed |
| SC1088-14E | 0.15 | 0.08 | 0.03 | 0.73 | 0.02 | mixed |
| SC109-14E | 1.00 | 0.00 | 0.00 | 0.00 | 0.00 | Caudatum |
| SC1097-3 | 0.00 | 0.00 | 0.62 | 0.38 | 0.00 | mixed |
| SC110-14E | 1.00 | 0.00 | 0.00 | 0.00 | 0.00 | Caudatum |
| SC1101-14E | 0.08 | 0.03 | 0.01 | 0.85 | 0.03 | mixed |
| SC1103-14E | 0.77 | 0.09 | 0.06 | 0.04 | 0.04 | mixed |
| SC1104-14E | 0.70 | 0.01 | 0.20 | 0.06 | 0.03 | mixed |
| SC1107-9 | 0.08 | 0.02 | 0.44 | 0.38 | 0.07 | mixed |
| SC1109-14E | 0.15 | 0.20 | 0.58 | 0.00 | 0.07 | mixed |
| SC111-14E | 1.00 | 0.00 | 0.00 | 0.00 | 0.00 | Caudatum |
| SC1111-14E | 0.60 | 0.13 | 0.07 | 0.12 | 0.07 | mixed |
| SC1114-6 | 0.35 | 0.16 | 0.35 | 0.14 | 0.01 | mixed |
| SC1116-14E | 0.46 | 0.06 | 0.42 | 0.06 | 0.01 | mixed |
| SC1117-14E | 0.40 | 0.33 | 0.11 | 0.13 | 0.04 | mixed |
| SC1118-14E | 0.71 | 0.03 | 0.17 | 0.06 | 0.03 | mixed |
| SC112-14E | 1.00 | 0.00 | 0.00 | 0.00 | 0.00 | Caudatum |
| SC1123-14E | 0.00 | 0.97 | 0.00 | 0.03 | 0.00 | Guinea |
| SC1124-14E | 0.00 | 1.00 | 0.00 | 0.00 | 0.00 | Guinea |
| SC1125-14E | 0.00 | 1.00 | 0.00 | 0.00 | 0.00 | Guinea |
| SC113-14 | 0.76 | 0.02 | 0.16 | 0.03 | 0.03 | mixed |
| SC1133-6 | 0.44 | 0.13 | 0.40 | 0.02 | 0.01 | mixed |
| SC114-14E | 0.92 | 0.03 | 0.01 | 0.03 | 0.01 | mixed |
| SC115-14E | 0.86 | 0.06 | 0.06 | 0.02 | 0.00 | mixed |
| SC1154-14E | 0.06 | 0.06 | 0.01 | 0.71 | 0.16 | mixed |
| SC1155-14E | 0.09 | 0.05 | 0.00 | 0.68 | 0.17 | mixed |
| SC1158-5bk | 0.01 | 0.11 | 0.66 | 0.14 | 0.09 | mixed |
| SC1160-14E | 0.08 | 0.00 | 0.13 | 0.01 | 0.79 | mixed |
| SC1166-6 | 0.00 | 0.00 | 0.20 | 0.80 | 0.00 | mixed |
| SC1170-6 | 0.02 | 0.03 | 0.31 | 0.48 | 0.17 | mixed |
| SC1172-14E | 0.45 | 0.01 | 0.46 | 0.08 | 0.00 | mixed |
| SC1177-14E | 0.00 | 0.01 | 0.04 | 0.00 | 0.95 | E African durras |
| SC1178-9 | 0.59 | 0.16 | 0.19 | 0.06 | 0.01 | mixed |
| SC1179-6 | 0.03 | 0.00 | 0.16 | 0.80 | 0.00 | mixed |
| SC118-14E | 0.88 | 0.02 | 0.06 | 0.04 | 0.00 | mixed |
| SC1184-14E | 0.03 | 0.05 | 0.11 | 0.45 | 0.36 | mixed |
| SC1186-14E | 0.11 | 0.11 | 0.09 | 0.59 | 0.10 | mixed |
| SC119-14E | 0.86 | 0.01 | 0.11 | 0.00 | 0.03 | mixed |
| SC1193-3 | 0.00 | 0.01 | 0.47 | 0.48 | 0.03 | mixed |
| SC12-14E | 0.00 | 0.00 | 0.02 | 0.28 | 0.70 | mixed |
| SC120-14E | 0.98 | 0.00 | 0.00 | 0.02 | 0.00 | Caudatum |
| SC1201-14E | 0.19 | 0.11 | 0.58 | 0.08 | 0.03 | mixed |

| | | | | | | |
|---|---|---|---|---|---|---|
| SC1201-6-3 | 0.13 | 0.22 | 0.38 | 0.21 | 0.05 | mixed |
| SC1203-14E | 0.50 | 0.15 | 0.28 | 0.03 | 0.03 | mixed |
| SC121-14E | 0.05 | 0.02 | 0.93 | 0.00 | 0.00 | mixed |
| SC1211-14E | 0.36 | 0.17 | 0.36 | 0.08 | 0.02 | mixed |
| SC1214-14E | 0.75 | 0.03 | 0.12 | 0.08 | 0.02 | mixed |
| SC1215-13E | 0.15 | 0.01 | 0.58 | 0.22 | 0.03 | mixed |
| SC1218 | 0.45 | 0.24 | 0.12 | 0.15 | 0.05 | mixed |
| SC123-14E | 0.00 | 0.06 | 0.16 | 0.44 | 0.34 | mixed |
| SC124-14E | 0.00 | 0.03 | 0.00 | 0.25 | 0.71 | mixed |
| SC1261-14E | 0.96 | 0.00 | 0.00 | 0.02 | 0.01 | Caudatum |
| SC1262-14E | 0.98 | 0.00 | 0.01 | 0.00 | 0.01 | Caudatum |
| SC1263-14E | 0.88 | 0.06 | 0.02 | 0.04 | 0.00 | mixed |
| SC1271-13E | 1.00 | 0.00 | 0.00 | 0.00 | 0.00 | Caudatum |
| SC1272-5 | 0.27 | 0.00 | 0.19 | 0.53 | 0.01 | mixed |
| SC1287-14E | 0.00 | 0.01 | 0.21 | 0.00 | 0.78 | mixed |
| SC1293-14E | 0.47 | 0.35 | 0.09 | 0.07 | 0.02 | mixed |
| SC13-14E | 0.00 | 0.00 | 0.17 | 0.02 | 0.82 | mixed |
| SC1300-14E | 1.00 | 0.00 | 0.00 | 0.00 | 0.00 | Caudatum |
| SC1302-14E | 1.00 | 0.00 | 0.00 | 0.00 | 0.00 | Caudatum |
| SC1305-14E | 0.92 | 0.00 | 0.08 | 0.00 | 0.00 | mixed |
| SC1307-14E | 1.00 | 0.00 | 0.00 | 0.00 | 0.00 | Caudatum |
| SC1313-14E | 0.98 | 0.00 | 0.02 | 0.00 | 0.00 | Caudatum |
| SC1314-14E | 0.52 | 0.00 | 0.00 | 0.00 | 0.48 | mixed |
| SC1316-14E | 0.81 | 0.00 | 0.19 | 0.00 | 0.00 | mixed |
| SC1317-14E | 0.98 | 0.00 | 0.00 | 0.02 | 0.00 | Caudatum |
| SC1318-14E | 1.00 | 0.00 | 0.00 | 0.00 | 0.00 | Caudatum |
| SC1319-14E | 1.00 | 0.00 | 0.00 | 0.00 | 0.00 | Caudatum |
| SC132-12E | 0.13 | 0.04 | 0.00 | 0.67 | 0.17 | mixed |
| SC1320-14E | 0.96 | 0.02 | 0.02 | 0.00 | 0.00 | Caudatum |
| SC1321-14E | 0.99 | 0.00 | 0.00 | 0.01 | 0.00 | Caudatum |
| SC1322-14E | 0.21 | 0.12 | 0.07 | 0.43 | 0.16 | mixed |
| SC1325-14E | 0.33 | 0.13 | 0.03 | 0.38 | 0.13 | mixed |
| SC1328-14E | 0.62 | 0.07 | 0.13 | 0.12 | 0.06 | mixed |
| SC1329-14E | 0.23 | 0.12 | 0.11 | 0.40 | 0.15 | mixed |
| SC1330-14E | 0.25 | 0.16 | 0.02 | 0.41 | 0.15 | mixed |
| SC1332-14E | 0.12 | 0.74 | 0.03 | 0.11 | 0.00 | mixed |
| SC1333-14E | 0.13 | 0.69 | 0.11 | 0.07 | 0.00 | mixed |
| SC1334-5 | 0.11 | 0.83 | 0.00 | 0.04 | 0.01 | mixed |
| SC1337-14E | 0.07 | 0.62 | 0.13 | 0.09 | 0.09 | mixed |
| SC1339-14E | 0.08 | 0.82 | 0.06 | 0.04 | 0.00 | mixed |
| SC134-14E | 0.02 | 0.09 | 0.06 | 0.59 | 0.25 | mixed |
| SC1341-14E | 0.08 | 0.83 | 0.06 | 0.03 | 0.01 | mixed |
| SC1342-14E | 0.08 | 0.84 | 0.04 | 0.02 | 0.01 | mixed |
| SC1345-14E | 0.92 | 0.02 | 0.01 | 0.03 | 0.02 | mixed |
| SC135-14E | 0.09 | 0.07 | 0.03 | 0.48 | 0.32 | mixed |
| SC1351-14E | 0.64 | 0.11 | 0.04 | 0.14 | 0.06 | mixed |
| SC1356-14E | 0.62 | 0.09 | 0.13 | 0.11 | 0.06 | mixed |

| | | | | | | |
|---|---|---|---|---|---|---|
| SC137-14E | 0.00 | 0.00 | 0.00 | 0.00 | 1.00 | E African durras |
| SC139-14E | 0.04 | 0.01 | 0.09 | 0.36 | 0.49 | mixed |
| SC140-14E | 0.05 | 0.03 | 0.02 | 0.35 | 0.54 | mixed |
| SC141-14E | 0.03 | 0.05 | 0.01 | 0.39 | 0.51 | mixed |
| SC142-14E | 0.19 | 0.07 | 0.28 | 0.38 | 0.09 | mixed |
| SC1424-2 | 0.01 | 0.03 | 0.28 | 0.52 | 0.15 | mixed |
| SC1426-10E | 0.09 | 0.85 | 0.03 | 0.03 | 0.00 | mixed |
| SC1440-8 | 0.06 | 0.39 | 0.21 | 0.01 | 0.32 | mixed |
| SC145-14E | 0.00 | 0.00 | 0.04 | 0.00 | 0.96 | E African durras |
| SC146-14E | 0.89 | 0.00 | 0.07 | 0.00 | 0.03 | mixed |
| SC147-14E | 0.01 | 0.01 | 0.05 | 0.00 | 0.93 | mixed |
| SC1476-8 | 0.65 | 0.09 | 0.11 | 0.10 | 0.05 | mixed |
| SC15-14E | 0.13 | 0.05 | 0.04 | 0.00 | 0.78 | mixed |
| SC150-14E | 0.01 | 0.02 | 0.03 | 0.35 | 0.59 | mixed |
| SC154-14E | 0.01 | 0.01 | 0.05 | 0.13 | 0.80 | mixed |
| SC155-14E | 0.02 | 0.01 | 0.02 | 0.36 | 0.59 | mixed |
| SC156-14E | 0.00 | 0.00 | 0.00 | 0.00 | 1.00 | E African durras |
| SC157-6 | 0.00 | 0.12 | 0.82 | 0.06 | 0.00 | mixed |
| SC158-14E | 0.00 | 0.00 | 0.00 | 0.47 | 0.53 | mixed |
| SC159-11E | 0.00 | 0.02 | 0.04 | 0.28 | 0.66 | mixed |
| SC16-14E | 0.05 | 0.02 | 0.11 | 0.08 | 0.74 | mixed |
| SC161-14E | 0.00 | 0.01 | 0.11 | 0.44 | 0.44 | mixed |
| SC165-14E | 0.65 | 0.07 | 0.12 | 0.08 | 0.09 | mixed |
| SC166-14E | 0.00 | 0.00 | 0.00 | 0.00 | 1.00 | E African durras |
| SC167-14E | 0.07 | 0.00 | 0.06 | 0.00 | 0.87 | mixed |
| SC17-14E | 0.03 | 0.04 | 0.12 | 0.08 | 0.74 | mixed |
| SC170-14E | 0.94 | 0.00 | 0.06 | 0.00 | 0.00 | mixed |
| SC170-6-17 | 0.84 | 0.01 | 0.13 | 0.01 | 0.01 | mixed |
| SC170-6-8 | 0.51 | 0.33 | 0.07 | 0.06 | 0.02 | mixed |
| SC171-14E | 0.56 | 0.03 | 0.38 | 0.03 | 0.00 | mixed |
| SC172-12E | 1.00 | 0.00 | 0.00 | 0.00 | 0.00 | Caudatum |
| SC173-14E | 1.00 | 0.00 | 0.00 | 0.00 | 0.00 | Caudatum |
| SC175-14E | 0.96 | 0.00 | 0.00 | 0.04 | 0.00 | Caudatum |
| SC176-14E | 0.88 | 0.00 | 0.12 | 0.00 | 0.00 | mixed |
| SC180-14E | 0.00 | 0.00 | 0.04 | 0.44 | 0.52 | mixed |
| SC182-14E | 0.00 | 0.00 | 0.00 | 0.26 | 0.74 | mixed |
| SC183-14E | 0.34 | 0.17 | 0.40 | 0.03 | 0.05 | mixed |
| SC184-14E | 0.90 | 0.00 | 0.10 | 0.00 | 0.00 | mixed |
| SC185-14E | 0.80 | 0.06 | 0.11 | 0.01 | 0.01 | mixed |
| SC186-14E | 0.16 | 0.68 | 0.03 | 0.12 | 0.01 | mixed |
| SC187-14E | 0.15 | 0.61 | 0.06 | 0.14 | 0.05 | mixed |
| SC188-14E | 0.19 | 0.57 | 0.09 | 0.13 | 0.02 | mixed |
| SC189-2 | 0.01 | 0.00 | 0.01 | 0.98 | 0.00 | Asian durras |
| SC19-14E | 0.00 | 0.00 | 0.00 | 0.00 | 1.00 | E African durras |
| SC190-6 | 0.00 | 0.00 | 0.29 | 0.71 | 0.00 | mixed |
| SC191-14E | 0.10 | 0.05 | 0.13 | 0.72 | 0.00 | mixed |
| SC192-14E | 0.03 | 0.05 | 0.11 | 0.80 | 0.00 | mixed |

| | | | | | | |
|---|---|---|---|---|---|---|
| SC193-14E | 0.08 | 0.78 | 0.03 | 0.10 | 0.00 | mixed |
| SC195-14E | 0.10 | 0.18 | 0.49 | 0.17 | 0.06 | mixed |
| SC196-14E | 0.04 | 0.03 | 0.03 | 0.88 | 0.03 | mixed |
| SC2-14E | 0.00 | 0.00 | 0.00 | 0.07 | 0.93 | mixed |
| SC20-14E | 0.00 | 0.00 | 0.00 | 0.00 | 1.00 | E African durras |
| SC200-14E | 0.00 | 0.00 | 0.00 | 1.00 | 0.00 | Asian durras |
| SC201-14E | 0.02 | 0.01 | 0.03 | 0.94 | 0.00 | mixed |
| SC202-14E | 0.00 | 0.00 | 0.04 | 0.96 | 0.00 | Asian durras |
| SC203-14E | 0.00 | 0.00 | 0.01 | 0.99 | 0.00 | Asian durras |
| SC206-14E | 0.00 | 0.00 | 0.00 | 1.00 | 0.00 | Asian durras |
| SC207-14E | 0.00 | 0.00 | 0.00 | 1.00 | 0.00 | Asian durras |
| SC208-14E | 0.00 | 0.03 | 0.00 | 0.97 | 0.00 | Asian durras |
| SC209-14E | 0.06 | 0.02 | 0.00 | 0.92 | 0.00 | mixed |
| SC21-14E | 0.00 | 0.00 | 0.01 | 0.00 | 0.99 | E African durras |
| SC210-14E | 0.00 | 0.03 | 0.00 | 0.97 | 0.00 | Asian durras |
| SC211-14E | 0.00 | 0.01 | 0.00 | 0.99 | 0.00 | Asian durras |
| SC212-14E | 0.00 | 0.01 | 0.00 | 0.99 | 0.00 | Asian durras |
| SC213-14E | 0.00 | 0.53 | 0.00 | 0.47 | 0.00 | mixed |
| SC214-14E | 0.01 | 0.01 | 0.12 | 0.86 | 0.00 | mixed |
| SC215-14E | 0.00 | 0.01 | 0.00 | 0.99 | 0.00 | Asian durras |
| SC216-14E | 0.05 | 0.01 | 0.07 | 0.85 | 0.02 | mixed |
| SC217-14E | 0.00 | 0.04 | 0.00 | 0.96 | 0.00 | Asian durras |
| SC218-14E | 0.04 | 0.01 | 0.00 | 0.94 | 0.01 | mixed |
| SC22-14E | 0.06 | 0.06 | 0.10 | 0.61 | 0.17 | mixed |
| SC220-14E | 0.02 | 0.01 | 0.00 | 0.96 | 0.00 | Asian durras |
| SC221-14E | 0.00 | 0.02 | 0.00 | 0.98 | 0.00 | Asian durras |
| SC223-14E | 0.77 | 0.05 | 0.10 | 0.02 | 0.05 | mixed |
| SC224-14E | 0.00 | 0.00 | 0.01 | 0.00 | 0.99 | E African durras |
| SC23-14E | 0.03 | 0.06 | 0.12 | 0.63 | 0.17 | mixed |
| SC230-14E | 0.03 | 0.08 | 0.00 | 0.90 | 0.00 | mixed |
| SC231-14E | 0.06 | 0.03 | 0.00 | 0.91 | 0.00 | mixed |
| SC233-14E | 0.00 | 0.00 | 0.00 | 1.00 | 0.00 | Asian durras |
| SC235-14E | 0.06 | 0.09 | 0.36 | 0.14 | 0.34 | mixed |
| SC236-3 | 0.00 | 0.00 | 0.46 | 0.51 | 0.03 | mixed |
| SC24-14E | 0.03 | 0.05 | 0.16 | 0.61 | 0.16 | mixed |
| SC240-14E | 0.00 | 0.00 | 0.00 | 1.00 | 0.00 | Asian durras |
| SC241-14E | 0.08 | 0.12 | 0.40 | 0.37 | 0.03 | mixed |
| SC242-14E | 0.12 | 0.17 | 0.55 | 0.11 | 0.05 | mixed |
| SC243-14E | 0.09 | 0.16 | 0.60 | 0.10 | 0.05 | mixed |
| SC244-14E | 0.14 | 0.11 | 0.55 | 0.15 | 0.05 | mixed |
| SC245-14E | 0.05 | 0.09 | 0.07 | 0.79 | 0.00 | mixed |
| SC247-14E | 0.09 | 0.15 | 0.53 | 0.20 | 0.04 | mixed |
| SC248-14E | 0.07 | 0.14 | 0.48 | 0.26 | 0.04 | mixed |
| SC249-14E | 0.00 | 0.67 | 0.30 | 0.00 | 0.03 | mixed |
| SC25-14E | 0.06 | 0.04 | 0.11 | 0.64 | 0.15 | mixed |
| SC250-14E | 0.11 | 0.19 | 0.49 | 0.16 | 0.04 | mixed |
| SC252-14E | 0.07 | 0.18 | 0.57 | 0.15 | 0.03 | mixed |

| | | | | | | |
|---|---|---|---|---|---|---|
| SC254-14E | 0.11 | 0.15 | 0.47 | 0.24 | 0.03 | mixed |
| SC256-14E | 0.00 | 1.00 | 0.00 | 0.00 | 0.00 | Guinea |
| SC257-14E | 0.47 | 0.07 | 0.35 | 0.11 | 0.00 | mixed |
| SC258-14E | 0.14 | 0.16 | 0.63 | 0.00 | 0.07 | mixed |
| SC259-14E | 0.96 | 0.00 | 0.04 | 0.00 | 0.00 | Caudatum |
| SC260-6 | 0.07 | 0.56 | 0.30 | 0.07 | 0.00 | mixed |
| SC261-14E | 0.24 | 0.66 | 0.06 | 0.04 | 0.00 | mixed |
| SC262-14E | 0.32 | 0.54 | 0.05 | 0.05 | 0.04 | mixed |
| SC265-14E | 0.05 | 0.67 | 0.22 | 0.03 | 0.03 | mixed |
| SC266-6 | 0.08 | 0.57 | 0.28 | 0.08 | 0.00 | mixed |
| SC267-14E | 0.27 | 0.49 | 0.16 | 0.06 | 0.02 | mixed |
| SC268-14E | 0.63 | 0.10 | 0.12 | 0.13 | 0.03 | mixed |
| SC269-14E | 0.30 | 0.06 | 0.54 | 0.10 | 0.00 | mixed |
| SC27-14E | 0.02 | 0.04 | 0.02 | 0.44 | 0.48 | mixed |
| SC270-14E | 0.02 | 0.95 | 0.00 | 0.03 | 0.00 | mixed |
| SC271-14E | 0.00 | 0.95 | 0.05 | 0.00 | 0.00 | Guinea |
| SC272-14E | 0.00 | 0.97 | 0.03 | 0.00 | 0.00 | Guinea |
| SC273-14E | 0.08 | 0.84 | 0.05 | 0.03 | 0.00 | mixed |
| SC276-14E | 0.32 | 0.39 | 0.22 | 0.07 | 0.00 | mixed |
| SC277-14E | 0.00 | 0.99 | 0.01 | 0.00 | 0.00 | Guinea |
| SC278-14E | 0.11 | 0.70 | 0.13 | 0.02 | 0.04 | mixed |
| SC279-14E | 0.00 | 1.00 | 0.00 | 0.00 | 0.00 | Guinea |
| SC28-14E | 0.05 | 0.06 | 0.08 | 0.64 | 0.17 | mixed |
| SC280-14E | 0.00 | 0.88 | 0.12 | 0.00 | 0.00 | mixed |
| SC281-14E | 0.10 | 0.42 | 0.32 | 0.00 | 0.15 | mixed |
| SC283-14E | 0.08 | 0.31 | 0.56 | 0.00 | 0.05 | mixed |
| SC284-14E | 0.59 | 0.39 | 0.01 | 0.00 | 0.00 | mixed |
| SC285-14E | 0.00 | 1.00 | 0.00 | 0.00 | 0.00 | Guinea |
| SC287-14E | 0.00 | 1.00 | 0.00 | 0.00 | 0.00 | Guinea |
| SC289-14E | 0.00 | 0.95 | 0.02 | 0.02 | 0.00 | Guinea |
| SC29-14E | 0.04 | 0.07 | 0.12 | 0.65 | 0.13 | mixed |
| SC290-14E | 0.00 | 1.00 | 0.00 | 0.00 | 0.00 | Guinea |
| SC291-14E | 0.00 | 1.00 | 0.00 | 0.00 | 0.00 | Guinea |
| SC293-14E | 0.02 | 0.96 | 0.00 | 0.02 | 0.00 | Guinea |
| SC295-11E | 0.00 | 1.00 | 0.00 | 0.00 | 0.00 | Guinea |
| SC296-14E | 0.09 | 0.84 | 0.07 | 0.00 | 0.00 | mixed |
| SC297-14E | 0.00 | 0.99 | 0.01 | 0.00 | 0.00 | Guinea |
| SC299-14E | 0.01 | 0.99 | 0.00 | 0.00 | 0.00 | Guinea |
| SC3-3 | 0.00 | 0.01 | 0.48 | 0.40 | 0.11 | mixed |
| SC30-14E | 0.01 | 0.01 | 0.09 | 0.20 | 0.68 | mixed |
| SC300-14E | 0.13 | 0.87 | 0.00 | 0.00 | 0.00 | mixed |
| SC301-14E | 0.03 | 0.92 | 0.04 | 0.00 | 0.00 | mixed |
| SC305-14E | 0.05 | 0.45 | 0.33 | 0.00 | 0.17 | mixed |
| SC306-14E | 0.00 | 0.00 | 0.00 | 0.00 | 1.00 | E African durras |
| SC307-14E | 0.00 | 0.00 | 0.00 | 1.00 | 0.00 | Asian durras |
| SC311-14E | 0.26 | 0.35 | 0.12 | 0.17 | 0.10 | mixed |
| SC314-6 | 0.09 | 0.08 | 0.79 | 0.00 | 0.03 | mixed |

| | | | | | | |
|---|---|---|---|---|---|---|
| SC319-14E | 0.31 | 0.25 | 0.16 | 0.15 | 0.14 | mixed |
| SC320-14E | 0.56 | 0.20 | 0.04 | 0.14 | 0.05 | mixed |
| SC323-14E | 0.39 | 0.25 | 0.25 | 0.05 | 0.07 | mixed |
| SC324-14 | 0.87 | 0.02 | 0.11 | 0.00 | 0.00 | mixed |
| SC325-14E | 0.28 | 0.26 | 0.23 | 0.05 | 0.18 | mixed |
| SC326-6 | 0.58 | 0.00 | 0.35 | 0.07 | 0.00 | mixed |
| SC328-14E | 0.97 | 0.00 | 0.03 | 0.00 | 0.00 | Caudatum |
| SC329-14E | 0.51 | 0.28 | 0.14 | 0.07 | 0.00 | mixed |
| SC330-14E | 0.90 | 0.01 | 0.07 | 0.01 | 0.00 | mixed |
| SC331-14E | 0.04 | 0.45 | 0.46 | 0.04 | 0.01 | mixed |
| SC333-14E | 0.64 | 0.18 | 0.02 | 0.11 | 0.04 | mixed |
| SC334-14E | 0.75 | 0.09 | 0.04 | 0.09 | 0.03 | mixed |
| SC335-14E | 0.49 | 0.19 | 0.10 | 0.19 | 0.04 | mixed |
| SC336-14E | 0.66 | 0.15 | 0.00 | 0.13 | 0.07 | mixed |
| SC338-14E | 0.57 | 0.11 | 0.04 | 0.20 | 0.08 | mixed |
| SC339-14E | 0.44 | 0.22 | 0.08 | 0.18 | 0.08 | mixed |
| SC340-14E | 0.51 | 0.16 | 0.04 | 0.22 | 0.08 | mixed |
| SC342-14E | 0.32 | 0.55 | 0.07 | 0.06 | 0.01 | mixed |
| SC343-9 | 0.54 | 0.04 | 0.23 | 0.15 | 0.03 | mixed |
| SC344-14E | 0.09 | 0.25 | 0.46 | 0.14 | 0.07 | mixed |
| SC345-14E | 0.32 | 0.25 | 0.19 | 0.20 | 0.04 | mixed |
| SC346-14E | 0.52 | 0.17 | 0.09 | 0.18 | 0.03 | mixed |
| SC347-14E | 0.19 | 0.54 | 0.07 | 0.17 | 0.04 | mixed |
| SC348-14E | 0.06 | 0.79 | 0.06 | 0.09 | 0.00 | mixed |
| SC349-14E | 0.36 | 0.25 | 0.13 | 0.21 | 0.05 | mixed |
| SC35-14E | 0.05 | 0.05 | 0.09 | 0.64 | 0.17 | mixed |
| SC350-14E | 0.96 | 0.00 | 0.04 | 0.00 | 0.00 | Caudatum |
| SC351-14E | 0.01 | 0.92 | 0.03 | 0.00 | 0.04 | mixed |
| SC352-14E | 0.57 | 0.10 | 0.03 | 0.20 | 0.10 | mixed |
| SC353-14E | 0.89 | 0.07 | 0.00 | 0.01 | 0.04 | mixed |
| SC354-14E | 0.01 | 0.91 | 0.06 | 0.00 | 0.02 | mixed |
| SC356-14E | 0.27 | 0.66 | 0.03 | 0.04 | 0.00 | mixed |
| SC358-14E | 0.05 | 0.80 | 0.04 | 0.11 | 0.00 | mixed |
| SC36-14E | 0.03 | 0.06 | 0.13 | 0.64 | 0.14 | mixed |
| SC362-14E | 0.10 | 0.78 | 0.00 | 0.10 | 0.02 | mixed |
| SC366-14E | 0.04 | 0.85 | 0.09 | 0.03 | 0.00 | mixed |
| SC367-14E | 0.08 | 0.81 | 0.03 | 0.08 | 0.00 | mixed |
| SC368-14E | 0.08 | 0.77 | 0.05 | 0.10 | 0.01 | mixed |
| SC369-14E | 0.07 | 0.76 | 0.06 | 0.09 | 0.02 | mixed |
| SC37-14E | 0.00 | 0.00 | 0.00 | 0.11 | 0.89 | mixed |
| SC370-14E | 0.09 | 0.74 | 0.07 | 0.11 | 0.00 | mixed |
| SC371-14E | 0.34 | 0.51 | 0.08 | 0.07 | 0.00 | mixed |
| SC372-14E | 0.03 | 0.80 | 0.13 | 0.04 | 0.00 | mixed |
| SC373-14E | 0.16 | 0.76 | 0.01 | 0.07 | 0.00 | mixed |
| SC374-14E | 0.03 | 0.96 | 0.00 | 0.00 | 0.00 | Guinea |
| SC377-14E | 0.00 | 0.98 | 0.02 | 0.00 | 0.00 | Guinea |
| SC38-14E | 0.05 | 0.07 | 0.02 | 0.67 | 0.20 | mixed |

| | | | | | | |
|---|---|---|---|---|---|---|
| SC380-14E | 0.36 | 0.35 | 0.28 | 0.01 | 0.00 | mixed |
| SC386-14E | 0.00 | 0.79 | 0.18 | 0.02 | 0.00 | mixed |
| SC387-14E | 0.01 | 0.78 | 0.02 | 0.00 | 0.18 | mixed |
| SC389-14E | 0.00 | 1.00 | 0.00 | 0.00 | 0.00 | Guinea |
| SC391-14E | 0.08 | 0.81 | 0.01 | 0.09 | 0.00 | mixed |
| SC392-14E | 0.06 | 0.81 | 0.03 | 0.08 | 0.01 | mixed |
| SC396-14E | 0.09 | 0.79 | 0.01 | 0.10 | 0.01 | mixed |
| SC397-14E | 0.14 | 0.63 | 0.12 | 0.10 | 0.02 | mixed |
| SC398-14E | 0.31 | 0.00 | 0.69 | 0.00 | 0.00 | mixed |
| SC399-14E | 0.16 | 0.66 | 0.06 | 0.11 | 0.01 | mixed |
| SC402-14E | 0.03 | 0.84 | 0.05 | 0.07 | 0.00 | mixed |
| SC403-14E | 0.00 | 1.00 | 0.00 | 0.00 | 0.00 | Guinea |
| SC405-14E | 0.90 | 0.04 | 0.00 | 0.03 | 0.02 | mixed |
| SC406-14E | 0.23 | 0.68 | 0.01 | 0.06 | 0.02 | mixed |
| SC407-14E | 0.33 | 0.44 | 0.07 | 0.08 | 0.08 | mixed |
| SC408-14E | 0.09 | 0.86 | 0.05 | 0.00 | 0.00 | mixed |
| SC409-14E | 0.20 | 0.57 | 0.08 | 0.14 | 0.00 | mixed |
| SC41-5 | 0.09 | 0.08 | 0.31 | 0.42 | 0.11 | mixed |
| SC410-14E | 0.00 | 0.00 | 0.00 | 1.00 | 0.00 | Asian durras |
| SC410-6 | 0.00 | 0.67 | 0.27 | 0.06 | 0.00 | mixed |
| SC411-14E | 0.52 | 0.12 | 0.17 | 0.10 | 0.09 | mixed |
| SC417-14E | 0.12 | 0.12 | 0.37 | 0.31 | 0.08 | mixed |
| SC418-14E | 0.71 | 0.12 | 0.12 | 0.02 | 0.04 | mixed |
| SC42-14E | 0.26 | 0.23 | 0.23 | 0.02 | 0.26 | mixed |
| SC420-14E | 0.76 | 0.06 | 0.06 | 0.08 | 0.04 | mixed |
| SC422-14E | 0.49 | 0.19 | 0.23 | 0.04 | 0.06 | mixed |
| SC423-14 | 0.95 | 0.00 | 0.05 | 0.00 | 0.00 | Caudatum |
| SC424-14E | 0.93 | 0.00 | 0.07 | 0.00 | 0.00 | mixed |
| SC425-14E | 0.55 | 0.20 | 0.05 | 0.16 | 0.04 | mixed |
| SC43-14E | 0.00 | 0.00 | 0.00 | 0.00 | 1.00 | E African durras |
| SC430-14E | 0.08 | 0.06 | 0.00 | 0.69 | 0.17 | mixed |
| SC435-14E | 0.00 | 0.00 | 0.96 | 0.04 | 0.00 | Kafir |
| SC436-9 | 0.01 | 0.02 | 0.11 | 0.87 | 0.00 | mixed |
| SC437-14E | 0.02 | 0.03 | 0.06 | 0.89 | 0.00 | mixed |
| SC438-3 | 0.00 | 0.00 | 0.64 | 0.32 | 0.03 | mixed |
| SC44-14E | 0.00 | 0.00 | 0.00 | 0.00 | 1.00 | E African durras |
| SC441-14E | 0.00 | 0.10 | 0.03 | 0.88 | 0.00 | mixed |
| SC442-14E | 0.00 | 0.04 | 0.03 | 0.93 | 0.00 | mixed |
| SC445-14E | 0.10 | 0.15 | 0.44 | 0.28 | 0.03 | mixed |
| SC449-14E | 0.10 | 0.12 | 0.43 | 0.32 | 0.04 | mixed |
| SC450-14E | 0.02 | 0.03 | 0.06 | 0.84 | 0.05 | mixed |
| SC451-14E | 0.00 | 0.06 | 0.22 | 0.72 | 0.00 | mixed |
| SC452-14E | 0.00 | 0.00 | 0.00 | 1.00 | 0.00 | Asian durras |
| SC454-14E | 0.07 | 0.04 | 0.47 | 0.32 | 0.10 | mixed |
| SC456-6 | 0.01 | 0.00 | 0.18 | 0.81 | 0.00 | mixed |
| SC457-14E | 0.03 | 0.00 | 0.14 | 0.82 | 0.01 | mixed |
| SC458-12E | 0.00 | 0.03 | 0.19 | 0.78 | 0.00 | mixed |

| | | | | | | |
|---|---|---|---|---|---|---|
| SC460-14E | 0.03 | 0.00 | 0.05 | 0.91 | 0.01 | mixed |
| SC462-14E | 0.00 | 0.02 | 0.01 | 0.96 | 0.00 | Asian durras |
| SC464-14E | 0.02 | 0.03 | 0.08 | 0.88 | 0.00 | mixed |
| SC465-14E | 0.18 | 0.28 | 0.40 | 0.07 | 0.07 | mixed |
| SC466-14E | 0.11 | 0.01 | 0.71 | 0.14 | 0.03 | mixed |
| SC467-14E | 0.00 | 0.00 | 0.25 | 0.75 | 0.00 | mixed |
| SC468-14E | 0.00 | 0.00 | 0.42 | 0.58 | 0.00 | mixed |
| SC469-14E | 0.01 | 0.00 | 0.01 | 0.98 | 0.00 | Asian durras |
| SC470-6 | 0.00 | 0.00 | 0.36 | 0.64 | 0.00 | mixed |
| SC471-14E | 0.00 | 0.00 | 0.00 | 1.00 | 0.00 | Asian durras |
| SC472-14E | 0.01 | 0.00 | 0.00 | 0.99 | 0.00 | Asian durras |
| SC473-14E | 0.02 | 0.00 | 0.07 | 0.91 | 0.00 | mixed |
| SC475-14E | 0.00 | 0.00 | 0.01 | 0.99 | 0.00 | Asian durras |
| SC477-14E | 0.00 | 0.00 | 0.06 | 0.94 | 0.00 | mixed |
| SC479-14E | 0.00 | 0.00 | 0.00 | 1.00 | 0.00 | Asian durras |
| SC48-14E | 0.44 | 0.18 | 0.13 | 0.19 | 0.06 | mixed |
| SC480-14E | 0.00 | 0.00 | 0.00 | 1.00 | 0.00 | Asian durras |
| SC482-14E | 0.00 | 0.00 | 0.00 | 1.00 | 0.00 | Asian durras |
| SC483-14E | 0.00 | 0.00 | 0.00 | 1.00 | 0.00 | Asian durras |
| SC484-14E | 0.00 | 0.00 | 0.00 | 0.87 | 0.13 | mixed |
| SC485-14E | 0.00 | 0.00 | 0.00 | 1.00 | 0.00 | Asian durras |
| SC489-14E | 0.00 | 0.00 | 0.00 | 1.00 | 0.00 | Asian durras |
| SC490-14E | 0.00 | 0.00 | 0.00 | 0.87 | 0.13 | mixed |
| SC492-14E | 0.00 | 0.00 | 0.00 | 0.99 | 0.00 | Asian durras |
| SC493-14E | 0.00 | 0.00 | 0.98 | 0.02 | 0.00 | Kafir |
| SC494-14E | 0.00 | 0.00 | 0.04 | 0.96 | 0.00 | Asian durras |
| SC497-14E | 0.00 | 0.00 | 0.03 | 0.97 | 0.00 | Asian durras |
| SC498-14E | 0.00 | 0.00 | 0.00 | 1.00 | 0.00 | Asian durras |
| SC499-14E | 0.00 | 0.00 | 0.00 | 1.00 | 0.00 | Asian durras |
| SC500-9 | 0.00 | 0.00 | 0.02 | 0.98 | 0.00 | Asian durras |
| SC501-14E | 0.51 | 0.08 | 0.00 | 0.37 | 0.05 | mixed |
| SC502-14E | 0.51 | 0.10 | 0.14 | 0.20 | 0.05 | mixed |
| SC505-14E | 0.97 | 0.00 | 0.00 | 0.00 | 0.03 | Caudatum |
| SC51-14E | 0.47 | 0.17 | 0.09 | 0.23 | 0.05 | mixed |
| SC512-14E | 0.08 | 0.06 | 0.38 | 0.45 | 0.03 | mixed |
| SC514-14E | 0.06 | 0.15 | 0.47 | 0.29 | 0.02 | mixed |
| SC515-14E | 0.00 | 1.00 | 0.00 | 0.00 | 0.00 | Guinea |
| SC516-14E | 0.09 | 0.15 | 0.46 | 0.25 | 0.04 | mixed |
| SC517-9 | 0.04 | 0.40 | 0.37 | 0.13 | 0.06 | mixed |
| SC519-14E | 0.00 | 1.00 | 0.00 | 0.00 | 0.00 | Guinea |
| SC520-14E | 0.00 | 0.96 | 0.04 | 0.00 | 0.00 | Guinea |
| SC521-6 | 0.00 | 0.81 | 0.18 | 0.01 | 0.00 | mixed |
| SC523-14E | 0.00 | 1.00 | 0.00 | 0.00 | 0.00 | Guinea |
| SC525-9 | 0.00 | 0.89 | 0.11 | 0.00 | 0.00 | mixed |
| SC526-14E | 0.00 | 0.96 | 0.04 | 0.00 | 0.00 | Guinea |
| SC527-14E | 0.00 | 0.92 | 0.06 | 0.02 | 0.00 | mixed |
| SC529-14E | 0.00 | 0.95 | 0.05 | 0.00 | 0.00 | Guinea |

| | | | | | | |
|---|---|---|---|---|---|---|
| SC532-14E | 0.00 | 1.00 | 0.00 | 0.00 | 0.00 | Guinea |
| SC536-14E | 0.00 | 1.00 | 0.00 | 0.00 | 0.00 | Guinea |
| SC537-14E | 0.00 | 0.93 | 0.03 | 0.02 | 0.02 | mixed |
| SC538-14E | 0.09 | 0.65 | 0.16 | 0.10 | 0.00 | mixed |
| SC54-14E | 0.50 | 0.13 | 0.07 | 0.22 | 0.08 | mixed |
| SC540-6 | 0.00 | 0.80 | 0.20 | 0.00 | 0.00 | mixed |
| SC544-14E | 0.00 | 0.97 | 0.03 | 0.00 | 0.00 | Guinea |
| SC545-14E | 0.00 | 0.96 | 0.02 | 0.01 | 0.00 | Guinea |
| SC546-14E | 0.00 | 0.95 | 0.05 | 0.00 | 0.00 | mixed |
| SC547-6 | 0.00 | 0.94 | 0.06 | 0.00 | 0.00 | mixed |
| SC55-14E | 0.53 | 0.12 | 0.05 | 0.25 | 0.05 | mixed |
| SC550-14E | 0.00 | 0.44 | 0.21 | 0.02 | 0.33 | mixed |
| SC553-14E | 0.00 | 0.99 | 0.00 | 0.01 | 0.00 | Guinea |
| SC556-3 | 0.00 | 0.00 | 0.97 | 0.03 | 0.00 | Kafir |
| SC557-14E | 0.14 | 0.26 | 0.50 | 0.00 | 0.10 | mixed |
| SC558-14E | 0.21 | 0.22 | 0.44 | 0.05 | 0.09 | mixed |
| SC56-14E | 0.63 | 0.06 | 0.11 | 0.15 | 0.04 | mixed |
| SC562-14E | 0.68 | 0.13 | 0.03 | 0.10 | 0.05 | mixed |
| SC563-14E | 0.47 | 0.18 | 0.19 | 0.14 | 0.03 | mixed |
| SC565-14E | 0.20 | 0.49 | 0.23 | 0.05 | 0.04 | mixed |
| SC566-14E | 0.12 | 0.84 | 0.02 | 0.03 | 0.00 | mixed |
| SC567-14E | 0.10 | 0.85 | 0.03 | 0.02 | 0.01 | mixed |
| SC57-14E | 0.98 | 0.00 | 0.01 | 0.00 | 0.01 | Caudatum |
| SC572-14E | 0.69 | 0.04 | 0.05 | 0.17 | 0.05 | mixed |
| SC574-14E | 0.85 | 0.05 | 0.09 | 0.02 | 0.00 | mixed |
| SC575-14E | 0.60 | 0.14 | 0.02 | 0.19 | 0.04 | mixed |
| SC578-14E | 0.13 | 0.19 | 0.61 | 0.00 | 0.07 | mixed |
| SC58-14E | 0.64 | 0.08 | 0.09 | 0.13 | 0.07 | mixed |
| SC580-14E | 0.00 | 0.00 | 0.02 | 0.98 | 0.00 | Asian durras |
| SC582-14E | 0.05 | 0.09 | 0.41 | 0.43 | 0.02 | mixed |
| SC586-14E | 0.81 | 0.00 | 0.17 | 0.01 | 0.00 | mixed |
| SC589-14E | 0.00 | 0.00 | 0.00 | 1.00 | 0.00 | Asian durras |
| SC59-12E | 0.31 | 0.22 | 0.32 | 0.08 | 0.07 | mixed |
| SC590-14E | 0.32 | 0.20 | 0.26 | 0.16 | 0.07 | mixed |
| SC593-14E | 0.38 | 0.27 | 0.18 | 0.08 | 0.09 | mixed |
| SC598-14E | 0.00 | 0.02 | 0.07 | 0.26 | 0.64 | mixed |
| SC599-11E | 0.59 | 0.00 | 0.41 | 0.00 | 0.00 | mixed |
| SC6-14E | 0.42 | 0.17 | 0.25 | 0.00 | 0.16 | mixed |
| SC60-14E | 0.97 | 0.01 | 0.00 | 0.02 | 0.00 | Caudatum |
| SC600-14E | 0.06 | 0.06 | 0.07 | 0.65 | 0.16 | mixed |
| SC601-14E | 0.46 | 0.11 | 0.42 | 0.00 | 0.01 | mixed |
| SC602-6 | 0.13 | 0.00 | 0.27 | 0.00 | 0.60 | mixed |
| SC603-14E | 0.00 | 0.00 | 0.95 | 0.04 | 0.00 | Kafir |
| SC604-6 | 0.30 | 0.13 | 0.44 | 0.10 | 0.02 | mixed |
| SC605-14E | 0.04 | 0.41 | 0.19 | 0.05 | 0.32 | mixed |
| SC606-14E | 0.46 | 0.02 | 0.36 | 0.10 | 0.06 | mixed |
| SC609-14E | 0.85 | 0.05 | 0.02 | 0.04 | 0.03 | mixed |

| | | | | | | |
|---|---|---|---|---|---|---|
| SC61-14E | 0.41 | 0.27 | 0.14 | 0.10 | 0.08 | mixed |
| SC614-14E | 0.00 | 0.00 | 0.85 | 0.13 | 0.02 | mixed |
| SC615-9 | 0.01 | 0.03 | 0.58 | 0.38 | 0.00 | mixed |
| SC618-6 | 0.00 | 0.00 | 0.22 | 0.78 | 0.00 | mixed |
| SC62-14E | 0.38 | 0.21 | 0.17 | 0.17 | 0.07 | mixed |
| SC620-14E | 0.06 | 0.17 | 0.50 | 0.25 | 0.02 | mixed |
| SC621-14E | 0.02 | 0.00 | 0.24 | 0.71 | 0.03 | mixed |
| SC623-14E | 0.14 | 0.15 | 0.50 | 0.15 | 0.05 | mixed |
| SC624-14E | 0.17 | 0.10 | 0.69 | 0.03 | 0.00 | mixed |
| SC625-14E | 0.00 | 0.00 | 0.88 | 0.12 | 0.00 | mixed |
| SC626-14E | 0.00 | 0.02 | 0.94 | 0.04 | 0.00 | mixed |
| SC627-14E | 0.00 | 0.00 | 0.93 | 0.07 | 0.00 | mixed |
| SC628-14E | 0.00 | 0.00 | 1.00 | 0.00 | 0.00 | Kafir |
| SC629-14E | 0.00 | 0.00 | 0.99 | 0.00 | 0.00 | Kafir |
| SC63-14E | 0.59 | 0.11 | 0.12 | 0.12 | 0.06 | mixed |
| SC630-14E | 0.00 | 0.00 | 1.00 | 0.00 | 0.00 | Kafir |
| SC631-6 | 0.00 | 0.00 | 1.00 | 0.00 | 0.00 | Kafir |
| SC632-14 | 0.00 | 0.00 | 1.00 | 0.00 | 0.00 | Kafir |
| SC634-14E | 0.00 | 0.00 | 0.89 | 0.10 | 0.01 | mixed |
| SC635-14E | 0.83 | 0.08 | 0.00 | 0.07 | 0.02 | mixed |
| SC636-6 | 0.74 | 0.00 | 0.26 | 0.00 | 0.00 | mixed |
| SC637-14E | 0.85 | 0.00 | 0.14 | 0.02 | 0.00 | mixed |
| SC639-6 | 0.61 | 0.00 | 0.36 | 0.03 | 0.00 | mixed |
| SC64-14E | 0.89 | 0.00 | 0.11 | 0.00 | 0.00 | mixed |
| SC641-14E | 0.91 | 0.00 | 0.09 | 0.00 | 0.00 | mixed |
| SC642-14E | 0.91 | 0.00 | 0.09 | 0.00 | 0.00 | mixed |
| SC643-14E | 0.88 | 0.00 | 0.12 | 0.00 | 0.00 | mixed |
| SC644-14E | 0.94 | 0.00 | 0.06 | 0.00 | 0.00 | mixed |
| SC645-14E | 0.82 | 0.00 | 0.17 | 0.01 | 0.00 | mixed |
| SC646-14E | 0.04 | 0.00 | 0.96 | 0.00 | 0.00 | Kafir |
| SC647-14E | 0.02 | 0.00 | 0.98 | 0.00 | 0.00 | Kafir |
| SC648-14E | 0.00 | 0.00 | 1.00 | 0.00 | 0.00 | Kafir |
| SC649-14E | 0.66 | 0.09 | 0.23 | 0.00 | 0.03 | mixed |
| SC650-14E | 0.00 | 0.02 | 0.00 | 0.98 | 0.00 | Asian durras |
| SC652-9 | 0.00 | 0.00 | 1.00 | 0.00 | 0.00 | Kafir |
| SC653-14E | 0.00 | 0.00 | 1.00 | 0.00 | 0.00 | Kafir |
| SC654-14E | 0.00 | 0.00 | 1.00 | 0.00 | 0.00 | Kafir |
| SC655-14E | 0.83 | 0.06 | 0.07 | 0.00 | 0.04 | mixed |
| SC657-14E | 0.00 | 0.00 | 1.00 | 0.00 | 0.00 | Kafir |
| SC659-14E | 0.35 | 0.03 | 0.47 | 0.13 | 0.02 | mixed |
| SC66-14E | 0.36 | 0.23 | 0.29 | 0.05 | 0.07 | mixed |
| SC663-14E | 0.00 | 0.00 | 1.00 | 0.00 | 0.00 | Kafir |
| SC67-14E | 0.35 | 0.25 | 0.22 | 0.10 | 0.08 | mixed |
| SC671-14E | 0.11 | 0.16 | 0.69 | 0.00 | 0.03 | mixed |
| SC672-14E | 0.03 | 0.00 | 0.97 | 0.00 | 0.00 | Kafir |
| SC673-14E | 0.00 | 0.00 | 1.00 | 0.00 | 0.00 | Kafir |
| SC679-14E | 0.49 | 0.16 | 0.29 | 0.03 | 0.03 | mixed |

| | | | | | | |
|---|---|---|---|---|---|---|
| SC680-14E | 0.00 | 0.01 | 0.91 | 0.07 | 0.01 | mixed |
| SC681-14E | 0.93 | 0.00 | 0.07 | 0.00 | 0.00 | mixed |
| SC686-14E | 0.96 | 0.00 | 0.04 | 0.00 | 0.00 | Caudatum |
| SC687-14E | 0.94 | 0.02 | 0.02 | 0.01 | 0.01 | mixed |
| SC69-14E | 0.38 | 0.19 | 0.32 | 0.07 | 0.04 | mixed |
| SC690-14E | 0.83 | 0.00 | 0.17 | 0.00 | 0.00 | mixed |
| SC691-14E | 0.71 | 0.07 | 0.12 | 0.07 | 0.04 | mixed |
| SC692-14E | 0.92 | 0.00 | 0.08 | 0.00 | 0.00 | mixed |
| SC693-14E | 0.91 | 0.00 | 0.09 | 0.00 | 0.00 | mixed |
| SC694-14E | 0.86 | 0.03 | 0.08 | 0.03 | 0.00 | mixed |
| SC695-9 | 0.87 | 0.00 | 0.13 | 0.00 | 0.00 | mixed |
| SC7-14E | 0.00 | 0.00 | 0.00 | 0.00 | 1.00 | E African durras |
| SC70-14E | 0.37 | 0.22 | 0.22 | 0.13 | 0.06 | mixed |
| SC700-14E | 0.48 | 0.23 | 0.15 | 0.07 | 0.07 | mixed |
| SC701-14E | 0.90 | 0.01 | 0.04 | 0.04 | 0.02 | mixed |
| SC702-14E | 0.71 | 0.01 | 0.08 | 0.15 | 0.05 | mixed |
| SC704-14E | 0.76 | 0.22 | 0.02 | 0.00 | 0.00 | mixed |
| SC705-14E | 0.78 | 0.05 | 0.10 | 0.03 | 0.03 | mixed |
| SC707-14E | 0.86 | 0.01 | 0.11 | 0.00 | 0.03 | mixed |
| SC708-14E | 0.94 | 0.02 | 0.02 | 0.01 | 0.00 | mixed |
| SC709-14E | 0.44 | 0.03 | 0.44 | 0.08 | 0.00 | mixed |
| SC712-14E | 0.85 | 0.07 | 0.00 | 0.06 | 0.02 | mixed |
| SC715-14E | 0.88 | 0.03 | 0.00 | 0.06 | 0.03 | mixed |
| SC716-14E | 0.00 | 0.50 | 0.50 | 0.00 | 0.00 | mixed |
| SC719-14E | 0.62 | 0.09 | 0.15 | 0.10 | 0.05 | mixed |
| SC72-9 | 0.35 | 0.21 | 0.26 | 0.08 | 0.10 | mixed |
| SC721-14E | 0.23 | 0.04 | 0.68 | 0.05 | 0.00 | mixed |
| SC723-14E | 0.81 | 0.09 | 0.01 | 0.04 | 0.04 | mixed |
| SC724-14E | 0.98 | 0.01 | 0.00 | 0.00 | 0.01 | Caudatum |
| SC725-14E | 0.74 | 0.08 | 0.09 | 0.08 | 0.00 | mixed |
| SC726-14E | 0.94 | 0.03 | 0.01 | 0.01 | 0.01 | mixed |
| SC727-14E | 0.91 | 0.02 | 0.06 | 0.01 | 0.00 | mixed |
| SC728-14E | 0.84 | 0.06 | 0.00 | 0.09 | 0.01 | mixed |
| SC730-14E | 0.98 | 0.01 | 0.01 | 0.00 | 0.00 | Caudatum |
| SC733-14E | 0.82 | 0.08 | 0.03 | 0.04 | 0.04 | mixed |
| SC734-14E | 0.03 | 0.00 | 0.93 | 0.04 | 0.00 | mixed |
| SC736-14E | 0.32 | 0.02 | 0.63 | 0.01 | 0.02 | mixed |
| SC737-14E | 0.94 | 0.02 | 0.00 | 0.02 | 0.02 | mixed |
| SC738-14E | 0.70 | 0.04 | 0.14 | 0.10 | 0.03 | mixed |
| SC74-3 | 0.06 | 0.01 | 0.74 | 0.16 | 0.02 | mixed |
| SC741-14E | 0.00 | 0.02 | 0.11 | 0.87 | 0.01 | mixed |
| SC748-14E | 0.83 | 0.00 | 0.13 | 0.04 | 0.00 | mixed |
| SC749-14E | 0.98 | 0.01 | 0.00 | 0.01 | 0.01 | Caudatum |
| SC751-14E | 0.00 | 0.37 | 0.63 | 0.00 | 0.00 | mixed |
| SC752-14 | 0.11 | 0.02 | 0.09 | 0.72 | 0.06 | mixed |
| SC753-3 | 0.36 | 0.00 | 0.57 | 0.07 | 0.00 | mixed |
| SC754-14 | 0.94 | 0.00 | 0.03 | 0.03 | 0.00 | mixed |

| | | | | | | |
|---|---|---|---|---|---|---|
| SC755-14E | 0.12 | 0.16 | 0.36 | 0.17 | 0.19 | mixed |
| SC756-14E | 0.04 | 0.00 | 0.03 | 0.90 | 0.02 | mixed |
| SC757-14E | 0.08 | 0.05 | 0.00 | 0.70 | 0.17 | mixed |
| SC759-6 | 0.68 | 0.00 | 0.24 | 0.08 | 0.00 | mixed |
| SC760-14E | 0.86 | 0.03 | 0.06 | 0.05 | 0.00 | mixed |
| SC761-14E | 0.00 | 0.00 | 1.00 | 0.00 | 0.00 | Kafir |
| SC762-14 | 0.97 | 0.02 | 0.00 | 0.00 | 0.02 | Caudatum |
| SC763-14E | 0.77 | 0.01 | 0.22 | 0.00 | 0.00 | mixed |
| SC764-14E | 0.80 | 0.01 | 0.20 | 0.00 | 0.00 | mixed |
| SC77-14E | 0.24 | 0.18 | 0.40 | 0.11 | 0.07 | mixed |
| SC770-14E | 0.89 | 0.04 | 0.00 | 0.04 | 0.03 | mixed |
| SC773-14E | 0.04 | 0.06 | 0.10 | 0.65 | 0.16 | mixed |
| SC774-14 | 0.88 | 0.01 | 0.05 | 0.03 | 0.03 | mixed |
| SC779-14E | 0.17 | 0.04 | 0.74 | 0.05 | 0.00 | mixed |
| SC78-14E | 0.23 | 0.17 | 0.42 | 0.10 | 0.08 | mixed |
| SC780-14E | 0.87 | 0.01 | 0.05 | 0.05 | 0.02 | mixed |
| SC781-14E | 0.14 | 0.04 | 0.04 | 0.75 | 0.04 | mixed |
| SC782-14E | 0.05 | 0.11 | 0.19 | 0.00 | 0.66 | mixed |
| SC784-14E | 0.01 | 0.04 | 0.07 | 0.88 | 0.00 | mixed |
| SC787-14E | 0.83 | 0.02 | 0.09 | 0.06 | 0.00 | mixed |
| SC79-14E | 0.23 | 0.18 | 0.39 | 0.12 | 0.08 | mixed |
| SC790-6 | 0.62 | 0.00 | 0.34 | 0.04 | 0.00 | mixed |
| SC797-14E | 0.04 | 0.02 | 0.94 | 0.00 | 0.00 | mixed |
| SC798-14E | 0.99 | 0.00 | 0.01 | 0.00 | 0.00 | Caudatum |
| SC80-14E | 0.61 | 0.04 | 0.14 | 0.16 | 0.06 | mixed |
| SC800-14E | 1.00 | 0.00 | 0.00 | 0.00 | 0.00 | Caudatum |
| SC803-14E | 0.96 | 0.01 | 0.00 | 0.03 | 0.00 | Caudatum |
| SC804-14E | 0.98 | 0.00 | 0.00 | 0.01 | 0.00 | Caudatum |
| SC805-14E | 0.95 | 0.00 | 0.03 | 0.01 | 0.01 | mixed |
| SC807-14E | 0.04 | 0.08 | 0.13 | 0.74 | 0.01 | mixed |
| SC808-14E | 0.58 | 0.05 | 0.25 | 0.10 | 0.01 | mixed |
| SC810-14E | 0.89 | 0.00 | 0.11 | 0.00 | 0.00 | mixed |
| SC814-9 | 0.00 | 0.00 | 0.36 | 0.64 | 0.00 | mixed |
| SC817-14E | 0.33 | 0.07 | 0.24 | 0.30 | 0.05 | mixed |
| SC819-14E | 0.00 | 0.01 | 0.02 | 0.97 | 0.00 | Asian durras |
| SC821-14E | 0.00 | 0.00 | 1.00 | 0.00 | 0.00 | Kafir |
| SC823-14E | 0.00 | 0.00 | 0.00 | 0.00 | 1.00 | E African durras |
| SC826-14E | 0.06 | 0.01 | 0.00 | 0.93 | 0.00 | mixed |
| SC827-14E | 0.08 | 0.01 | 0.04 | 0.85 | 0.02 | mixed |
| SC83-14E | 0.26 | 0.29 | 0.31 | 0.05 | 0.10 | mixed |
| SC830-14E | 0.05 | 0.02 | 0.07 | 0.83 | 0.02 | mixed |
| SC831-14E | 0.00 | 0.00 | 0.03 | 0.97 | 0.00 | Asian durras |
| SC832-14E | 0.01 | 0.01 | 0.00 | 0.99 | 0.00 | Asian durras |
| SC833-13E | 0.00 | 0.00 | 0.03 | 0.97 | 0.00 | Asian durras |
| SC834-13E | 0.05 | 0.01 | 0.00 | 0.92 | 0.02 | mixed |
| SC837-6 | 0.00 | 0.00 | 0.24 | 0.76 | 0.00 | mixed |
| SC839-14E | 0.00 | 0.00 | 0.06 | 0.93 | 0.00 | mixed |

| | | | | | | |
|---|---|---|---|---|---|---|
| SC84-14E | 0.29 | 0.29 | 0.30 | 0.03 | 0.09 | mixed |
| SC841-14E | 0.00 | 0.00 | 0.48 | 0.48 | 0.04 | mixed |
| SC842-14E | 0.08 | 0.13 | 0.35 | 0.22 | 0.23 | mixed |
| SC846-14 | 0.98 | 0.00 | 0.00 | 0.02 | 0.00 | Caudatum |
| SC847-13E | 0.00 | 0.01 | 0.04 | 0.95 | 0.00 | mixed |
| SC848-13 | 0.00 | 0.00 | 0.01 | 0.99 | 0.00 | Asian durras |
| SC85-6 | 0.42 | 0.19 | 0.25 | 0.07 | 0.07 | mixed |
| SC851-14E | 0.00 | 0.00 | 0.83 | 0.13 | 0.04 | mixed |
| SC852-14E | 0.12 | 0.08 | 0.08 | 0.59 | 0.14 | mixed |
| SC854-6 | 0.00 | 0.00 | 0.42 | 0.58 | 0.00 | mixed |
| SC855-14E | 0.13 | 0.10 | 0.17 | 0.43 | 0.17 | mixed |
| SC858-3 | 0.00 | 0.00 | 0.50 | 0.50 | 0.00 | mixed |
| SC859-14E | 0.02 | 0.02 | 0.01 | 0.96 | 0.00 | Asian durras |
| SC86-14E | 0.29 | 0.27 | 0.32 | 0.06 | 0.06 | mixed |
| SC863-14E | 0.00 | 0.00 | 1.00 | 0.00 | 0.00 | Kafir |
| SC865-14E | 0.01 | 0.00 | 0.16 | 0.82 | 0.01 | mixed |
| SC868-3 | 0.23 | 0.00 | 0.63 | 0.13 | 0.01 | mixed |
| SC87-14E | 0.15 | 0.16 | 0.65 | 0.00 | 0.04 | mixed |
| SC871-6 | 0.00 | 0.00 | 0.18 | 0.82 | 0.00 | mixed |
| SC875-14E | 0.00 | 0.00 | 1.00 | 0.00 | 0.00 | Kafir |
| SC876-14E | 0.00 | 0.00 | 0.00 | 1.00 | 0.00 | Asian durras |
| SC877-3 | 0.00 | 0.00 | 0.48 | 0.52 | 0.00 | mixed |
| SC888-14E | 0.01 | 0.00 | 0.01 | 0.98 | 0.00 | Asian durras |
| SC891-14E | 0.03 | 0.00 | 0.00 | 0.97 | 0.00 | Asian durras |
| SC893-3 | 0.00 | 0.00 | 0.56 | 0.44 | 0.00 | mixed |
| SC895-3 | 0.00 | 0.00 | 0.29 | 0.71 | 0.00 | mixed |
| SC90-14E | 0.40 | 0.26 | 0.12 | 0.13 | 0.08 | mixed |
| SC902-14E | 0.03 | 0.01 | 0.95 | 0.00 | 0.00 | Kafir |
| SC905-14E | 0.03 | 0.00 | 0.04 | 0.92 | 0.00 | mixed |
| SC906-14E | 0.95 | 0.00 | 0.00 | 0.05 | 0.00 | mixed |
| SC91-14E | 0.03 | 0.93 | 0.01 | 0.00 | 0.03 | mixed |
| SC910-14E | 0.00 | 0.00 | 1.00 | 0.00 | 0.00 | Kafir |
| SC913-14E | 0.52 | 0.03 | 0.45 | 0.00 | 0.00 | mixed |
| SC919-14E | 0.04 | 0.01 | 0.02 | 0.93 | 0.00 | mixed |
| SC921-3 | 0.00 | 0.00 | 0.41 | 0.59 | 0.00 | mixed |
| SC923-3 | 0.00 | 0.00 | 0.45 | 0.55 | 0.00 | mixed |
| SC924-14E | 0.00 | 0.00 | 0.12 | 0.88 | 0.00 | mixed |
| SC929-14E | 0.00 | 0.00 | 0.00 | 1.00 | 0.00 | Asian durras |
| SC93-14E | 0.73 | 0.09 | 0.04 | 0.12 | 0.03 | mixed |
| SC935-14E | 0.22 | 0.08 | 0.08 | 0.53 | 0.09 | mixed |
| SC94-14E | 0.00 | 0.98 | 0.02 | 0.00 | 0.00 | Guinea |
| SC941-14E | 0.07 | 0.05 | 0.53 | 0.15 | 0.20 | mixed |
| SC942-14E | 0.00 | 0.00 | 0.00 | 0.00 | 0.99 | E African durras |
| SC947-13E | 0.57 | 0.25 | 0.05 | 0.09 | 0.05 | mixed |
| SC949-14E | 0.00 | 0.01 | 0.82 | 0.12 | 0.04 | mixed |
| SC950-14E | 0.02 | 0.01 | 0.88 | 0.09 | 0.00 | mixed |
| SC951-14E | 0.03 | 0.13 | 0.20 | 0.49 | 0.15 | mixed |

| | | | | | | |
|---|---|---|---|---|---|---|
| SC956-14E | 0.00 | 0.86 | 0.11 | 0.01 | 0.02 | mixed |
| SC96-14E | 0.77 | 0.10 | 0.02 | 0.03 | 0.07 | mixed |
| SC963-14E | 0.85 | 0.06 | 0.06 | 0.00 | 0.03 | mixed |
| SC964-14E | 0.97 | 0.00 | 0.03 | 0.00 | 0.00 | Caudatum |
| SC969-14E | 0.37 | 0.23 | 0.28 | 0.06 | 0.05 | mixed |
| SC97-14 | 0.00 | 1.00 | 0.00 | 0.00 | 0.00 | #N/A |
| SC97-14E | 0.00 | 1.00 | 0.00 | 0.00 | 0.00 | Guinea |
| SC971-14E | 0.40 | 0.15 | 0.35 | 0.02 | 0.07 | mixed |
| SC972-14E | 0.98 | 0.00 | 0.00 | 0.02 | 0.00 | Caudatum |
| SC975-14E | 0.13 | 0.12 | 0.25 | 0.26 | 0.25 | mixed |
| SC979-14E | 0.96 | 0.00 | 0.04 | 0.00 | 0.00 | Caudatum |
| SC98-14E | 0.00 | 1.00 | 0.00 | 0.00 | 0.00 | Guinea |
| SC982-14 | 0.98 | 0.02 | 0.00 | 0.00 | 0.00 | Caudatum |
| SC987-14E | 0.03 | 0.05 | 0.18 | 0.13 | 0.61 | mixed |
| SC99-14E | 0.88 | 0.00 | 0.06 | 0.03 | 0.03 | mixed |
| SC998-14E | 0.12 | 0.10 | 0.16 | 0.49 | 0.12 | mixed |
| SC999-14E | 0.00 | 0.00 | 0.00 | 0.00 | 1.00 | E African durras |
| IS13848 | 0.97 | 0.01 | 0.00 | 0.00 | 0.02 | Caudatum |
| IS22287 | 0.04 | 0.00 | 0.60 | 0.27 | 0.09 | mixed |
| IS25733 | 0.11 | 0.84 | 0.00 | 0.04 | 0.01 | mixed |
| IS27390 | 0.10 | 0.66 | 0.00 | 0.13 | 0.11 | mixed |
| IS8525 | 0.00 | 0.00 | 1.00 | 0.00 | 0.00 | Kafir |
| Karper 669 | 0.07 | 0.46 | 0.33 | 0.12 | 0.03 | mixed |
| Kuyuma | 0.99 | 0.00 | 0.01 | 0.00 | 0.00 | Caudatum |
| L1999B-5 | 0.87 | 0.00 | 0.13 | 0.00 | 0.00 | mixed |
| LR9198 | 0.05 | 0.00 | 0.11 | 0.80 | 0.04 | mixed |
| Macia | 0.99 | 0.00 | 0.01 | 0.00 | 0.00 | Caudatum |
| PI525695 | 0.01 | 0.49 | 0.14 | 0.03 | 0.34 | mixed |
| PI563516 | 0.07 | 0.39 | 0.52 | 0.03 | 0.00 | mixed |
| PI609477 | 0.09 | 0.89 | 0.00 | 0.00 | 0.02 | mixed |
| PI656046 | 0.12 | 0.02 | 0.23 | 0.57 | 0.06 | mixed |
| QL12 | 0.00 | 0.51 | 0.43 | 0.04 | 0.02 | mixed |
| R9188 | 0.38 | 0.00 | 0.62 | 0.00 | 0.00 | mixed |
| R9247 | 0.44 | 0.00 | 0.51 | 0.04 | 0.00 | mixed |
| R931945-2-2 | 0.66 | 0.00 | 0.26 | 0.07 | 0.01 | mixed |
| R986087-2-4-1 | 0.69 | 0.00 | 0.23 | 0.04 | 0.03 | mixed |
| R993396 | 0.65 | 0.00 | 0.18 | 0.14 | 0.02 | mixed |
| R995248 | 0.62 | 0.03 | 0.14 | 0.17 | 0.04 | mixed |
| SC1012-8BK | 1.00 | 0.00 | 0.00 | 0.00 | 0.00 | Caudatum |
| SC1013-11Ebk | 0.99 | 0.00 | 0.00 | 0.01 | 0.00 | Caudatum |
| SC1018-11Ebk | 0.85 | 0.04 | 0.00 | 0.08 | 0.03 | mixed |
| SC1047-11Ebk | 0.06 | 0.07 | 0.05 | 0.66 | 0.16 | mixed |
| SC1048-8bk | 0.10 | 0.01 | 0.16 | 0.59 | 0.14 | mixed |
| SC1053-8bk | 0.08 | 0.05 | 0.10 | 0.63 | 0.15 | mixed |
| SC1061-8bk | 0.20 | 0.01 | 0.76 | 0.03 | 0.00 | mixed |
| SC1068-11Ebk | 0.73 | 0.01 | 0.21 | 0.04 | 0.01 | mixed |
| SC1075-8bk | 0.05 | 0.79 | 0.11 | 0.05 | 0.00 | mixed |

| | | | | | | |
|---|---|---|---|---|---|---|
| SC1079-11Ebk | 0.89 | 0.03 | 0.01 | 0.05 | 0.02 | mixed |
| SC1081-8bk | 0.00 | 0.88 | 0.12 | 0.00 | 0.00 | mixed |
| SC1106-8bk | 0.01 | 0.00 | 0.10 | 0.00 | 0.89 | mixed |
| SC1108-11Ebk | 0.11 | 0.15 | 0.57 | 0.15 | 0.03 | mixed |
| SC1112-8bk | 0.81 | 0.19 | 0.00 | 0.00 | 0.00 | mixed |
| SC1113-11Ebk | 0.69 | 0.09 | 0.07 | 0.09 | 0.06 | mixed |
| SC1115-11Ebk | 0.20 | 0.59 | 0.08 | 0.13 | 0.00 | mixed |
| SC1119-8bk | 0.00 | 0.87 | 0.10 | 0.01 | 0.01 | mixed |
| SC1120-8bk | 0.37 | 0.18 | 0.25 | 0.16 | 0.04 | mixed |
| SC1136-11Ebk | 0.52 | 0.11 | 0.27 | 0.05 | 0.05 | mixed |
| SC1139-5bk | 0.77 | 0.00 | 0.22 | 0.01 | 0.00 | mixed |
| SC1152-5bk | 0.25 | 0.07 | 0.31 | 0.29 | 0.08 | mixed |
| SC1156-11Ebk | 0.03 | 0.10 | 0.02 | 0.66 | 0.19 | mixed |
| SC1157-11Ebk | 0.05 | 0.10 | 0.05 | 0.63 | 0.17 | mixed |
| SC1159-11Ebk | 0.11 | 0.08 | 0.02 | 0.64 | 0.16 | mixed |
| SC1162-8bk | 0.07 | 0.00 | 0.13 | 0.00 | 0.80 | mixed |
| SC1174-5bk | 0.00 | 0.00 | 0.21 | 0.00 | 0.79 | mixed |
| SC1175-5bk | 0.00 | 0.00 | 0.37 | 0.23 | 0.40 | mixed |
| SC1185-11Ebk | 0.23 | 0.15 | 0.37 | 0.21 | 0.04 | mixed |
| SC1189-8bk | 0.00 | 0.00 | 0.10 | 0.12 | 0.77 | mixed |
| SC1190-5bk | 0.02 | 0.00 | 0.40 | 0.45 | 0.12 | mixed |
| SC1192-8bk | 0.38 | 0.00 | 0.28 | 0.27 | 0.07 | mixed |
| SC1196-8bk | 0.05 | 0.03 | 0.22 | 0.65 | 0.04 | mixed |
| SC1197-8bk | 0.01 | 0.03 | 0.32 | 0.56 | 0.08 | mixed |
| SC1205-5bk | 0.84 | 0.11 | 0.00 | 0.03 | 0.01 | mixed |
| SC1209-5bk | 0.18 | 0.15 | 0.66 | 0.00 | 0.01 | mixed |
| SC1212-5bk | 0.81 | 0.00 | 0.19 | 0.00 | 0.00 | mixed |
| SC1220-11Ebk | 0.53 | 0.16 | 0.12 | 0.14 | 0.05 | mixed |
| SC1222-11Ebk | 0.57 | 0.13 | 0.12 | 0.13 | 0.06 | mixed |
| SC1226-2bk | 0.33 | 0.08 | 0.48 | 0.10 | 0.02 | mixed |
| SC1227-2bk | 0.00 | 0.50 | 0.45 | 0.05 | 0.00 | mixed |
| SC1228-2bk | 0.38 | 0.00 | 0.55 | 0.06 | 0.00 | mixed |
| SC1230-5bk | 0.46 | 0.10 | 0.24 | 0.17 | 0.03 | mixed |
| SC1233-2bk | 0.00 | 0.65 | 0.35 | 0.00 | 0.00 | mixed |
| SC1234-2bk | 0.36 | 0.01 | 0.49 | 0.11 | 0.03 | mixed |
| SC1236-2bk | 0.01 | 0.00 | 0.91 | 0.08 | 0.00 | mixed |
| SC1241-2bk | 0.40 | 0.22 | 0.27 | 0.07 | 0.04 | mixed |
| SC1244-11Ebk | 0.47 | 0.17 | 0.25 | 0.09 | 0.03 | mixed |
| SC1246-11Ebk | 0.69 | 0.10 | 0.05 | 0.14 | 0.03 | mixed |
| SC1251-2bk | 0.05 | 0.00 | 0.92 | 0.04 | 0.00 | mixed |
| SC1252-5bk | 0.05 | 0.01 | 0.27 | 0.62 | 0.06 | mixed |
| SC1255-8bk | 0.91 | 0.01 | 0.07 | 0.01 | 0.00 | mixed |
| SC1256-8bk | 0.44 | 0.09 | 0.17 | 0.23 | 0.06 | mixed |
| SC1258-8bk | 0.73 | 0.00 | 0.26 | 0.01 | 0.00 | mixed |
| SC1260-5bk | 0.46 | 0.00 | 0.51 | 0.03 | 0.00 | mixed |
| SC1264-5bk | 0.67 | 0.20 | 0.09 | 0.04 | 0.00 | mixed |
| SC1267-8bk | 0.73 | 0.17 | 0.08 | 0.02 | 0.00 | mixed |

| | | | | | | |
|---|---|---|---|---|---|---|
| SC1268-5bk | 0.74 | 0.00 | 0.26 | 0.00 | 0.00 | mixed |
| SC1274-8bk | 1.00 | 0.00 | 0.00 | 0.00 | 0.00 | Caudatum |
| SC1278-5bk | 0.21 | 0.00 | 0.24 | 0.55 | 0.00 | mixed |
| SC1280-5bk | 0.62 | 0.01 | 0.24 | 0.11 | 0.02 | mixed |
| SC1281-5bk | 0.77 | 0.00 | 0.20 | 0.03 | 0.00 | mixed |
| SC1282-5bk | 0.54 | 0.02 | 0.20 | 0.18 | 0.05 | mixed |
| SC1286-2bk | 0.44 | 0.00 | 0.47 | 0.09 | 0.00 | mixed |
| SC1288-5bk | 0.27 | 0.00 | 0.73 | 0.00 | 0.00 | mixed |
| SC1290-5bk | 0.85 | 0.00 | 0.15 | 0.00 | 0.00 | mixed |
| SC1291-5bk | 0.70 | 0.00 | 0.30 | 0.00 | 0.00 | mixed |
| SC1294-8bk | 0.36 | 0.01 | 0.63 | 0.00 | 0.00 | mixed |
| SC1295-8bk | 0.45 | 0.31 | 0.15 | 0.10 | 0.00 | mixed |
| SC1296-5bk | 0.09 | 0.19 | 0.67 | 0.01 | 0.05 | mixed |
| SC1297-5bk | 0.31 | 0.00 | 0.69 | 0.01 | 0.00 | mixed |
| SC1301-5bk | 0.65 | 0.00 | 0.35 | 0.00 | 0.00 | mixed |
| SC1303-2bk | 0.54 | 0.00 | 0.40 | 0.06 | 0.00 | mixed |
| SC1304-5bk | 0.74 | 0.00 | 0.24 | 0.02 | 0.00 | mixed |
| SC1308-2bk | 0.00 | 0.00 | 0.53 | 0.41 | 0.06 | mixed |
| SC1309-5bk | 0.65 | 0.00 | 0.35 | 0.00 | 0.00 | mixed |
| SC1323-5bk | 0.16 | 0.02 | 0.33 | 0.39 | 0.10 | mixed |
| SC1326-5bk | 0.19 | 0.11 | 0.27 | 0.32 | 0.11 | mixed |
| SC1327-2bk | 0.06 | 0.02 | 0.59 | 0.29 | 0.04 | mixed |
| SC133-8BK | 0.09 | 0.02 | 0.09 | 0.50 | 0.30 | mixed |
| SC1338-11Ebk | 0.07 | 0.72 | 0.14 | 0.05 | 0.02 | mixed |
| SC1340-2bk | 0.50 | 0.23 | 0.25 | 0.00 | 0.02 | mixed |
| SC1343-2bk | 0.01 | 0.38 | 0.54 | 0.07 | 0.00 | mixed |
| SC1344-2bk | 0.56 | 0.00 | 0.44 | 0.00 | 0.00 | mixed |
| SC1347-2bk | 0.15 | 0.07 | 0.38 | 0.34 | 0.05 | mixed |
| SC1348-5bk | 0.52 | 0.06 | 0.27 | 0.10 | 0.05 | mixed |
| SC1349-5bk | 0.47 | 0.09 | 0.23 | 0.16 | 0.04 | mixed |
| SC1350-5bk | 0.42 | 0.14 | 0.26 | 0.14 | 0.04 | mixed |
| SC1352-5bk | 0.42 | 0.04 | 0.39 | 0.11 | 0.04 | mixed |
| SC1353-5bk | 0.49 | 0.13 | 0.19 | 0.16 | 0.03 | mixed |
| SC1355-5bk | 0.46 | 0.13 | 0.19 | 0.18 | 0.03 | mixed |
| SC1359-2bk | 0.49 | 0.00 | 0.47 | 0.03 | 0.00 | mixed |
| SC1360-2bk | 0.11 | 0.12 | 0.43 | 0.29 | 0.05 | mixed |
| SC1361-5bk | 0.37 | 0.13 | 0.30 | 0.17 | 0.03 | mixed |
| SC1362-2bk | 0.65 | 0.10 | 0.25 | 0.00 | 0.00 | mixed |
| SC1368-2bk | 0.62 | 0.10 | 0.21 | 0.06 | 0.00 | mixed |
| SC1369-2bk | 0.73 | 0.13 | 0.10 | 0.04 | 0.00 | mixed |
| SC1371-2bk | 0.76 | 0.00 | 0.24 | 0.00 | 0.00 | mixed |
| SC1374-2bk | 0.42 | 0.12 | 0.46 | 0.00 | 0.00 | mixed |
| SC1375-2bk | 0.76 | 0.03 | 0.20 | 0.01 | 0.00 | mixed |
| SC1377-2bk | 0.31 | 0.18 | 0.46 | 0.03 | 0.02 | mixed |
| SC1380-11Ebk | 0.51 | 0.10 | 0.23 | 0.14 | 0.02 | mixed |
| SC1381-2bk | 0.52 | 0.06 | 0.42 | 0.00 | 0.00 | mixed |
| SC1382-2bk | 0.24 | 0.00 | 0.73 | 0.03 | 0.00 | mixed |

| | | | | | | |
|---|---|---|---|---|---|---|
| SC1386-5bk | 0.26 | 0.01 | 0.67 | 0.06 | 0.00 | mixed |
| SC1387-2bk | 0.38 | 0.01 | 0.57 | 0.04 | 0.00 | mixed |
| SC1388-2bk | 0.43 | 0.00 | 0.57 | 0.00 | 0.00 | mixed |
| SC1390-2bk | 0.73 | 0.00 | 0.27 | 0.00 | 0.00 | mixed |
| SC1392-5bk | 0.10 | 0.06 | 0.04 | 0.64 | 0.15 | mixed |
| SC1393-2bk | 0.54 | 0.10 | 0.31 | 0.04 | 0.01 | mixed |
| SC1394-2bk | 0.78 | 0.00 | 0.22 | 0.00 | 0.00 | mixed |
| SC1395-2bk | 0.50 | 0.07 | 0.41 | 0.00 | 0.01 | mixed |
| SC1396-2bk | 0.50 | 0.06 | 0.41 | 0.02 | 0.00 | mixed |
| SC1397-2bk | 0.46 | 0.09 | 0.40 | 0.05 | 0.01 | mixed |
| SC1398-2bk | 0.42 | 0.07 | 0.48 | 0.04 | 0.00 | mixed |
| SC1410-2bk | 0.10 | 0.04 | 0.51 | 0.32 | 0.04 | mixed |
| SC1411-2bk | 0.09 | 0.06 | 0.47 | 0.30 | 0.07 | mixed |
| SC1412-2bk | 0.10 | 0.04 | 0.55 | 0.26 | 0.05 | mixed |
| SC1413-2bk | 0.11 | 0.05 | 0.55 | 0.25 | 0.04 | mixed |
| SC1415-2bk | 0.10 | 0.02 | 0.51 | 0.30 | 0.07 | mixed |
| SC1421-2bk | 0.05 | 0.05 | 0.38 | 0.46 | 0.06 | mixed |
| SC1423-2bk | 0.00 | 0.05 | 0.53 | 0.36 | 0.06 | mixed |
| SC1430-2bk | 0.32 | 0.00 | 0.59 | 0.08 | 0.02 | mixed |
| SC1431-2bk | 0.41 | 0.00 | 0.52 | 0.08 | 0.00 | mixed |
| SC219-8BK | 0.00 | 0.00 | 0.19 | 0.81 | 0.00 | mixed |
| SC355-8BK | 0.27 | 0.52 | 0.13 | 0.08 | 0.00 | mixed |
| SC400-5BK | 0.00 | 0.28 | 0.30 | 0.42 | 0.00 | mixed |
| C45-14TempB | 0.00 | 0.02 | 0.02 | 0.25 | 0.71 | mixed |
| SC513-11EBK | 0.08 | 0.22 | 0.56 | 0.13 | 0.01 | mixed |
| SC610-5BK | 0.01 | 0.00 | 0.35 | 0.57 | 0.08 | mixed |
| SC674-5BK | 0.73 | 0.00 | 0.26 | 0.02 | 0.00 | mixed |
| SC676-5BK | 0.34 | 0.00 | 0.66 | 0.00 | 0.00 | mixed |
| SC678-5BK | 0.80 | 0.00 | 0.20 | 0.00 | 0.00 | mixed |
| SC688-8BK | 0.88 | 0.00 | 0.12 | 0.00 | 0.00 | mixed |
| SC711-11EBK | 0.73 | 0.04 | 0.21 | 0.03 | 0.00 | mixed |
| SC758-8BK | 0.72 | 0.04 | 0.22 | 0.02 | 0.00 | mixed |
| SC809-11EBK | 0.72 | 0.00 | 0.26 | 0.03 | 0.00 | mixed |
| SC824-8BK | 0.00 | 0.05 | 0.53 | 0.34 | 0.09 | mixed |
| SC843-8BK | 0.00 | 0.00 | 0.42 | 0.58 | 0.00 | mixed |
| SC844-8BK | 0.15 | 0.00 | 0.08 | 0.76 | 0.01 | mixed |
| SC850-8BK | 0.25 | 0.00 | 0.13 | 0.62 | 0.00 | mixed |
| SC916-11EBK | 0.23 | 0.04 | 0.18 | 0.43 | 0.11 | mixed |
| SC92-4 | 0.00 | 0.00 | 0.61 | 0.36 | 0.03 | mixed |
| SC961-2BK | 0.00 | 0.00 | 0.21 | 0.79 | 0.00 | mixed |
| SC962-2BK | 0.00 | 0.00 | 0.08 | 0.92 | 0.00 | mixed |
| Sureno | 1.00 | 0.00 | 0.00 | 0.00 | 0.00 | Caudatum |
