## Supplemental Table 5 for "Large-scale GWAS in sorghum reveals common genetic control of grain size among cereals"

Table S5 List of reported grain size QTL in previous studies

| QTL ID | LG | CI start | CI end | Population | Population size | Publication | Overlap with QTL in this study |
|---|---|---|---|---|---|---|---|
| QGrnWgt1.1 | 1 | 0.00 | 5.46 | BTxARG-1/PI656056 | 279 | Boyles et al 2017 | Y |
| QGrnWgt1.2 | 1 | 0.00 | 8.61 | BTx642/BTxARG-1 | 191 | Boyles et al 2017 | Y |
| QGrnWgt1.3 | 1 | 4.47 | 23.67 | ATx623/SA2313 | 72 | Han et al 2015 | Y |
| QGrnWgt1.4 | 1 | 4.58 | 18.42 | IS2807/379 | 110 | Rami et al 1998 | Y |
| QGrnWgt1.5 | 1 | 5.34 | 22.80 | SA2313/Hiro-1 | 138 | Han et al 2015 | Y |
| QGrnWgt1.6 | 1 | 6.07 | 22.07 | K-385/SA2313 | 109 | Han et al 2015 | Y |
| QGrnWgt1.7 | 1 | 8.00 | 15.00 | Tx7078/B35 | 98 | Tuinstra et al 1997 | Y |
| QGrnWgt1.8 | 1 | 9.18 | 13.82 | BTx623/Rio | 176 | Murray et al 2008a | Y |
| QGrnWgt1.9 | 1 | 9.27 | 18.87 | SA2313/Hiro-1 | 138 | Han et al 2015 | Y |
| QGrnWgt1.10 | 1 | 19.71 | 26.04 | IS8525/R931945-2-2 | 250 | Spagnolli et al 2016 | Y |
| QGrnWgt1.11 | 1 | 20.16 | 25.59 | IS8525/R931945-2-2 | 250 | Spagnolli et al 2016 | Y |
| QGrnWgt1.12 | 1 | 20.32 | 30.32 | E-Tian/Ji2731 | 209 | Mocoeur et al 2015 | Y |
| QGrnWgt1.13 | 1 | 26.72 | 33.28 | 296B/IS18551 | 168 | Srinivas et al 2009 | Y |
| QGrnWgt1.14 | 1 | 37.75 | 44.37 | R931945–2-2/S. verticilliflorun | 217 | Tao et al., 2018 | Y |
| QGrnWgt1.15 | 1 | 61.35 | 77.65 | BTx623/S. propinquum | 370 | Feltus et al 2006 | N |
| QGrnWgt1.16 | 1 | 116.24 | 123.19 | CK60/China17 | 131 | Gelli et al 2016 | Y |
| QGrnWgt1.17 | 1 | 147.88 | 163.00 | M35-1/B35 | 245 | Nagaraja Reddy et al 2013 | Y |
| QGrnWgt1.18 | 1 | 185.56 | 194.45 | BTxARG-1/PI656056 | 279 | Boyles et al 2017 | N |
| QGrnWgt2.1 | 2 | 48.35 | 60.72 | Red Kafir/Takakibi (F2) | 118 | Shehzad and Okuno 2014 | Y |
| QGrnWgt2.2 | 2 | 140.98 | 157.09 | BTx642/BTxARG-1 | 191 | Boyles et al 2017 | Y |
| QGrnWgt2.3 | 2 | 160.00 | 170.00 | BTx623/S. propinquum | 370 | Paterson et al 1995a | Y |
| QGrnWgt2.4 | 2 | 162.77 | 173.86 | IS2449/IS1488 | 100 | Phuong et al 2013 | Y |
| QGrnWgt2.5 | 2 | 167.39 | 167.86 | R931945–2-2/S. verticilliflorun | 217 | Tao et al., 2018 | N |
| QGrnWgt2.6 | 2 | 173.53 | 187.75 | SA2313/Hiro-1 | 138 | Han et al 2015 | Y |
| QGrnWgt2.7 | 2 | 179.28 | 185.62 | IS8525/R931945-2-2 | 250 | Spagnolli et al 2016 | Y |
| QGrnWgt2.8 | 2 | 181.60 | 189.40 | IS2807/249 | 90 | Rami et al 1998 | Y |
| QGrnWgt3.1 | 3 | 4.47 | 13.98 | E-Tian/Ji2731 | 209 | Mocoeur et al 2015 | Y |
| QGrnWgt3.2 | 3 | 32.36 | 43.68 | E36-1/SPV570 | 184 | Rajkumar et al 2013 | N |
| QGrnWgt3.3 | 3 | 37.22 | 48.78 | BTx623/S. propinquum | 370 | Feltus et al 2006 | Y |
| QGrnWgt3.4 | 3 | 48.61 | 61.39 | IS2807/379 | 110 | Rami et al 1998 | Y |
| QGrnWgt3.5 | 3 | 60.99 | 74.85 | M35-1/B35 | 245 | Nagaraja Reddy et al 2013 | Y |
| QGrnWgt3.6 | 3 | 101.76 | 102.54 | R931945–2-2/S. verticilliflorun | 217 | Tao et al., 2018 | Y |
| QGrnWgt3.7 | 3 | 106.56 | 111.06 | R931945–2-2/S. verticilliflorun | 217 | Tao et al., 2018 | N |
| QGrnWgt3.8 | 3 | 145.00 | 155.00 | BTx623/S. propinquum | 370 | Paterson et al 1995a | Y |
| QGrnWgt3.9 | 3 | 149.77 | 156.12 | R931945–2-2/S. verticilliflorun | 217 | Tao et al., 2018 | Y |
| QGrnWgt4.1 | 4 | 53.08 | 59.93 | E-Tian/Ji2731 | 209 | Mocoeur et al 2015 | Y |
| QGrnWgt4.2 | 4 | 73.53 | 73.58 | R931945–2-2/S. verticilliflorun | 217 | Tao et al., 2018 | Y |
| QGrnWgt4.3 | 4 | 78.02 | 84.36 | IS8525/R931945-2-2 | 250 | Spagnolli et al 2016 | Y |
| QGrnWgt4.4 | 4 | 79.89 | 90.11 | 296B/IS18551 | 168 | Srinivas et al 2009 | Y |
| QGrnWgt4.5 | 4 | 91.08 | 99.29 | M35-1/B35 | 245 | Nagaraja Reddy et al 2013 | Y |
| QGrnWgt4.6 | 4 | 106.91 | 108.48 | R931945–2-2/S. verticilliflorun | 217 | Tao et al., 2018 | Y |
| QGrnWgt4.7 | 4 | 117.83 | 125.17 | BTx623/IS3620C | 137 | Feltus et al 2006 | Y |
| QGrnWgt4.8 | 4 | 127.08 | 135.92 | BTx623/IS3620C | 137 | Brown et al 2006 | N |
| QGrnWgt4.9 | 4 | 132.97 | 145.77 | M35-1/B35 | 245 | Nagaraja Reddy et al 2013 | N |

| | | | | | | | |
|---|---|---|---|---|---|---|---|
| QGrnWgt5.1 | 5 | 23.15 | 24.12 | R931945–2-2/S. verticilliflorun | 217 | Tao et al., 2018 | Y |
| QGrnWgt5.2 | 5 | 61.83 | 70.13 | CK60/China17 | 131 | Gelli et al 2016 | Y |
| QGrnWgt5.3 | 5 | 73.33 | 73.97 | R931945–2-2/S. verticilliflorun | 217 | Tao et al., 2018 | Y |
| QGrnWgt5.4 | 5 | 90.08 | 94.92 | BTxARG-1/PI656056 | 279 | Boyles et al 2017 | N |
| QGrnWgt6.1 | 6 | 31.97 | 33.77 | R931945–2-2/S. verticilliflorun | 217 | Tao et al., 2018 | Y |
| QGrnWgt6.2 | 6 | 46.61 | 58.39 | BTx623/S. propinquum | 370 | Feltus et al 2006 | Y |
| QGrnWgt6.3 | 6 | 59.14 | 70.67 | BTx642/BTxARG-1 | 191 | Boyles et al 2017 | N |
| QGrnWgt6.4 | 6 | 76.03 | 88.97 | BTx623/IS3620C | 137 | Feltus et al 2006 | N |
| QGrnWgt6.5 | 6 | 77.07 | 90.93 | 296B/IS18551 | 168 | Srinivas et al 2009 | Y |
| QGrnWgt6.6 | 6 | 79.41 | 85.59 | BTx623/Rio | 176 | Murray et al 2008a | N |
| QGrnWgt6.7 | 6 | 86.36 | 87.88 | R931945–2-2/S. verticilliflorun | 217 | Tao et al., 2018 | N |
| QGrnWgt6.8 | 6 | 99.17 | 101.49 | R931945–2-2/S. verticilliflorun | 217 | Tao et al., 2018 | N |
| QGrnWgt6.9 | 6 | 155.47 | 165.22 | E-Tian/Ji2731 | 209 | Mocoeur et al 2015 | Y |
| QGrnWgt7.1 | 7 | 59.46 | 60.16 | R931945–2-2/S. verticilliflorun | 217 | Tao et al., 2018 | N |
| QGrnWgt7.2 | 7 | 60.66 | 67.00 | IS8525/R931945-2-2 | 250 | Spagnolli et al 2016 | N |
| QGrnWgt7.3 | 7 | 67.14 | 75.35 | M35-1/B35 | 245 | Nagaraja Reddy et al 2013 | Y |
| QGrnWgt7.4 | 7 | 69.34 | 76.87 | BTx623/Rio | 189 | Bai et al 2017 | Y |
| QGrnWgt7.5 | 7 | 71.13 | 75.87 | IS2807/379 | 110 | Rami et al 1998 | Y |
| QGrnWgt7.6 | 7 | 71.40 | 75.60 | IS2807/379 | 110 | Rami et al 1998 | Y |
| QGrnWgt7.7 | 7 | 72.20 | 75.01 | R931945–2-2/S. verticilliflorun | 217 | Tao et al., 2018 | Y |
| QGrnWgt7.8 | 7 | 99.39 | 108.90 | E-Tian/Ji2731 | 209 | Mocoeur et al 2015 | Y |
| QGrnWgt7.9 | 7 | 111.42 | 118.85 | IS8525/R931945-2-2 | 250 | Spagnolli et al 2016 | Y |
| QGrnWgt7.10 | 7 | 117.38 | 134.62 | BTx623/S. propinquum | 370 | Feltus et al 2006 | Y |
| QGrnWgt8.1 | 8 | 5.10 | 14.99 | IS8525/R931945-2-2 | 250 | Spagnolli et al 2016 | N |
| QGrnWgt8.2 | 8 | 6.33 | 13.76 | IS8525/R931945-2-2 | 250 | Spagnolli et al 2016 | N |
| QGrnWgt8.3 | 8 | 7.33 | 12.76 | IS8525/R931945-2-2 | 250 | Spagnolli et al 2016 | N |
| QGrnWgt8.4 | 8 | 29.75 | 41.94 | BTx642/BTxARG-1 | 191 | Boyles et al 2017 | N |
| QGrnWgt8.5 | 8 | 45.89 | 59.11 | BTx623/S. propinquum | 370 | Feltus et al 2006 | Y |
| QGrnWgt8.6 | 8 | 65.79 | 74.21 | BTx623/Rio | 176 | Murray et al 2008a | Y |
| QGrnWgt8.7 | 8 | 66.18 | 77.32 | BTx623/IS3620C | 137 | Brown et al 2006 | Y |
| QGrnWgt9.1 | 9 | 21.33 | 28.78 | BTx623/IS3620C | 137 | Brown et al 2006 | N |
| QGrnWgt9.2 | 9 | 67.93 | 78.07 | BTx623/IS3620C | 137 | Brown et al 2006 | Y |
| QGrnWgt9.3 | 9 | 93.41 | 97.53 | R931945–2-2/S. verticilliflorun | 217 | Tao et al., 2018 | Y |
| QGrnWgt9.4 | 9 | 107.18 | 111.35 | BTx623/Rio | 189 | Bai et al 2017 | Y |
| QGrnWgt9.5 | 9 | 118.67 | 123.11 | M35-1/B35 | 245 | Nagaraja Reddy et al 2013 | N |
| QGrnWgt10.1 | 10 | 25.58 | 35.98 | E-Tian/Ji2731 | 209 | Mocoeur et al 2015 | Y |
| QGrnWgt10.2 | 10 | 29.16 | 42.84 | BTx623/IS3620C | 137 | Feltus et al 2006 | Y |
| QGrnWgt10.3 | 10 | 52.34 | 60.66 | BTx623/IS3620C | 137 | Feltus et al 2006 | Y |
| QGrnWgt10.4 | 10 | 55.00 | 65.00 | BTx623/S. propinquum | 370 | Paterson et al 1995a | Y |
| QGrnWgt10.5 | 10 | 59.86 | 72.75 | R931945–2-2/S. verticilliflorun | 217 | Tao et al., 2018 | N |
| QGrnWgt10.6 | 10 | 79.10 | 80.87 | R931945–2-2/S. verticilliflorun | 217 | Tao et al., 2018 | N |
