## Supplemental Table 6 for "Large-scale GWAS in sorghum reveals common genetic control of grain size among cereals"

Table S6 Lis of reported grain size genes in rice and maize

| Species | Gene name | Encoded protein | Methods | Gene ID | Reference |
|---|---|---|---|---|---|
| Maize | Bt2 | AGP small subunit | Transgenic experiment | Zm00001d050032 | Jiang et al. 2013 |
| Maize | CYP724B3 | CYP724B3 | Transgenic experiment | Zm00001d003349 | Wu et al. 2008 |
| Maize | GbssIIa | UDP-Glycosyltransferase superfamily protein | Transgenic experiment | Zm00001d019479 | Jiang et al. 2013 |
| Maize | Gln-4 | Glutamine synthetase isoenzymes | Mutant analysis | Zm00001d051804 | Martin et al. 2006 |
| Maize | Mn1 | NA | Mutant analysis | Zm00001d003776 | Miller 1992 |
| Maize | O1 | myosin XI motor protein | Mutant analysis | Zm00001d052110 | Wang et al. 2012 |
| Maize | O2 | Regulatory protein opaque-2 | Mutant analysis | Zm00001d018971 | Hartings et al. 1989 |
| Maize | PBF1 | endosperm-specific transcription factor | NIL analysis | Zm00001d005100 | Lang et al. 2014 |
| Maize | SbeI | Uncharacterized protein | Transgenic experiment | Zm00001d036361 | Jiang et al. 2013 |
| Maize | SbeIIb | 1,4-alpha-glucan-branching enzyme | Transgenic experiment | Zm00001d016684 | Jiang et al. 2013 |
| Maize | Sh1 | Sucrose synthase 1 | Transgenic experiment | Zm00001d045042 | Jiang et al. 2013 |
| Maize | Sh2 | AGP large subunit | tant analysis/Transgenic experir | Zm00001d044129 | Jiang et al. 2013 |
| Maize | SMK1 | pentatricopeptide repeat protein | Mutant analysis | Zm00001d007100 | Li et al. 2014 |
| Maize | ZmIPT2 | cytokinin biosynthetic enzyme | Association study | Zm00001d003869 | Weng et al. 2013 |
| Maize | ZmSWEET4c | Transepithelial hexose transporter | Mutant analysis | Zm00001d015912 | Sosso et al. 2015 |
| Rice | DEP2 | novel plant-specific protein | QTL Cloning | Os07g0616000 | Li et al. 2010 |
| Rice | An-1 | a basic helix-loop-helix protein | QTL Cloning | Os04g0350700 | Luo et al. 2013 |
| Rice | APG | Typical bHLH protein | Transgenic experiment | Os05g0139100 | Heang and Sassa 2012 |
| Rice | BC14 | Golgi-localized nucleotide sugar transporters | Mutant analysis | Os02g0614100 | Zhang et al. 2011 |
| Rice | BG1 | Primary auxin response gene | Mutant analysis | Os03g0175800 | Liu et al. 2015 |
| Rice | BG2 | Cytochrome P450 | Transgenic experiment | Os07g0603700 | Xu et al. 2015 |
| Rice | *BG3* | a plasma membrane transporter of cytokinin | Mutant analysis | Os01g0680200 | Xiao et al. 2018 |
| Rice | BRD1 | Brassinosteroid-6-oxidase | Mutant analysis | Os03g0602300 | Mori et al. 2002 |
| Rice | BRD2 | homologous to Arabidopsis DIM1/DWF1 | Mutant analysis | Os10g0397400 | Hong et al. 2005 |
| Rice | Bu1 | a helix-loop-helix protein | Transgenic experiment | Os06g0226500 | Tanaka et al. 2009 |
| Rice | Cyc-T1;3 | Cyclin-T1;3 | Transgenic experiment | Os11g0157100 | Qi et al. 2012 |
| Rice | CYP704A3 | a putative cytochrome P450 | Transgenic experiment | Os04g0573900 | Tang et al. 2016 |
| Rice | CYP90B2 | CYP90B2 | Transgenic experiment | Os03g0227700 | Wu et al. 2008 |
| Rice | D11 | Novel Cytochrome P450 | Mutant analysis | Os04g0469800 | Tanabe et al. 2005 |
| Rice | D2 | CYP90D2 | Mutant analysis | Os01g0197100 | Hong et al. 2003 |
| Rice | D61 | OsBRI1 | Mutant analysis | Os01g0718300 | Morinaka et al. 2006 |
| Rice | DEP1 | PEBP-like domain protein | QTL Cloning | Os09g0441900 | Huang et al. 2009 |
| Rice | DEP3 | Patatin-like phospholipase A2 | Association study | Os06g0677000 | Qiao et al. 2011 |
| Rice | DLT | GRAS family protein | Transgenic experiment | Os06g0127800 | Tong et al. 2012 |
| Rice | FLO2 | Protein with a tetratricopeptide repeat motif | Mutant analysis | Os04g0645100 | She et al. 2010 |
| Rice | FUWA | NHL domain-containing protein | Transgenic experiment | Os02g0234200 | Chen et al. 2015 |
| Rice | GAD1 | DERMAL PATTERNING FAC- TOR-LIKE pep | QTL Cloning | Os08g0485500 | Jin et al. 2016 |
| Rice | *GGC2* | Gγ protein | Transgenic experiment | Os08g0456600 | Sun et al. 2018 |
| Rice | GIF1 | Cell-wall invertase | Mutant analysis | Os04g0413500 | Wang et al. 2008 |
| Rice | GL3.1/qGL3 | Phosphatase with Kelch-like repeat domain | QTL Cloning | Os03g0646900 | Qi et al. 2012; Zhang 2012 |
| Rice | GL4 | Myb-like protein | QTL Cloning | ORGLA04G025430C | Wu et al. 2017 |
| Rice | GLW7 | Plant-specific transcription factor OsSPL13 | QTL Cloning | Os07g0505200 | Si et al. 2016 |
| Rice | Grain Length3.2 | Cytochrome P450 | Transgenic experiment | Os03g0417700 | Xu et al. 2015 |
| Rice | GS2/GL2 | Growth regulating factor 4 | QTL Cloning | Os02g0701300 | Che et al. 2015; Hu et al. 2015 |

| | | | | | |
|---|---|---|---|---|---|
| Rice | GS3 | Trans-membrane protein | QTL Cloning | OS03G0407400 | Mao et al. 2010 |
| Rice | GS5 | Putative serine carboxypeptidase | QTL Cloning | Os05g0158500 | Li et al. 2011 |
| Rice | GS9 | an unknown expressed protein | QTL Cloning | LOC_Os09g27590 | Zhao et al. 2018 |
| Rice | GSK2 | signaling kinase | Transgenic experiment | Os05g0207500 | Tong et al. 2012 |
| Rice | *gsn1/LARGE8* | OsMKP1 | Mutant analysis | Os05g0115800 | Guo et al. 2018; Xu etal., 2018 |
| Rice | GW2 | RING-type E3 ubiquitin ligase | QTL Cloning | Os02g0244100 | Song et al. 2007 |
| Rice | GW5/qSW5 | calmodulin binding protein | QTL Cloning | Os05g0187500 | Liu et al. 2017 |
| Rice | GW6 | GNAT-like protein | QTL | Os06g0650300 | Song et al. 2015 |
| Rice | GW8/OsPL16 | Squamosa promoter-binding protein like 16 | QTL Cloning | Os08g0531600 | Wang et al. 2012 |
| Rice | HGW | ubiquitin-associated domain protein | Mutant analysis | Os06g0160400 | Li et al. 2012 |
| Rice | MHZ7 | a membrane protein homologous to EIN2 | Mutant analysis | Os07g0155600 | Ma et al. 2013 |
| Rice | IYB transcription fa | A MYB transcription factor | Association study | Os07G0497500 | Huang et al. 2012 |
| Rice | A Zinc finger protei | A Zinc finger protein | Association study | Os02G0192300 | Huang et al. 2012 |
| Rice | A Expressed protei | A Expressed protein | Association study | Os03G0604566 | Huang et al. 2012 |
| Rice | A transport protein | A transport protein | Association study | Os03G0604600 | Huang et al. 2012 |
| Rice | A Expressed protei | A Expressed protein | Association study | Os03G0574600 | Huang et al. 2012 |
| Rice | Nuf2 family protei | A Nuf2 family protein | Association study | Os03G0577100 | Huang et al. 2012 |
| Rice | Nuf2 family protei | A Nuf2 family protein | Association study | Os03G0577100 | Huang et al. 2012 |
| Rice | Nuf2 family protei | A Nuf2 family protein | Association study | Os03G0577100 | Huang et al. 2012 |
| Rice | A GASR7 | A GASR7 | Association study | Os06G0266800 | Huang et al. 2012 |
| Rice | A transcription facto | A transcription factor | Association study | Os06G0265400 | Huang et al. 2012 |
| Rice | Areceptor-like kinas | Areceptor-like kinase | Association study | Os07G0501800 | Huang et al. 2012 |
| Rice | OsAGSW1 | Chloroplast-localized ABC1 protein kinase | Transgenic experiment | Os05g0323800 | Li et al. 2015 |
| Rice | *OsARF4* | an auxin response factor (ARF) family protein | Transgenic experiment | Os01g0927600 | Hu et al. 2018 |
| Rice | OsBAK1 | Leucine-rich repeat receptor-like kinase | Mutant analysis | Os08g0174700 | Yuan et al. 2017 |
| Rice | OsBUL1 | bHLH transcriptional activator | Transgenic experiment | Os02g0747900 | Jang and Li 2017 |
| Rice | OsBZR1 | Transcription factor mediating BRresponses | Transgenic experiment | Os07g0580500 | Zhu et al. 2015 |
| Rice | OsCCS52B | rice cell cycle switch 52 B | Mutant analysis | Os01g0972900 | Su'udi et al. 2012 |
| Rice | OsFBK12 | F-box protein containing a Kelch repeat motif | Transgenic experiment | Os03g0171600 | Chen et al. 2013 |
| Rice | OsFIE1 | Putative Polycomb group protein FIE2 | Transgenic experiment | Os08g0137250 | Folsom et al. 2014 |
| Rice | OsFIE2 | Putative Polycomb group protein FIE2 | Transgenic experiment | Os08g0137100 | Na et al. 2012 |
| Rice | OsGIF1 | Rice GRF-interacting protein 1 | Transgenic experiment | Os03g0733600 | He et al. 2017 |
| Rice | OsMAPK6 | Mitogen-activated protein kinase 6 | Mutant analysis | Os06g0154500 | Liu et al. 2015b |
| Rice | *OsMKK4* | MKK4 | Mutant analysis | Os02g0787300 | Xu etal., 2018 |
| Rice | *OsMKKK10* | MKKK10 | Mutant analysis | Os04g0559800 | Xu etal., 2018 |
| Rice | *OsMPK6* | MPK6 | Mutant analysis | Os10g0533600 | Guo et al. 2018 |
| Rice | OsPPKL2 | Phosphatase with Kelch-like repeat domain | Transgenic experiment | Os05g0144400 | Zhan et al. 2012 |
| Rice | OsPPKL3 | Phosphatase with Kelch-like repeat domain | Transgenic experiment | Os12g0617900 | Zhan et al. 2012 |
| Rice | *ospup7* | a plasma membrane transporter of cytokinin | Transgenic experiment | Os05g0556800 | Xiao et al. 2018 |
| Rice | OsSAMS1 | -ADENOSYL-L-METHIONINE SYNTHETASI | Transgenic experiment | Os05g0135700 | Chen et al. 2013 |
| Rice | OsSGL | DUF1645 family protein | Transgenic experiment | Os02g0134200 | Wang et al. 2016 |
| Rice | *OsSPMS1* | a SPMS encoding gene | Transgenic experiment | Os06g0528600 | Tao etal. 2018 |
| Rice | OsSUT2 | Sucrose transporter | Mutant analysis | Os12g0641400 | Eom et al. 2011 |
| Rice | PGL1 | Atypical bHLH protein | Transgenic experiment | Os03g0171300 | Heang and Sassa 2012 |
| Rice | PGL2 | Atypical bHLH protein | Transgenic experiment | Os02g0747900 | Heang and Sassa 2012b |
| Rice | qGL3-2/qLGY3 | MADS-box transcription factor 1 | QTL Cloning | Os03g0215400 | Liu etal. 2018; Yu etal., 2018 |
| Rice | qGW7/GL7 | TONNEAU1-recruiting motif protein | QTL Cloning | LOC_Os07g41200 | Wang et al. 2015a; Wang et al. 2015b |

| | | | | | |
|---|---|---|---|---|---|
| Rice | *TGW3/GL3.3/TGW* | GSK3/SHAGGY-Like Kinase OsGSK5/OsSK41 | QTL Cloning | Os03g0841800 | u et a. 2018; Xiao et al. 2018; Ying et al. 201 |
| Rice | RGA1/D1 | heterotrimeric G protein α subunit | Mutant analysis | Os05g0333200 | Ashikari 1999 |
| Rice | RGB1 | G protein β subunit | Transgenic experiment | Os03g0669100 | Utsunomiya et al. 2011 |
| Rice | RSR1 | RICE STARCH REGULATOR1 | Mutant analysis | Os05g0121600 | Fu and Xue 2010 |
| Rice | SDG725 | H3K36 methyltransferase | Transgenic experiment | Os02g0554000 | Sui et al. 2012 |
| Rice | SERF1 | transcription factor SALT-RESPONSIVE ERF1 | Mutant analysis | Os05g0420300 | Schmidt et al. 2013 |
| Rice | SG1 | Novel protein | Mutant analysis | Os09g0459200 | Nakagawa et al. 2012 |
| Rice | SGL1 | Novel protein | Transgenic experiment | OS02G0762600 | Nakagawa et al. 2012 |
| Rice | SLG | BAHD acyltransferase-like protein | Mutant analysis | Os08g0562500 | Feng et al. 2016 |
| Rice | SMG1 | mitogen-activated protein kinase kinase 4 | Mutant analysis | Os02g0787300 | Duan et al. 2014 |
| Rice | SMOS1 | ıusual APETALA2 (AP2)-type tran- scription fac | Mutant analysis | Os05g0389000 | Hirano et al. 2017 |
| Rice | SRS3 | Kinesin 13 protein | Mutant analysis | Os05g0154700 | Kitagawa et al. 2010 |
| Rice | SRS5 | Alpha-tubulin protein | Mutant analysis | Os11g0247300 | Segami 2012 |
| Rice | TGW6 | Iindole-3-acetic acid (IAA)-glucose hydrolase | QTL Cloning | Os06g0623700 | Ishimaru et al. 2013 |
| Rice | TH1 | a DUF640 domain-like gene | Mutant analysis | Os02g0811000 | Li et al. 2012c |
| Rice | TIFY 11b | TIFY gene | Transgenic experiment | Os03g0181100 | Hakata et al. 2012 |
| Rice | TUD1 | U-box E3 ubiquitin ligase | Mutant analysis | Os03g0232600 | Hu et al. 2013 |
| Rice | WTG1 | Deubiquitinating enzyme | Mutant analysis | Os08g0537800 | Wang et al. 2017 |
