## Supplemental Table 7 for "Large-scale GWAS in sorghum reveals common genetic control of grain size among cereals"

Table S7 Effect of population structure on grain size

| Racial groups | TKW | Volume | Length | Width | Thickness |
|---|---|---|---|---|---|
| E African durras | 20.03[a] | 21.10[a] | 4.06[a] | 3.11[a] | 3.19[a] |
| Asian durras | 25.57[b] | 24.11[b] | 4.24[b] | 3.37[b] | 3.20[a] |
| Kafir | 27.47[b] | 25.59[b] | 4.42[c] | 3.41[b] | 3.23[a] |
| Guinea | 31.22[c] | 27.98[c] | 4.71[d] | 3.58[c] | 3.16[a] |
| Caudatum | 32.16[c] | 27.84[c] | 4.50[c] | 3.62[c] | 3.24[ab] |
