## Supplemental Table 8 for "Large-scale GWAS in sorghum reveals common genetic control of grain size among cereals"

Table S8 Marker Trait associations in the diversity panel

| SNP | Chr | Position | Predicted cM | Length | Thickness | TKW | Volume | Width | PC1 | PC2 | PC3 |
|---|---|---|---|---|---|---|---|---|---|---|---|
| M000615 | Chr01 | 1197378 | 3.18 | 2.54E-06 | 4.14E-01 | 3.27E-01 | 2.19E-01 | 5.47E-03 | 4.62E-07 | 4.89E-01 | 2.22E-01 |
| M002136 | Chr01 | 3551559 | 10.35 | 9.30E-01 | 2.98E-01 | 1.17E-01 | 9.01E-07 | 3.12E-01 | 7.52E-01 | 1.33E-01 | 1.74E-01 |
| M002601 | Chr01 | 4235340 | 10.90 | 8.61E-13 | 8.36E-01 | 4.21E-02 | 1.09E-01 | 2.36E-02 | 4.70E-06 | 2.50E-01 | 3.19E-01 |
| M003249 | Chr01 | 5405218 | 12.65 | 1.21E-01 | 2.56E-01 | 2.95E-01 | 8.38E-07 | 3.01E-02 | 2.38E-01 | 1.68E-02 | 6.34E-03 |
| M003474 | Chr01 | 5800441 | 13.43 | 8.68E-01 | 7.40E-01 | 2.85E-01 | 1.26E-02 | 1.86E-05 | 2.37E-01 | 5.83E-04 | 3.55E-01 |
| M003550 | Chr01 | 5918322 | 13.66 | 6.73E-02 | 1.41E-02 | 1.34E-02 | 5.65E-01 | 9.33E-01 | 4.35E-02 | 3.52E-01 | 2.02E-08 |
| M004378 | Chr01 | 7312772 | 18.27 | 3.61E-01 | 2.36E-01 | 1.95E-08 | 4.12E-01 | 2.82E-02 | 1.84E-01 | 1.26E-03 | 4.05E-02 |
| M005704 | Chr01 | 9747369 | 25.45 | 2.80E-01 | 6.14E-10 | 2.65E-02 | 4.90E-01 | 2.20E-01 | 2.55E-01 | 2.37E-02 | 2.06E-03 |
| M005731 | Chr01 | 9819334 | 25.48 | 9.91E-01 | 8.89E-01 | 1.28E-03 | 1.78E-01 | 1.91E-01 | 3.22E-01 | 5.09E-06 | 6.29E-02 |
| M007294 | Chr01 | 12613599 | 31.25 | 4.61E-02 | 6.89E-02 | 2.32E-03 | 7.68E-07 | 3.59E-03 | 6.14E-01 | 2.48E-02 | 2.82E-01 |
| M007595 | Chr01 | 13193540 | 33.54 | 4.77E-01 | 6.71E-01 | 6.43E-02 | 9.06E-02 | 8.27E-02 | 7.35E-01 | 2.72E-01 | 1.15E-05 |
| M009214 | Chr01 | 17427840 | 43.49 | 2.37E-01 | 1.89E-01 | 1.68E-07 | 2.81E-01 | 1.15E-01 | 6.86E-02 | 6.15E-02 | 4.53E-01 |
| M009345 | Chr01 | 17638047 | 43.60 | 1.71E-05 | 5.17E-01 | 1.40E-01 | 1.51E-01 | 1.32E-01 | 1.11E-02 | 5.07E-02 | 2.28E-01 |
| M010716 | Chr01 | 21347404 | 46.68 | 2.45E-03 | 7.96E-01 | 1.30E-05 | 3.31E-06 | 1.99E-02 | 1.46E-05 | 3.61E-01 | 1.26E-01 |
| M013480 | Chr01 | 53202019 | 59.15 | 9.29E-01 | 3.81E-08 | 1.03E-02 | 6.11E-02 | 2.79E-01 | 4.58E-01 | 2.26E-01 | 1.12E-01 |
| M016956 | Chr01 | 62899310 | 94.76 | 5.84E-06 | 4.17E-01 | 1.47E-01 | 2.38E-04 | 5.38E-04 | 1.80E-01 | 1.41E-02 | 1.63E-01 |
| M019957 | Chr01 | 68372498 | 125.38 | 7.91E-01 | 2.53E-10 | 2.51E-02 | 3.52E-02 | 2.69E-01 | 1.17E-01 | 3.40E-01 | 3.48E-01 |
| M020793 | Chr01 | 69804026 | 129.89 | 2.87E-02 | 2.84E-09 | 9.96E-01 | 6.57E-01 | 1.72E-01 | 1.24E-01 | 4.21E-01 | 8.57E-02 |
| M037788 | Chr02 | 57512339 | 93.17 | 3.23E-01 | 3.16E-08 | 9.16E-03 | 3.35E-02 | 2.24E-01 | 1.42E-01 | 3.41E-02 | 3.12E-05 |
| M038579 | Chr02 | 59194781 | 106.68 | 3.79E-01 | 2.07E-01 | 1.34E-02 | 3.60E-08 | 1.92E-03 | 2.42E-02 | 1.68E-02 | 8.06E-02 |
| M042976 | Chr02 | 66573634 | 149.23 | 1.99E-01 | 8.70E-01 | 1.89E-03 | 2.44E-02 | 9.66E-06 | 4.46E-02 | 3.62E-02 | 8.14E-01 |
| M043684 | Chr02 | 67673733 | 152.63 | 9.82E-01 | 1.62E-03 | 5.27E-01 | 8.52E-01 | 3.46E-01 | 7.91E-01 | 1.77E-01 | 6.31E-06 |
| M045553 | Chr02 | 70379906 | 166.28 | 8.07E-07 | 4.22E-01 | 3.14E-01 | 7.21E-04 | 6.95E-03 | 3.96E-02 | 1.38E-02 | 9.11E-01 |
| M047672 | Chr02 | 73588718 | 181.30 | 2.32E-03 | 5.45E-02 | 5.64E-02 | 5.42E-02 | 1.02E-02 | 1.74E-06 | 3.79E-01 | 1.52E-01 |
| M052341 | Chr03 | 2972967 | 9.19 | 1.77E-01 | 1.67E-01 | 9.44E-02 | 1.11E-03 | 2.06E-07 | 6.47E-02 | 1.37E-01 | 1.01E-01 |
| M052809 | Chr03 | 3597468 | 11.06 | 6.16E-01 | 3.04E-01 | 6.23E-08 | 7.26E-02 | 7.43E-01 | 6.42E-03 | 7.91E-02 | 4.22E-03 |
| M055343 | Chr03 | 7459293 | 31.12 | 4.81E-01 | 2.44E-01 | 8.98E-01 | 8.98E-01 | 7.48E-01 | 3.05E-01 | 7.76E-01 | 1.90E-08 |
| M057478 | Chr03 | 11842434 | 46.52 | 2.42E-06 | 3.61E-01 | 4.99E-03 | 1.86E-02 | 1.30E-01 | 4.25E-03 | 1.58E-01 | 2.95E-01 |
| M057498 | Chr03 | 11867983 | 46.57 | 8.04E-02 | 8.33E-02 | 4.24E-02 | 2.78E-01 | 2.02E-05 | 5.04E-02 | 7.57E-02 | 3.27E-01 |
| M062129 | Chr03 | 54907966 | 70.31 | 7.34E-02 | 4.75E-01 | 1.28E-08 | 2.23E-05 | 2.09E-05 | 2.40E-03 | 6.78E-02 | 1.61E-05 |
| M076774 | Chr04 | 4559812 | 29.99 | 3.12E-02 | 8.40E-01 | 7.88E-03 | 3.17E-05 | 1.64E-06 | 6.04E-07 | 4.42E-02 | 6.73E-01 |
| M078413 | Chr04 | 7553709 | 55.81 | 1.51E-01 | 2.16E-08 | 2.55E-01 | 7.29E-01 | 5.42E-01 | 2.08E-01 | 8.04E-01 | 2.92E-03 |
| M080340 | Chr04 | 13834059 | 71.37 | 8.17E-02 | 7.43E-06 | 5.30E-01 | 2.90E-01 | 6.44E-01 | 9.10E-01 | 4.85E-01 | 4.00E-01 |
| M082318 | Chr04 | 49255658 | 73.60 | 2.00E-02 | 3.57E-01 | 5.32E-03 | 2.04E-01 | 9.27E-02 | 1.41E-08 | 3.31E-01 | 6.15E-01 |
| M083044 | Chr04 | 51716417 | 83.25 | 7.09E-01 | 5.70E-01 | 1.73E-01 | 6.81E-01 | 6.36E-01 | 6.42E-01 | 1.78E-01 | 8.04E-06 |
| M083997 | Chr04 | 54000525 | 94.49 | 3.06E-01 | 1.24E-01 | 6.75E-06 | 9.95E-05 | 6.20E-04 | 1.90E-02 | 3.28E-02 | 1.33E-01 |
| M088370 | Chr04 | 61446722 | 122.03 | 1.77E-02 | 2.27E-01 | 1.90E-05 | 1.10E-04 | 1.35E-01 | 7.37E-07 | 1.39E-01 | 2.79E-01 |
| M091666 | Chr04 | 66127237 | 152.27 | 1.28E-07 | 6.22E-01 | 6.09E-02 | 4.64E-03 | 1.25E-01 | 2.06E-04 | 6.11E-03 | 2.11E-01 |
| M095512 | Chr05 | 3575186 | 23.25 | 9.27E-01 | 5.80E-02 | 1.38E-06 | 2.07E-01 | 4.98E-01 | 6.74E-01 | 2.46E-01 | 2.76E-01 |
| M100240 | Chr05 | 38548979 | 64.58 | 4.45E-01 | 2.94E-01 | 9.37E-02 | 8.23E-02 | 6.23E-06 | 8.52E-03 | 2.88E-01 | 6.61E-02 |
| M101012 | Chr05 | 54107794 | 65.76 | 1.86E-01 | 3.49E-01 | 1.45E-02 | 1.61E-05 | 1.89E-01 | 1.77E-02 | 5.18E-01 | 4.68E-01 |
| M102450 | Chr05 | 61218281 | 70.70 | 5.95E-07 | 4.41E-01 | 4.24E-01 | 1.83E-02 | 8.41E-03 | 3.93E-03 | 3.95E-01 | 8.40E-02 |
| M103195 | Chr05 | 63339533 | 73.89 | 2.26E-06 | 2.15E-01 | 2.40E-02 | 9.26E-03 | 2.61E-01 | 1.78E-02 | 8.05E-02 | 4.63E-01 |
| M107294 | Chr05 | 71211474 | 119.00 | 1.45E-04 | 4.11E-01 | 6.94E-07 | 7.55E-02 | 3.98E-03 | 3.30E-01 | 1.02E-01 | 9.13E-01 |

| | | | | | | | | | | | |
|---|---|---|---|---|---|---|---|---|---|---|---|
| M109041 | Chr06 | 3594243 | 33.68 | 1.31E-01 | 1.04E-03 | 5.99E-01 | 4.74E-01 | 9.54E-01 | 4.45E-01 | 3.14E-01 | 7.04E-06 |
| M118469 | Chr06 | 55406074 | 125.64 | 2.83E-01 | 3.34E-01 | 1.82E-01 | 6.78E-01 | 8.00E-01 | 7.72E-01 | 9.58E-01 | 1.11E-05 |
| M123536 | Chr07 | 1463227 | 16.76 | 2.00E-04 | 1.78E-01 | 3.15E-01 | 1.71E-01 | 3.36E-01 | 6.28E-08 | 6.12E-01 | 1.17E-01 |
| M125276 | Chr07 | 4810904 | 58.13 | 7.39E-02 | 3.94E-02 | 2.99E-01 | 7.10E-06 | 5.36E-02 | 1.53E-03 | 3.19E-01 | 4.34E-01 |
| M128159 | Chr07 | 24798629 | 72.25 | 3.27E-01 | 7.33E-03 | 9.80E-01 | 6.76E-03 | 8.08E-07 | 3.17E-02 | 4.19E-01 | 1.38E-02 |
| M128561 | Chr07 | 38985304 | 72.30 | 4.25E-01 | 7.03E-01 | 1.08E-01 | 1.25E-01 | 2.80E-05 | 1.94E-03 | 3.46E-01 | 5.15E-01 |
| M129315 | Chr07 | 52822173 | 74.35 | 1.33E-06 | 6.01E-03 | 5.02E-01 | 1.26E-01 | 5.50E-01 | 5.48E-02 | 5.89E-02 | 1.48E-01 |
| M129967 | Chr07 | 55675216 | 78.67 | 6.41E-01 | 3.88E-01 | 1.23E-02 | 5.64E-05 | 1.53E-08 | 9.56E-02 | 1.16E-01 | 4.00E-02 |
| M132056 | Chr07 | 60263992 | 106.63 | 9.92E-10 | 2.94E-02 | 5.12E-01 | 8.43E-01 | 1.93E-01 | 6.05E-01 | 4.78E-03 | 2.68E-01 |
| M143723 | Chr08 | 58069324 | 87.75 | 8.54E-01 | 8.02E-07 | 7.57E-01 | 8.38E-01 | 9.80E-01 | 7.94E-01 | 3.02E-01 | 5.57E-02 |
| M145080 | Chr08 | 60306822 | 102.58 | 2.25E-03 | 7.93E-01 | 2.83E-01 | 7.62E-05 | 1.67E-04 | 6.60E-07 | 1.84E-01 | 8.09E-02 |
| M146586 | Chr08 | 62329307 | 127.16 | 6.46E-02 | 9.36E-01 | 2.34E-02 | 2.52E-01 | 5.18E-02 | 9.30E-07 | 3.98E-02 | 8.04E-02 |
| M147523 | Chr09 | 1188274 | 14.45 | 3.67E-01 | 3.53E-04 | 1.15E-06 | 9.13E-02 | 3.27E-03 | 7.77E-02 | 5.98E-01 | 3.60E-03 |
| M151212 | Chr09 | 8955726 | 67.98 | 4.90E-08 | 7.57E-01 | 8.25E-02 | 4.26E-02 | 5.01E-02 | 2.64E-03 | 2.97E-01 | 1.86E-01 |
| M151613 | Chr09 | 11497840 | 68.55 | 8.04E-06 | 9.43E-01 | 8.70E-01 | 8.38E-03 | 8.32E-01 | 2.72E-06 | 2.48E-01 | 2.41E-01 |
| M153043 | Chr09 | 42995650 | 75.64 | 8.10E-02 | 1.24E-09 | 2.52E-01 | 8.31E-01 | 8.17E-01 | 8.74E-01 | 6.16E-01 | 5.54E-02 |
| M153348 | Chr09 | 45347219 | 76.17 | 2.46E-01 | 1.32E-06 | 6.50E-01 | 2.16E-01 | 5.55E-01 | 5.65E-01 | 3.73E-01 | 2.76E-01 |
| M156558 | Chr09 | 52757898 | 104.82 | 4.42E-01 | 2.67E-06 | 2.44E-01 | 4.50E-01 | 6.95E-01 | 6.19E-01 | 2.37E-01 | 1.05E-02 |
| M156618 | Chr09 | 52795485 | 104.93 | 3.24E-02 | 9.74E-01 | 1.51E-05 | 1.33E-01 | 7.02E-01 | 2.49E-01 | 5.86E-01 | 7.23E-01 |
| M158516 | Chr09 | 56099639 | 113.99 | 6.44E-07 | 9.78E-01 | 8.58E-01 | 3.21E-01 | 7.39E-01 | 1.78E-01 | 2.49E-01 | 5.76E-01 |
| M161240 | Chr10 | 777136 | 13.22 | 3.69E-01 | 2.24E-07 | 1.15E-01 | 4.28E-01 | 1.66E-01 | 9.18E-01 | 1.06E-02 | 4.60E-02 |
| M162890 | Chr10 | 3445247 | 32.55 | 7.41E-01 | 3.13E-02 | 4.68E-12 | 1.56E-04 | 7.05E-08 | 5.04E-02 | 1.40E-02 | 8.65E-04 |
| M167660 | Chr10 | 18074250 | 57.96 | 2.19E-01 | 1.03E-18 | 5.11E-02 | 7.72E-01 | 3.64E-01 | 5.90E-01 | 1.83E-02 | 2.59E-03 |
| M168658 | Chr10 | 40511389 | 58.58 | 3.45E-06 | 3.15E-01 | 8.05E-01 | 1.67E-02 | 8.73E-03 | 3.27E-01 | 6.42E-02 | 4.45E-01 |
| M174724 | Chr10 | 58653874 | 102.30 | 8.38E-01 | 2.39E-01 | 4.03E-07 | 4.36E-06 | 1.56E-05 | 8.93E-01 | 1.54E-05 | 2.14E-03 |
| M175480 | Chr10 | 60146528 | 103.55 | 3.58E-01 | 7.79E-01 | 8.17E-06 | 1.39E-03 | 6.12E-03 | 2.39E-02 | 2.10E-01 | 2.37E-01 |
| M175501 | Chr10 | 60172927 | 103.56 | 6.02E-01 | 8.52E-01 | 5.40E-08 | 8.74E-01 | 1.28E-01 | 2.91E-01 | 4.18E-01 | 3.13E-01 |
