## Supplemental Table 9 for "Large-scale GWAS in sorghum reveals common genetic control of grain size among cereals"

Table S9 Effects of QTL in the diversity panel

| QTL ID | Chr | start | end | Traits affected in DP | | | | | $r^2$ | | | | |
|---|---|---|---|---|---|---|---|---|---|---|---|---|---|
| | | | | TKW | Volume | Length | Width | Thickness | TKW | Volume | Length | Width | Thickness |
| qGS1.1 | 1 | 3.18 | 3.18 | N | Y | Y | N | N | NA | 8.20E-03 | 2.38E-02 | NA | NA |
| qGS1.2 | 1 | 10.35 | 13.66 | Y | Y | Y | Y | Y | 5.14E-02 | 3.84E-02 | 1.91E-02 | 3.03E-02 | 4.28E-02 |
| qGS1.3 | 1 | 16.89 | 18.27 | Y | Y | Y | Y | Y | 5.44E-02 | 3.26E-02 | 1.98E-02 | 2.47E-02 | 2.51E-02 |
| qGS1.4 | 1 | 25.45 | 25.48 | Y | Y | Y | Y | Y | 1.33E-02 | 1.62E-02 | 1.25E-02 | 6.84E-03 | 2.85E-02 |
| qGS1.5 | 1 | 31.25 | 33.54 | N | Y | N | N | N | NA | 2.64E-03 | NA | NA | NA |
| qGS1.6 | 1 | 43.49 | 43.60 | Y | Y | Y | Y | N | 2.00E-02 | 2.31E-02 | 3.23E-02 | 1.47E-02 | NA |
| qGS1.7 | 1 | 46.68 | 46.73 | Y | Y | Y | Y | N | 2.86E-02 | 1.33E-02 | 1.47E-02 | 1.65E-02 | NA |
| qGS1.8 | 1 | 59.15 | 60.07 | Y | Y | N | N | Y | 8.80E-03 | 8.33E-03 | NA | NA | 2.42E-02 |
| qGS1.9 | 1 | 94.75 | 94.76 | Y | Y | Y | Y | N | 1.64E-02 | 3.01E-02 | 2.91E-02 | 2.72E-02 | 2.46E-03 |
| qGS1.12 | 1 | 123.06 | 125.38 | N | N | N | N | Y | NA | NA | NA | NA | 1.65E-02 |
| qGS1.13 | 1 | 129.89 | 129.89 | N | N | N | N | Y | NA | NA | NA | NA | 4.20E-03 |
| qGS2.6 | 2 | 93.17 | 93.17 | Y | Y | N | Y | Y | 9.91E-03 | 1.20E-02 | NA | 9.60E-03 | 2.11E-02 |
| qGS2.7 | 2 | 106.68 | 106.68 | N | Y | Y | N | N | NA | 7.95E-03 | 1.23E-02 | NA | NA |
| qGS2.10 | 2 | 148.73 | 149.23 | N | N | N | Y | N | NA | NA | NA | 2.71E-03 | NA |
| qGS2.11 | 2 | 152.32 | 152.63 | N | N | Y | N | N | NA | NA | 7.22E-03 | NA | NA |
| qGS2.13 | 2 | 166.28 | 166.28 | Y | Y | Y | Y | N | 1.26E-02 | 1.68E-02 | 1.25E-02 | 1.81E-02 | NA |
| qGS2.14 | 2 | 180.56 | 182.44 | Y | Y | Y | Y | Y | 3.33E-02 | 3.18E-02 | 1.70E-02 | 1.92E-02 | 1.93E-02 |
| qGS3.3 | 3 | 9.19 | 9.19 | Y | Y | N | Y | N | 9.35E-03 | 9.76E-03 | NA | 1.91E-02 | NA |
| qGS3.4 | 3 | 11.06 | 11.06 | Y | Y | Y | Y | Y | 2.56E-02 | 1.36E-02 | 7.52E-03 | 2.32E-02 | 7.36E-03 |
| qGS3.5 | 3 | 31.12 | 31.12 | Y | Y | N | Y | N | 1.22E-02 | 6.81E-03 | 4.83E-03 | 1.38E-02 | NA |
| qGS3.6 | 3 | 46.52 | 48.31 | Y | Y | Y | Y | N | 1.72E-02 | 2.36E-02 | 1.49E-02 | 3.11E-02 | NA |
| qGS3.8 | 3 | 70.31 | 72.25 | Y | Y | Y | Y | Y | 5.30E-02 | 4.12E-02 | 1.95E-02 | 3.59E-02 | 3.89E-02 |
| qGS4.2 | 4 | 29.99 | 29.99 | Y | Y | Y | Y | Y | 1.84E-02 | 2.42E-02 | 1.20E-02 | 2.41E-02 | 1.15E-02 |
| qGS4.3 | 4 | 55.81 | 55.81 | N | N | N | N | Y | NA | NA | NA | NA | 1.66E-02 |
| qGS4.4 | 4 | 71.37 | 73.60 | Y | Y | Y | Y | N | 1.75E-02 | 1.79E-02 | 2.85E-02 | 2.08E-02 | NA |
| qGS4.5 | 4 | 82.14 | 84.01 | N | N | N | N | Y | NA | NA | NA | NA | 8.39E-03 |
| qGS4.6 | 4 | 94.49 | 94.71 | Y | Y | Y | Y | N | 9.40E-03 | 9.56E-03 | 9.28E-03 | 1.15E-02 | NA |
| qGS4.8 | 4 | 121.78 | 123.63 | Y | Y | Y | Y | N | 1.87E-02 | 1.88E-02 | 1.80E-02 | 1.77E-02 | NA |
| qGS4.9 | 4 | 152.27 | 152.27 | Y | Y | Y | Y | N | 8.10E-03 | 7.81E-03 | 1.90E-02 | 6.66E-03 | NA |
| qGS5.1 | 5 | 22.87 | 23.25 | Y | Y | Y | Y | Y | 4.81E-02 | 3.31E-02 | 1.40E-02 | 2.77E-02 | 1.50E-02 |
| qGS5.2 | 5 | 64.58 | 65.76 | N | Y | Y | Y | N | 5.61E-03 | 6.00E-03 | 6.97E-03 | 1.30E-02 | NA |
| qGS5.3 | 5 | 70.70 | 70.70 | N | Y | Y | N | N | NA | 9.44E-03 | 1.18E-02 | NA | NA |
| qGS5.4 | 5 | 73.38 | 73.89 | Y | Y | Y | Y | Y | 2.22E-02 | 2.70E-02 | 2.47E-02 | 1.57E-02 | 2.16E-02 |
| qGS5.5 | 5 | NA | NA | Y | Y | Y | Y | N | 7.33E-03 | 1.19E-02 | 1.66E-02 | 1.09E-02 | NA |
| qGS6.2 | 6 | 32.68 | 33.68 | Y | N | N | N | Y | NA | NA | NA | NA | 9.07E-03 |
| qGS6.5 | 6 | 125.43 | 125.64 | N | N | Y | Y | Y | NA | NA | 6.38E-03 | 8.11E-03 | 1.13E-02 |
| qGS7.1 | 7 | 16.76 | 16.76 | Y | Y | Y | Y | N | 2.19E-02 | 2.66E-02 | 2.59E-02 | 2.62E-02 | NA |
| qGS7.2 | 7 | 58.13 | 58.36 | Y | Y | Y | Y | Y | 1.27E-02 | 2.38E-02 | 1.72E-02 | 1.79E-02 | 1.34E-02 |
| qGS7.3 | 7 | 72.21 | 72.30 | Y | Y | Y | Y | Y | 1.65E-02 | 1.90E-02 | 4.11E-03 | 3.51E-02 | 1.85E-02 |
| qGS7.4 | 7 | 74.35 | 75.24 | N | N | Y | Y | N | NA | NA | 9.09E-03 | 6.40E-03 | NA |
| qGS7.5 | 7 | 78.67 | 78.67 | Y | Y | Y | Y | Y | 2.27E-02 | 3.75E-02 | 1.52E-02 | 2.90E-02 | 1.48E-02 |
| qGS7.6 | 7 | 106.63 | 106.63 | Y | Y | Y | Y | N | 1.07E-02 | 2.05E-02 | 3.37E-02 | 8.38E-03 | NA |
| qGS8.3 | 8 | 87.75 | 87.75 | N | N | N | N | Y | NA | NA | NA | NA | 1.27E-02 |

| | | | | | | | | | | | | | |
|---|---|---|---|---|---|---|---|---|---|---|---|---|---|
| qGS8.5 | 8 | 102.58 | 103.12 | Y | Y | Y | Y | N | 1.49E-02 | 2.27E-02 | 2.67E-02 | 1.70E-02 | NA |
| qGS8.10 | 8 | 127.16 | 127.16 | N | N | N | N | N | NA | NA | NA | NA | NA |
| qGS9.1 | 9 | 14.45 | 14.45 | Y | Y | N | Y | Y | 8.01E-03 | 7.41E-03 | NA | 7.36E-03 | 7.15E-03 |
| qGS9.2 | 9 | 67.33 | 69.39 | Y | Y | Y | Y | N | 7.22E-03 | 1.03E-02 | 1.16E-02 | 9.39E-03 | NA |
| qGS9.3 | 9 | 75.64 | 76.17 | Y | Y | Y | Y | Y | 1.33E-02 | 1.43E-02 | 8.55E-03 | 1.02E-02 | 7.92E-03 |
| qGS9.5 | 9 | 104.82 | 104.93 | Y | N | N | N | Y | 1.46E-02 | NA | NA | NA | 3.55E-02 |
| qGS9.7 | 9 | 113.99 | 113.99 | N | Y | Y | N | N | NA | 1.15E-02 | 2.34E-02 | 5.30E-03 | NA |
| qGS10.1 | 10 | 13.22 | 13.22 | N | Y | N | N | Y | NA | 8.22E-03 | NA | NA | 2.16E-02 |
| qGS10.2 | 10 | 31.94 | 32.55 | Y | Y | Y | Y | Y | 4.81E-02 | 4.12E-02 | 1.98E-02 | 3.39E-02 | 1.50E-02 |
| qGS10.4 | 10 | 57.96 | 58.58 | Y | Y | Y | Y | Y | 3.23E-02 | 3.35E-02 | 1.20E-02 | 9.26E-03 | 6.16E-02 |
| qGS10.6 | 10 | 102.30 | 103.56 | Y | Y | Y | Y | Y | 4.97E-02 | 3.87E-02 | 1.83E-02 | 4.03E-02 | 2.05E-02 |
