## Supplemental Table 10 for "Large-scale GWAS in sorghum reveals common genetic control of grain size among cereals"

Table S10 Marker Trait associations in BC-NAM

| SNP | Chr | Position | Predicted cM | Length | TKW | Volume | Width | Thickness | PC1 | PC2 | PC3 |
|---|---|---|---|---|---|---|---|---|---|---|---|
| MN00756 | Chr01 | 5067479 | 11.98 | 5.61E-01 | 2.02E-01 | 1.84E-03 | 8.78E-01 | 5.35E-06 | 9.49E-01 | 4.82E-03 | 1.63E-01 |
| MN01078 | Chr01 | 6907667 | 16.89 | 8.75E-01 | 3.93E-05 | 1.28E-03 | 2.64E-01 | 4.12E-02 | 5.80E-01 | 6.45E-02 | 5.24E-02 |
| MN02210 | Chr01 | 12725754 | 31.69 | 5.06E-02 | 7.70E-03 | 4.39E-02 | 4.21E-02 | 9.92E-01 | 2.33E-05 | 4.43E-01 | 6.44E-01 |
| MN02253 | Chr01 | 13016060 | 32.84 | 2.99E-04 | 5.43E-02 | 2.26E-06 | 4.65E-05 | 1.48E-01 | 4.81E-01 | 6.90E-03 | 4.22E-02 |
| MN03588 | Chr01 | 21451374 | 46.73 | 6.59E-05 | 4.20E-02 | 1.70E-04 | 9.38E-02 | 3.22E-02 | 4.40E-01 | 5.49E-02 | 6.19E-01 |
| MN03592 | Chr01 | 21457153 | 46.73 | 8.32E-01 | 2.50E-01 | 7.61E-01 | 1.30E-01 | 2.18E-01 | 9.27E-05 | 4.76E-01 | 8.46E-01 |
| MN04864 | Chr01 | 53668800 | 59.74 | 3.24E-01 | 2.20E-13 | 1.06E-02 | 2.88E-01 | 6.07E-01 | 2.32E-01 | 8.23E-01 | 4.61E-01 |
| MN04865 | Chr01 | 53766098 | 59.86 | 5.42E-01 | 2.44E-01 | 2.73E-02 | 5.06E-01 | 1.37E-06 | 4.33E-01 | 7.08E-07 | 5.33E-01 |
| MN04889 | Chr01 | 53936232 | 60.07 | 3.73E-01 | 1.67E-01 | 1.40E-02 | 1.29E-05 | 8.20E-01 | 7.12E-02 | 3.77E-01 | 5.11E-01 |
| MN06097 | Chr01 | 62886536 | 94.75 | 5.54E-06 | 6.92E-02 | 1.56E-01 | 1.20E-02 | 5.58E-01 | 5.85E-03 | 4.41E-01 | 1.11E-01 |
| MN06408 | Chr01 | 64100803 | 98.57 | 5.18E-01 | 1.93E-01 | 2.84E-01 | 3.65E-01 | 6.40E-01 | 6.30E-07 | 3.56E-01 | 9.14E-02 |
| MN06928 | Chr01 | 67340819 | 115.03 | 8.07E-01 | 9.93E-01 | 6.39E-01 | 6.27E-01 | 1.79E-06 | 1.25E-02 | 7.39E-03 | 6.02E-01 |
| MN07028 | Chr01 | 67567101 | 116.76 | 5.81E-01 | 6.73E-02 | 5.11E-01 | 8.36E-01 | 4.95E-05 | 3.61E-01 | 1.62E-02 | 9.01E-01 |
| MN07029 | Chr01 | 67575793 | 116.85 | 8.15E-01 | 7.97E-02 | 3.82E-01 | 7.51E-01 | 3.20E-05 | 4.02E-01 | 1.25E-02 | 9.74E-01 |
| MN07153 | Chr01 | 68155793 | 123.06 | 5.64E-01 | 9.04E-01 | 8.43E-01 | 4.22E-01 | 3.82E-01 | 3.43E-03 | 1.92E-08 | 2.27E-01 |
| MN07204 | Chr01 | 68344410 | 125.08 | 3.80E-01 | 5.70E-01 | 9.15E-01 | 2.97E-01 | 2.78E-15 | 6.66E-03 | 8.44E-02 | 5.74E-02 |
| MN08517 | Chr01 | 75732030 | 156.66 | 4.91E-01 | 3.14E-06 | 1.71E-01 | 3.66E-01 | 4.48E-01 | 6.66E-01 | 2.22E-01 | 4.58E-01 |
| MN09955 | Chr02 | 1044797 | 0.10 | 2.53E-01 | 2.25E-01 | 1.01E-01 | 1.76E-05 | 6.49E-01 | 9.81E-01 | 5.89E-01 | 2.40E-01 |
| MN10386 | Chr02 | 2902033 | 13.86 | 8.26E-01 | 1.44E-01 | 1.47E-05 | 9.06E-02 | 9.54E-01 | 8.01E-01 | 9.31E-01 | 3.24E-01 |
| MN10400 | Chr02 | 2977717 | 13.93 | 3.67E-08 | 3.85E-02 | 4.32E-02 | 5.34E-02 | 6.58E-01 | 1.66E-05 | 8.51E-01 | 4.74E-01 |
| MN11929 | Chr02 | 10134590 | 56.44 | 8.81E-01 | 7.21E-02 | 8.91E-02 | 3.98E-01 | 1.15E-02 | 9.89E-01 | 2.18E-05 | 7.56E-02 |
| MN11994 | Chr02 | 10821630 | 59.55 | 3.06E-01 | 2.96E-06 | 1.70E-01 | 1.36E-01 | 3.24E-01 | 3.41E-02 | 2.54E-01 | 3.64E-01 |
| MN13645 | Chr02 | 57031547 | 88.94 | 6.11E-01 | 4.26E-01 | 5.97E-01 | 9.28E-01 | 1.36E-05 | 9.35E-01 | 3.40E-03 | 8.93E-01 |
| MN14324 | Chr02 | 60589262 | 121.99 | 9.84E-01 | 1.84E-01 | 1.33E-01 | 2.99E-01 | 9.77E-06 | 3.63E-01 | 3.84E-04 | 9.06E-01 |
| MN14666 | Chr02 | 61587299 | 123.92 | 3.72E-02 | 1.81E-01 | 3.14E-03 | 2.79E-01 | 3.38E-05 | 1.49E-01 | 6.28E-02 | 1.29E-01 |
| MN14870 | Chr02 | 62351035 | 134.53 | 9.37E-01 | 4.18E-02 | 6.49E-01 | 2.42E-01 | 1.27E-01 | 8.40E-01 | 4.57E-06 | 1.90E-01 |
| MN15726 | Chr02 | 66339434 | 148.73 | 8.41E-03 | 8.80E-03 | 5.90E-06 | 3.68E-03 | 8.31E-02 | 2.22E-01 | 4.36E-01 | 3.39E-01 |
| MN15918 | Chr02 | 67288480 | 152.32 | 1.42E-01 | 7.11E-03 | 4.74E-01 | 6.41E-01 | 2.63E-02 | 6.33E-06 | 5.82E-02 | 2.66E-01 |
| MN16268 | Chr02 | 69048996 | 159.86 | 9.86E-07 | 6.94E-07 | 6.44E-11 | 5.72E-10 | 4.28E-01 | 7.37E-02 | 2.12E-02 | 2.79E-04 |
| MN17175 | Chr02 | 73307212 | 180.56 | 8.35E-02 | 7.03E-01 | 2.46E-02 | 2.72E-02 | 1.37E-02 | 2.90E-05 | 6.57E-01 | 5.19E-01 |
| MN17236 | Chr02 | 73494298 | 181.05 | 8.47E-01 | 1.31E-01 | 3.90E-01 | 9.38E-01 | 3.87E-03 | 1.27E-01 | 7.42E-08 | 6.61E-01 |
| MN17477 | Chr02 | 74155670 | 182.44 | 2.57E-01 | 1.92E-05 | 5.60E-01 | 2.85E-01 | 1.98E-01 | 5.50E-08 | 8.37E-02 | 7.94E-02 |
| MN17678 | Chr02 | 75213838 | 188.18 | 7.09E-01 | 5.95E-01 | 4.16E-01 | 3.11E-01 | 3.92E-05 | 9.96E-01 | 5.60E-02 | 4.06E-01 |
| MN17695 | Chr02 | 75310779 | 189.09 | 8.34E-01 | 1.06E-01 | 1.29E-05 | 8.03E-02 | 1.57E-02 | 5.24E-01 | 9.39E-03 | 6.26E-03 |
| MN18177 | Chr03 | 617265 | 0.68 | 2.23E-01 | 3.41E-01 | 3.38E-03 | 2.74E-01 | 1.00E-01 | 1.60E-01 | 5.57E-05 | 2.85E-02 |
| MN18445 | Chr03 | 1678233 | 4.04 | 4.05E-06 | 1.86E-01 | 2.56E-01 | 1.76E-02 | 4.67E-01 | 1.29E-02 | 7.90E-02 | 1.50E-01 |
| MN18446 | Chr03 | 1678879 | 4.05 | 2.58E-01 | 1.25E-02 | 3.08E-06 | 7.00E-03 | 6.27E-01 | 3.65E-02 | 3.12E-02 | 2.57E-01 |
| MN18447 | Chr03 | 1681078 | 4.05 | 6.80E-01 | 1.10E-05 | 5.96E-01 | 1.32E-01 | 9.58E-01 | 8.97E-02 | 2.69E-01 | 8.78E-02 |
| MN21114 | Chr03 | 13099674 | 48.31 | 5.36E-05 | 1.77E-02 | 1.13E-07 | 2.93E-02 | 1.02E-02 | 8.61E-03 | 1.16E-01 | 1.47E-01 |
| MN21707 | Chr03 | 27806521 | 56.79 | 4.04E-02 | 2.24E-01 | 9.15E-02 | 5.64E-03 | 7.88E-03 | 5.14E-01 | 3.97E-05 | 5.43E-01 |
| MN22076 | Chr03 | 50560344 | 57.86 | 3.43E-01 | 7.44E-01 | 9.97E-01 | 8.14E-01 | 4.39E-06 | 6.86E-01 | 5.54E-02 | 8.50E-01 |
| MN22172 | Chr03 | 51503726 | 58.00 | 3.71E-05 | 9.74E-01 | 1.13E-02 | 4.03E-01 | 6.58E-01 | 3.46E-01 | 4.67E-01 | 6.67E-01 |
| MN45817 | Chr03 | 52373135 | 58.13 | 3.15E-01 | 1.17E-01 | 5.81E-05 | 4.95E-01 | 7.46E-01 | 3.77E-02 | 9.26E-01 | 1.93E-02 |
| MN22591 | Chr03 | 55299513 | 72.25 | 1.06E-04 | 1.63E-01 | 3.82E-06 | 1.99E-05 | 2.39E-01 | 1.45E-03 | 7.09E-01 | 4.90E-02 |

| | | | | | | | | | | | |
|---|---|---|---|---|---|---|---|---|---|---|---|
| MN23179 | Chr03 | 58595865 | 102.06 | 8.32E-01 | 6.99E-02 | 1.67E-01 | 3.78E-01 | 1.16E-06 | 2.82E-01 | 7.90E-08 | 7.66E-01 |
| MN25537 | Chr03 | 70185379 | 149.33 | 6.59E-01 | 6.22E-01 | 1.85E-01 | 2.66E-01 | 5.92E-10 | 5.61E-01 | 4.51E-05 | 1.87E-02 |
| MN25819 | Chr03 | 70935839 | 151.89 | 6.17E-01 | 6.80E-02 | 8.87E-01 | 9.78E-01 | 4.72E-04 | 9.97E-01 | 2.74E-05 | 7.80E-01 |
| MN26009 | Chr03 | 71518284 | 154.39 | 9.28E-01 | 9.38E-06 | 3.42E-02 | 1.30E-01 | 2.89E-01 | 6.15E-02 | 3.41E-01 | 3.02E-01 |
| MN26079 | Chr03 | 71802920 | 155.80 | 6.03E-04 | 3.74E-03 | 1.94E-05 | 9.34E-06 | 9.40E-02 | 8.61E-03 | 5.86E-01 | 6.91E-01 |
| MN27530 | Chr04 | 4223843 | 27.71 | 1.28E-05 | 2.36E-03 | 3.29E-02 | 1.95E-01 | 2.09E-01 | 6.16E-02 | 7.24E-01 | 1.00E+00 |
| MN27534 | Chr04 | 4229836 | 27.75 | 4.79E-01 | 1.48E-02 | 2.18E-06 | 8.45E-06 | 6.30E-01 | 3.30E-02 | 7.49E-01 | 7.48E-01 |
| MN28574 | Chr04 | 13834059 | 71.37 | 4.13E-08 | 9.76E-01 | 3.86E-08 | 8.11E-01 | 7.23E-02 | 5.74E-06 | 4.88E-01 | 4.61E-04 |
| MN28575 | Chr04 | 13840285 | 71.38 | 3.70E-01 | 3.59E-05 | 1.04E-01 | 4.06E-02 | 3.09E-02 | 4.78E-02 | 6.86E-01 | 5.95E-02 |
| MN28847 | Chr04 | 23092693 | 71.62 | 1.13E-01 | 2.20E-01 | 1.10E-02 | 4.48E-02 | 2.17E-01 | 8.26E-07 | 7.41E-01 | 1.60E-02 |
| MN29446 | Chr04 | 51365107 | 82.14 | NA | 3.05E-01 | NA | 5.80E-01 | 1.81E-01 | 1.39E-09 | 7.87E-01 | 3.84E-03 |
| MN29451 | Chr04 | 51427526 | 82.36 | 2.08E-05 | 3.05E-01 | 1.69E-05 | 5.80E-01 | 1.81E-01 | NA | 7.87E-01 | 3.84E-03 |
| MN29587 | Chr04 | 52404340 | 84.01 | 6.95E-03 | 7.62E-02 | 5.98E-03 | 2.58E-05 | 1.77E-01 | 4.06E-02 | 4.05E-01 | 3.30E-02 |
| MN29839 | Chr04 | 54019798 | 94.50 | 3.04E-01 | 1.82E-06 | 1.80E-01 | 5.48E-01 | 1.97E-01 | 4.44E-01 | 6.79E-01 | 1.52E-02 |
| MN29933 | Chr04 | 54337785 | 94.71 | 5.49E-01 | 3.36E-08 | 6.80E-01 | 7.38E-01 | 9.80E-01 | 4.66E-01 | 2.78E-01 | 1.57E-01 |
| MN30743 | Chr04 | 57705122 | 106.83 | 3.39E-02 | 4.95E-02 | 6.35E-01 | 1.69E-05 | 8.04E-01 | 4.14E-01 | 9.37E-01 | 1.42E-02 |
| MN30889 | Chr04 | 58285783 | 106.95 | 2.37E-03 | 1.79E-02 | 1.25E-10 | 1.51E-01 | 4.63E-01 | 8.20E-01 | 3.57E-01 | 7.99E-02 |
| MN30901 | Chr04 | 58338621 | 106.96 | 2.70E-03 | 2.95E-02 | 4.83E-01 | 2.49E-01 | 6.93E-01 | 8.14E-06 | 4.15E-01 | 1.44E-01 |
| MN30942 | Chr04 | 58486658 | 106.99 | 1.80E-09 | 3.91E-01 | 3.47E-01 | 5.69E-02 | 2.17E-01 | 1.23E-01 | 1.90E-01 | 1.54E-01 |
| MN31566 | Chr04 | 61396457 | 121.78 | 1.67E-01 | 7.03E-02 | 9.24E-04 | 2.76E-05 | 2.71E-01 | 2.63E-02 | 4.87E-01 | 4.52E-01 |
| MN31652 | Chr04 | 61772402 | 123.63 | 3.14E-01 | 3.98E-01 | 9.32E-01 | 7.74E-01 | 1.80E-07 | 8.30E-02 | 1.32E-07 | 1.01E-01 |
| MN33638 | Chr05 | 3301237 | 22.87 | 2.92E-01 | 1.35E-05 | 5.54E-02 | 4.12E-02 | 7.86E-01 | 6.14E-02 | 5.23E-01 | 4.32E-02 |
| MN36911 | Chr05 | 62745841 | 73.38 | 5.00E-02 | 7.76E-01 | 1.15E-05 | 3.26E-04 | 4.35E-01 | 1.67E-01 | 8.69E-01 | 1.33E-01 |
| MN38691 | Chr05 | 70963748 | 118.00 | 4.90E-02 | 6.59E-02 | 4.65E-01 | 1.09E-01 | 7.03E-05 | 5.48E-01 | 1.95E-02 | 2.67E-01 |
| MN39249 | Chr06 | 2325352 | 28.20 | 9.65E-01 | 2.78E-01 | 2.92E-01 | 3.43E-02 | 2.00E-03 | 8.34E-01 | 2.15E-06 | 3.69E-03 |
| MN39278 | Chr06 | 2491448 | 30.03 | 1.07E-01 | 2.98E-05 | 8.38E-03 | 5.50E-03 | 1.31E-04 | 8.13E-02 | 1.64E-01 | 1.15E-02 |
| MN39365 | Chr06 | 3407622 | 32.68 | 4.90E-04 | 1.02E-01 | 1.05E-06 | 4.41E-03 | 1.31E-02 | 9.83E-07 | 1.19E-01 | 4.76E-02 |
| MN40133 | Chr06 | 39213141 | 50.50 | 7.54E-01 | 6.88E-01 | 4.11E-01 | 8.62E-02 | 3.98E-11 | 9.99E-01 | 5.57E-01 | 2.99E-01 |
| MN40138 | Chr06 | 39236984 | 50.60 | 3.33E-01 | 8.24E-05 | 2.53E-01 | 4.90E-02 | 9.92E-02 | 4.59E-01 | 1.24E-02 | 1.81E-02 |
| MN40363 | Chr06 | 43198230 | 52.55 | 8.28E-01 | 6.38E-01 | 9.82E-01 | 9.54E-01 | 1.48E-01 | 9.25E-01 | 1.79E-05 | 2.70E-02 |
| MN41467 | Chr06 | 49660992 | 89.27 | 1.33E-01 | 5.67E-03 | 3.86E-02 | 1.89E-01 | 7.22E-08 | 4.47E-01 | 6.84E-01 | 7.67E-01 |
| MN41468 | Chr06 | 49663694 | 89.27 | 3.40E-01 | 1.15E-04 | 7.01E-02 | 5.73E-02 | 1.62E-01 | 5.72E-01 | 4.75E-02 | 8.69E-01 |
| MN41469 | Chr06 | 49672518 | 89.28 | 3.40E-01 | 1.15E-04 | 7.01E-02 | 5.73E-02 | 1.62E-01 | 5.72E-01 | 4.75E-02 | 8.69E-01 |
| MN42533 | Chr06 | 54391353 | 125.43 | 1.43E-01 | 3.46E-03 | 1.19E-01 | 5.61E-03 | 5.96E-01 | 2.72E-05 | 4.07E-01 | 8.98E-01 |
| MN43454 | Chr06 | 58659106 | 157.96 | 4.71E-03 | 1.29E-01 | 3.86E-06 | 7.81E-03 | 8.76E-01 | 8.31E-01 | 7.63E-01 | 9.40E-01 |
| MN43772 | Chr06 | 59994048 | 162.94 | 3.15E-01 | 3.27E-06 | 5.04E-03 | 6.12E-01 | 1.66E-07 | 1.28E-02 | 1.45E-06 | 3.62E-02 |
| MN44944 | Chr07 | 5140651 | 58.36 | 1.10E-04 | 1.94E-01 | 1.34E-01 | 3.30E-02 | 5.87E-02 | 1.27E-01 | 1.10E-01 | 9.62E-01 |
| MN45713 | Chr07 | 14300338 | 72.21 | 5.33E-01 | 7.14E-06 | 8.55E-01 | 8.93E-01 | 6.69E-01 | 1.00E-01 | 9.19E-01 | 9.28E-03 |
| MN46311 | Chr07 | 54307547 | 75.24 | 5.98E-02 | 7.08E-02 | 1.79E-01 | 6.78E-05 | 9.87E-01 | 5.16E-02 | 9.68E-01 | 1.02E-01 |
| MN49182 | Chr08 | 5474884 | 58.65 | 1.67E-05 | 2.40E-01 | 2.03E-01 | 5.43E-02 | 7.76E-01 | 1.89E-01 | 8.53E-01 | 7.06E-01 |
| MN49262 | Chr08 | 5997613 | 59.32 | 6.20E-01 | 3.70E-02 | 3.59E-05 | 8.80E-02 | 8.86E-02 | 5.41E-02 | 1.02E-01 | 1.73E-01 |
| MN49453 | Chr08 | 10712721 | 67.33 | 6.71E-04 | 1.20E-01 | 3.48E-07 | 5.74E-03 | 3.56E-01 | 1.65E-01 | 5.80E-01 | 8.39E-01 |
| MN51078 | Chr08 | 59873076 | 99.35 | 5.02E-01 | 3.66E-06 | 3.50E-01 | 7.88E-01 | 6.13E-04 | 2.62E-01 | 4.05E-04 | 3.02E-01 |
| MN51169 | Chr08 | 60394198 | 103.12 | 2.92E-01 | 7.08E-01 | 7.85E-01 | 6.52E-05 | 6.75E-02 | 5.80E-02 | 6.15E-01 | 1.30E-02 |
| MN51281 | Chr08 | 60911383 | 106.38 | 4.50E-03 | 2.11E-01 | 2.60E-05 | 5.28E-02 | 1.36E-02 | 3.59E-02 | 4.51E-01 | 4.14E-02 |
| MN51462 | Chr08 | 61382939 | 109.44 | 4.38E-01 | 4.26E-02 | 8.85E-02 | 4.75E-01 | 4.88E-02 | 2.66E-01 | 9.56E-09 | 5.25E-01 |
| MN51660 | Chr08 | 61773908 | 111.98 | 8.59E-01 | 6.51E-07 | 5.10E-01 | 3.15E-01 | 8.00E-01 | 1.15E-01 | 8.58E-01 | 2.61E-01 |

| | | | | | | | | | | | |
|---|---|---|---|---|---|---|---|---|---|---|---|
| MN51698 | Chr08 | 61868146 | 112.60 | 1.76E-03 | 7.28E-01 | 3.10E-02 | 6.84E-01 | 9.75E-01 | 7.98E-06 | 7.17E-01 | 7.41E-02 |
| MN51776 | Chr08 | 61977227 | 118.53 | 8.66E-01 | 9.35E-01 | 9.36E-01 | 9.27E-01 | 3.46E-09 | 5.49E-01 | 1.82E-01 | 5.48E-01 |
| MN53585 | Chr05 | 7768366 | 67.33 | 8.17E-02 | 1.13E-05 | 1.04E-06 | 5.42E-04 | 1.49E-05 | 8.41E-04 | 4.08E-04 | 2.19E-02 |
| MN53904 | Chr09 | 15253577 | 69.39 | 7.24E-03 | 1.37E-03 | 6.14E-03 | 7.76E-05 | 5.71E-02 | 1.50E-02 | 8.85E-02 | 9.59E-02 |
| MN55398 | Chr09 | 51441691 | 94.86 | 1.89E-02 | 6.14E-01 | 1.92E-01 | 3.11E-02 | 4.64E-01 | 9.86E-06 | 4.28E-02 | 9.97E-01 |
| MN55969 | Chr09 | 53947707 | 108.19 | 1.59E-01 | 2.03E-03 | 1.65E-01 | 3.69E-01 | 6.81E-05 | 3.00E-01 | 9.75E-01 | 4.36E-02 |
| MN55971 | Chr09 | 53959893 | 108.23 | 2.07E-01 | 7.11E-01 | 7.00E-01 | 2.72E-01 | 4.12E-01 | 4.64E-01 | 1.41E-05 | 6.97E-02 |
| MN56922 | Chr09 | 58170646 | 128.11 | 9.19E-02 | 3.57E-01 | 1.40E-02 | 6.51E-03 | 5.11E-01 | 2.72E-10 | 7.84E-02 | 8.43E-01 |
| MN56939 | Chr09 | 58198140 | 128.33 | 1.64E-05 | 2.44E-01 | 2.03E-02 | 1.11E-01 | 6.55E-01 | 5.44E-03 | 1.25E-01 | 6.65E-02 |
| MN57963 | Chr10 | 3158439 | 31.94 | 1.13E-01 | 2.13E-01 | 8.39E-01 | 3.54E-01 | 1.92E-02 | 3.08E-01 | 2.64E-10 | 4.05E-01 |
| MN57975 | Chr10 | 3207954 | 32.13 | 9.69E-01 | 2.10E-01 | 1.54E-01 | 9.82E-01 | 1.07E-04 | 8.78E-01 | 1.08E-01 | 7.32E-01 |
| MN59291 | Chr10 | 9752448 | 53.93 | 1.36E-05 | 6.84E-02 | 2.21E-01 | 7.47E-01 | 9.43E-01 | 9.14E-01 | 4.47E-01 | 2.63E-03 |
| MN59473 | Chr10 | 12131889 | 56.00 | 6.62E-01 | 2.36E-01 | 9.62E-01 | 4.36E-01 | 5.31E-06 | 5.20E-01 | 1.00E-04 | 9.57E-01 |
| MN61917 | Chr10 | 56447102 | 89.00 | 4.48E-01 | 9.21E-03 | 1.04E-01 | 1.16E-01 | 1.00E-06 | 8.17E-01 | 1.13E-06 | 6.36E-01 |
| MN62419 | Chr10 | 59315321 | 103.27 | 4.93E-01 | 7.15E-06 | 1.55E-01 | 8.01E-02 | 8.34E-01 | 7.93E-02 | 7.09E-01 | 1.64E-01 |
