## Supplemental Table 11 for "Large-scale GWAS in sorghum reveals common genetic control of grain size among cereals"

Table S11 Effects of QTL in BC-NAM

| QTL ID | Chr | start | end | Traits affected in BC-NAM | | | | | $r^2$ | | | | |
|---|---|---|---|---|---|---|---|---|---|---|---|---|---|
| | | | | TKW | Volume | Length | Width | Thickness | TKW | Volume | Length | Width | Thickness |
| qGS1.2 | 1 | 10.35 | 13.66 | Y | Y | Y | Y | Y | 3.56E-02 | 1.98E-02 | 1.18E-02 | 1.38E-02 | 2.39E-02 |
| qGS1.3 | 1 | 16.89 | 18.27 | Y | Y | Y | Y | Y | 3.34E-02 | 1.63E-02 | 1.02E-02 | 1.18E-02 | 1.32E-02 |
| qGS1.5 | 1 | 31.25 | 33.54 | Y | Y | Y | Y | Y | 2.10E-02 | 1.95E-02 | 1.28E-02 | 1.49E-02 | 1.37E-02 |
| qGS1.7 | 1 | 46.68 | 46.73 | N | N | Y | N | Y | NA | NA | 6.03E-03 | NA | 4.86E-03 |
| qGS1.8 | 1 | 59.15 | 60.07 | Y | Y | Y | Y | Y | 4.16E-02 | 1.38E-02 | 5.41E-03 | 1.40E-02 | 1.52E-02 |
| qGS1.9 | 1 | 94.75 | 94.76 | Y | Y | Y | Y | Y | 1.33E-02 | 7.33E-03 | 1.31E-02 | 1.18E-02 | 6.22E-03 |
| qGS1.10 | 1 | 98.57 | 98.57 | Y | Y | Y | Y | N | 1.86E-02 | 1.22E-02 | 2.03E-02 | 1.33E-02 | NA |
| qGS1.11 | 1 | 115.03 | 116.85 | N | Y | N | N | Y | NA | 7.45E-03 | NA | NA | 3.54E-02 |
| qGS1.12 | 1 | 123.06 | 125.38 | Y | Y | Y | Y | Y | 6.44E-03 | 7.83E-03 | 6.34E-03 | 6.46E-03 | 5.07E-02 |
| qGS1.14 | 1 | 156.66 | 156.66 | Y | Y | Y | Y | N | 1.22E-02 | 6.57E-03 | 6.32E-03 | 4.88E-03 | NA |
| qGS2.1 | 2 | 0.10 | 0.10 | N | Y | Y | Y | N | NA | 1.20E-02 | 7.97E-03 | 1.30E-02 | NA |
| qGS2.2 | 2 | 13.86 | 13.93 | Y | Y | Y | Y | Y | 6.90E-03 | 1.37E-02 | 1.44E-02 | 1.12E-02 | 4.25E-03 |
| qGS2.3 | 2 | 56.44 | 56.44 | Y | Y | N | N | Y | 8.45E-03 | 4.49E-03 | NA | NA | 6.72E-03 |
| qGS2.4 | 2 | 59.55 | 59.55 | Y | Y | Y | Y | Y | 1.49E-02 | 7.02E-03 | 6.28E-03 | 5.96E-03 | 4.60E-03 |
| qGS2.5 | 2 | 88.94 | 88.94 | N | N | N | N | Y | NA | NA | NA | NA | 5.98E-04 |
| qGS2.8 | 2 | 121.89 | 123.92 | Y | Y | Y | Y | Y | 1.44E-02 | 1.36E-02 | 7.24E-03 | 7.33E-03 | 1.76E-02 |
| qGS2.9 | 2 | 134.53 | 134.53 | N | N | Y | Y | Y | NA | NA | 8.58E-03 | 5.38E-03 | 5.32E-03 |
| qGS2.10 | 2 | 148.73 | 149.23 | N | N | N | N | Y | NA | NA | NA | NA | 5.60E-03 |
| qGS2.11 | 2 | 152.32 | 152.63 | Y | Y | Y | Y | N | 9.34E-03 | 1.33E-02 | 1.89E-02 | 8.15E-03 | NA |
| qGS2.12 | 2 | 159.86 | 159.86 | Y | Y | Y | Y | N | 1.31E-02 | 2.05E-02 | 2.95E-02 | 1.47E-02 | NA |
| qGS2.14 | 2 | 180.56 | 182.44 | N | N | Y | Y | Y | NA | NA | 9.85E-03 | 1.08E-02 | 9.73E-03 |
| qGS2.15 | 2 | 188.18 | 189.09 | Y | Y | Y | Y | Y | 1.99E-02 | 2.76E-02 | 2.85E-02 | 1.57E-02 | 7.19E-03 |
| qGS3.1 | 3 | 0.68 | 0.68 | Y | Y | Y | Y | Y | 3.89E-02 | 2.99E-02 | 2.49E-02 | 1.92E-02 | 1.34E-02 |
| qGS3.2 | 3 | 4.04 | 4.05 | Y | Y | Y | Y | N | 8.72E-03 | 5.86E-03 | 4.34E-03 | 7.06E-03 | NA |
| qGS3.6 | 3 | 46.52 | 48.31 | Y | Y | Y | Y | Y | 7.90E-03 | 1.78E-02 | 1.31E-02 | 1.31E-02 | 8.74E-03 |
| qGS3.7 | 3 | 56.79 | 58.00 | Y | N | Y | Y | Y | 4.47E-03 | NA | 5.90E-03 | 8.74E-03 | 5.03E-03 |
| qGS3.8 | 3 | 70.31 | 72.25 | Y | Y | Y | Y | N | 5.21E-03 | 1.04E-02 | 1.10E-02 | 7.96E-03 | NA |
| qGS3.9 | 3 | 102.06 | 102.06 | Y | N | Y | N | Y | 5.31E-03 | NA | 4.68E-03 | NA | 1.19E-02 |
| qGS3.10 | 3 | 149.33 | 149.33 | Y | Y | N | Y | Y | 4.09E-02 | 1.82E-02 | NA | 1.30E-02 | 5.65E-02 |
| qGS3.11 | 3 | 151.89 | 151.89 | Y | N | N | N | Y | 1.09E-02 | NA | NA | NA | 2.00E-02 |
| qGS3.12 | 3 | 154.39 | 155.80 | Y | Y | Y | Y | Y | 8.65E-03 | 5.36E-03 | 5.86E-03 | 7.77E-03 | 5.71E-03 |
| qGS4.1 | 4 | 27.71 | 27.75 | Y | Y | Y | N | N | 1.46E-03 | 2.29E-03 | 2.65E-03 | NA | NA |
| qGS4.4 | 4 | 71.37 | 73.60 | Y | Y | Y | Y | Y | 1.37E-02 | 2.09E-02 | 2.28E-02 | 1.72E-02 | 7.01E-03 |
| qGS4.5 | 4 | 82.14 | 84.01 | Y | Y | Y | Y | Y | 3.26E-02 | 3.48E-02 | 3.92E-02 | 2.02E-02 | 8.42E-03 |
| qGS4.6 | 4 | 94.49 | 94.71 | Y | Y | Y | Y | Y | 4.34E-02 | 2.72E-02 | 2.33E-02 | 1.49E-02 | 1.58E-02 |
| qGS4.7 | 4 | 106.83 | 106.99 | Y | Y | Y | Y | N | 3.06E-02 | 3.01E-02 | 3.42E-02 | 2.29E-02 | NA |
| qGS4.8 | 4 | 121.78 | 123.63 | Y | Y | Y | Y | Y | 3.04E-02 | 1.51E-02 | 4.53E-03 | 1.13E-02 | 3.12E-02 |
| qGS5.1 | 5 | 22.87 | 23.25 | Y | Y | Y | Y | N | 1.35E-02 | 6.26E-03 | 6.95E-03 | 4.70E-03 | NA |
| qGS5.4 | 5 | 73.38 | 73.89 | Y | Y | Y | Y | N | 1.01E-02 | 2.37E-02 | 2.61E-02 | 1.83E-02 | NA |
| qGS5.5 | 5 | NA | NA | Y | N | N | N | N | NA | NA | NA | NA | NA |
| qGS6.1 | 6 | 28.20 | 30.03 | Y | Y | Y | Y | Y | 4.03E-02 | 1.63E-02 | 6.81E-03 | 6.55E-03 | 4.03E-02 |
| qGS6.2 | 6 | 32.68 | 33.68 | Y | Y | Y | Y | Y | 4.88E-02 | 1.16E-02 | 4.82E-03 | 2.36E-03 | 4.89E-02 |
| qGS6.3 | 6 | 50.50 | 52.75 | Y | Y | Y | Y | Y | 5.33E-02 | 2.53E-02 | 1.11E-02 | 1.11E-02 | 5.84E-02 |

| | | | | | | | | | | | | | |
|---|---|---|---|---|---|---|---|---|---|---|---|---|---|
| qGS6.4 | 6 | 89.27 | 89.28 | N | N | N | N | Y | NA | NA | NA | NA | 8.98E-03 |
| qGS6.5 | 6 | 125.43 | 125.64 | N | N | N | N | Y | NA | NA | NA | NA | 7.59E-03 |
| qGS6.6 | 6 | 157.96 | 157.96 | N | N | Y | Y | N | NA | NA | 5.40E-03 | 4.18E-03 | NA |
| qGS6.7 | 6 | 162.94 | 162.94 | Y | Y | Y | Y | Y | 3.84E-02 | 2.33E-02 | 1.17E-02 | 1.38E-02 | 3.38E-02 |
| qGS7.2 | 7 | 58.13 | 58.36 | N | Y | Y | Y | N | NA | 5.17E-03 | 4.42E-03 | 7.81E-03 | NA |
| qGS7.3 | 7 | 72.21 | 72.30 | Y | Y | N | Y | N | 1.15E-02 | 5.28E-03 | NA | 8.75E-03 | NA |
| qGS7.4 | 7 | 74.35 | 75.24 | Y | Y | N | Y | N | 6.57E-03 | 4.79E-03 | NA | 6.27E-03 | NA |
| qGS8.1 | 8 | 58.65 | 59.32 | Y | Y | Y | Y | Y | 2.21E-02 | 1.79E-02 | 1.08E-02 | 1.73E-02 | 9.75E-03 |
| qGS8.2 | 8 | 67.33 | 67.33 | N | Y | N | N | N | NA | 5.54E-05 | NA | NA | NA |
| qGS8.4 | 8 | 99.35 | 99.35 | Y | Y | N | Y | Y | 1.96E-02 | 6.47E-03 | NA | 8.11E-03 | 5.48E-03 |
| qGS8.5 | 8 | 102.58 | 103.12 | Y | Y | Y | Y | Y | 3.10E-02 | 2.99E-02 | 1.70E-02 | 2.56E-02 | 1.51E-02 |
| qGS8.6 | 8 | 106.38 | 106.38 | Y | Y | Y | Y | Y | 2.70E-02 | 2.48E-02 | 1.64E-02 | 2.11E-02 | 8.55E-03 |
| qGS8.7 | 8 | 109.44 | 109.44 | Y | N | N | N | N | NA | NA | NA | NA | NA |
| qGS8.8 | 8 | 111.98 | 112.60 | Y | Y | Y | Y | Y | 2.54E-02 | 2.44E-02 | 2.20E-02 | 1.57E-02 | 1.39E-02 |
| qGS8.9 | 8 | 118.53 | 118.53 | Y | Y | Y | N | Y | 1.26E-02 | 9.93E-03 | 6.83E-03 | NA | 1.67E-02 |
| qGS9.2 | 9 | 67.33 | 69.39 | Y | Y | Y | Y | Y | 3.92E-02 | 2.55E-02 | 1.14E-02 | 2.03E-02 | 2.98E-02 |
| qGS9.4 | 9 | 94.86 | 94.86 | N | N | Y | N | Y | NA | NA | NA | NA | NA |
| qGS9.6 | 9 | 108.19 | 108.23 | Y | Y | Y | Y | Y | 3.37E-02 | 1.98E-02 | 9.94E-03 | 1.79E-02 | 2.05E-02 |
| qGS9.8 | 9 | 128.11 | 128.33 | Y | Y | Y | Y | N | 5.34E-03 | 5.34E-03 | 6.72E-03 | 6.06E-03 | NA |
| qGS10.2 | 10 | 31.94 | 32.55 | Y | N | N | N | Y | 3.90E-03 | NA | NA | NA | NA |
| qGS10.3 | 10 | 53.93 | 56.00 | Y | Y | Y | Y | Y | 8.19E-03 | 2.27E-02 | 3.45E-02 | 1.79E-02 | 9.76E-03 |
| qGS10.5 | 10 | 89.00 | 89.00 | Y | N | N | N | Y | 7.91E-03 | NA | NA | NA | 1.66E-02 |
| qGS10.6 | 10 | 102.30 | 103.56 | Y | Y | Y | Y | N | 2.66E-02 | 1.30E-02 | 1.70E-02 | 9.00E-03 | NA |
