## Supplemental Table 12 for "Large-scale GWAS in sorghum reveals common genetic control of grain size among cereals"

Table S12 Tests of associations between population-specific QTL with grain size parameters in the alternative population

| QTL.ID | Chr | start | end | Pop | associations with grain size-rlated traits in alternative population | | | | | No. SNP | Significant threshold |
|---|---|---|---|---|---|---|---|---|---|---|---|
| | | | | | length | TKW | Thickness | Volume | Width | | |
| qGS1.1 | 1 | 3.18 | 3.18 | DP | 2.46E-05 | 2.14E-04 | NS | 2.39E-04 | NS | 43 | 1.16E-03 |
| qGS1.4 | 1 | 25.45 | 25.48 | DP | NS | NS | NS | NS | NS | 116 | 4.31E-04 |
| qGS1.6 | 1 | 43.49 | 43.60 | DP | NS | 2.00E-07 | 1.84E-04 | 1.48E-04 | NS | 127 | 3.94E-04 |
| qGS1.10 | 1 | 98.57 | 98.57 | NAM | 2.91E-04 | 5.31E-05 | 8.21E-09 | 3.08E-05 | 8.31E-05 | 93 | 5.38E-04 |
| qGS1.11 | 1 | 115.03 | 116.85 | NAM | 3.53E-04 | 1.69E-04 | NS | NS | NS | 82 | 6.10E-04 |
| qGS1.13 | 1 | 129.89 | 129.89 | DP | 9.34E-04 | NS | 2.79E-09 | NS | NS | 19 | 2.63E-03 |
| qGS1.14 | 1 | 156.66 | 156.66 | NAM | 7.11E-04 | 2.82E-04 | 2.30E-04 | NS | 2.16E-04 | 44 | 1.14E-03 |
| qGS2.1 | 2 | 0.10 | 0.10 | NAM | NS | 8.55E-05 | 8.38E-11 | 3.65E-07 | 9.74E-05 | 188 | 2.66E-04 |
| qGS2.2 | 2 | 13.86 | 13.93 | NAM | NS | 8.06E-07 | 2.28E-04 | 1.97E-05 | 7.83E-06 | 114 | 4.39E-04 |
| qGS2.3 | 2 | 56.44 | 56.44 | NAM | NS | NS | 2.74E-03 | NS | NS | 12 | 4.17E-03 |
| qGS2.4 | 2 | 59.55 | 59.55 | NAM | 1.53E-04 | 2.81E-04 | 1.69E-05 | 4.20E-05 | 4.28E-06 | 37 | 1.35E-03 |
| qGS2.5 | 2 | 88.94 | 88.94 | NAM | NS | 4.02E-04 | 1.43E-08 | 2.13E-04 | 1.49E-04 | 26 | 1.92E-03 |
| qGS2.6 | 2 | 93.17 | 93.17 | DP | NS | 1.74E-05 | NS | 1.08E-04 | 1.70E-05 | 105 | 4.76E-04 |
| qGS2.7 | 2 | 106.68 | 106.68 | DP | 3.28E-04 | 1.21E-04 | 1.04E-03 | 2.05E-04 | 1.57E-04 | 22 | 2.27E-03 |
| qGS2.8 | 2 | 121.89 | 123.92 | NAM | 5.30E-05 | 1.06E-08 | 1.25E-04 | 6.72E-05 | 1.90E-05 | 230 | 2.17E-04 |
| qGS2.9 | 2 | 134.53 | 134.53 | NAM | NS | 5.54E-04 | NS | NS | NS | 26 | 1.92E-03 |
| qGS2.12 | 2 | 159.86 | 159.86 | NAM | NS | 2.70E-05 | 3.37E-05 | 6.16E-04 | 9.10E-06 | 77 | 6.49E-04 |
| qGS2.13 | 2 | 166.28 | 166.28 | DP | 8.82E-07 | 4.54E-05 | NS | 4.08E-06 | 7.71E-04 | 17 | 2.94E-03 |
| qGS2.15 | 2 | 188.18 | 189.09 | NAM | NS | 2.13E-05 | 1.64E-05 | 1.47E-04 | 3.36E-04 | 37 | 1.35E-03 |
| qGS3.1 | 3 | 0.68 | 0.68 | NAM | NS | NS | NS | NS | NS | 37 | 1.35E-03 |
| qGS3.2 | 3 | 4.04 | 4.05 | NAM | NS | NS | NS | NS | NS | 72 | 6.94E-04 |
| qGS3.3 | 3 | 9.19 | 9.19 | DP | NS | 2.97E-04 | 5.03E-04 | NS | NS | 90 | 5.56E-04 |
| qGS3.4 | 3 | 11.06 | 11.06 | DP | 8.93E-04 | 3.43E-07 | 7.88E-04 | 8.58E-04 | 7.03E-06 | 26 | 1.92E-03 |
| qGS3.5 | 3 | 31.12 | 31.12 | DP | 1.89E-04 | 5.02E-04 | 7.42E-06 | 1.14E-04 | NS | 81 | 6.17E-04 |
| qGS3.7 | 3 | 56.79 | 58.00 | NAM | NS | 7.47E-10 | 1.05E-12 | 2.12E-09 | 2.97E-05 | 870 | 5.75E-05 |
| qGS3.9 | 3 | 102.06 | 102.06 | NAM | NS | 1.32E-06 | 3.49E-05 | NS | 6.04E-04 | 61 | 8.20E-04 |
| qGS3.10 | 3 | 149.33 | 149.33 | NAM | NS | 1.28E-05 | 5.08E-05 | 1.53E-05 | 1.17E-04 | 118 | 4.24E-04 |
| qGS3.11 | 3 | 151.89 | 151.89 | NAM | 2.04E-05 | 4.52E-05 | 3.72E-04 | 1.62E-08 | 5.94E-05 | 71 | 7.04E-04 |
| qGS3.12 | 3 | 154.39 | 155.80 | NAM | 7.41E-05 | 3.30E-05 | 7.74E-10 | 2.23E-07 | 4.73E-06 | 134 | 3.73E-04 |
| qGS4.1 | 4 | 27.71 | 27.75 | NAM | NS | 6.72E-07 | 6.01E-08 | 7.88E-06 | 3.32E-04 | 51 | 9.80E-04 |
| qGS4.2 | 4 | 29.99 | 29.99 | DP | NS | NS | NS | NS | NS | 14 | 3.57E-03 |
| qGS4.3 | 4 | 55.81 | 55.81 | DP | NS | NS | NS | NS | NS | 9 | 5.56E-03 |
| qGS4.7 | 4 | 106.83 | 106.99 | NAM | NS | 1.78E-08 | 2.75E-09 | 1.43E-06 | 1.77E-07 | 380 | 1.32E-04 |
| qGS4.9 | 4 | 152.27 | 152.27 | DP | 1.42E-03 | NS | NS | NS | NS | 17 | 2.94E-03 |
| qGS5.2 | 5 | 64.58 | 65.76 | DP | 2.77E-07 | 2.24E-06 | NS | 1.07E-06 | 5.67E-06 | 434 | 1.15E-04 |
| qGS5.3 | 5 | 70.70 | 70.70 | DP | 6.74E-06 | NS | 4.36E-04 | 2.86E-05 | NS | 62 | 8.06E-04 |
| qGS6.1 | 6 | 28.20 | 30.03 | NAM | NS | NS | NS | NS | NS | 52 | 9.62E-04 |
| qGS6.3 | 6 | 50.50 | 52.75 | NAM | NS | NS | NS | NS | 9.46E-05 | 221 | 2.26E-04 |
| qGS6.4 | 6 | 89.27 | 89.28 | NAM | 2.65E-04 | NS | NS | NS | NS | 168 | 2.98E-04 |
| qGS6.6 | 6 | 157.96 | 157.96 | NAM | NS | NS | 5.86E-04 | NS | 3.82E-05 | 64 | 7.81E-04 |
| qGS6.7 | 6 | 162.94 | 162.94 | NAM | NS | 3.04E-04 | 4.85E-04 | NS | 3.09E-04 | 71 | 7.04E-04 |
| qGS7.1 | 7 | 16.76 | 16.76 | DP | NS | NS | NS | NS | NS | 8 | 6.25E-03 |
| qGS7.5 | 7 | 78.67 | 78.67 | DP | 9.12E-04 | 6.26E-04 | NS | 3.22E-04 | NS | 12 | 4.17E-03 |

| | | | | | | | | | | | |
|---|---|---|---|---|---|---|---|---|---|---|---|
| qGS7.6 | 7 | 106.63 | 106.63 | DP | NS | NS | NS | NS | 1.43E-03 | 12 | 4.17E-03 |
| qGS8.1 | 8 | 58.65 | 59.32 | NAM | 6.85E-05 | 8.58E-07 | 1.15E-06 | 2.71E-05 | 1.97E-05 | 173 | 2.89E-04 |
| qGS8.2 | 8 | 67.33 | 67.33 | NAM | NS | 2.48E-07 | 4.74E-05 | 3.35E-06 | 9.11E-07 | 247 | 2.02E-04 |
| qGS8.3 | 8 | 87.75 | 87.75 | DP | 1.03E-04 | NS | NS | 3.94E-06 | 1.46E-04 | 67 | 7.46E-04 |
| qGS8.4 | 8 | 99.35 | 99.35 | NAM | NS | 1.39E-03 | 3.24E-07 | 6.85E-04 | NS | 22 | 2.27E-03 |
| qGS8.6 | 8 | 106.38 | 106.38 | NAM | NS | 6.10E-04 | NS | NS | NS | 35 | 1.43E-03 |
| qGS8.7 | 8 | 109.44 | 109.44 | NAM | NS | NS | 3.36E-04 | NS | NS | 48 | 1.04E-03 |
| qGS8.8 | 8 | 111.98 | 112.60 | NAM | NS | NS | NS | NS | NS | 60 | 8.33E-04 |
| qGS8.9 | 8 | 118.53 | 118.53 | NAM | NS | NS | 6.36E-04 | NS | NS | 27 | 1.85E-03 |
| qGS8.1[illegible] | 8 | 127.16 | 127.16 | DP | NS | 1.43E-04 | 1.45E-03 | 4.06E-04 | NS | 12 | 4.17E-03 |
| qGS9.1 | 9 | 14.45 | 14.45 | DP | NS | 3.65E-03 | NS | NS | NS | 10 | 5.00E-03 |
| qGS9.3 | 9 | 75.64 | 76.17 | DP | 6.01E-05 | 9.90E-07 | 3.27E-06 | 5.93E-06 | 1.84E-06 | 153 | 3.27E-04 |
| qGS9.4 | 9 | 94.86 | 94.86 | NAM | NS | 9.09E-06 | 3.01E-07 | 1.05E-07 | 7.46E-05 | 34 | 1.47E-03 |
| qGS9.5 | 9 | 104.82 | 104.93 | DP | NS | 2.82E-05 | 1.46E-04 | NS | NS | 49 | 1.02E-03 |
| qGS9.6 | 9 | 108.19 | 108.23 | NAM | NS | 3.57E-04 | NS | NS | 4.15E-05 | 57 | 8.77E-04 |
| qGS9.7 | 9 | 113.99 | 113.99 | DP | 2.33E-04 | 1.26E-06 | 3.85E-05 | 2.28E-06 | 1.03E-05 | 40 | 1.25E-03 |
| qGS9.8 | 9 | 128.11 | 128.33 | NAM | NS | 8.70E-07 | 1.81E-10 | 6.61E-04 | 1.07E-03 | 45 | 1.11E-03 |
| qGS10.[illegible] | 10 | 13.22 | 13.22 | DP | 1.77E-04 | 7.32E-04 | 3.34E-03 | NS | NS | 8 | 6.25E-03 |
| qGS10.[illegible] | 10 | 53.93 | 56.00 | NAM | 6.57E-05 | 8.88E-06 | 2.44E-05 | 8.25E-06 | 6.86E-06 | 333 | 1.50E-04 |
| qGS10.[illegible] | 10 | 57.96 | 58.58 | DP | 2.66E-07 | 1.47E-07 | 3.60E-06 | 4.54E-06 | 4.72E-05 | 418 | 1.20E-04 |
| qGS10.[illegible] | 10 | 89.00 | 89.00 | NAM | NS | NS | 3.45E-07 | NS | NS | 217 | 2.30E-04 |
