## Supplemental Table 13 for "Large-scale GWAS in sorghum reveals common genetic control of grain size among cereals"

Table S13 Comparison of grain size regions identified in rice and maize with grain size QTL identified in this study

| SNPs identified in rice and maize | | | | Predicted position in sorghum | | Closest QTL in this study | |
|---|---|---|---|---|---|---|---|
| Rice | Trait | Chr | Position | LG | cM | Closest QTL | Distance to closest QTL |
| Rice | Grain length | 3 | 23344870 | 1 | 35.63 | qGS1.5 | 2.10 |
| Rice | Grain length | 3 | 21838912 | 1 | 37.85 | qGS1.5 | 4.31 |
| Rice | Grain width | 10 | 19050181 | 1 | 44.18 | qGS1.6 | 0.58 |
| Rice | Grain length | 3 | 17379260 | 1 | 94.40 | qGS1.9 | 0.35 |
| Rice | Grain length | 3 | 17370294 | 1 | 94.41 | qGS1.9 | 0.34 |
| Rice | Grain length | 7 | 19506576 | 2 | 159.74 | qGS2.12 | 0.12 |
| Rice | Grain weight | 7 | 19574163 | 2 | 159.83 | qGS2.12 | 0.03 |
| Rice | Grain length | 7 | 19745179 | 2 | 160.02 | qGS2.12 | 0.17 |
| Rice | Grain weight | 2 | 5049913 | 4 | 40.11 | qGS4.2 | 10.12 |
| Rice | Grain width | 2 | 35545283 | 4 | 169.00 | qGS4.9 | 16.73 |
| Rice | Grain width | 2 | 35644379 | 4 | 169.00 | qGS4.9 | 16.73 |
| Rice | Grain width | 5 | 4806714 | 9 | 64.19 | qGS9.2 | 3.14 |
| Rice | Grain width | 5 | 4850026 | 9 | 64.57 | qGS9.2 | 2.76 |
| Rice | Grain weight | 5 | 4943999 | 9 | 65.29 | qGS9.2 | 2.05 |
| Rice | Grain width | 5 | 5270611 | 9 | 67.21 | qGS9.2 | 0.12 |
| Rice | Grain width | 5 | 5341575 | 9 | 67.69 | qGS9.2 | 0.00 |
| Rice | Grain weight | 5 | 5343564 | 9 | 67.71 | qGS9.2 | 0.00 |
| Rice | Grain width | 5 | 5359253 | 9 | 67.80 | qGS9.2 | 0.00 |
| Rice | Grain length | 6 | 9040983 | 10 | 55.89 | qGS10.3 | 0.00 |
| Rice | Grain length | 6 | 9282785 | 10 | 56.03 | qGS10.3 | 0.03 |
| Maize | Kernel length | 1 | 289546922 | 1 | 10.51 | qGS1.2 | 0.00 |
| Maize | Kernel length | 5 | 15323770 | 1 | 29.26 | qGS1.4 | 3.78 |
| Maize | Kernel length | 5 | 33978605 | 1 | 55.18 | qGS1.8 | 3.96 |
| Maize | Kernel length | 1 | 115012268 | 1 | 58.84 | qGS1.8 | 0.31 |
| Maize | Kernel length | 1 | 115012268 | 1 | 58.84 | qGS1.8 | 0.31 |
| Maize | Kernel length | 1 | 45780048 | 1 | 127.55 | qGS1.12 | 2.16 |
| Maize | Kernel thickness | 7 | 18426957 | 2 | 43.23 | qGS2.3 | 13.21 |
| Maize | Kernel thickness | 7 | 137701592 | 2 | 143.84 | qGS2.10 | 4.90 |
| Maize | Kernel thickness | 8 | 172351409 | 3 | 128.01 | qGS3.10 | 21.32 |
| Maize | Kernel thickness | 4 | 232139827 | 4 | 50.62 | qGS4.3 | 5.19 |
| Maize | Kernel length | 5 | 167062100 | 4 | 80.27 | qGS4.5 | 1.87 |
| Maize | Kernel length | 2 | 143268372 | 5 | 23.48 | qGS5.1 | 0.23 |
| Maize | Kernel thickness | 10 | 106051168 | 6 | 44.24 | qGS6.3 | 6.26 |
| Maize | Kernel thickness | 10 | 120118588 | 6 | 70.82 | qGS6.3 | 18.06 |
| Maize | Kernel thickness | 10 | 65089069 | 7 | 68.38 | qGS7.3 | 3.83 |
| Maize | Kernel length | 4 | 54509254 | 7 | 103.86 | qGS7.6 | 2.76 |
| Maize | Kernel width | 4 | 198929997 | 7 | 131.25 | qGS7.6 | 24.63 |
| Maize | Kernel length | 3 | 111867963 | 8 | 72.02 | qGS8.2 | 4.70 |
| Maize | Kernel thickness | 1 | 160819237 | 8 | 81.73 | qGS8.2 | 6.02 |
| Maize | Kernel length | 1 | 164640160 | 8 | 87.86 | qGS8.3 | 0.11 |
| Maize | Kernel length | 1 | 164640160 | 8 | 87.86 | qGS8.3 | 0.11 |
| Maize | Kernel length | 9 | 479709 | 10 | 53.73 | qGS10.3 | 0.20 |
| Maize | Kernel width | 5 | 50343804 | 10 | 98.15 | qGS10.6 | 4.15 |
