## Supplemental Table 14 for "Large-scale GWAS in sorghum reveals common genetic control of grain size among cereals"

Table S14 List of grain size candidate genes in sorghum

| Gene ID | chr | Start | End | Predicted cM | Original gene | species | Oppsite effect on grain number | colsest QTL |
|---|---|---|---|---|---|---|---|---|
| Sobic.001G016200 | Chr01 | 1423660 | 1428078 | 4.15 | Nuf2 family protein | Rice | NA | qGS1.1 |
| Sobic.001G018100 | Chr01 | 1526528 | 1531907 | 4.62 | qTGW3/GL3.3/TGW3 | Rice | N | qGS1.1 |
| Sobic.001G056700 | Chr01 | 4275459 | 4279430 | 10.94 | O2 | Maize | N | qGS1.2 |
| Sobic.001G101700 | Chr01 | 7782002 | 7785807 | 22.82 | OsGIF1 | Rice | NA | qGS1.4 |
| Sobic.001G107100 | Chr01 | 8265620 | 8268721 | 23.72 | SRS5 | Rice | NA | qGS1.4 |
| Sobic.001G154900 | Chr01 | 12437580 | 12447250 | 30.59 | GL3.1/qGL3 | Rice | N | qGS1.5 |
| Sobic.001G170800 | Chr01 | 14278553 | 14283299 | 36.21 | Transport protein | Rice | NA | qGS1.5 |
| Sobic.001G172400 | Chr01 | 14434718 | 14438560 | 36.48 | BRD1 | Rice | NA | qGS1.5 |
| Sobic.001G184900 | Chr01 | 15806732 | 15807894 | 40.22 | Expressed protein | Rice | NA | qGS1.6 |
| Sobic.001G254100 | Chr01 | 28215025 | 28215913 | 55.23 | PGL1 | Rice | NA | qGS1.8 |
| Sobic.001G254200 | Chr01 | 28465427 | 28472547 | 55.24 | OsFBK12 | Rice | Y | qGS1.8 |
| Sobic.001G315700 | Chr01 | 60385344 | 60390035 | 78.50 | OsMPK6 | Rice | Y | qGS1.9 |
| Sobic.001G335800 | Chr01 | 62429238 | 62434197 | 94.34 | qGW7/GL7 | Rice | N | qGS1.9 |
| Sobic.001G336200 | Chr01 | 62458876 | 62461021 | 94.36 | Grain Length3.2 | Rice | NA | qGS1.9 |
| Sobic.001G341700 | Chr01 | 62910779 | 62916258 | 94.78 | GS3 | Rice | N | qGS1.9 |
| Sobic.001G445900 | Chr01 | 72260202 | 72266394 | 137.49 | CYP90B2 | Rice | N | qGS1.13 |
| Sobic.001G448700 | Chr01 | 72598605 | 72601731 | 138.58 | TUD1 | Rice | NA | qGS1.13 |
| Sobic.001G455900 | Chr01 | 73188105 | 73199446 | 138.82 | qGL3-2/qLGY3 | Rice | N | qGS1.13 |
| Sobic.001G482600 | Chr01 | 75415876 | 75416893 | 154.53 | TIFY 11b | Rice | NA | qGS1.14 |
| Sobic.001G484200 | Chr01 | 75526130 | 75530742 | 155.31 | RGA1/D1 | Rice | NA | qGS1.14 |
| Sobic.001G485400 | Chr01 | 75624275 | 75627243 | 155.94 | BG1 | Rice | Y | qGS1.14 |
| Sobic.001G488400 | Chr01 | 75826748 | 75828379 | 157.31 | PGL1 | Rice | NA | qGS1.14 |
| Sobic.001G488500 | Chr01 | 75844162 | 75849593 | 157.45 | OsFBK12 | Rice | Y | qGS1.14 |
| Sobic.002G054800 | Chr02 | 5243140 | 5247362 | 22.67 | O2 | Maize | N | qGS2.2 |
| Sobic.002G116000 | Chr02 | 14334143 | 14341029 | 72.23 | GbssIIa | Maize | NA | qGS2.4 |
| Sobic.002G216600 | Chr02 | 60869170 | 60876291 | 122.40 | DEP1 | Rice | Y | qGS2.8 |
| Sobic.002G220700 | Chr02 | 61230574 | 61233249 | 123.12 | GS9 | Rice | N | qGS2.8 |
| Sobic.002G226500 | Chr02 | 61857257 | 61859003 | 131.16 | SG1 | Rice | NA | qGS2.9 |
| Sobic.002G257900 | Chr02 | 64368406 | 64372901 | 144.27 | GW8/OsPL16 | Rice | N | qGS2.10 |
| Sobic.002G308400 | Chr02 | 68272394 | 68274070 | 153.09 | MYB transcription factor | Rice | NA | qGS2.11 |
| Sobic.002G311000 | Chr02 | 68477396 | 68483354 | 154.13 | Receptor-like kinase | Rice | NA | qGS2.11 |
| Sobic.002G312200 | Chr02 | 68569338 | 68574778 | 155.43 | GLW7 | Rice | N | qGS2.11 |
| Sobic.002G353200 | Chr02 | 71635214 | 71637226 | 175.92 | OsBZR1 | Rice | N | qGS2.14 |
| Sobic.002G367300 | Chr02 | 72705386 | 72710986 | 179.07 | qGW7/GL7 | Rice | N | qGS2.14 |
| Sobic.002G367600 | Chr02 | 72744933 | 72746890 | 179.14 | BG2 | Rice | Y | qGS2.14 |
| Sobic.002G374400 | Chr02 | 73220257 | 73228268 | 180.36 | DEP2 | Rice | N | qGS2.14 |
| Sobic.003G014500 | Chr03 | 1285914 | 1293809 | 2.82 | MHZ7 | Rice | NA | qGS3.2 |
| Sobic.003G030600 | Chr03 | 2718024 | 2725425 | 9.09 | D2 | Rice | NA | qGS3.3 |
| Sobic.003G035400 | Chr03 | 3244200 | 3246825 | 9.29 | GW5/qSW5 | Rice | N | qGS3.3 |
| Sobic.003G140000 | Chr03 | 13615713 | 13618680 | 48.92 | OsSAMS1 | Rice | Y | qGS3.6 |
| Sobic.003G230500 | Chr03 | 57000120 | 57007815 | 87.43 | Sh2 | Maize | NA | qGS3.9 |
| Sobic.003G257400 | Chr03 | 59557217 | 59558392 | 110.27 | BG3 | Rice | Y | qGS3.9 |
| Sobic.003G277900 | Chr03 | 61388359 | 61393032 | 115.47 | D61 | Rice | Y | qGS3.9 |
| Sobic.003G380900 | Chr03 | 69444094 | 69445096 | 146.35 | SERF1 | Rice | Y | qGS3.10 |

| | | | | | | | | |
|---|---|---|---|---|---|---|---|---|
| Sobic.003G406600 | Chr03 | 71419053 | 71424347 | 153.93 | GW8/OsPL16 | Rice | N | qGS3.12 |
| Sobic.003G411900 | Chr03 | 71918718 | 71925752 | 156.40 | OsARF4 | Rice | N | qGS3.12 |
| Sobic.003G444100 | Chr03 | 74252301 | 74256013 | 173.00 | OsCCS52B | Rice | NA | qGS3.12 |
| Sobic.004G032300 | Chr04 | 2604252 | 2605787 | 17.11 | OsSGL | Rice | N | qGS4.1 |
| Sobic.004G065400 | Chr04 | 5288865 | 5292249 | 34.95 | GW6 | Rice | N | qGS4.2 |
| Sobic.004G075600 | Chr04 | #N/A | #N/A | 40.80 | Zinc finger protein | Rice | NA | qGS4.2 |
| Sobic.004G102900 | Chr04 | 9666921 | 9672711 | 62.26 | FUWA | Rice | Y | qGS4.3 |
| Sobic.004G107300 | Chr04 | 10267538 | 10272971 | 63.58 | GW2 | Rice | Y | qGS4.3 |
| Sobic.004G113100 | Chr04 | 11149537 | 11155410 | 65.60 | OsSPMS1 | Rice | N | qGS4.4 |
| Sobic.004G133600 | Chr04 | 21285737 | 21291316 | 71.53 | ZmSWEET4c | Maize | N | qGS4.4 |
| Sobic.004G163700 | Chr04 | 51292092 | 51304326 | 81.93 | SbeIIb | Maize | NA | qGS4.5 |
| Sobic.004G176000 | Chr04 | 52832596 | 52846810 | 84.50 | SDG725 | Rice | NA | qGS4.5 |
| Sobic.004G214100 | Chr04 | 56389819 | 56395087 | 101.70 | BC14 | Rice | NA | qGS4.7 |
| Sobic.004G237000 | Chr04 | 58488864 | 58490438 | 106.99 | PGL2 | Rice | NA | qGS4.7 |
| Sobic.004G247000 | Chr04 | 59472640 | 59476805 | 107.32 | Gln-4 | Maize | NA | qGS4.7 |
| Sobic.004G269900 | Chr04 | 61416646 | 61421827 | 121.91 | GS2/GL2 | Rice | N | qGS4.8 |
| Sobic.004G307800 | Chr04 | 64546000 | 64546994 | 139.89 | SGL1 | Rice | NA | qGS4.9 |
| Sobic.004G314100 | Chr04 | 65063734 | 65067245 | 142.57 | GS9 | Rice | N | qGS4.9 |
| Sobic.004G317300 | Chr04 | 65318890 | 65336694 | 144.50 | O1 | Maize | N | qGS4.9 |
| Sobic.004G323600 | Chr04 | 65819835 | 65821680 | 149.26 | SMG1 | Rice | NA | qGS4.9 |
| Sobic.004G330200 | Chr04 | 66383113 | 66384153 | 154.79 | TGW6 | Rice | NA | qGS4.9 |
| Sobic.004G338400 | Chr04 | 67036246 | 67041813 | 155.92 | TH1 | Rice | N | qGS4.9 |
| Sobic.005G001500 | Chr05 | 119063 | 122213 | 0.00 | PBF1 | Maize | NA | qGS5.5 |
| Sobic.006G059900 | Chr06 | 41007893 | 41008915 | 51.20 | ZmIPT2 | Maize | NA | qGS6.3 |
| Sobic.006G114600 | Chr06 | 48245595 | 48250231 | 84.48 | D11 | Rice | NA | qGS6.4 |
| Sobic.006G176800 | Chr06 | 53195854 | 53204792 | 115.31 | OsMKKK10 | Rice | Y | qGS6.5 |
| Sobic.006G186000 | Chr06 | 54122260 | 54124881 | 125.37 | CYP704A3 | Rice | NA | qGS6.5 |
| Sobic.006G203400 | Chr06 | 55407501 | 55411315 | 125.65 | GS2/GL2 | Rice | N | qGS6.5 |
| Sobic.006G239000 | Chr06 | 57969070 | 57983024 | 153.58 | FLO2 | Rice | N | qGS6.6 |
| Sobic.006G261200 | Chr06 | 59608893 | 59612102 | 161.52 | GL4 | Rice | NA | qGS6.7 |
| Sobic.007G059600 | Chr07 | 6233483 | 6239958 | 59.12 | OsBAK1 | Rice | NA | qGS7.2 |
| Sobic.007G101500 | Chr07 | 25707028 | 25712978 | 72.25 | Bt2 | Maize | NA | qGS7.3 |
| Sobic.007G149200 | Chr07 | 58010697 | 58016328 | 88.19 | DEP1/GGC2 | Rice | Y | qGS7.5 |
| Sobic.007G153100 | Chr07 | 58560223 | 58563102 | 90.47 | GS9 | Rice | N | qGS7.5 |
| Sobic.007G156800 | Chr07 | 59120488 | 59121768 | 97.77 | SGL1 | Rice | NA | qGS7.6 |
| Sobic.007G165800 | Chr07 | 60102559 | 60105535 | 105.51 | SLG | Rice | NA | qGS7.6 |
| Sobic.007G189000 | Chr07 | 62168848 | 62174483 | 119.92 | WTG1 | Rice | N | qGS7.6 |
| Sobic.007G193500 | Chr07 | 62605971 | 62612183 | 122.59 | GW8/OsPL16 | Rice | N | qGS7.6 |
| Sobic.008G001700 | Chr08 | 152798 | 157364 | 0.71 | PBF1 | Maize | NA | qGS8.1 |
| Sobic.008G100400 | Chr08 | 47585592 | 47589188 | 70.27 | SMK1 | Maize | NA | qGS8.2 |
| Sobic.008G173900 | Chr08 | 60836806 | 60845885 | 105.95 | OsPPKL3 | Rice | NA | qGS8.6 |
| Sobic.008G193300 | Chr08 | 62665983 | 62671469 | 134.00 | OsSUT2 | Rice | NA | qGS8.10 |
| Sobic.009G017400 | Chr09 | 1616925 | 1621888 | 20.24 | gsn1/LARGE8 | Rice | Y | qGS9.1 |
| Sobic.009G024600 | Chr09 | 2179301 | 2183620 | 28.31 | RSR1 | Rice | N | qGS9.1 |
| Sobic.009G033600 | Chr09 | 3045194 | 3048685 | 46.48 | OsSAMS1 | Rice | Y | qGS9.2 |
| Sobic.009G036400 | Chr09 | 3351538 | 3354991 | 47.68 | APG | Rice | NA | qGS9.2 |
| Sobic.009G040700 | Chr09 | 3913451 | 3922888 | 49.91 | OsPPKL2 | Rice | NA | qGS9.2 |

| | | | | | | | | |
|---|---|---|---|---|---|---|---|---|
| Sobic.009G049400 | Chr09 | 4902297 | 4908926 | 54.66 | SRS3 | Rice | Y | qGS9.2 |
| Sobic.009G053600 | Chr09 | 5403633 | 5408821 | 58.10 | GS5 | Rice | N | qGS9.2 |
| Sobic.009G070000 | Chr09 | 8184500 | 8186318 | 67.81 | GW5/qSW5 | Rice | N | qGS9.2 |
| Sobic.009G093100 | Chr09 | 22875390 | 22878858 | 71.10 | OsAGSW1 | Rice | Y | qGS9.2 |
| Sobic.009G124200 | Chr09 | 47743018 | 47750375 | 84.09 | SMOS1 | Rice | NA | qGS9.3 |
| Sobic.009G141500 | Chr09 | 49879082 | 49879738 | 87.27 | SERF1 | Rice | Y | qGS9.4 |
| Sobic.009G227300 | Chr09 | 56873995 | 56875316 | 117.49 | BG3 | Rice | Y | qGS9.7 |
| Sobic.010G017400 | Chr10 | 1413401 | 1415917 | 23.59 | DLT | Rice | NA | qGS10.2 |
| Sobic.010G043200 | Chr10 | 3339728 | 3345825 | 32.46 | OsMAPK6 | Rice | NA | qGS10.2 |
| Sobic.010G047400 | Chr10 | 3668202 | 3673695 | 32.75 | HGW | Rice | N | qGS10.2 |
| Sobic.010G069600 | Chr10 | 5582591 | 5586127 | 34.53 | SMG1 | Rice | NA | qGS10.2 |
| Sobic.010G072300 | Chr10 | 5859074 | 5867276 | 35.19 | Sh1 | Maize | NA | qGS10.2 |
| Sobic.010G091700 | Chr10 | 8099384 | 8100834 | 44.85 | PGL2/OsBUL1 | Rice | NA | qGS10.3 |
| Sobic.010G110100 | Chr10 | 11004657 | 11007712 | 55.67 | A transcription factor | Rice | NA | qGS10.3 |
| Sobic.010G111200 | Chr10 | 11197868 | 11198820 | 55.78 | GASR7 | Rice | NA | qGS10.3 |
| Sobic.010G153900 | Chr10 | 44759504 | 44767453 | 58.73 | OsSPMS1 | Rice | N | qGS10.4 |
| Sobic.010G210100 | Chr10 | 55370341 | 55372973 | 80.38 | GW6 | Rice | N | qGS10.5 |
| Sobic.010G228100 | Chr10 | 57117625 | 57119973 | 91.21 | DEP3 | Rice | Y | qGS10.5 |
| Sobic.010G273900 | Chr10 | 60695280 | 60700347 | 114.95 | SbeI | Maize | NA | qGS10.6 |
| Sobic.010G277300 | Chr10 | 61014032 | 61018074 | 118.00 | BRD2 | Rice | N | qGS10.6 |

| Distance to closest QTL | Referencese |
|---|---|
| 0.97 | Huang et al., 2012b |
| 1.44 | Hu et al., 2018/Xia et al., 2018/Ying et al., 2018; |
| 0.00 | Hartings et al., 1989 |
| 2.63 | He et al. (2017) |
| 1.73 | Segami, 2012 |
| 0.66 | Qi et al., 2012; Zhang et al., 2012 |
| 2.67 | Huang et al., 2012b |
| 2.94 | Mori et al., 2002 |
| 3.27 | Huang et al., 2012b |
| 3.92 | Heang and Sassa, 2012a |
| 3.91 | Chen et al., 2013 |
| 16.25 | Gou et al., 2018 |
| 0.41 | Wang et al., 2015a; Wang et al., 2015b |
| 0.39 | Adamski et al., 2009; Xu et al., 2015 |
| 0.02 | Mao et al., 2010; Li et al., 2010c |
| 7.60 | Wu et al., 2008 |
| 8.69 | Hu et al., 2013 |
| 8.93 | Yu et al., 2018/Liu et al., 2018 |
| 2.13 | Hakata et al., 2012 |
| 1.36 | Ashikari, 1999 |
| 0.72 | Liu et al., 2015b |
| 0.65 | Heang and Sassa, 2012a |
| 0.79 | Chen et al., 2013 |
| 8.73 | Hartings et al., 1989 |
| 12.68 | Jiang et al., 2013a |
| 0.00 | Huang et al., 2009; Li et al., 2012b |
| 0.00 | Zhao et al.(2018) |
| 3.36 | Nakagawa et al., 2012 |
| 4.46 | Wang et al., 2012b |
| 0.46 | Huang et al., 2012b |
| 1.50 | Huang et al., 2012b |
| 2.79 | Si et al., 2016 |
| 4.64 | Zhu et al. (2015) |
| 1.50 | Wang et al., 2015a; Wang et al., 2015b |
| 1.42 | Xu et al., 2015 |
| 0.21 | Li et al., 2010a |
| 1.22 | Ma et al., 2013 |
| 0.10 | Hong et al., 2003 |
| 0.11 | Liu et al., 2017 |
| 0.61 | Chen et al., 2013 |
| 14.63 | Jiang et al., 2013a |
| 8.21 | Xiao et al., 20018 |
| 13.41 | Morinaka et al., 2006; Jiang et al., 2013b |
| 2.98 | Schmidt et al., 2013 |

| | |
|---|---|
| 0.47 | Wang et al., 2012 |
| 0.60 | Hu et al., 2018 |
| 18.61 | Su’udi et al., 2012 |
| 10.60 | Wang et al. (2016) |
| 4.97 | Song et al., 2015 |
| 10.82 | Huang et al., 2012b |
| 6.45 | Chen 2015 |
| 7.77 | Song et al., 2007;Li et al., 2010b |
| 5.77 | Tao et al., 2018 |
| 0.00 | Sosso et al., 2015;Zhang et al., 2015 |
| 0.21 | Jiang et al., 2013a |
| 0.49 | Sui et al., 2012 |
| 5.13 | Zhang et al., 2011 |
| 0.00 | Heang and Sassa, 2012b;Jang and Li, 2017 |
| 0.33 | Martin et al., 2006 |
| 0.00 | Che et al., 2015; Hu et al., 2015 |
| 12.38 | Nakagawa et al., 2012 |
| 9.70 | Zhao et al.(2018) |
| 7.76 | Wang et al., 2012a |
| 3.00 | Duan et al., 2014 |
| 2.53 | Ishimaru et al., 2013 |
| 3.65 | Li et al., 2012c |
| 22.87 | Lang et al., 2014 |
| 0.00 | Weng et al., 2013 |
| 4.79 | Tanabe et al., 2005;Wu et al., 2008 |
| 10.11 | Gou et al., 2018 |
| 0.06 | Tang et al. (2016) |
| 0.00 | Che et al., 2015; Hu et al., 2015 |
| 4.38 | She et al., 2010 |
| 1.42 | Wu et al. (2017) |
| 0.77 | Yuan et al. (2017) |
| 0.00 | Jiang et al., 2013a |
| 9.51 | Huang et al., 2009; Li et al., 2012b; Sun et al., 2018 |
| 11.80 | Zhao et al.(2018) |
| 8.85 | Nakagawa et al., 2012 |
| 1.11 | Feng et al. (2016) |
| 13.29 | Wang et al. (2017) |
| 15.97 | Wang et al., 2012b |
| 57.94 | Lang et al., 2014 |
| 2.94 | Li et al., 2014 |
| 0.42 | Zhan et al., 2012 |
| 6.85 | Eom et al., 2011 |
| 5.78 | Gou et al., 2018; Xu et al., 2018 |
| 13.85 | Fu and Xue,, 2010 |
| 20.85 | Chen et al., 2013 |
| 19.65 | Heang and Sassa, 2012b |
| 17.42 | Zhan et al., 2012 |

| | |
|---|---|
| 12.68 | Kitagawa et al., 2010 |
| 9.24 | Li et al., 2011 |
| 0.00 | Liu et al., 2017 |
| 1.71 | Li and Li 2015 |
| 7.93 | Hirano et al. (2017) |
| 7.60 | Schmidt et al., 2013 |
| 3.50 | Xiao et al., 20018 |
| 8.34 | Tong et al. (2012) |
| 0.00 | Liu et al. (2015b) |
| 0.20 | Li et al., 2012a |
| 1.98 | Duan et al., 2014 |
| 2.64 | Jiang et al., 2013a |
| 9.08 | Heang and Sassa, 2012b;Jang and Li, 2017 |
| 0.00 | Huang et al., 2012b |
| 0.00 | Huang et al., 2012b |
| 0.15 | Tao et al., 2018 |
| 8.62 | Song et al., 2015 |
| 2.21 | Qiao et al., 2011 |
| 11.39 | Jiang et al., 2013a |
| 15.70 | Hong et al., 2005 |
