## Supplemental Table 15 for "Large-scale GWAS in sorghum reveals common genetic control of grain size among cereals"

Table S15 Summary of candidate genes within close vicinities of grain size QTL

| | No. candidates with opposite effect on grain number | No. candidates with no opposite effect on grain number | Candidates no information available | Total number |
|---|---|---|---|---|
| Total number | 21 | 37 | 53 | 111 |
| Within 0.1 cM vicinity of grain size and weight QTL | 1 | 7 | 8 | 16 |
| Within 1 cM vicinity of grain size and weight QTL | 4 | 15 | 17 | 36 |
