## Supplemental Table 16 for "Large-scale GWAS in sorghum reveals common genetic control of grain size among cereals"

Table S16 Correspondence between numbers on x-axis of Figure 5 and identities of QTL identified in BC-NAM

| Numbers on x-axis of figure 4 | QTL ID |
|---|---|
| 1 | qGS9.8 |
| 2 | qGS6.7 |
| 3 | qGS1.10 |
| 4 | qGS9.4 |
| 5 | MN09216 |
| 6 | qGS1.3 |
| 7 | qGS6.1 |
| 8 | qGS2.1 |
| 9 | qGS1.8 |
| 10 | qGS3.11 |
| 11 | qGS5.1 |
| 12 | qGS3.10 |
| 13 | qGS8.4 |
| 14 | qGS2.5 |
| 15 | qGS10.5 |
| 16 | qGS3.2 |
| 17 | qGS3.12 |
| 18 | qGS9.2 |
| 19 | qGS4.1 |
| 20 | qGS1.2 |
| 21 | qGS1.5 |
| 22 | qGS3.6 |
| 23 | qGS6.6 |
| 24 | qGS2.14 |
| 25 | qGS9.6 |
| 26 | qGS10.2 |
| 27 | qGS3.1 |
| 28 | qGS7.3 |
| 29 | qGS8.1 |
| 30 | qGS2.2 |
| 31 | qGS7.4 |
| 32 | qGS8.2 |
| 33 | qGS3.7 |
| 34 | qGS2.9 |
| 35 | qGS7.2 |
| 36 | qGS2.8 |
| 37 | qGS6.3 |
| 38 | qGS3.9 |
| 39 | qGS8.7 |
| 40 | qGS10.3 |
| 41 | qGS10.6 |
| 42 | qGS5.4 |
| 43 | qGS6.4 |
| 44 | qGS6.2 |

| | |
|---|---|
| 45 | qGS1.7 |
| 46 | qGS3.8 |
| 47 | qGS2.4 |
| 48 | qGS1.12 |
| 49 | qGS1.14 |
| 50 | qGS8.5 |
| 51 | qGS6.5 |
| 52 | qGS2.10 |
| 53 | qGS1.9 |
| 54 | qGS8.9 |
| 55 | qGS2.12 |
| 56 | qGS1.11 |
| 57 | qGS4.5 |
| 58 | qGS2.11 |
| 59 | qGS4.7 |
| 60 | qGS4.8 |
| 61 | qGS8.8 |
| 62 | qGS8.6 |
| 63 | qGS2.3 |
| 64 | qGS4.4 |
| 65 | qGS4.6 |
| 66 | qGS2.15 |
