## Supplemental Figures for "Large-scale GWAS in sorghum reveals common genetic control of grain size among cereals"

### Figure S1  PCA and LD of NAM

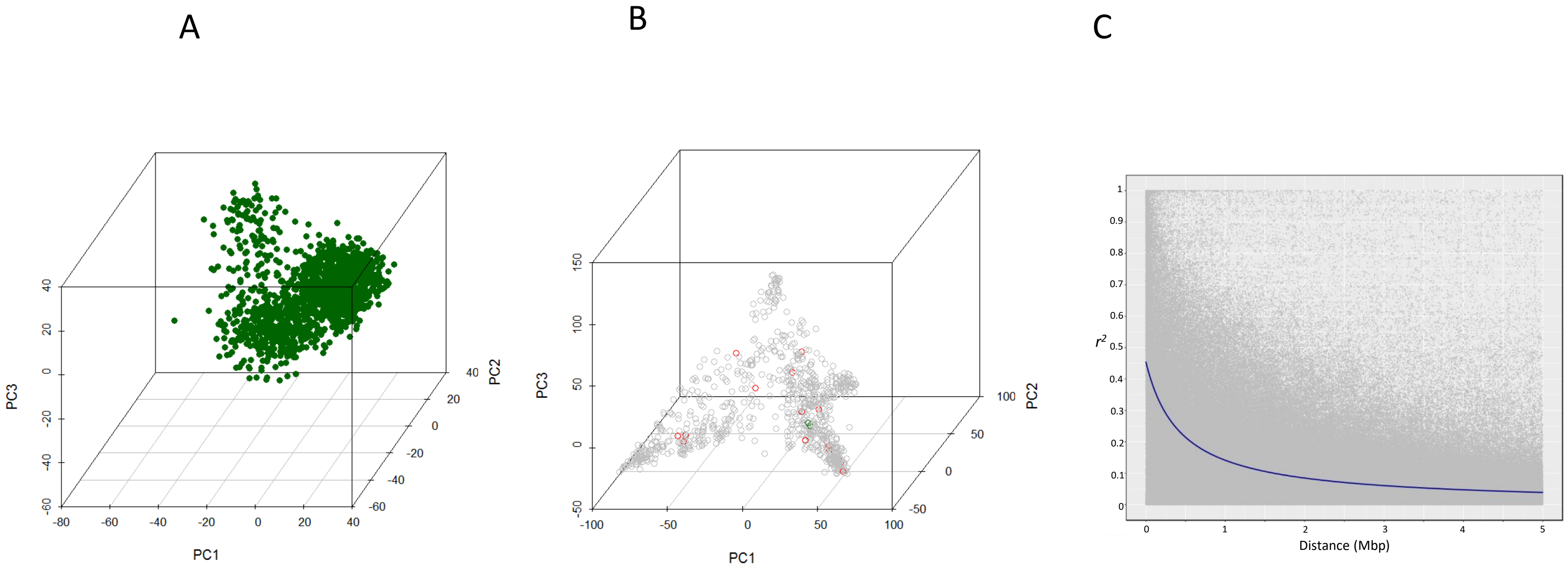

### Figure S2 PCA analysis of grain size variation in DP

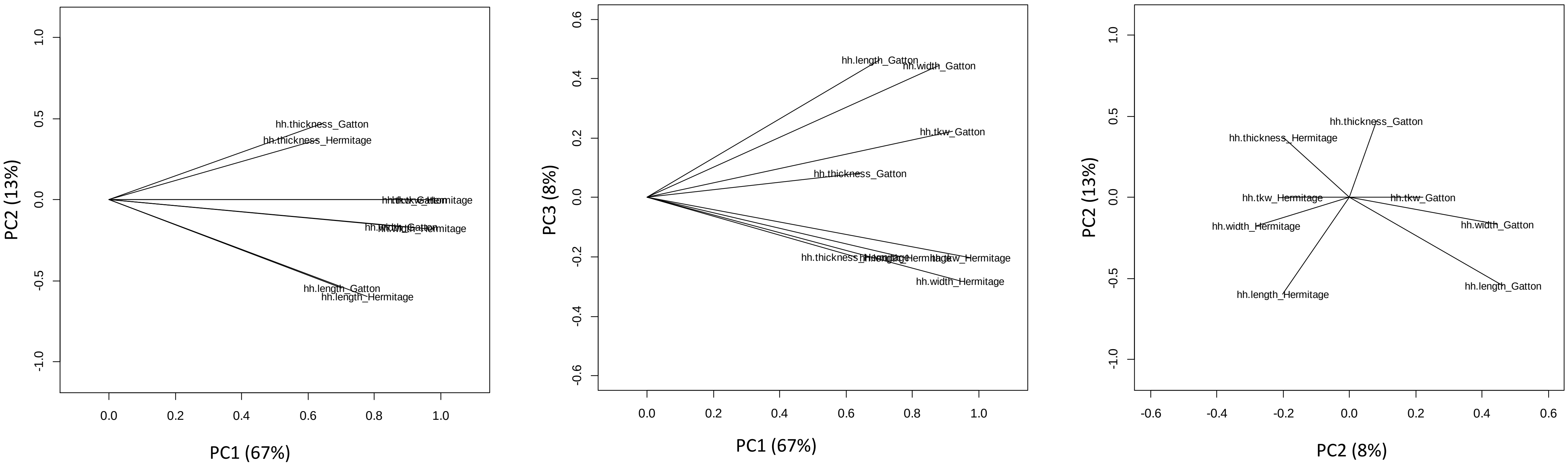

### Figure S3 PCA analysis of grain size variation in BC-NAM

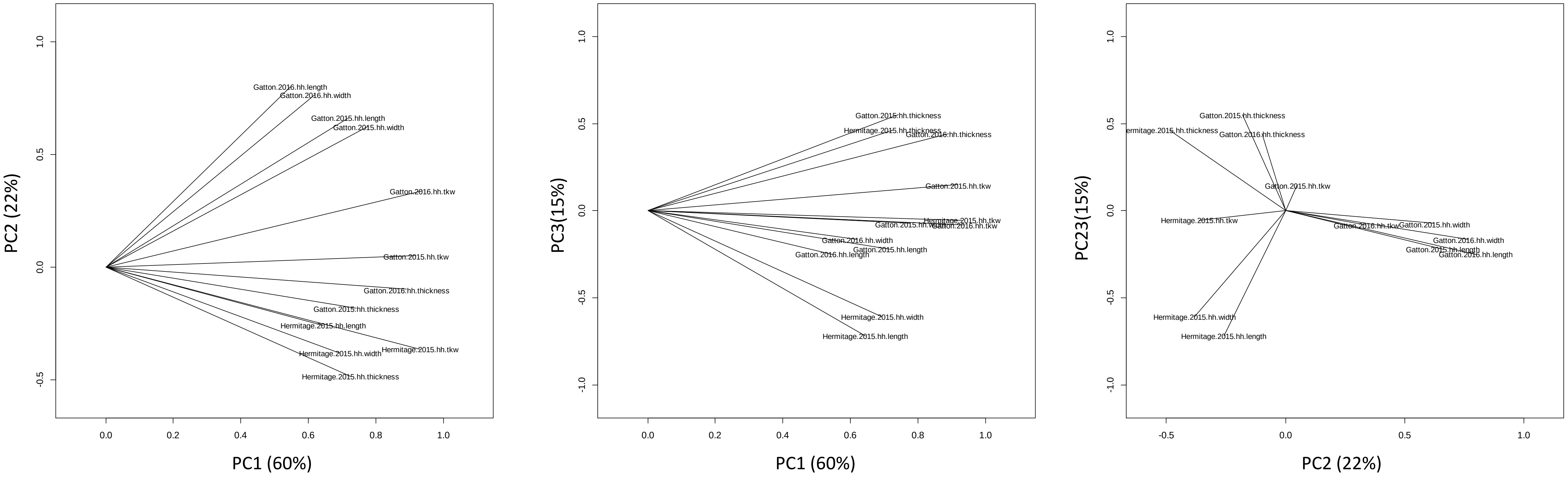

### Figure S4  population structure, LD and PCA of DP

A

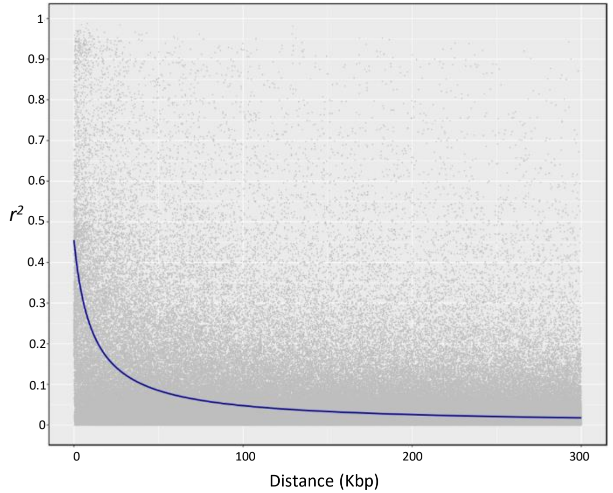


B

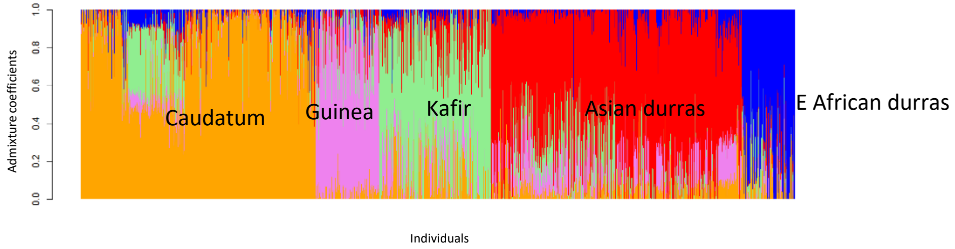


C

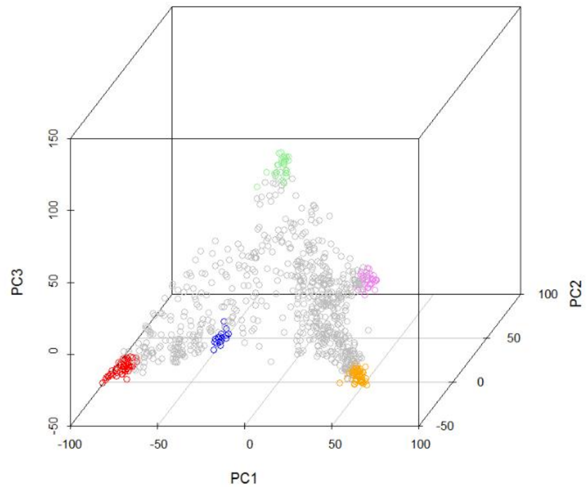


| | |
|---|---|
| Caudatum | Orange |
| Guinea | Violet |
| Kafir | Lightgreen |
| Asian durras | red |
| E African durras | blue |

### Figure S5 Distribution of SNPs across sorghum genome in DP

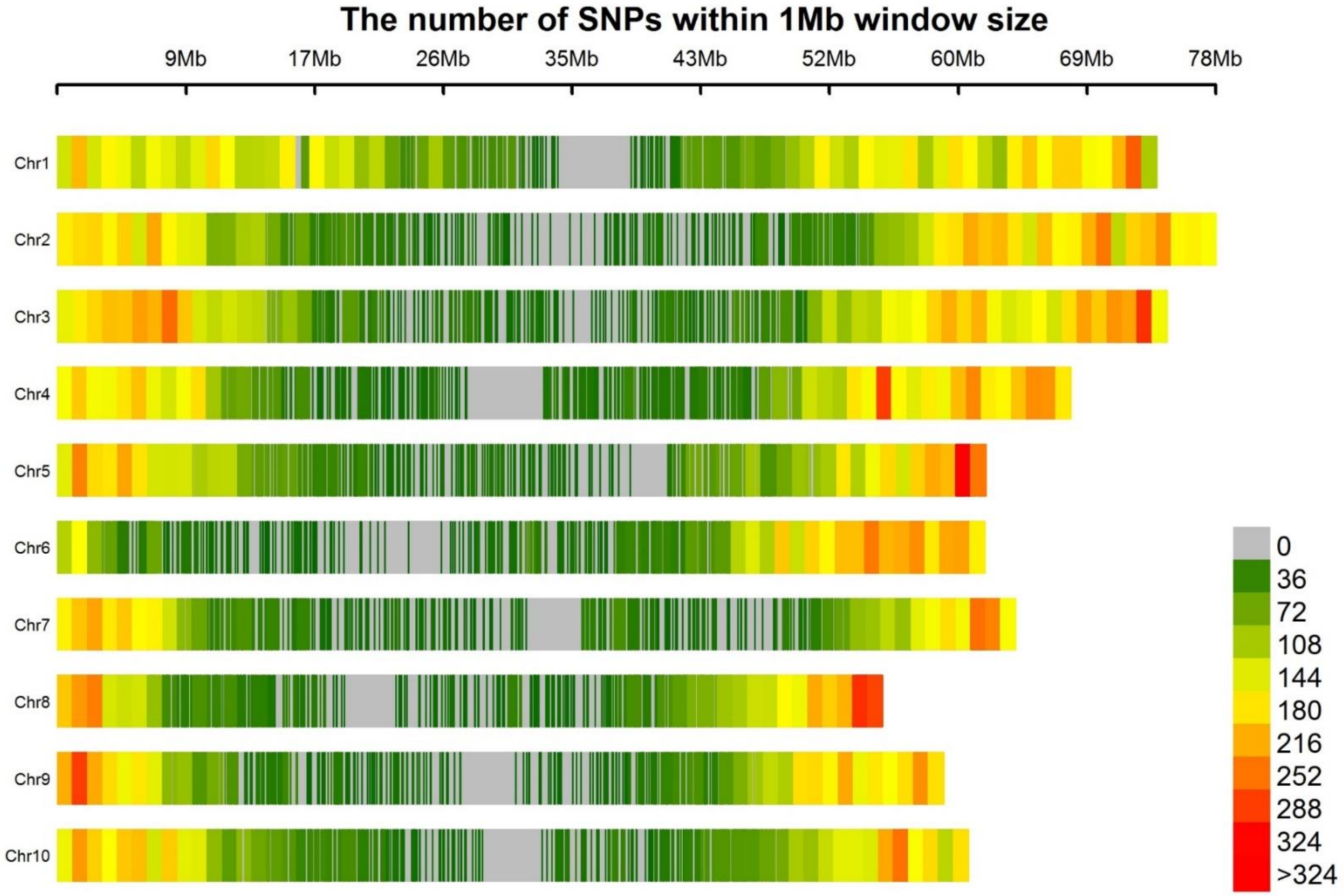

Figure S6 Distribution of SNPs across sorghum genome in NAM

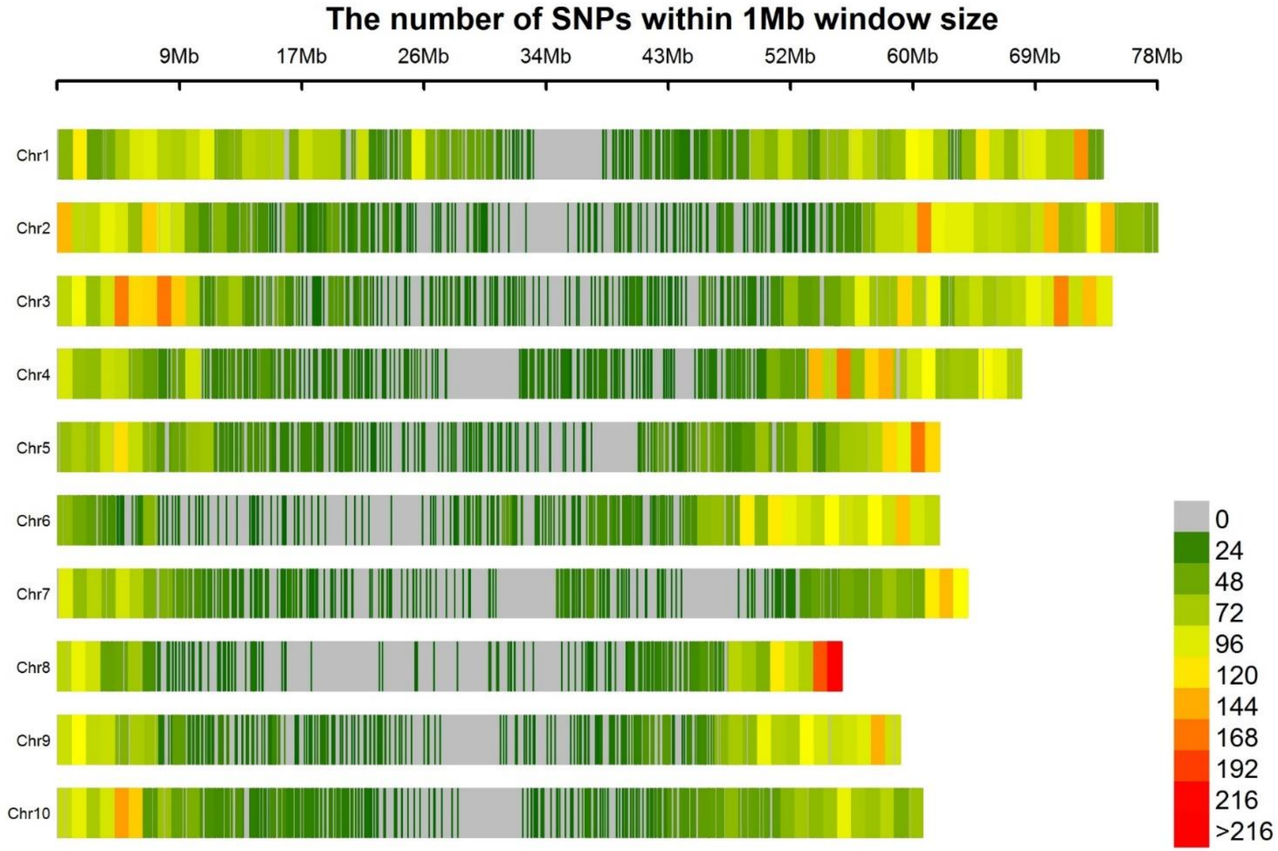

Figure S7

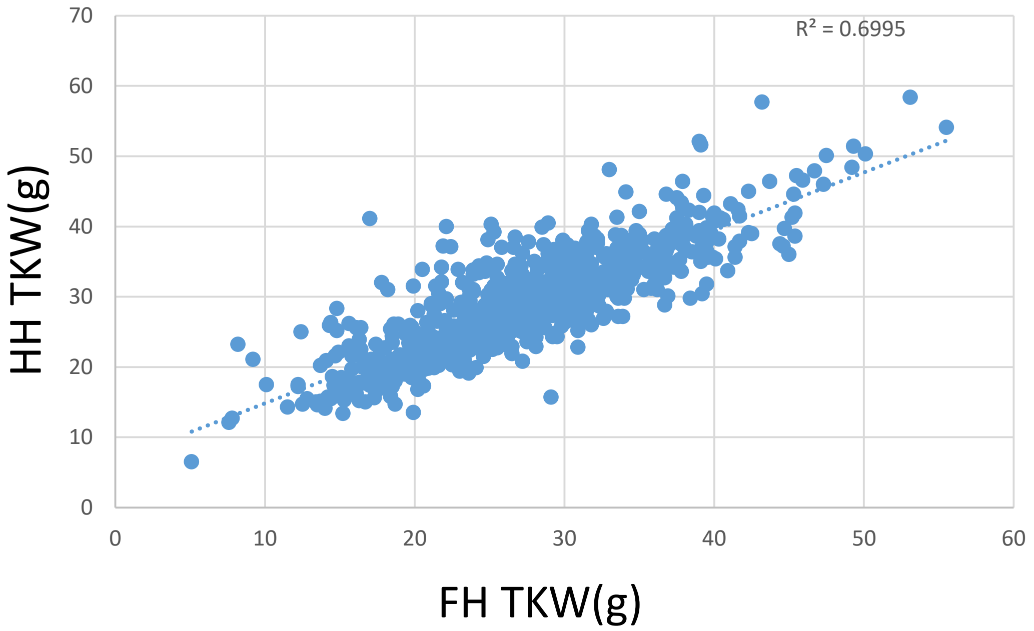

HH TKW(g)
FH TKW(g)
R² = 0.6995
70
60
50
40
30
20
10
0
0
10
20
30
40
50
60


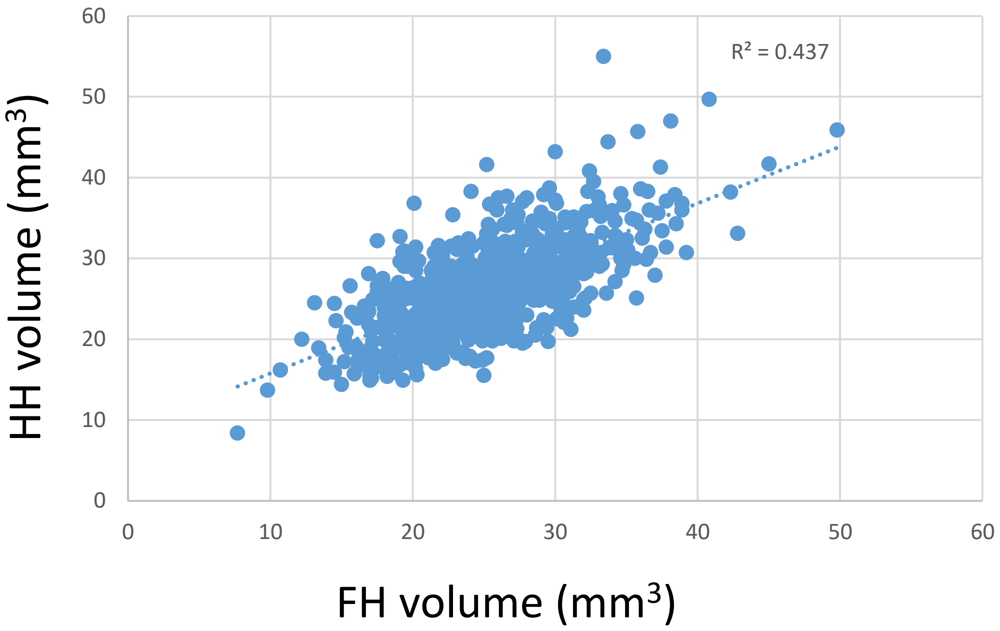

HH volume (mm³)
FH volume (mm³)
R² = 0.437
60
50
40
30
20
10
0
0
10
20
30
40
50
60

Figure S8 DP PCs

PC1

PC2

PC3

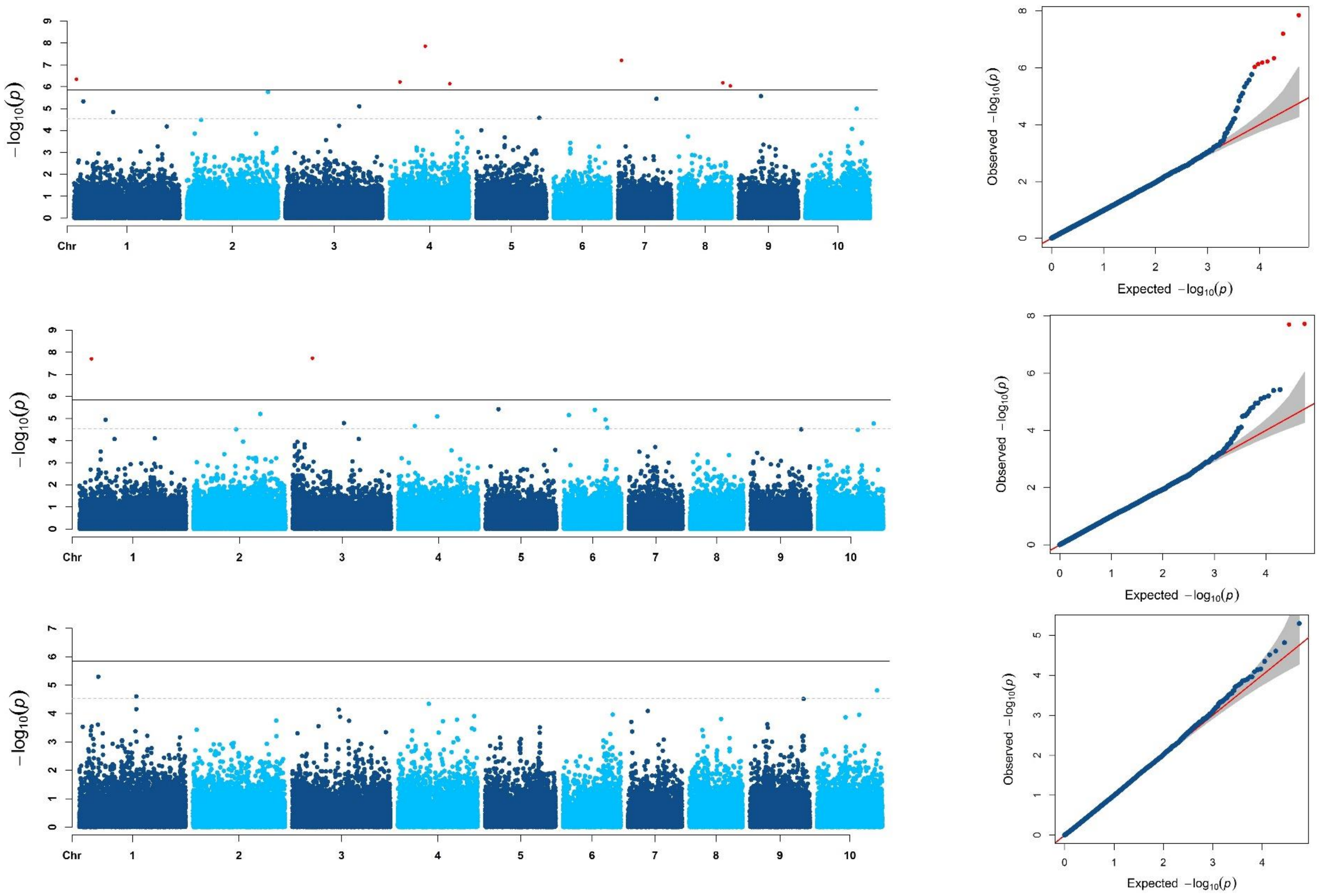

### Figure S9 NAM pcs

PC1

PC2

PC3

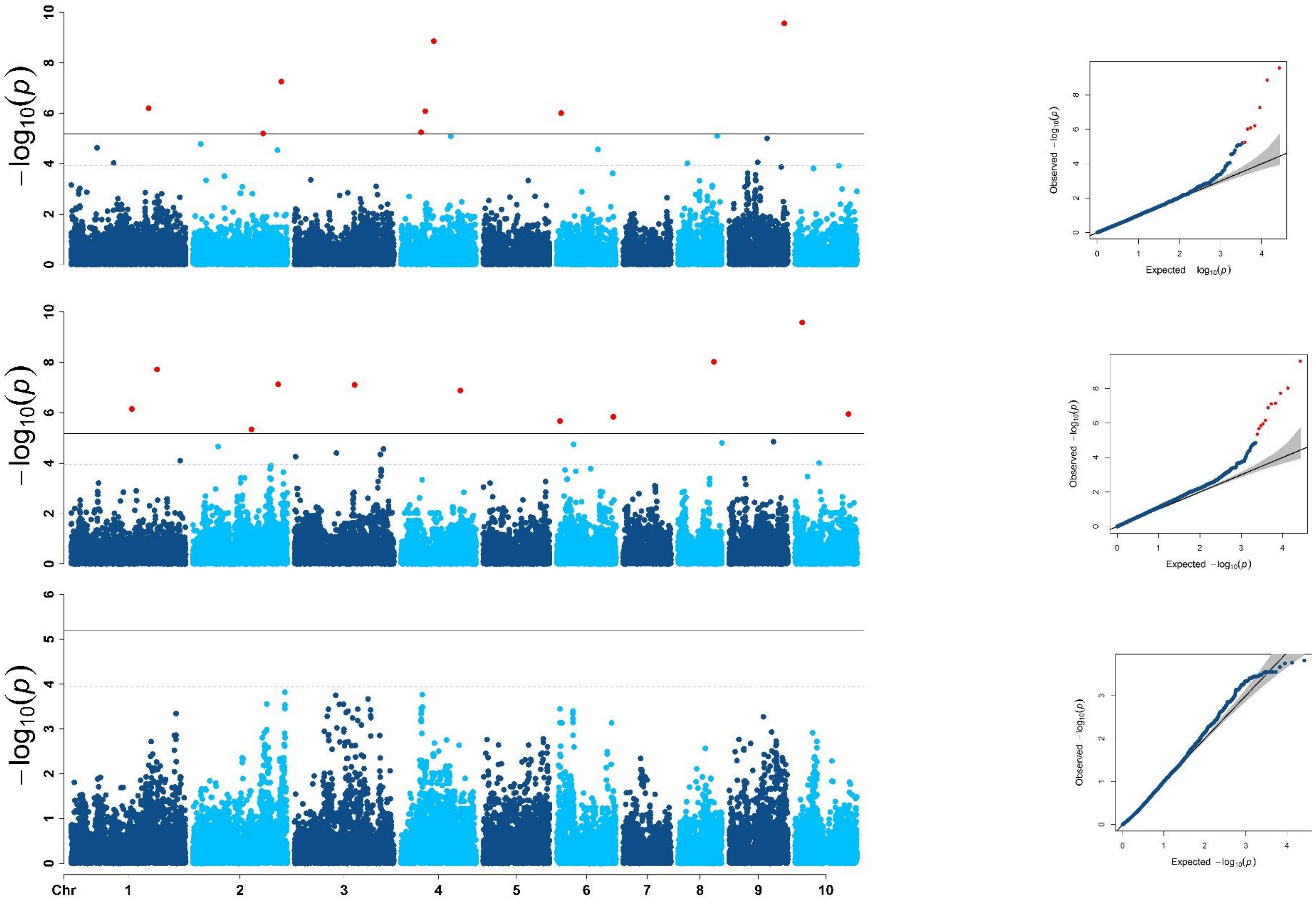

### Figure S10

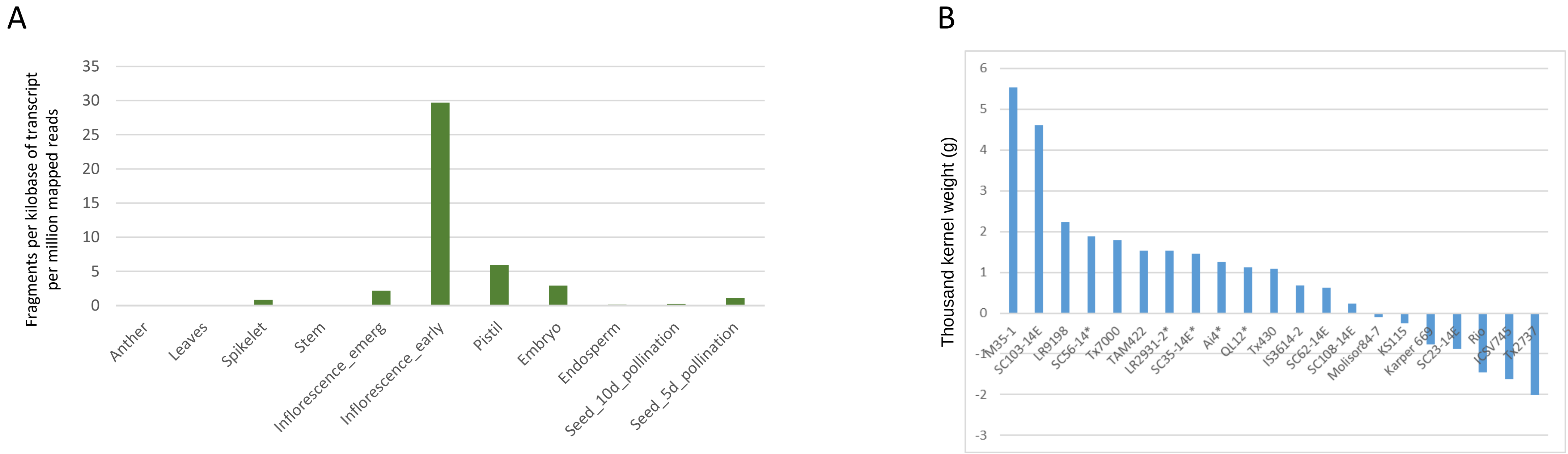
A
Fragments per kilobase of transcript per million mapped reads
35
30
25
20
15
10
5
0
Anther
Leaves
Spikelet
Stem
Inflorescence_emerg
Inflorescence_early
Pistil
Embryo
Endosperm
Seed_10d_pollination
Seed_5d_pollination
B
Thousand kernel weight (g)
6
5
4
3
2
1
0
-1
-2
-3
M35-1
SC103-14E
LR9198
SC56-14*
Tx7000
TAM422
LR2931-2*
SC35-14E*
Ai4*
QL12*
Tx430
IS3614-2
SC62-14E
SC108-14E
Molisor84-7
KS115
Karper 669
SC23-14E
Rio
ICSV745
Tx2737
